# Sensitivity optimization of a rhodopsin-based fluorescent voltage indicator

**DOI:** 10.1101/2021.11.09.467909

**Authors:** Ahmed S Abdelfattah, Jihong Zheng, Daniel Reep, Getahun Tsegaye, Arthur Tsang, Benjamin J Arthur, Monika Rehorova, Carl VL Olson, Yi-Chieh Huang, Yichun Shuai, Minoru Koyama, Maria V Moya, Timothy D Weber, Andrew L Lemire, Christopher A Baker, Natalie Falco, Qinsi Zheng, Jonathan B Grimm, Mighten C Yip, Deepika Walpita, Craig R Forest, Martin Chase, Luke Campagnola, Gabe Murphy, Allan M Wong, Jerome Mertz, Michael N Economo, Glenn Turner, Bei-Jung Lin, Tsai-Wen Chen, Ondrej Novak, Luke D Lavis, Karel Svoboda, Wyatt Korff, Eric R Schreiter, Jeremy P Hasseman, Ilya Kolb

**Author notes:** corresponding authors: ASA, ERS, JPH, IK. contributed equally to this work (Ahmed S Abdelfattah, Jihong Zheng). co-supervised this work (Eric R Schreiter, Jeremy P Hasseman, Ilya Kolb).

## Abstract

The ability to optically image cellular transmembrane voltage at millisecond-timescale resolution can offer unprecedented insight into the function of living brains in behaving animals. The chemigenetic voltage indicator Voltron is bright and photostable, making it a favorable choice for long *in vivo* imaging of neuronal populations at cellular resolution. Improving the voltage sensitivity of Voltron would allow better detection of spiking and subthreshold voltage signals. We performed site saturation mutagenesis at 40 positions in Voltron and screened for increased ΔF/F_0_ in response to action potentials (APs) in neurons. Using a fully automated patch-clamp system, we discovered a Voltron variant (Voltron.A122D) that increased the sensitivity to a single AP by 65% compared to Voltron. This variant (named Voltron2) also exhibited approximately 3-fold higher sensitivity in response to sub-threshold membrane potential changes. Voltron2 retained the sub-millisecond kinetics and photostability of its predecessor, with lower baseline fluorescence. Introducing the same A122D substitution to other Ace2 opsin-based voltage sensors similarly increased their sensitivity. We show that Voltron2 enables improved sensitivity voltage imaging in mice, zebrafish and fruit flies. Overall, we have discovered a generalizable mutation that significantly increases the sensitivity of Ace2 rhodopsin-based sensors, improving their voltage reporting capability.

## Introduction

Genetically encoded voltage indicators (GEVIs) have served as an enabling technology for visualizing neuronal activity at unprecedented spatiotemporal resolution(Hochbaum et al. 2014; Lin and Schnitzer 2016; Xu et al. 2017). Nevertheless, optical imaging of voltage using GEVIs presents many challenges for the design of these proteins. An ideal voltage sensor must concurrently fulfill many requirements, including but not limited to: (1) high sensitivity to membrane potential changes of a neuron, (2) fluorescence changes that are fast enough to follow and accurately report APs and (3) high degree of localization to neuron outer membranes. Further requirements may be desirable depending on application, such as sensitivity to sub-threshold membrane potential changes, photostability, and compatibility with two-photon excitation.

One approach to engineering GEVIs involves exploiting the native voltage sensitivity of microbial rhodopsins. The opsin Archaerhodopsin 3 (Arch) was first successfully used to optically record APs in neuronal culture (Kralj et al. 2012); however, it was found to be too dim at physiologically tolerable imaging powers for *in vivo* applications. Subsequent protein engineering efforts of Arch yielded improvements in brightness as well as sensitivity, kinetics, and reduced photocurrents (Chien et al. 2021; Flytzanis et al. 2014; Gong et al. 2013; Hochbaum et al. 2014; McIsaac et al. 2014; Piatkevich et al. 2018). An alternative strategy to develop bright rhodopsin-based GEVIs is to create a Förster resonance energy transfer (FRET) pair between a bright fluorescent protein (FP) and the rhodopsin protein (Gong et al. 2015; Zou et al. 2014). In this strategy, the bright FP is the reporter fluorophore, and the rhodopsin is used as the voltage sensitive domain. This strategy was successfully implemented to develop Ace2N-mNeon, a bright fast GEVI that was able to report single APs *in vivo* (Gong et al. 2015).

The Ace2N-mNeon member of the rhodopsin family of GEVIs has been used as a scaffold to create GEVIs with other favorable characteristics. A red GEVI called VARNAM consisting of Ace2N fused to a red FP mRuby3 displayed high sensitivity, good *in vivo* performance, and a spectral shift that made it compatible with blue-shifted optogenetic probes (Kannan et al. 2018). Our group has previously replaced the FP in Ace2N-mNeon with a HaloTag protein (Los et al. 2008) covalently bound to a small-molecule fluorophore (JaneliaFluor or JF (Grimm et al. 2015, 2017)) to create a chemigenetic sensor called Voltron (Abdelfattah et al. 2019). The introduction of three point mutations to the rhodopsin domain of Voltron led to Positron, a positive-going GEVI with sensitivity and kinetics comparable to the original Voltron (Abdelfattah et al. 2020).

Encouraged by the ability of point mutations in the rhodopsin domain to alter function, we performed a large-scale screen of point mutations to find improved versions of Voltron. We discovered that the introduction of an A122D mutation increased the sensitivity of Voltron, particularly in the sub-threshold range, without compromising kinetics, membrane trafficking or photobleaching. Thus Voltron.A122D was named Voltron2 as a next-generation version of the sensor. Consistent with the observation in culture, *in vivo* imaging in flies, zebrafish and mice revealed an increased signal-to-noise ratio (SNR) of Voltron2 compared to Voltron.

## Methods

### Reagent availability

The following plasmids used in this study are available on Addgene:

- pAAV-syn-FLEX-Ace2N-4AA-mNeon-ST A122D WPRE (#172908)
- pGP-pcDNA3.1 Puro-CAG-Voltron2 (#172909)
- pGP-CAG-Ace2N-4AA-mNeon A122D-WPRE-bGH-polyA (#172911)
- pGP-CAG-Ace2N-4AA-mNeon-ST A122D-WPRE-bGH-polyA (#172912)

The JF_549_-HaloTag ligand is available from Promega; all other dyes can be requested at dyes.janelia.org.

### Single-site directed mutagenesis

The cloning vector pcDNA3.1/Puro-CAG-Ace2N_HaloTag expression vector (Invitrogen) was modified by moving the KpnI site from outside of the insert to the junction between Ace2N domain and the Halo-tag. The subsequent vector was digested by NheI/KpnI cleaving out the Ace2N domain. End PCR primers were designed 30bp upstream of NheI site (5’-GCTCACAAATACCACT-3’) and 38 bp downstream of new KpnI site (5’-CCAGGACTTCCACATAA-3’). Overlapping internal primers were designed for each of 40 targeted amino acid residues in the Ace2N domain. One primer of the pair contained the degenerate codon NNS and the other primer a 27-30bp complementary overhang. When paired with the end primers two amplicons were created (Phusion High-Fidelity DNA Polymerase; NEB) that overlap with each other and the digested vector ends. Each set of overlapping paired amplicons (37.5 fmol each) were assembled with the digested pcDNA3.1/Puro-CAG backbone (25 fmol) using an isothermal assembly reaction (Gibson et al. 2009). Each 20 uL reaction mix consisted of 5X isothermal assembly buffer (25% PEG-8000, 500 mM Tris-HCl pH 7.5, 50mM MgCl2, 50mM DTT, 1mM each dNTP and 5mM NAD), T5 exonuclease (0.08 U, NEB), Taq DNA Ligase (80 U, NEB), Phusion HF DNA Polymerase (0.5 U, NEB). The reactions were incubated at 50oC for 30-60 minutes. Reactions were transformed into STABL2 chemically competent E. coli cells (ThermoFisher) and plated on LB/Amp agar plates and incubated at 37oC for 16-20 hours.

For each site library, 96 colonies were picked into 2.6 mL of 2x-YT media (2 × 1.3 mL in 2mL deep-well culture plates) and grown for 24 hours, 225 rpm @37C with Ampicillin (100mg/L). The cultures were pelleted at 3200 × g and frozen at −80C. For each plate plasmids were extracted using the E-Z 96 FastFilter Kit (Omega BioTek) and eluted into a half-area UV transparent 96 well plate (Corning Costar). Each of the plasmid plates was concentration normalized to 60ng/ul by reading the 260nm absorbance (Tecan Infinite M1000Pro) followed by custom dilution (Hamilton Nimbus). Variant plasmids were arrayed along with controls for high-throughput electroporation of neuronal cell culture (Hamilton STAR). Top performing variants from the subsequent neuronal culture screen were Sanger-sequenced to determine their mutation as well as the entire library being sequenced using a next-generation deep-sequencing approach (Supplementary Methods).

### Combinatorial mutagenesis

Top-performing single-site mutations (Y63L, N69E, V74E/W, R78H, N81S, L89A/C/G/T, A122D/H, V196P) were recombined to test all possible combinations (1423). All combinations could be recapitulated using two overlapping amplicons covering the Ace2N domain. Some mutations (Y63L, A122D/H, V196P) were introduced as part of the PCR template and others (N69E, V74E/W, R78H, N81S, L89A/C/G/T) by PCR primer. For the N-term amplicon (305bp) twenty-four reverse primers were designed based on the wild-type anti-sense sequence (5’- AGTGGTGTGGTCAGCACCCAGTTAATATATCTTGCGTAGACCACCTGCCTTTCACCATTCATTGTCAGGTCC-3’) and included every combination of N69E, V74E/W, R78H, N81S. Forty-eight unique N-term amplicons were created by combining these twenty-four reverse mutagenic primers, the upstream end primer (5’- GCTCACAAATACCACT-3’) and templates with and without Y63L. For the C-term amplicon (493bp) 10 forward primers were designed based on the wild-type sense sequence (5’- ATATTAACTGGGTGCTGACCACACCACTGCTCCTGCTCGATCTCATCGTCATGACCAAGATGGGCGGAGTGA -3’) and included every combination of N81S, L89A/C/G/T. Sixty unique C-term amplicons were created by combining 10 forward mutagenic primers, the downstream end primer (5’- CCAGGACTTCCACATAA-3’) and templates each containing a combination of A122D/H and V196P. The N-term and C-term amplicon libraries overlapped by 28 bp (5’-ATATTAACTGGGTGCTGACCACACCACT-3’) and included the N81 site in both. The PCR products were gel extracted, quantified and normalized to 18.75 fmol/ul. The NheI/KpnI digested pcDNA3.1/Puro-CAG backbone was normalized to 12.5 fmol/ul. Using a liquid-handling robot (Hamilton STAR) the N-term and C-term amplicons sets were pairwise combined (2uL each) along with the NheI/KpnI digested pcDNA3.1/Puro-CAG vector (2uL) to create 1423 unique isothermal assembly reactions in 96 well thermocyler plates. The plates were reacted and transformed as above in 96 well plates. Approx. 35uL of each transformant was robotically dispensed into the corresponding wells of two 48 well Q-trays (Genetix) containing LB/Amp agar. Q-trays were incubated for 16-20 hours at 37°C and two colonies were picked from each well and separately cultured, pelleted and frozen in 96 well deep well plates. Plasmids were extracted from the 96 well pellets and concentration normalized as above. Once verified by Sanger sequencing the combinatorial variants were arrayed for electroporation of neuronal cell culture and the subsequent field stimulation assay.

### Neuronal cell culture

Experiments were conducted in accordance with guidelines for animal research approved by the Janelia Research Campus Institutional Animal Care and Use Committee. Neonatal rat pups (Charles River Laboratory) were euthanized and neocortices (for field stimulation experiments) or hippocampi (for patch-clamp experiments), were isolated. Tissue was dissociated using papain (Worthington) in 10 mM HEPES pH 7.4 in Hanks’ Balanced Salt Solution for 30 min at 37°C. Suspensions were triturated with a Pasteur pipette and passed through a 40-μm strainer. Cells were transfected by combining 5×10^5^ viable cells with 400 ng plasmid DNA and nucleofection solution in a 25-μL electroporation cuvette (Lonza). Cells were electroporated according to the manufacturer’s protocol.

For the field stimulation screen, neurons were plated onto poly-D-lysine (PDL) coated, 96-well, glass bottom (#1.5 cover glass) plates (MatTek) at ~1×10^5^ cells per well in 100 μL of a 4:1 mixture of NbActiv4 (BrainBits) and plating medium (28 mM glucose, 2.4 mM NaHCO3, 100 μg/mL transferrin, 25 μg/mL insulin, 2 mM L-glutamine, 100 U/mL penicillin, 10 μg/mL streptomycin, 10% FBS in MEM). The next day, 190 μL of NbActiv4 medium was added to each well. Plates were incubated at 37°C and 5% CO2, to be imaged after 12-15 days in culture. Typically, 8 wells of a 96-well plate were electroporated with Voltron (as a control) and the remaining wells were electroporated with constructs of interest (4 wells per construct). The first and last columns of the plate were not used.

For patch-clamp, ~2×10^5^ cells were plated onto PDL-coated, 35-mm glass bottom plates (Mattek, #0 cover glass) in 120 μL of a 1:1 mixture of NbActiv4 and plating medium in the center of the plate. The next day, 2 mL of NbActiv4 medium was added to each plate. Plates were incubated for 7-13 days prior to beginning experiments.

### Field stimulation assay in neuronal culture

To prepare the neurons for field stimulation, they were first incubated for 1 hour in NbActiv4 media supplemented with 2 nM JF_525_-HaloTag at 37°C. They were then rinsed three times with imaging buffer containing (in mM) 140 NaCl, 0.2 KCl, 10 HEPES, 30 glucose (pH 7.3-7.4) and left in a solution containing imaging buffer with added receptor blockers (10 μM CNQX, 10 μM (R)-CPP, 10 μM gabazine, 1 mM (S)-MCPG, Tocris) to reduce spontaneous activity (Wardill et al. 2013).

The field stimulation assay for GEVIs was adapted from our existing screening pipeline (Dana et al. 2016, 2019). Fluorescence was excited with a white LED (Cairn Research) through a custom filter cube (Excitation: 512/25 nm, Emission: 555/20 nm, dichroic: 525 nm, Chroma) and imaged using a 40X/0.6 NA objective (Olympus) with an EMCCD camera (Ixon Ultra DU897, Andor). To enable high-speed imaging, an Optomask (Cairn Research) was used to mask out camera pixels outside a 256×256 center square. Reference images of each field of view (FOV) were taken at full sensor frame, 100 ms exposure. For high-speed imaging during stimulation, we applied 8x binning, 0.01 ms exposure, and 25 EM gain for a resulting frame rate of 1,497 Hz.

For each well in the 96-well plate, either 9 FOVs surrounding the center of the well were chosen, or a machine vision function utilizing ilastik (Berg et al. 2019) was used to automatically focus on cell somata. For each FOV, first, a reference image was acquired, and then, neuronal APs were evoked by field stimulation (8 pulses, 40 V, 1 ms, 8.3 Hz; S-48, Grass Instruments) concurrently with high-speed imaging. The camera ‘fire’ signal and the stimulator sense line were used to determine the frame at which the stimulation occurred.

To correct for photobleaching, a single exponential with three free parameters was fit to the time series for each pixel. Frames succeeding each electrical stimulus (during which the response nominally occurred) were excluded from the fit. The value of the fitted bleach function at the first frame was taken as the baseline fluorescence for that pixel. Background fluorescence was computed as the 1^st^ percentile of the baseline fluorescence across all pixels.

Responses to the eight electrical pulses within each recording were averaged using the timings derived from the camera and electrode triggers. For each pixel, a Mann-Whitney U test was performed between the frames preceding the average response (20 ms) and 10, 20, and 40 ms of frames succeeding it. Pixels with a p value < 0.001 for any of these three tests were considered responsive and averaged together to contribute to the ΔF/F_0_ trace. Traces were fit with the product of a rising and decaying exponential to capture both the on and off kinetics. The fit was used to calculate the characteristics of the variant such as maximum ΔF/F_0_ and kinetics (10-90% rise and decay times).

Pixel statistics were pooled across all the wells in each plate that contained the construct of interest. Wells with fewer than four responsive pixels were considered to be unresponsive and discarded from analysis.

For every plate in the field stimulation assay, a percent detectable improvement (PDI) statistic was calculated to answer the question: “Given the variability of Voltron control wells in the plate, what is the minimum improvement in ΔF/F_0_ that can be reliably detected?”. That is, a PDI of 20% for a plate indicates that a ≥ 20% improvement in ΔF/F_0_ over Voltron can be considered statistically meaningful. Large PDI values are undesirable because they indicate high variability in the control responses. PDI is calculated as follows: 100*(mean(x)-quantile(x,0.01))/mean(x), where x is the ΔF/F_0_ of Voltron control well pixels, sampled 10,000 times with replacement. The PDI was one of the parameters used to screen out poorly responding variants.

Normalization to in-plate Voltron controls was useful to reduce the effects of within-plate and within-week variability. Pixels from each variant were pooled across wells. For each variant, the median was taken from this pool and divided by the median from the control pool to perform the normalization. Significance values for each variant were determined using a Mann-Whitney U test between the pools.

### Automated whole-cell electrophysiology

Cultured neurons were patch-clamped at 7-13 DIV at room temperature (23°C). On the day of the experiment, cell culture medium was first rinsed with imaging buffer consisting of (in mM): 145 NaCl, 2.5 KCl, 10 D-Glucose, 10 HEPES, 2 CaCl2, 1 MgCl2 (pH 7.3, adjusted to 310 mOsm with sucrose). The cells were then incubated with 100 nM JF_525_ dye for 10 minutes (for Voltron mutant screening only), rinsed twice, and kept in imaging buffer. For voltage clamp recordings, 1 μM TTX was added to the bath to suppress the generation of APs. Micropipettes were pulled on a horizontal puller (P-97, Sutter Instruments) to a tip resistance of 3 to 6 MΩ. For voltage clamp experiments, pipettes were filled with cesium-based internal solution containing (in mM): 115 CsMeSO4, 15 CsCl, 3.5 Mg-ATP, 5 NaF, 10 EGTA, 10 HEPES, 3 QX-314 (pH 7.3-7.4, 280-290 mOsm). For current clamp experiments, pipettes were filled with 130 KMeSO4,10 HEPES, 5 NaCl, 1 MgCl2, 1 Mg-ATP, 0.4 Na-GTP, 14 Tris-phosphocreatine (pH 7.3-7.4, 280-290 mOsm).

To perform automated patch-clamp screening of the top-performing hits from the field stimulation screen, we used a custom-built Automated uM Workstation, manufactured by Sensapex (Oulu, Finland), based on the PatcherBot (Kolb et al. 2019). The system is built around an AxioObserver 7 inverted microscope (Zeiss), outfitted with a computer-controlled stage, micromanipulators, and pipette pressure controllers. Pipettes were automatically cleaned between every patch-clamp attempt with Tergazyme and reused, enabling higher throughput than possible with manual patch-clamp (Kolb et al. 2016, 2019). Electrophysiology recordings were performed with a Multiclamp 700B amplifier (Molecular Devices), and digitized with a multifunction data acquisition board (National Instruments PCIe-6259). Neurons were imaged using a 40×/1.3 NA oil immersion objective (Zeiss), illuminated with high-power LEDs (Spectra-X light engine, Lumencor) and imaged with a digital sCMOS camera (Hamamatsu Orca Flash 4.0). To image Voltron_525_, we used a filter cube containing 510/25 nm excitation filter, 545/40 emission filter, 525 nm dichroic (Chroma), with a measured power of 14.7 mW/mm^2^ in the imaging plane. To image Ace2N-mNeon, the filter cube contained a 470/24 nm excitation filter, 525/40 nm emission filter, 506 nm dichroic with a measured power of 18.1 mW/mm^2^ in the imaging plane. To image VARNAM, the filter cube contained 575/25 nm excitation filter, 610LP emission filter, 594 nm dichroic, with a measured power of 32.8 mW/mm^2^.

The uM Workstation was controlled by the Python platform Acq4 (Campagnola et al. 2014), modified to perform fully automated electrophysiology (https://www.acq4.org). To generate fluorescence/voltage curves, the membrane potential was stepped from +50 to −110 mV in 20 mV increments from a resting potential of −70 mV (0.5 s baseline, 1 s step). For current clamp recordings, a short current pulse was injected (2 nA, 2 ms) to evoke APs.

Stimulus timing, baseline fluorescence calculation, background subtraction, and photobleaching correction was performed the same way as for the field stimulation assay. To identify responsive pixels, a Mann-Whitney U test was performed between the baseline and voltage step segments of the recording. The P value criterion to identify responsive pixels was empirically set to 1e-10.

The onset of each step was fit with the product of a rising and decaying exponential to capture the transient response (if any), summed with a single rising exponential to capture the steady-state response. The decay response was fit with a single exponential. Peak ΔF/F_0_ as well as onset and decay kinetics were calculated at each voltage step as was done for field stimulation.

### Imaging and whole-cell recording in brain slices

All animal work was performed according to Institutional Animal Care and Use Committee approved protocols. Stereotaxic injections were made into right visual cortex (3.8 mm posterior and 3.0 mm lateral from bregma) of ~4-week-old Sst-IRES-Cre driver mice under isofluorane anesthesia. Two injections of 200 nL each of AAV2/1-syn-Flex-Voltron_585_-ST and AAV2/1-syn-Flex-Voltron2_585_-ST and were targeted to 300 and 600 μm below the cortical surface.

Four weeks later, isoflurane anesthetized mice were transcardially perfused with ice-cold NMDG slicing solution containing (in mM): 98 HCl, 96 N-methyl-d-glucamine (NMDG), 2.5 KCl, 25 D-Glucose, 25 NaHCO3, 17.5 HEPES, 12 N-acetylcysteine, 10 MgSO4, 5 Na-L-Ascorbate, 3 Myo-inositol, 3 Na Pyruvate, 2 Thiourea, 1.25 NaH2PO4·H2O, 0.5 CaCl2, and 0.01 taurine. Acute 350 μm parasagittal slices containing primary visual cortex from the right hemisphere were prepared with a Compresstome (Precisionary Instruments) in ice-cold NMDG slicing solution at a slice angle of 17° relative to the sagittal plane. Slices were incubated for 10min in NMDG slicing solution at 34°C and then transferred to artificial CSF (aCSF; in mM): 94 NaCl, 25 D-Glucose, 25 NaHCO3, 14 HEPES, 12.3 N-acetylcysteine, 5 Na-L-Ascorbate, 3 Myo-inositol, 3 Na Pyruvate, 2.5 KCl, 2 CaCl2, 2 MgSO4, 2 Thiourea, 1.25 NaH2PO4 H20, 0.01 Taurine. All solutions were maintained under constant carbogen (95% O2; 5% CO2).

To complete fluorescent labeling of Voltron-expressing cells, 1 nM of JF_585_ was dissolved in 20μl dimethyl sulfoxide (DMSO) and 20 μL of 20% Pluronic F-127 (w/w in DMSO). The solubilized dye was then added to 20 mL of oxygenated aCSF and incubated with the acute brain slices for 1h at room temperature, after which the slices were removed to a holding chamber (BSK 12, Scientific Systems Design) containing 500 mL oxygenated aCSF without dye. Slices were kept in this latter solution for at least one hour at room temperature prior to any experiment.

Slices were visualized using oblique (Olympus; WI-OBCD) infrared illumination using 20× or 4× objectives (Olympus). Recording pipettes were pulled from filamented borosilicate glass (Sutter Instruments) to a tip resistance of 3–8 MΩ using a DMZ Zeitz-Puller (Zeitz). Electrophysiology, image collection and subsequent analysis were performed using Acq4. Signals were amplified using Multiclamp 700B amplifiers (Molecular Devices) and digitized at 50–200 kHz using ITC-1600 digitizers (Heka). Neurons were held in whole-cell patch clamp with an internal solution containing (in mM): 130 K-gluconate, 10 HEPES, 0.3 ethylene glycol-bis(β-aminoethyl ether)-N,N,N’,N’-tetraacetic acid (EGTA), 3 KCl, 0.23 Na2GTP, 6.35 Na2Phosphocreatine, 3.4 Mg-ATP, 13.4 biocytin, and 50 μM Cascade Blue dye.

Voltron_585_-associated fluorescence was examined using a 595 nm LED (Thorlabs) at 6.9 μW/mm^2^ power and 598/25 nm excitation and 650/54 nm emission filters (Semrock). Images were collected by sampling a 675μm × 137μm region of the slice with a digital sCMOS camera (Hamamatsu; Flash 4.0 V2) at 500 Hz and 4×4 pixel binning. Image analysis was performed by custom routines written in Python. For each camera frame, average fluorescence intensity over an elliptical region of interest (ROI) over neuropil adjacent to a cell was subtracted from an identically shaped region containing the cell itself. Synthetic post-synaptic potentials (synPSPs) of −15mV to +15mV in 5mV increments, repeated for a total of 10 trials per cell were injected in voltage clamp mode. Two adjacent 10 ms-long temporal windows prior to the onset of the current injection were designated as “noise” and “baseline” epochs, and a third 10 ms “signal” temporal window was centered over the fluorescence peak (from 4 ms to 14 ms after the onset of the change in membrane potential). The ΔF/F was calculated for each trial as the average change in fluorescence between the “signal” and “baseline” windows. To determine SNR, the least-squares regression line for the ΔF/F was used to determine the average change in signal per mV. The noise for this ratio was calculated by determining the standard deviation of the dF/F between the “noise” and “baseline” windows (when the membrane potential in the cell was held constant), and then dividing the signal per mV by that value. Thus, a value of 0.5 indicates that the fluorescence change associated with a 4 mV alteration in membrane potential is equal in magnitude to 2× the st.dev. of fluorescence values during a period in which the membrane potential is unchanged.

### Simultaneous voltage imaging and optogenetic stimulation in brain slices

Voltron2 and Channelrhodopsin2 were expressed throughout the motor cortex using injections of a mixture of (1) rAAVretro-hSyn-Cre-WPRE (2×10^9^ g.c.; Addgene #105553-AAVrg), (2) AAV1-Syn-FLEX-Voltron2585-WPRE (1×10^9^ g.c.), (3) AAV8-Syn-ChR2(H134R)-GFP (3×10^8^ g.c.; Addgene #58880-AAV8), and (4) 0.05% Trypan Blue in 1 μL of sterile PBS into the lateral ventricle of C57Bl/6N mice (Charles River) at postnatal day 1 (Kim et al. 2014). At least 14 days following virus injection, mice were transcardially perfused with 15 mL of chilled and carbogen-bubbled (95% O_2_/5% CO_2_) NMDG aCSF solution (in mM: 92 NMDG, 2.5 KCl, 1.25 NaH_2_PO_4_, 30 NaHCO_3_, 20 HEPES, 25 glucose, 2 thiourea, 5 Na-ascorbate, 3 Na-pyruvate, 0.5 CaCl_2_·4H_2_O and 10 MgSO_4_·7H_2_O, pH 7.3-7.4, 300-310 mOsm). Acute slices through motor cortex were made in chilled NMDG aCSF with constant bubbling (Ting et al. 2014). Following re-introduction of sodium in 37°C NMDG aCSF, slices were transferred to a holding chamber containing 25 nM JF_585_ dye in 5 mL bubbled, room temperature HEPES aCSF buffer (in mM: 92 NaCl, 2.5 KCl, 1.25 NaH_2_PO_4_, 30 NaHCO_3_, 20 HEPES, 25 glucose, 2 thiourea, 5 Na-ascorbate, 3 Na-pyruvate, 2 CaCl_2_·4H_2_O and 2 MgSO_4_·7H_2_O, pH 7.3-7.4, 300-310 mOSm). Slices were incubated in dye solution for 1 hour, and moved to fresh HEPES aCSF for 1 hour to wash out excess dye. Experiments were performed at room temperature in HEPES aCSF solution. Whole-cell recordings were made using filamented glass pipettes (Sutter #BF150-86-10) pulled to 3-8 MOhm resistance (Sutter P-1000 Micropipette Puller), and intracellular recording buffer containing (in mM) 145 K-Gluconate, 10 HEPES, 1 EGTA, 2 Mg-ATP, 0.3 Na_2_-GTP, and 2 MgCl_2_ (pH 7.3, 290-300 mOsm). A patch-clamp headstage (Molecular Devices #1-CV-7B) mounted on a motorized 4-axis Siskiyou MX7600 manipulator, and Axon Instruments MultiClamp 700b amplifier were used for all recordings.

Imaging was performed using a custom-built confocal microscope at a frame rate of 458 Hz using a 16X/0.8 NA water-immersion objective lens (Nikon CFI75 LWD 16X W). High frame rates were achieved using a system similar to that described previously (Badon et al. 2019) but with a 128-facet polygonal scanner (Cambridge Technology SA34) substituted for the x-axis scanner. Voltron2_585_ was excited with a 561 nm laser diode (Vortran Stradus). The time-averaged irradiance at the sample was 33 mW/mm^2^ and fluorescence was epi-collected with a dichroic mirror and emission filter (Chroma T570lpxr and ET570lp), and detected with a silicon photomultiplier (Hamamatsu S14420-1550MG, V_BIAS_ = 50 V) and amplified on-board with a custom circuit (https://github.com/tweber225/simple-sipm). A blue LED (Thorlabs M470L4) was used to provide full-field ChR2 stimulation. The LED was filtered and coupled into the confocal beam path with an excitation filter and dichroic mirror (Thorlabs MF475-35 and DMLP505R). Additionally, the LED was attenuated such that the desired irradiance levels (10-50 μW/mm^2^) were within the analog control range of the LED driver (Thorlabs LEDD1B). Image acquisition and stimulus timing were managed with ScanImage (Pologruto et al. 2003) and WaveSurfer (https://wavesurfer.janelia.org/).

### Lattice lightsheet imaging in zebrafish

*In vivo* light sheet microscopy of zebrafish was performed as previously described (Liu et al. 2018). Briefly, zebrafish transgenic lines expressing soma-tagged Voltron (Tg[vglut2a:Gal4; UAS:Voltron_552_-ST]) and Voltron2 (Tg[vglut2a:Gal4;UAS:Voltron2_552_-ST]) were generated. At three days post-fertilization (dpf), fish were incubated in a water solution containing 3 μM JF_552_ for 2 h. The fish at 4 to 6 dpf were then paralyzed by a-bungarotoxin (1 mg/mL) and mounted in low melting point agarose for imaging. The custom microscope used for imaging was described previously (Liu et al. 2018). Here it was used without the adaptive optics (AO) subsystem since optical aberration was negligible in the structure we imaged. A 740 nm thick light sheet was created from a 560 nm laser source using a multi-Bessel lattice with an outer and inner NA of 0.38 and 0.36, respectively, for a measured power of 100 μW at the back pupil of the excitation objective. Single-plane imaging was performed at an effective 108 × 108 nm XY resolution, with an FOV of 256×512 pixels, at a framerate of 400 Hz. Approximately 4-10 Voltron-expressing neurons were present in each field of view.

Fluorescent signal was recorded for 5 min. For analysis, the automated voltage imaging analysis package Volpy was used (Cai et al. 2020). To perform an unbiased comparison of Voltron_552_ and Voltron2_552_ populations, every spiking cell detected by Volpy was included in the analyzed dataset, irrespective of AP amplitude. ΔF/F_0_ and SNR for each cell were calculated by Volpy. For SNR calculations, the noise was defined as the standard deviation of the residual after subtracting spike and subthreshold components, as detected by Volpy.

### Voltron imaging in adult flies

Experiments were performed as described previously (Abdelfattah et al. 2019). Briefly, crosses of Voltron_552_ (*UAS-IVS-syn21-Ace2NHalo-p10* Su(Hw)attP8) or Voltron2_552_ (*UAS-IVS-syn21-Ace2N(A122D)Halo-p10* Su(Hw)attP8) reporters with split Gal4 drivers were raised on standard cornmeal food supplemented with all-trans-retinal (0.2 mM before eclosion and then 0.4 mM). 2- to 10-day old female progeny were collected for experiments. To prepare the fly for imaging, a small hole was dissected in the head capsule, and air sacs and fat tissue were removed but we did not intentionally remove the perineural sheath. The exposed brain was then bathed in a drop (~200 μL) of dye-containing saline (1 μM for JF_552_-Halotag ligand) for 1 hr. Saline contains (in mM): NaCl, 103; KCl, 3; CaCl_2_, 1.5; MgCl_2_, 4; NaHCO_3_, 26; N-tris(hydroxymethyl)methyl-2-aminoethanesulfonic acid, 5; NaH_2_PO_4_, 1; trehalose, 10; glucose, 10 (pH 7.3 when bubbled with 95% O_2_ and 5% CO_2_, 275 mOsm). The dye was then washed-out by rinsing three times with ~10 mL of fresh saline each time over a 1-hr period. Imaging was performed on a widefield fluorescence microscope (SOM, Sutter Instruments) equipped with a 60x, NA 1.0, water-immersion objective (LUMPlanFl/IR; Olympus) and an sCMOS camera (Orca Flash 4.0 V3, Hamamatsu). Images were acquired at 800 frames per second with 4×4 binning through the Hamamatsu imaging software (HCImage Live). For JF_552_, illumination was provided by a 561-nm LED (SA-561-1PLUS, Sutter) with an excitation filter (FF01-549/12-25, Semrock); intensity at the sample plane was 2-11 mW/mm^2^ for typical recordings. Emission was separated from excitation light using a dichroic mirror (FF562-Di03-25×36, Semrock) and an emission filter (FF01-590/36-25, Semrock). We found that JF_552_ allows for longer-duration imaging compared with JF_549_ and JF_525_, which we used previously (Abdelfattah et al. 2019). At the aforementioned illumination levels, spiking activity was detectable for over 20 min in PPL1-γ1pedc and over 10 min in MBON-γ1pedc>α/β.

For MBON-γ1pedc>α/β, both left and right hemispheres were sampled, while for PPL1-γ1pedc, whose axons project bilaterally, only one hemisphere was imaged. Each experiment at one illumination level consists of a recording of 15 s. Data were analyzed with custom-written scripts in MATLAB (Mathworks). Regions of interest (ROIs) corresponding to the γ1 region were manually selected, and the mean pixel intensity within the ROI was calculated. The raw fluorescence trace was de-trended by median filtering with a 50 ms time window. F_0_ was calculated from the filtered trace as the mean over the first 1 s of imaging session. Spike sorting and SNR quantification were performed on the de-trended trace. Spikes were automatically detected by finding local minima and verified by visual inspection. SNR was quantified as peak amplitude over the standard deviation of the trace excluding a 50 ms time window around any spikes.

### Imaging Parvalbumin (PV) neurons in mouse hippocampus

Hippocampal PV neuron imaging was performed using adult PV-Cre mice (JAX 008069). The imaging window was implanted using procedures similar to those previously described (Dombeck et al. 2010). In short, a circular craniotomy (3 mm diameter) was made centered at 2.0 mm caudal and 2.0 mm lateral to bregma. The surface of CA1 was exposed by gently removing the overlying cortex with aspiration. AAV2/1-syn-FLEX-Voltron_552_-ST (Voltron_552_; 4 mice) and either AAV2/1-syn-FLEX-Voltron2_552_-ST (2 mice) or AAV2/1-CAG-FLEX-Voltron2_552_-ST (Voltron2_552_, 3 mice) virus was diluted to 1.4×10^13^, 4.1×10^13^ and 8.26×10^11^ GC/mL, respectively. Diluted viruses were injected at four locations (separated by 700 μm, 50 nL per location) at a depth of 200 μm from CA1 surface (injection rate, 1 nL/s). The imaging window (constructed by gluing a 3 mm diameter cover glass to a stainless steel cannula of 3 mm diameter and 1.5 mm height) was placed onto the hippocampus and glued to the skull using super-bond C&B (Sun Medical). A titanium head bar was glued to the skull for head fixation during imaging.

Imaging experiments started 3 weeks after surgery. 100 nM of JF_552_ were dissolved in 20μL of DMSO (Sigma) and diluted in 20 μL Pluronic™ F-127 (20% w/v in DMSO, P3000MP, Invitrogen) and in 80 μL PBS. The dye solution was delivered using retro-orbital injection (Yardeni et al. 2011) with a 30 gauge needle. Three hours after dye injection, animals were placed under the microscope and labeled PV+ cells (47 – 137 μm deep) were illuminated using a 532 nm laser (Opus 532, Laser Quantum) through an excitation filter (FF02-520-28, Semrock). Fluorescence was collected using a 16X/0.8 NA objective (Nikon), separated from excitation light using a dichroic mirror (540lpxr, Chroma) and an emission filter (FF01-596/83, Semrock), and imaged onto a CMOS camera (DaVinci-1K, RedShirt) using a 50mm camera lens (Nikkor 50mm f1.2, Nikon) as the tube lens. For patterned illumination, the laser beam was expanded using a pair of lenses (C280TMD-A and AC254-150-A, Thorlabs) and directed to a digital micromirror device or DMD (V7000, ViALUX). The DMD was imaged to the sample using an 80 mm lens (AC254-080-A, Thorlabs) and the microscope objective. A reference image of labeled cells was first acquired using widefield illumination. Bright and in-focus neurons were selected manually and their coordinates were used to generate an illumination mask consisting of 64 μm diameter discs centered on each selected cell. The illumination intensity was ~70-140 mW/mm^2^ (i.e. ~0.22 – 0.45 mW per cell) at the sample plane. Images (190 × 160 pixels, corresponding to an area of 1.4 × 1.2 mm) were collected at 2000 Hz using Turbo-SM64 software (Sci-Measure) for three minutes (360,000 images).

Brain motion was corrected using rigid registration. The fluorescence F(t) of each cell was measured by averaging pixel values within a 10-pixel region covering the cell body. To correct for bleaching and other slow fluctuations, a baseline fluorescence trace F_0_ (t) was computed from F(t) by a moving average with 1s windows. Since Voltron fluorescence decreases with membrane depolarization, we define ΔF/F_0_ (t)=(-(F(t)-F_0_(t)))/(F_0_(t)) as an estimate of cells’ membrane potential. To detect APs, a high pass filtered version of ΔF/F_0_, (ΔF/F_0_)_hp_, was computed by subtracting a median-filtered (5 ms window) ΔF/F_0_. Positive peaks of the (ΔF/F_0_)_hp_ trace were detected and considered as candidate spike locations (with t_k_ and p_k_ being the locations and the amplitudes, respectively, of the k^th^ candidate peak). To choose a threshold, the distribution of p_k_, P(x), was estimated by kernel density method (‘ksdensity’ function in MATLAB). The same procedure was applied to the inverted (ΔF/F_0_)_hp_ trace to detect ‘noise’ peaks, and the amplitudes of those peaks were used to construct a noise distribution, P_noise_(x). The distribution of spike amplitudes was estimated as S(x)=P(x)-P_noise_(x), and a threshold value *th1* was chosen at the location where S(*th1*)=P_noise_(*th1*) in order to minimize the sum of type I and type II error. This approach works well in cells with good signal to noise ratio (SNR), but in low SNR cells it often leads to substantial false positive detections. We estimated the number of false positive detections (*nFP*), at any given threshold value, by counting the number supra-threshold ‘noise’ peaks in the inverted (ΔF/F_0_)_hp_ trace. If *nFP* at *th1* exceeds 18 over the 180 s recording period (i.e. false positive rate > 0.1Hz), the threshold was replaced by a higher value, *th2*, that allowed a maximum of 18 false positive detections. Candidate peaks larger than the threshold were used for an initial estimate of spike times, and segments of the (ΔF/F_0_)_hp_ trace around these peaks were averaged to generate an initial estimate of the AP waveform, AP(t). Since AP waveforms exhibit finite rise and decay times, the occurrence of a spike interferes with the detection of spikes within its immediate neighborhood. To correct for this effect, if a candidate peak p_i_ was surrounded by a larger peak pj within ± 2 ms, its amplitude was corrected by assuming that a spike occurred at t_j_ and by subtracting the contribution of that spike, i.e. p_i,corrected_=p_i_-AP(t_i_-t_j_). This procedure was used to correct the amplitudes of all candidate peaks. Finally, a candidate peak was detected as an AP if its corrected amplitude exceeded the above mentioned threshold.

To quantify the recording quality and the fidelity of spike detection, we first estimated the spike amplitude A by averaging the amplitudes of all detected spikes. The noise of the recording σ was estimated as the standard deviation of the (ΔF/F_0_)_hp_ trace excluding regions 2 ms before and 4 ms after each detected spike. The signal to noise ratio was measured for each cell as SNR=A/σ. A cell was included into our analysis if (1) its SNR exceeded 5, (2) the number of detected spikes in the cell exceeded 90 (i.e. spike rate > 0.5 Hz), (3) less than 1% of detected spikes had an inter-spike-interval less than 2.0 ms, and (4) the half-width of the spike waveform was shorter than 0.85ms. To compare the density of labeled neurons, a z-stack of images was acquired at the end of the recording session and cell bodies in a 1280 ×1280×200 μm^3^ volume were counted manually.

### Imaging in mouse visual cortex

All procedures were approved by the Institutional Animal Care and Use Committee at the Second Faculty of Medicine, Charles University in Prague. The procedures were carried out in accordance with the relevant guidelines and regulations.

Layer 2/3 pyramidal neurons in the visual cortex of mice (C57BI/6NCrl; Charles River Laboratories) were sparsely labeled with the indicators – either Voltron_525_-ST or Voltron2_525_-ST. We prepared four mice per group. In an anesthetized mouse (isoflurane in pure oxygen; 4% for induction, 1–2% for maintenance), we first glued a ring-shaped titanium headbar to the skull of the animal using a gel-form cyanoacrylate and then fully closed the skin around the headbar. A craniotomy (4.5 mm in diameter) was drilled over the left parietal cortex, centered on −2.5 mm lateral, +0.5 mm anterior from lambda (visual cortex). Using beveled, pulled-glass capillaries (tip size <12 μm), we injected a mixture of two viruses: high-titer AAV carrying the cassette for conditional expression of the voltage indicator (AAV2/1-syn-FLEX-Voltron_525_-ST or AAV2/1-syn-FLEX-Voltron2_525_-ST; titer 10^12^ GC/mL) and low-titer AAV carrying the transcription permissive signal (AAV9-CamKIIa-Cre; titer 108 GC/mL). Six to eight 40 nL injections at a depth of 150 μm were performed in each mouse. The craniotomy was coversliped and the cranial window was secured using cyanoacrylate. A standard analgesia protocol (ketoprofen) followed. Approximately seven weeks after surgery, the animal was prepared for imaging. One day prior to imaging, JF_525_ dye was administered intravenously. To prepare the JF dye for injection, 100 nM of lyophilized JF_525_ was dissolved in 20 μL of DMSO, 20 μL Pluronic F-127 (20% w/v in DMSO), and 60–80 μL of PBS. Mice were briefly anesthetized and 100 μL of the dye solution was injected into the retro-orbital sinus of the right eye using a 30-gauge needle. We used the same design of wide-field fluorescence microscopy with structured illumination as described in (Abdelfattah et al. 2019). Illumination was delivered using a 525 nm LED (Mightex, LCS-0525-60-22) and shaped using a digital mirror device (Texas Instruments, LightCrafter). The microscope was equipped with a water immersion objective (20X, NA 1.0, Olympus XLUMPLFLN) and a CMOS camera (Hamamatsu Orca Flash v3). Excitation and emission were separated using a standard filter cube (Chroma 49014; excitation 530/30, dichroic 550, emission 575/40). The illumination was restricted to single neurons using DMD. The illuminated spot was 80 μm in diameter and the intensity was kept at 18.5 mW/mm2 in the sample plane. Small fields of view (40 μm × 40 μm) containing single neurons were typically captured. Native 2048×2048 resolution of the camera was binned by a factor of 4. During imaging, we recorded only from neurons that produced at least ~120 photons per frame and per pixel as this was expected to lead to approximately 1% standard deviation of the raw signal (quantum efficiency of the camera 82%, neuron covered with ~100 pixels, noise dominated by shot noise). Three-minute time series at 500 frames per second were captured for most of the recordings; one minute at 1000 frames per second was used only for comparison of AP-related fluorescence changes. Mice were imaged fully awake without any visual stimulation.

To process the recordings, we first removed the in-plane motion artifacts using the fast rigid registration algorithm NoRMCorre (Pnevmatikakis and Giovannucci 2017). Neurons (n = 107 expressing Voltron_525_-ST in 4 mice, n = 102 expressing Voltron2_525_-ST in 4 mice) were then segmented manually. The signal was taken as the mean intensity over the region of the interest. The in vitro data showed a substantial difference in brightness of the two indicators. Since voltage-independent background autofluorescence (presumed to also be independent of the chosen indicator) would comprise different fractions of the signal and decrease the observed relative fluorescence changes, we subtracted the mean intensity of the neuropil surrounding each particular neuron from its signal (I_n_). To detrend the signal and extract the fluorescence changes related to both APs and slower membrane voltage changes (EPSPs, oscillations), we calculated a baseline (B_5s_) using a 5s median filter. The ΔF/F trace was then defined as ΔF/F = 100*(I_n_-B_5s_)/B_5s_. To extract only the AP-related fluorescence spikes, we calculated another baseline (B20) using a 20 ms median filter; ΔF/F_APs_ = 100*(I_n_-B_20ms_)/B20ms. We estimated the noise directly from the ΔF/F_APs_ trace. Based on the fact that the AP-related spikes are all negative-going and APs are generally sparse, positive values of the trace ΔF/F_APs_ can be considered as noise. We removed all negative data points and then randomly assigned positive/negative signs to the rest of the points. We calculated the standard deviation of these values (SD_noise_) for each neuron and set it as a threshold to detect spikes; THR = −4*SD_noise_. If the threshold was crossed at two neighboring time points, such doublet was considered as a single AP, the time point with higher amplitude was chosen by the algorithm and the spike was ascribed to this time point. Using four standard deviations leads to false positivity rate of 1– 2 false spikes per minute in our recordings.

To detect the periods of 3–5 Hz oscillations, we applied a bandpass filter (3–5Hz, MATLAB [bandpass]) to the ΔF/F trace and then detected the pronounced oscillations as outliers from the values’ variance. The bandpassed trace was thresholded by 3*MAD (median absolute deviations) using the MATLAB function [isoutlier]. Absolute values of these outliers were averaged for each neuron and later averaged over all neurons in both groups.

## Results

### High throughput screening of Voltron mutants in neuron culture

Voltron variants were generated using site saturation mutagenesis (SSM) performed at 40 positions within the rhodopsin domain. All screening was performed on Voltron mutants labeled with JF_525_ (Voltron_525_). Positions were chosen based on: (1) previous reports of analogous positions in other opsins that affected their thermal stability (Curnow et al. 2011; Faham et al. 2004; McIsaac et al. 2014; Perálvarez-Marín et al. 2008; Wagner et al. 2013), (2) amino acids in close proximity to the retinal chromophore that we reasoned might affect the environment of the Schiff base, or (3) positions that were found to be important in mutagenesis of Archaerhodopsin into a voltage sensor (Hochbaum et al. 2014) (Fig 1a). We performed two rounds of screening (Fig. 1b). In the first, we screened individual point mutations using a field stimulation assay in primary neuron cultures (Fig. 1c-e). For each variant, parameters relevant to the performance of the sensor *in vivo* were measured: AP sensitivity (ΔF/F_0_), AP rise and decay kinetics (*τ_ON_* and *τ_OFF_*), and baseline fluorescence (F_0_). To control for biological variability, the measured parameters of each construct were also normalized to an in-plate Voltron_525_ control. The control was also used to monitor the quality and consistency of the screen. For a construct screened in a 96-well plate, results were discarded if at least one of the following quality control (QC) criteria (empirically determined) were violated: (1) the average |ΔF/F_0_| of the in-plate Voltron_525_ controls was < 3.6%, (2) the PDI of the plate was > 30% (see Methods), or (3) the construct had < 100 pixels with a significant change in ΔF/F_0_ during the stimulation (“responsive pixels”).

**Figure 1:**
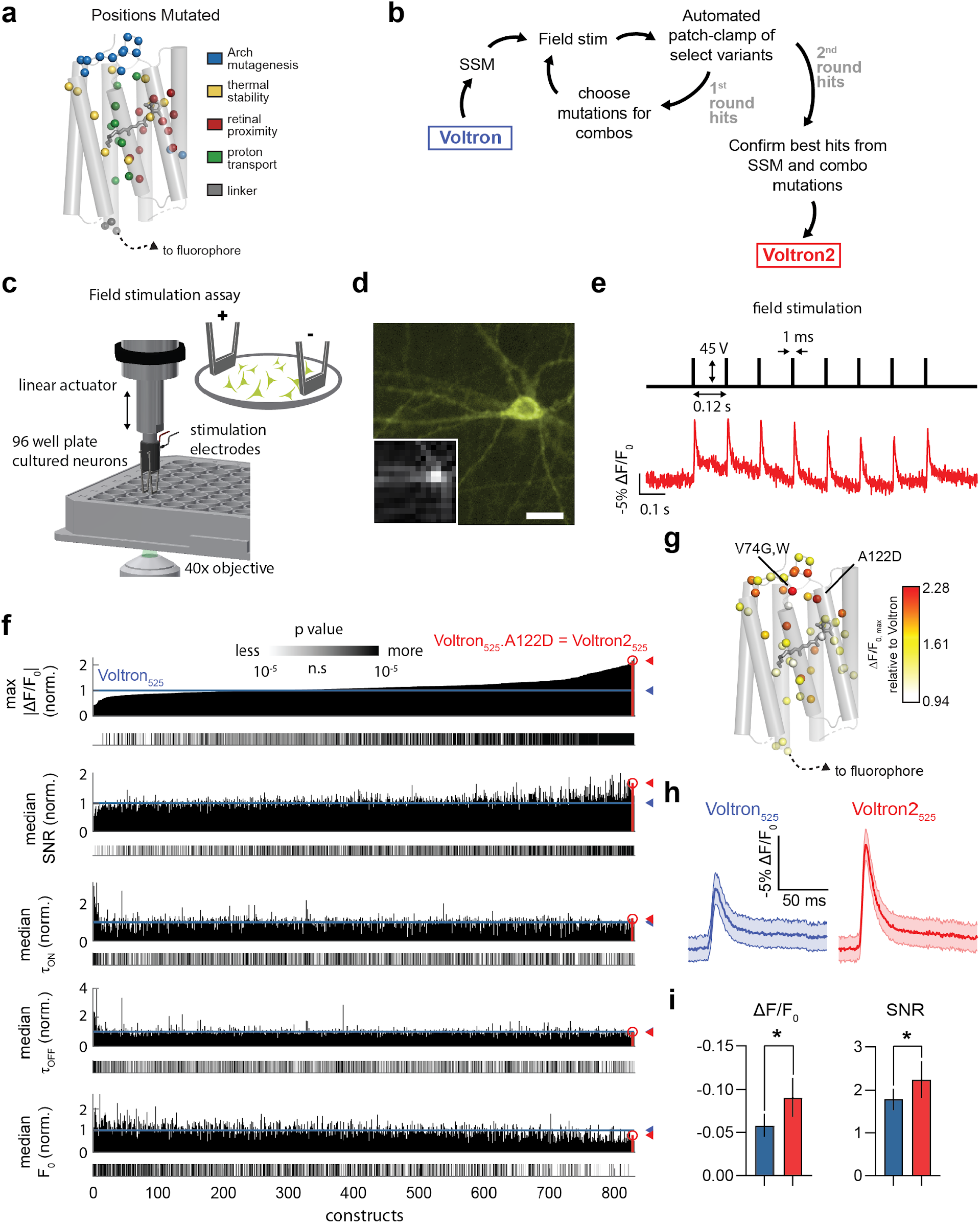
Mutagenesis and screening of Voltron in cultured neurons. a. Residues targeted for SSM in the Ace2N rhodopsin domain of Voltron, colored by the rationale for targeting them. b. Mutagenesis and screening workflow. c. Diagram of field stimulation assay performed in 96-well plates. d. Representative image of neuron from the screen expressing Voltron2 labeled with JF_525_, Voltron2_525_. Inset shows representative frame during fast (1,497 Hz) stream acquisition. Scale bar: 10 μm. e. Field stimulation parameters (top, black) and acquired fluorescence response of the neuron shown in d (bottom, red). All imaging in the screen was performed at a light density of 1.14 mW/mm^2^ measured in the image plane. f. Results of single-mutation Voltron_525_ screen, ranked by maximum |ΔF/F_0_| for each variant, normalized to in-plate Voltron_525_ controls (top). Color-coded p values are shown below each plot, indicating significant difference compared to in-plate controls. g. Mutated residues colored by the maximum increase in |ΔF/F_0_| achieved in that position. Top three mutations are labeled. h. Representative traces (mean ±s.e.m.) from a single plate containing Voltron_525_ (8 wells) and Voltron2_525_ (8 wells). i. Single AP ΔF/F_0_ (Voltron_525_, −.059 ± 0.001, n=338 wells; Voltron2_525_: −0.090 ± 0.002, n=130 wells; P<0.0001, Mann-Whitney U test) and SNR of Voltron2_525_ and SNR (Voltron_525_: 1.80 ± 0.013; Voltron2_525_: 2.24 ± 0.040; P<0.0001, Mann-Whitney *U* test).

Of the 2,727 variants that were screened in 199 plates, 2,314 (84%) passed the above QC criteria. Variants that failed QC were re-screened again and 34% of them passed QC on the second round of screening and were added to the main QC-passed pool. The majority (66%) of the libraries were then sequenced and results from the same mutation were grouped, resulting in 819 QC-passing mutants. We found 422 mutants (51%) with significantly improved ΔF/F_0_, 310 mutants (38%) with increased SNR, 233 mutants (27%) with reduced *τ_ON_*, 256 mutants (31%) with reduced *τ_OFF_*, and 307 (37%) with increased F_0_ compared to Voltron (Fig. 1f, P<0.01, Mann-Whitney *U* test). The key feature of Voltron we desired to optimize was ΔF/F_0_; therefore, we ranked all variants based on |ΔF/F_0_|_max_ (maximum of |ΔF/F_0_|) normalized to in-plate Voltron controls.

Although many variants had improved |ΔF/F_0_|_max_ over Voltron_525_, there was no single top-performing variant in this first round of screening. Instead, the difference in |ΔF/F_0_|_max_ of the top 3 variants was only ~10%, which was under our PDI metric (14±5.2% across the first screening round), indicating that the ranking of the top variants may not be accurate. The top two hits in the screen were Voltron_525_.V74G (|ΔF/F_0_|_max_ relative to Voltron_525_ = 2.28) and Voltron_525_.V74W (|ΔF/F_0_|_max_ relative to Voltron_525_ = 2.21; Fig. 1g, Supplementary Table 1). However, subsequent analysis with patch-clamp revealed that Voltron_525_.A122D (3^rd^ in the ranked |ΔF/F_0_|_max_ list, |ΔF/F_0_|_max_ relative to Voltron_525_ = 2.18) had superior properties as a voltage sensor. The Voltron_525_.A122D mutant (which we named Voltron2_525_) exhibited |ΔF/F_0_|_max_ and SNR that was 52% and 25%, respectively, higher than Voltron_525_ (Fig. 1 h,i).

The first-round SSM screen revealed many mutations that moderately increased ΔF/F_0_. We therefore embarked on a second round of combinatorial (combo) screening, hoping that combining 13 of the top performing mutations (Y63L, N69E, V74E/W, R78H, N81S, L89A/C/G/T, A122D/H, V196P) would further improve the sensor. Of the 1,232 constructs screened in 106 plates, 77% passed QC. Surprisingly, only 28 of 848 combo mutants (3.3%) had significantly improved |ΔF/F_0_|_max_ over Voltron2_525_ (P<0.01, Mann-Whitney *U* test; Supplementary Fig. 1, Supplementary Table 2). Similarly, only a few variants had increased SNR (20 of 848, 2.4%). The A122D substitution was present in 34% of the combo variants passing QC (Supplementary Table 3); nevertheless, the combo screen revealed that combining it with other mutations resulted in less sensitive variants. Subsequent automated patch-clamp analysis confirmed that Voltron2_525_, containing the sole A122D substitution, outperformed all combo mutants (Supplementary Fig. 4).

### Screening and characterization with automated whole-cell electrophysiology

Many single and combo mutation hits from the neuron culture screen had improved |ΔF/F_0_|_max_ over Voltron but had very similar ΔF/F_0_ characteristics among them. We deemed the field stimulation screen to be insufficiently sensitive to find the one variant with the best performance, so we used the uM Workstation, a fully automated whole-cell electrophysiology platform based on the PatcherBot to perform a secondary screen on top single and combinatorial mutant hits.

We first validated the throughput and performance of the automated electrophysiology platform. To mimic a small-scale screen, 10 35-mm Mattek dishes of cultured neurons were transfected with variants of the voltage sensor ASAP (St-Pierre et al. 2014). The uM Workstation made 103 patch-clamp attempts in 7.1 hours, with a 78% whole-cell success rate. The system operated unattended for ~5 hours during that day of screening. Thus, the uM Workstation allowed us to screen ~10 constructs per day, assuming 5-10 neurons per construct.

The uM Workstation achieves high throughput by automatically cleaning and reusing patch-clamp pipettes (Fig. 2a); however, it is conceivable that the cleaning process is imperfect and whole-cell success rate degrades over subsequent attempts. To address this, we evaluated pipette performance after multiple patch-clamp attempts. Whole-cell success rate decreased over time, but likely due to cell health degradation, not due to an accumulation of debris on the reused pipette, since replacing the pipette did not recover the success rate (Supplementary Fig. 2a). In a separate experiment we replaced the dish without replacing the pipette, and found that the success rate recovered, further suggesting that cell health degradation, not pipette debris is responsible for the apparent decrease in success rate (Supplementary Fig. 2b). To explore the limits of pipette cleaning, we patch-clamped cells with the same pipette, replacing the plate as needed, until the time to form a gigaohm seal increased, indicating a contaminated pipette. Consistent with previous observations, a single pipette could be used for patch-clamping ~50 neurons (Kolb et al. 2019) (Supplementary Fig. 2c). Last, we evaluated the quality of the recordings and found 85.6% (143 out of 167) of the successful whole-cell recordings had a holding current greater than –100 pA and access resistance less than 30 MΩ, which meets the criteria for most of the published data acquired with manual patch clamp. Together, we found that the automated uM Workstation successfully increased our throughput, enabling large-scale patch-clamp studies, without compromising data quality.

**Figure 2:**
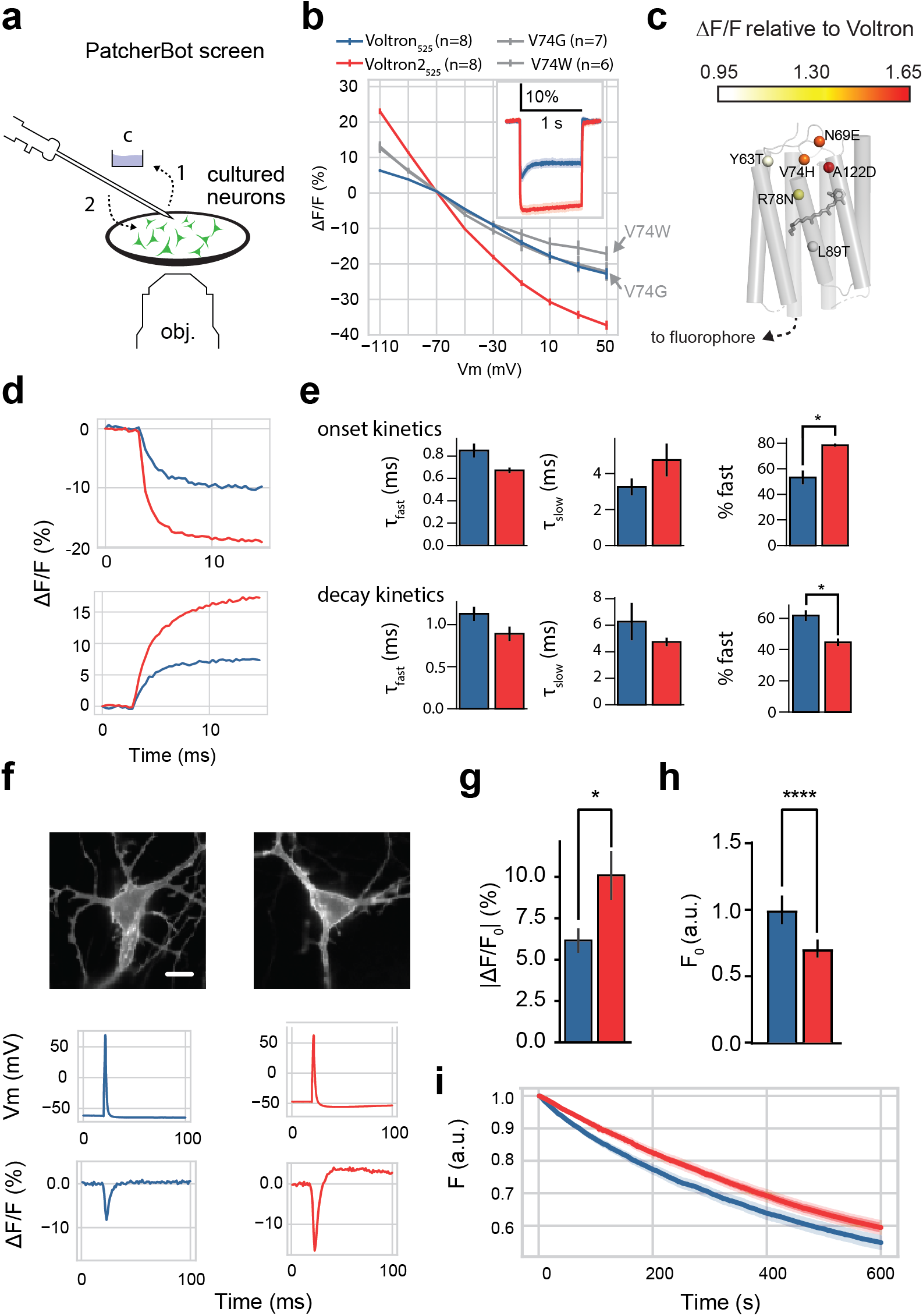
Automated patch-clamp screening and Voltron2 characterization in cultured neurons. a. Fully automated uM workstation screening platform, based on PatcherBot. The pipette cleaning procedure is shown where a used pipette is dipped into a reservoir of cleaning solution (step 1, “c”) and back to the neuronal culture for a subsequent patch-clamp attempt without the need for replacing the pipette (step 2). b. Peak fluorescence response to voltage steps from −70 mV of Voltron_525_, Voltron2_525_ and the top two variants from the field stimulation assay (Voltron2_525_ vs. Voltron_525_: P=0.012; Voltron2_525_ vs. Voltron_525_.V74G: P=0.015; Voltron2_525_ vs. Voltron_525_.V74W: P=0.0003, one-way ANOVA followed by Dunnett’s post-hoc test). Inset: Voltron_525_ and Voltron2_525_ fluorescence traces (mean ± s.e.m.) in response to −70 to +30 mV voltage steps. c. Mutated residues from 1^st^ screening round (single sites) colored by the maximum ΔF/F_0_ response to 100 mV (−70 to +30 mV) voltage steps, measured with the uM workstation. Top mutations at each position are labeled. d. Representative onset (top) and decay (bottom) fluorescence kinetics of Voltron_525_ and Voltron2_525_ in response to a +100 mV voltage step from −70 mV. e. Onset and decay kinetics. Onset kinetics: *P=0.03, Mann-Whitney U test. Decay kinetics: *P=0.03, Mann-Whitney U test. f. Representative fluorescence responses to single evoked APs in current clamp. Scale bar: 10 μm. g. ΔF/F_0_ in response to single AP stimulation in current clamp mode (*P = 0.03, Student’s unpaired t test). h. Normalized resting fluorescence relative to mTagBFP2 fused to the C terminus (****P<0.0001; Voltron_525_: n=105 cells, Voltron2_525_: n=115 cells). i. Photobleaching comparison of Voltron_525_ and Voltron2_525_ over 10 mins.

Using the uM Workstation we then screened top-performing single-position mutants from the field stimulation screen, including Voltron as a control. While Voltron_525_.V74G and Voltron_525_.V74W were the top performers from the field stimulation screen, their fluorescence response to a 100 mV voltage step was lower than that of Voltron2_525_ (Fig. 2b). The other mutants were also 8% to 55% less sensitive to 100 mV voltage steps than Voltron2 (Supplementary Fig. 3). Meanwhile, Voltron2_525_ was found to be 65% more sensitive than Voltron, consistent with the field stimulation screen. Furthermore, in the physiologically relevant sub-threshold voltage range (−90 to −50 mV), Voltron2_525_ exhibited a significantly steeper slope than Voltron_525_ (0.54+0.01 and 0.21+0.01%/mV, respectively; P = 0.0009, Mann-Whitney U test), making it a higher-fidelity optical reporter of changes in sub-threshold membrane potential.

Surprisingly, the combo mutation screen (second round of the field stimulation assay, Fig. 1b) yielded few variants with improved sensitivities. We nevertheless screened the 34 variants with sensitivities marginally better than Voltron2_525_ using the uM Workstation. As was the case with the single-position mutants, we found no combo mutants that out-performed Voltron2_525_ (Fig. 2c, Supplementary Fig. 4). Therefore, for the remainder of this study, we focused on characterization of Voltron2_525_.

Voltron2_525_ exhibited fast onset and decay kinetics that were best fit with a double exponential (Fig. 2d). Interestingly, the A122D mutation completely eliminated the transient peak in the fluorescence response of Voltron_525_ (Fig. 2b inset). The fast component of the onset and decay kinetics was slightly shorter for Voltron2_525_ (onset: 0.67±0.03 ms, decay: 0.89±0.09 ms) compared to Voltron_525_ (onset: 0.85±0.06 ms, decay: 1.13±0.08 ms), though not significantly different. The slow components were likewise similar between the two sensors (Voltron_525_: onset 3.26±0.47 ms, decay 6.27±1.41 ms; Voltron2_525_: onset 4.76±0.92 ms, decay 4.74±0.32 ms). The fast component of Voltron2_525_ accounted for a larger percentage of the overall response in the onset but not decay response (Fig. 2e). Overall, the kinetic properties of Voltron_525_ and Voltron2_525_ were found to be similar.

Consistent with the improved sensitivity of Voltron2_525_ in response to voltage steps, it was also superior in its sensitivity to APs. Voltron2_525_ reported single APs with ΔF/F_0_ of 10.09+1.47%, significantly higher than for Voltron_525_ (6.16+0.74%, Fig. 2 f,g). The baseline fluorescence of Voltron2_525_ was ~30% lower than Voltron_525_, which may be beneficial in some experiments but detrimental in others (Fig. 2h). The same trend was observed with the addition of the soma localization tag (Supplementary Fig. 5). Nevertheless, both Voltron2_525_ and Voltron2_525_-ST showed good membrane localization, qualitatively similar to their Voltron counterparts (Supplementary Fig. 6, 7). The addition of a soma localization tag to Voltron2_525_ increased its sensitivity to a 100 mV depolarization pulse by ~18% (Supplementary Fig. 8). In culture, Voltron2_525_ photobleached slightly, but not significantly, slower than Voltron_525_ (Voltron_525_: 45±2%, Voltron2_525_: 41±1% reduction in fluorescence; P=0.11, Mann-Whitney U test; Fig. 2i).

We reasoned that the A122D mutation responsible for the increased sensitivity of Voltron2_525_ could have beneficial properties when grafted onto other Ace rhodopsin-based GEVIs. We tested this hypothesis in Ace2N-mNeon and VARNAM. As expected, adding the A122D mutation to both GEVIs increased their sensitivity to depolarizing and hyperpolarizing voltage pulses (Supplementary Fig. 9). Similar to Voltron2_525_, A122D significantly increased the slope of the sensors in the sub-threshold range (Ace2N-mNeon: 0.091±0.012 %/mV, Ace2N-mNeon.A122D: 0.303±0.012%/mV, P=0.006; VARNAM: 0.104±0.012%/mV, VARNAM.A122D: 0.147±0.010/mV, P=0.045; Mann-Whitney U test). The mutation eliminated the transient peak from VARNAM but not from Ace2N-mNeon. Grafting A122D onto Positron did not result in increased sensitivity (Supplementary Fig. 10); however this was not surprising given that the proton transport pathway in Positron is different from Voltron (Abdelfattah et al. 2020). Together, the results suggest that the A122D mutation appears to generalize across different FRET donors.

### Voltage imaging and stimulation in acute brain slices

The high sensitivity of Voltron2 in the sub-threshold range of voltages should make it a suitable GEVI for detecting low-amplitude voltage fluctuations, such as those arising as a result of synaptic activity. To test this, synthetic PSPs (synPSPs) were injected into neurons expressing Voltron_585_ and Voltron2_585_ in acute mouse brain slices (Fig. 3a). Optically captured responses to PSPs were ~40% larger for Voltron2_585_ than Voltron_585_, consistent with the improved sensitivity of Voltron2_585_ in the sub-threshold range (Fig. 3b,c). The overall ΔF/F_0_ in response to ±15 mV synPSPs was 5.9% for Voltron2_585_, compared to 3.2% for Voltron_585_ (Fig. 3d top). Due to the increased sensitivity, the detectability of synPSPs was found to be significantly improved for Voltron2_585_ (Fig. 3d bottom). Together, we found that Voltron2_585_ could be used to image millivolt-scale synaptic events.

**Figure 3:**
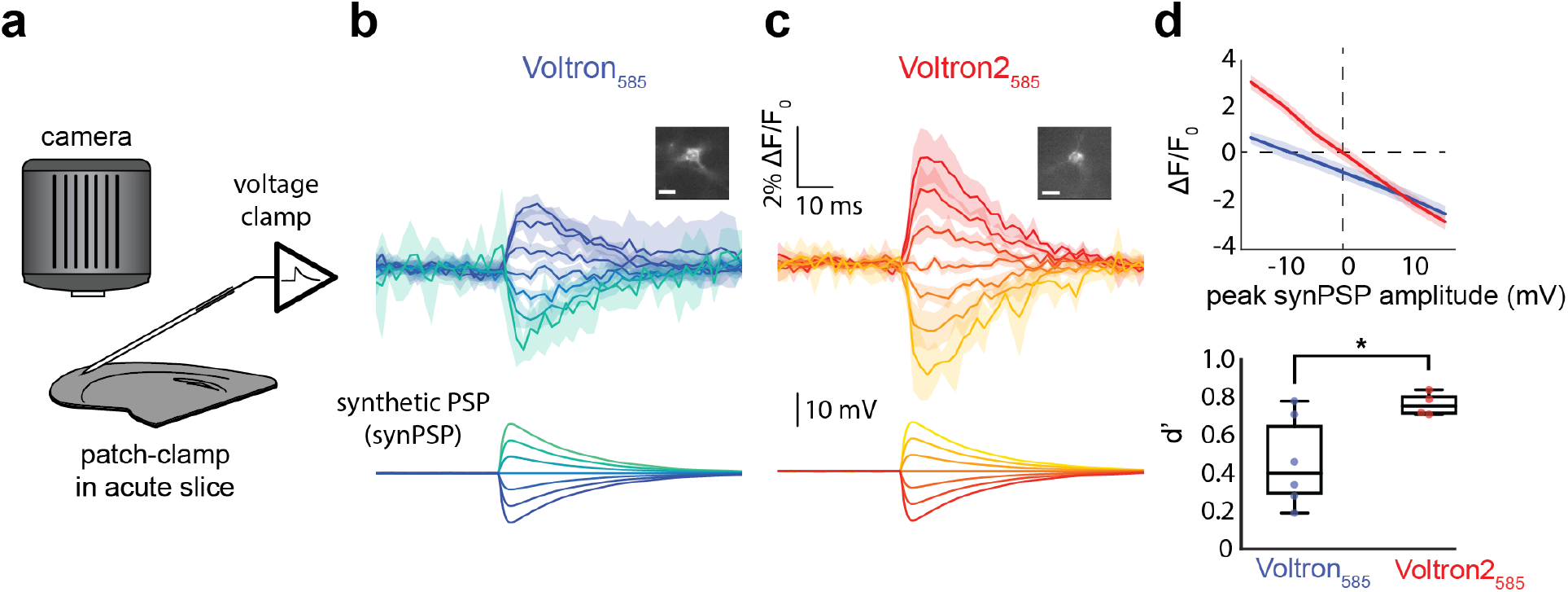
PSP detection using Voltron2 in mouse brain slices. a. Synthetic PSP (synPSP) experimental setup in acute mouse brain slice. b,c. Average ± SD of percent change in fluorescence over time for Voltron_585_ (b; n=6 cells) or Voltron2_585_ (c, n=4 cells) in response to changes from resting membrane potential of −15mV to +15mV in 5mV increments (lower panels), intended to mimic typical inhibitory or excitatory synaptic transmission. A representative cell for each construct is shown in the inset (scale bar = 10μm). d. Top: average ± SD of percent change in fluorescence as a function of the peak amplitude of the synthetic postsynaptic potential (synPSP) applied to the cell. Voltron does not cross the 0,0 point due to photobleaching in the course of the experiment. Bottom: sensitivity index (d’) of Voltron2_585_ is significantly higher than that of Voltron_585_ (P=0.025, Welch’s t-test).

We then evaluated the ability of Voltron2 to be used in the context of all-optical electrophysiology. Here, we expressed Voltron2_585_-ST along with ChR2-GFP (Boyden et al. 2005) in acute slices of mouse motor cortex (Fig. 4a). We confirmed with whole-cell electrophysiology that ChR2 could reliably elicit spiking activity when illuminated with moderate blue light intensity (30 μW/mm^2^) and that Voltron2_585_ accurately tracked the membrane voltage. Increasing the green excitation light intensity improved the SNR as expected but resulted in well-documented cross-excitation issues (Klapoetke et al. 2014; Packer et al. 2015) (Fig. 4b). Using the same illumination intensity, we imaged a FOV with 10 neurons during repeated ChR2 activation and found robust Voltron2_585_ signals that reported expected increases in spiking activity during ChR2 stimulation (Fig. 4c,d). These experiments suggest that Voltron2_585_ can be used with optogenetic actuators for all-optical interrogation of brain circuitry.

**Figure 4:**
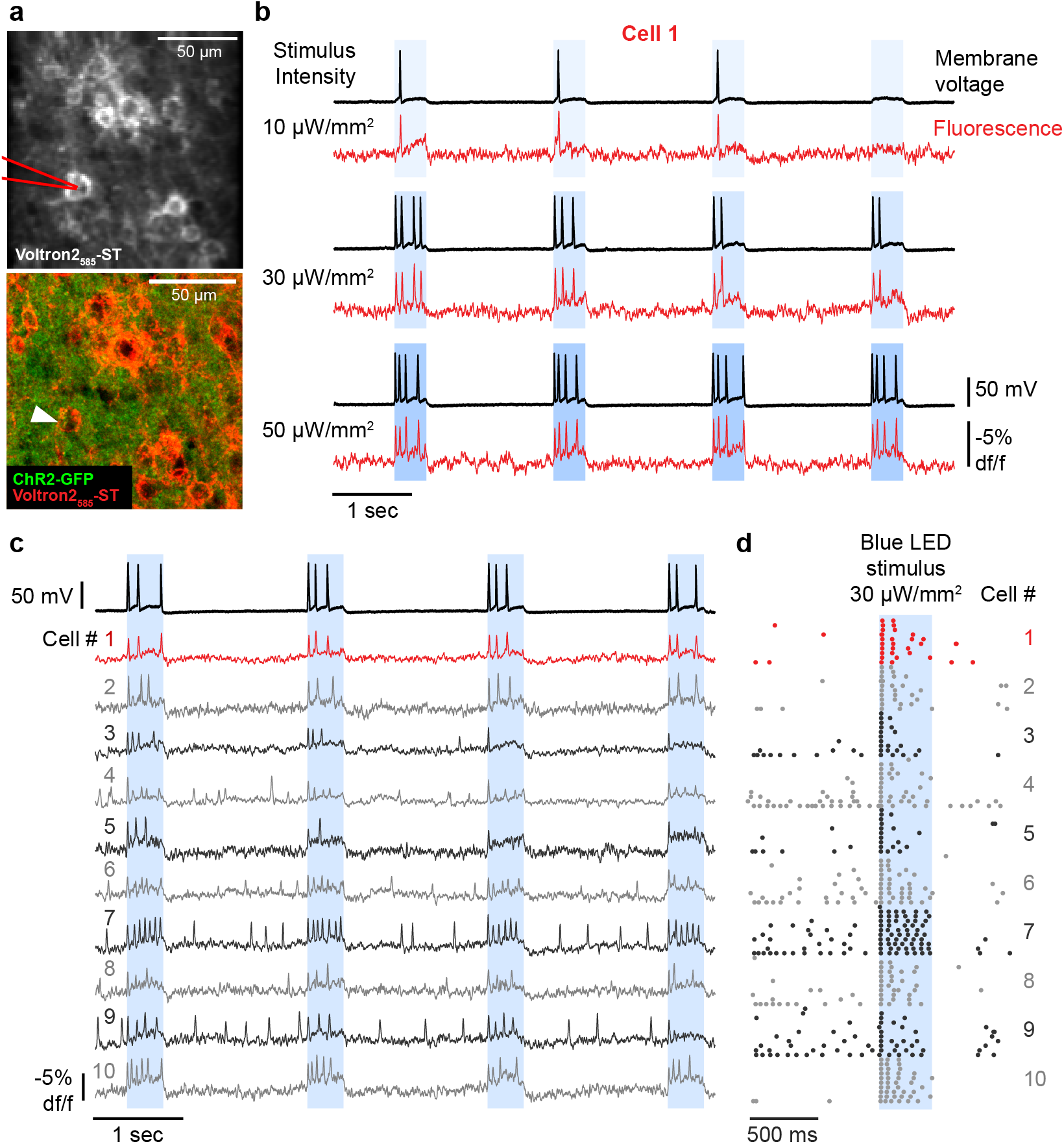
Simultaneous voltage imaging and optogenetic stimulation. **a.** (Top) Average intensity projection of 457 Hz confocal images showing Voltron2_585_-expressing cells labeled with JF585 in an acute slice of motor cortex. Pipette used for whole-cell recordings illustrated in red. (Bottom) Post-hoc confocal image showing pan-neuronal expression of ChR2-GFP in the same field of view (FOV) shown in top panel, with patched cell #1 indicated by white arrow. **b**. Whole-cell membrane voltage (black traces) and corresponding Voltron2 fluorescent signal (red traces) from patched cell #1 shown in **a**, showing responses to 400 ms stimulation with 10 (top), 30 (middle), and 50 μW/mm^2^ (bottom) blue light. **c.** Voltron2_585_ signals (red and gray traces) recorded across 10 distinct cells in the FOV shown in **a** in response to 400 ms stimulation with 30 μW/mm2 blue light. Corresponding membrane voltage is shown for patched cell # 1 (upper black trace). **d.** Raster plots show trial-aligned APs detected in fluorescent signals from cells #1-10 shown in **a** and **c**, across 10 repeated 400 ms blue stimulus trials.

### *In vivo* voltage imaging of olfactory sensory neurons in zebrafish

We next tested Voltron2_552_ side-by-side with Voltron in olfactory sensory neurons in larval zebrafish using a lattice lightsheet microscope (Fig. 5a). Volton2552 exhibited higher-amplitude spontaneous spiking and subthreshold activity than Voltron_552_ (Fig. 5b). The ΔF/F_0_ and SNR of detected spikes was significantly higher for Voltron2_552_, measured across hundreds of cells (Fig. 5c). Both Voltron2_552_ and Voltron_552_ were imaged over 5 minutes, with voltage signals still clearly visible at the end of the experiment, suggesting that longer recording sessions are also possible.

**Figure 5:**
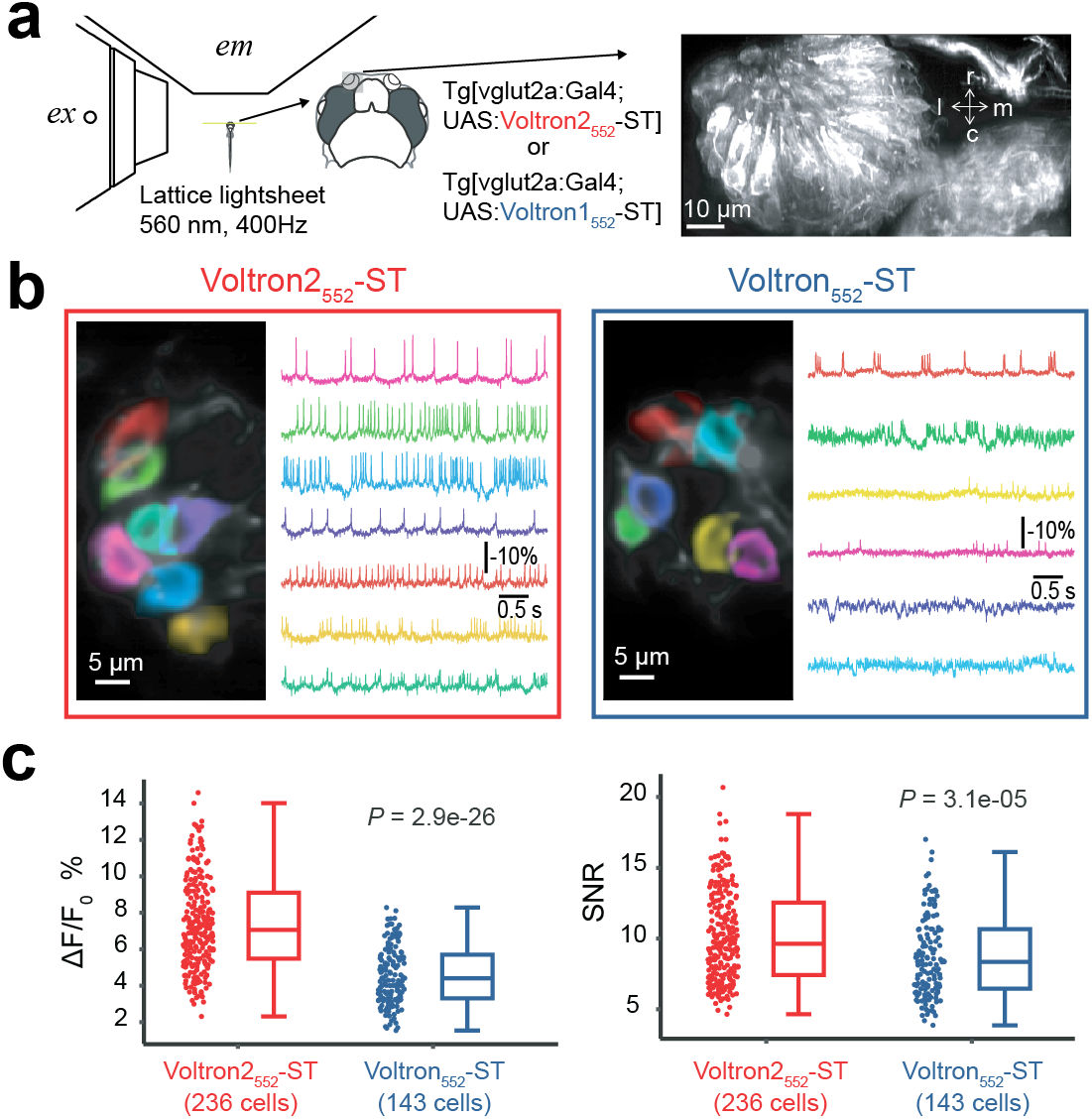
*In vivo* comparison of Voltron2-ST and Voltron-ST in zebrafish olfactory sensory neurons. a. Experimental setup. Left: Olfactory sensory neurons expressing Voltron2-ST or Voltron-ST, labeled with JF_552_ and imaged at 400 Hz using a lattice-lightsheet microscope. ex: excitation objective lens, em: imaging objective lens. Right: Volumetric rendering of olfactory sensory neurons in the nasal cavity. r, rostral; c, cadual; m, medial; l, lateral b. Representative FOVs and recordings. Spatial weights optimized for individual spiking neurons are shown in distinct colors over the structural image (left). The activity trace of corresponding neurons is shown in the same color (right). c. Performance comparisons of Voltron2_552_-ST and Voltron_552_-ST. Left: Distribution of spike-related fluorescence change of Voltron2_552_-ST and Voltron_552_-ST. Right: Distribution of SNR of Voltron2_552_-ST and Voltron_552_-ST. Statistical differences were assessed with the Wilcoxon rank-sum test.

### *In vivo* voltage imaging in adult *Drosophila melanogaster*

We tested Voltron2 in voltage recordings of spontaneous activity from two neuron types in the mushroom body (MB) circuit of adult *Drosophila melanogaster*, the output neuron MBON-γ1pedc>α/β and the dopaminergic neuron PPL1-γ1pedc (Aso et al. 2014). The expression of Voltron2 was driven by split Gal4 lines (*MB112C* and *MB320C*), which uniquely target these neurons, enabling a well-matched comparison of sensor performance across different flies. We imaged both cell types in the γ1 compartment, which contains the dendritic processes of MBON-γ1pedc>α/β and the axonal terminals of PPL1-γ1pedc. We used JF_552_-HaloTag conjugate (Fig. 6a). Among several JF dyes we tried in *Drosophila* neurons, we found that JF_552_ allowed for prolonged Voltron imaging, which in PPL1-γ1pedc can last over 20 min without significant deterioration of the health of the cell (unpublished observations, YS). JF_552_ is a JF_549_ analogue with fluorine substitution on the xanthene ring, which shows improved cell and tissue permeability (Zheng et al. 2019). Spike amplitudes (ΔF/F_0_) measured with Voltron2_552_ were significantly larger when compared to Voltron_552_ (Fig. 6b,d). The mean spike size is 74% larger in MBON-γ1pedc>α/β (Fig. 6c), and 57% larger in PPL1-γ1pedc (Fig. 6e). The SNR is also increased, most notably in MBON-γ1pedc>α/β (Fig. 6f,h). The basal florescence levels are lower with Voltron2 though (Fig. 6g,i), which contributes to the more moderate improvement of SNR as compared to ΔF/F.

**Figure 6:**
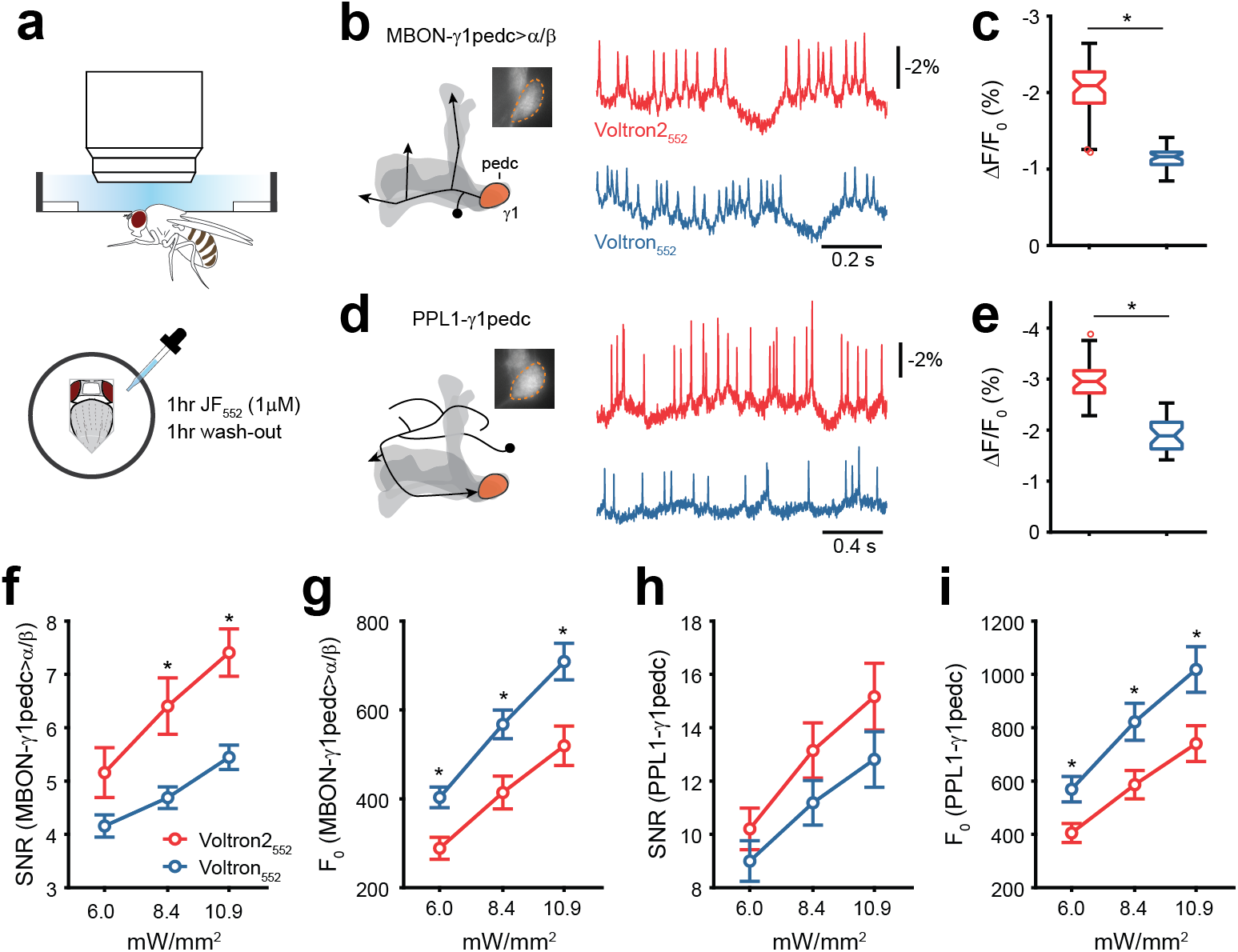
a. Experimental setup. A head-fixed fly is imaged using an sCMOS camera at 800 frames per second. Voltron is loaded with JF_552_-Halotag ligand via a 1hr incubation/1hr wash-out protocol. b,d. Voltage recordings in MBON-γ1pedc>α/β (*MB112C-Gal4*) and PPL1-γ1pedc (*MB320C-Gal4*). Neuron schematics are shown for the left hemisphere with the MB in shaded gray (arrowheads indicate axonal outputs). Fluorescence images were acquired from the γ1 compartment (inset, 50 μm × 50 μm), which contains dendrites of MBON-γ1pedc>α/β and axon terminals of PPL1-γ1pedc. Single-trial recordings of ΔF/F traces are shown (8.4 and 6.0 mW/mm^2^ for b and d respectively). c. Spike amplitude with Voltron2_552_ and Voltron_552_ in MBON-γ1pedc>α/β. P < 0.001, Wilcoxon rank sum test. For Voltron2_552_, the data set was from 15 hemispheres (8 flies) at three levels of illumination for a total of 45 experiments, for Voltron_552_, 13 hemispheres (7 flies) with 39 experiments. e. Spike amplitude in PPL1-γ1pedc. P < 0.001, Wilcoxon rank sum test. For both Voltron2_552_ and Voltron_552_, the dataset was from 10 flies at three levels of illumination for 30 total experiments. f,h. SNR calculated as spike amplitude over standard deviation of the spike-free zones of the trace. P = 0.07, 0.006, 0.003 between Voltron2_552_ and Voltron_552_ in MBON-γ1pedc>α/β, P = 0.28, 0.16, 0.17 in PPL1-γ1pedc, two-sample t-test. g,i. Lower basal florescence levels with Voltron2_552_. P < 0.01 in MBON-γ1pedc>α/β, P < 0.05 in PPL1-γ1pedc, two-sample t-test. * indicates P < 0.05.

### *In vivo* voltage imaging in mouse hippocampus and visual cortex

We next tested Voltron2-ST *in vivo* in parvalbumin (PV) expressing interneurons in the CA1 region of the mouse hippocampus. Cells expressing soma-targeted Voltron2_552_-ST and labeled with JF_552_ were individually illuminated using a DMD-based patterned illumination microscope, and the fluorescence responses of up to 34 neurons were imaged simultaneously at 2000 frames per second (Fig. 7a,b). Spontaneous APs in PV-positive interneurons induced nearly two-fold larger fluorescence changes in Voltron2_552_ compared to Voltron_552_ expressing neurons (Fig. 7c,d). The baseline fluorescence was dimmer for Voltron2_552_ (Fig. 7e), leading to slightly larger recording noise (Fig. 7f). Yet the overall SNR was still significantly improved compared to Voltron_552_ (Fig. 7g). The number of visually identifiable neurons was comparable despite the dimmer baseline fluorescence (Supplementary Fig. 11a). Furthermore, photobleaching was significantly slower in Voltron2_552_ compared to Voltron_552_-expressing cells (Supplementary Fig. 11b; 9.9%±5.8% vs. 15.6±4.6 % in 3 minutes, respectively).

**Figure 7:**
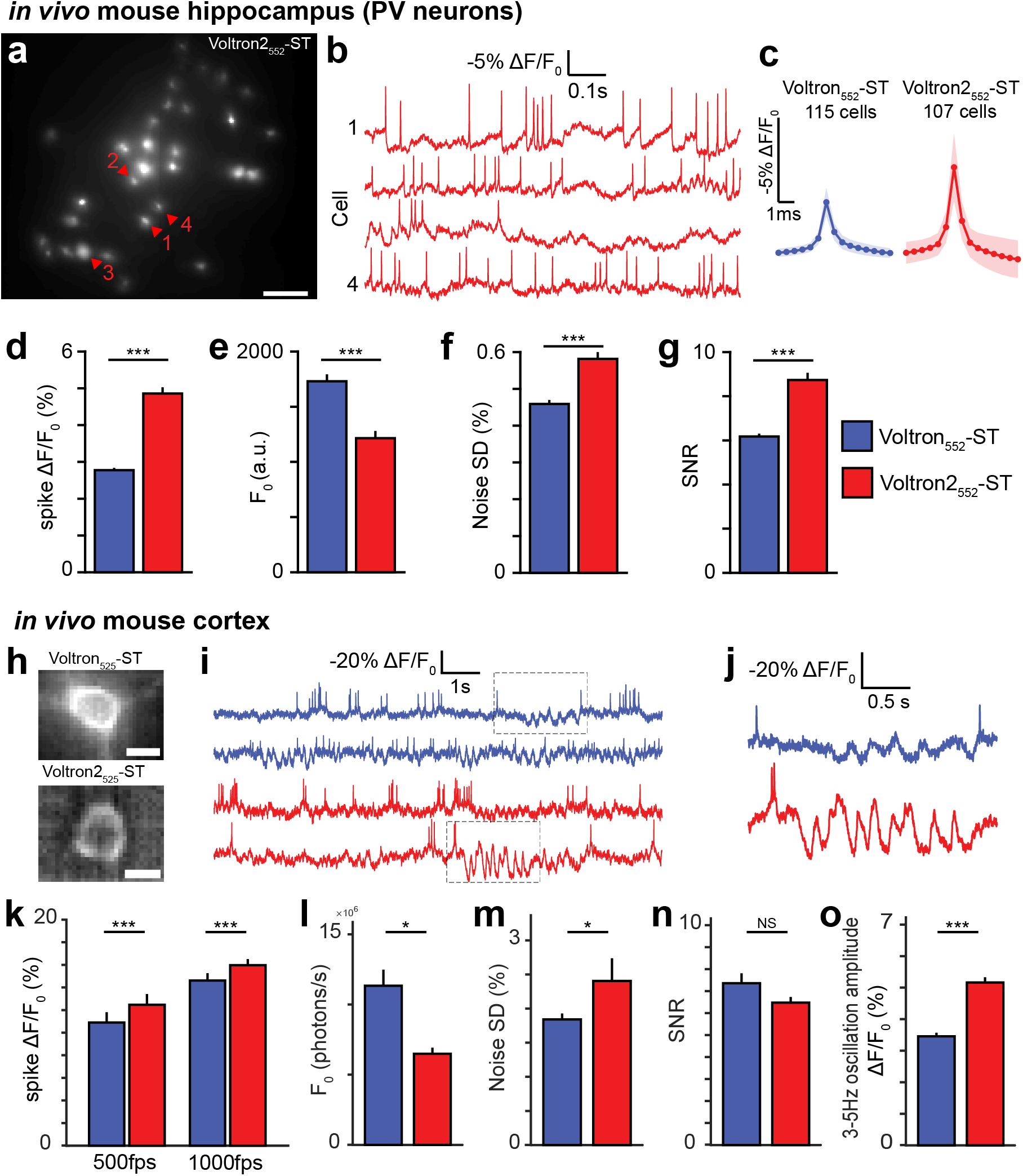
Imaging of voltage activity *in vivo* in mouse hippocampus and cortex with Voltron and Voltron2. a. Example image of hippocampal PV neurons expressing Voltron2-ST labeled with JF_552_. b. Sample fluorescence traces of cell 1 to 4 in (a). c. Average spike waveforms of cells expressing Voltron_552_-ST or Voltron2_552_-ST. d-g. Comparison of Voltron_552_-ST and Voltron2_552_-ST spike amplitude (d), baseline fluorescence (e), noise standard deviation (f), and SNR (g) in hippocampal PV neurons. h. Example images of cortical pyramidal neurons expressing Voltron-ST (top) or Voltron2-ST (bottom) labeled with JF_525_. Scale bar: 10 μm. i. Example fluorescence traces from individual neurons recorded using Voltron_525_-ST (blue) and Voltron2_525_-ST (red) detrended using a 5s median filter. Grey dashed boxes indicate detection of 3-5Hz oscillations shown in (j) and quantified in (o). j. Zoomed portions of the fluorescence traces in (i) showing spikes and 3-5Hz oscillations. k-o. Comparison of Voltron_525_-ST and Voltron2_525_-ST spike amplitude at both 500 and 1000 frames per second imaging rates (k), baseline fluorescence (l), noise standard deviation (m), SNR (n), and 3-5Hz oscillation amplitude (o) in cortical pyramidal neurons. For all plots: Statistically significant differences between groups were determined by Wilcoxon rank-sum test. * p < 0.05, ** p < 0.01, *** p < 0.001. Error bars indicate SEM.

Voltron2_525_-ST was then evaluated and benchmarked against Voltron_525_-ST in the mouse primary visual cortex. We used one-photon epifluorescence microscopy with structured illumination and the same protocol as previously (Abdelfattah et al. 2019). The chronic cranial window and a mixture of Cre-dependent Voltron and dilute CaMKIIa-Cre viruses enabled sparse, but very bright, labeling of pyramidal neurons (Fig. 7h). APs and subthreshold fluctuations were clearly observable using both sensors (Fig. 7i,j). Voltron2_552_-ST produced larger ΔF/F responses from spikes in cortical pyramidal neurons (Fig. 7k), and higher imaging rate led to larger responses from both indicators. Similar to other preparations, we observed that Voltron_552_-ST was significantly brighter than Voltron2_552_-ST (Fig. 7l). As shot noise is dominant in high-speed imaging, we observed smaller relative noise in the brighter Voltron_552_-ST-expressing neurons compared to Voltron2_552_-ST-expressing neurons (Fig. 7m). There was no significant difference in SNR between the two sensors in this preparation (Fig. 7n), likely because the improved ΔF/F of Voltron2_552_ was offset by its higher noise. We subsequently focused on the improved sensitivity of Voltron2_552_ around resting membrane potential. Low-frequency membrane voltage oscillations in individual cortical neurons in awake mice have previously been observed in the barrel (Crochet and Petersen 2006), auditory (Zhou et al. 2014) and visual cortices (Bennett et al. 2013). We focused on brief (1–2 s long) periods of 3–5 Hz oscillations around a ~12 mV hyperpolarized baseline, exhibiting a peak-to-peak amplitude of ~17 mV (Einstein et al. 2017). Due to the enhanced sensitivity of Voltron2_552_ in the subthreshold range, 3-5Hz oscillations were significantly more pronounced when imaging Voltron2_552_-ST, exhibiting ~50% larger amplitude (Fig. 7j,o). Together, these data indicate that Voltron2_552_ significantly improves the quality of in vivo voltage imaging in multiple regions of the mouse brain.

## Discussion

Here we present Voltron2 which contains a mutation to Voltron that increased the sensitivity of the GEVI by > 50% to APs in culture and *in vivo*. Moreover, Voltron2 is approximately 3-fold more sensitive to subthreshold changes due to its steeper slope around the resting membrane potential. As with our efforts to engineer positive-going FRET sensors (Abdelfattah et al. 2020), the mutation we discovered generalized to other Ace-based GEVIs with fluorescent protein reporters. In both Ace2N-mNeon and VARNAM, the A122D substitution increased sensitivity, particularly in the sub-threshold voltage range. Perhaps the sensitivity-improving mutations identified in our screen will also be useful for optimization of rhodopsin-only GEVIs, such as those based on Arch, that rely on imaging the dim retinal fluorescence directly.

Engineering improved GEVIs has been more challenging than GECIs. In this study, we screened >2,700 variants to attain ~50% increase in ΔF/F_0_ in the Voltron scaffold. Applying the same mutagenesis and screening strategy to GCaMP, RCaMP and R-GECO1 calcium indicators yielded >500% increases in ΔF/F_0_ with <1,000 screened variants (Dana et al. 2016, 2019). Further, combining mutations in GCaMP scaffolds has often yielded additive benefits, while doing so in the current context of the Ace2N rhodopsin ultimately did not produce any variants with significant improvements over the best single A122D mutation. It is possible that there are biophysical phenomena that impose a ceiling on the sensitivity of this scaffold. For example, it is expected that the FRET efficiency between the HaloTag-dye or FP donor and the rhodopsin retinal acceptor will limit the maximum fluorescence change. Each of these chromophores resides on or in a bulky protein domain, limiting their closest approach distance. We were intrigued to observe that the A122D mutation improved the sensitivity of the negative-going Voltron, but also decreased its fluorescence at resting membrane potential. It seems possible that additional mutations could restore the original resting fluorescence of Voltron while maintaining the improved sensitivity of A122D, leading to improved SNR, but our screens failed to identify such a variant. Mutations at the equivalent position of Ace2 A122D in bacteriorhodopsin changed the thermal stability of that protein (Wagner et al. 2013).

Various high-throughput platforms have been developed that have been used to screen for improved GEVIs (Chien et al. 2015; Kannan et al. 2018; Park et al. 2013; Piatkevich et al. 2018). The majority of these platforms utilize bacteria or tissue culture cells for screening. We instead opted to perform our high-content primary screen in dissociated neurons, a costlier and more time-consuming strategy, but one that maximized the compatibility of the resulting sensor with *in vivo* neuronal imaging. Even still, our field stimulation screen was insufficiently sensitive to disambiguate the top-performing sensors. We therefore relied on the automated patch-clamp system that afforded us the ability to screen dozens of sensors faster than possible manually, without compromising data quality. The system had a lower throughput than the field stimulation screen but enabled us to characterize the sensitivity and kinetics of many variants with much higher fidelity. The combination of both field stimulation and patch-clamp screens provided a high-quality assessment of top-performing variants.

We show that like its predecessor, Voltron2 can be readily used for *in vivo* imaging in mice, flies, and fish, as well as acute brain slice imaging in mice. These experiments generally confirm the characteristics of Voltron2 that we discovered in our cell culture screen; namely increased ΔF/F_0_ (particularly in the sub-threshold range), improved SNR, reduced baseline fluorescence, and reduced photobleaching in some preparations. These improvements were consistent among the dye ligands that were tested in vivo (JF_525_, JF_552_, and JF585). The increased sensitivity of Voltron2 in the subthreshold range was shown to significantly improve the detectability of 10-20 mV oscillations. Given the richness of information contained within subthreshold activity, including excitatory and inhibitory PSPs, oscillations of various frequencies, spikelets, and other features, Voltron2 can be a valuable tool for unveiling neuronal computations in intact preparations. In addition, in multiple preparations, Voltron2 extended the functional imaging time, which will push the limits of the biology that can be studied.

Increasing the sensitivity of GEVIs (the difference in photon flux per millivolt change in membrane potential) and reducing photobleaching still remain the main challenges to increase the adoption of GEVIs for *in vivo* experimentation. Protein engineering efforts devoted to creating two-photon-compatible GEVIs will also be required to address the emerging trend in the field to image deep in the brain while maintaining single-cell resolution. Chemigenetic indicators like Voltron2 continue to be promising scaffolds to address these goals.

## Author contributions

We use the CRediT taxonomy to clarify author contributions.

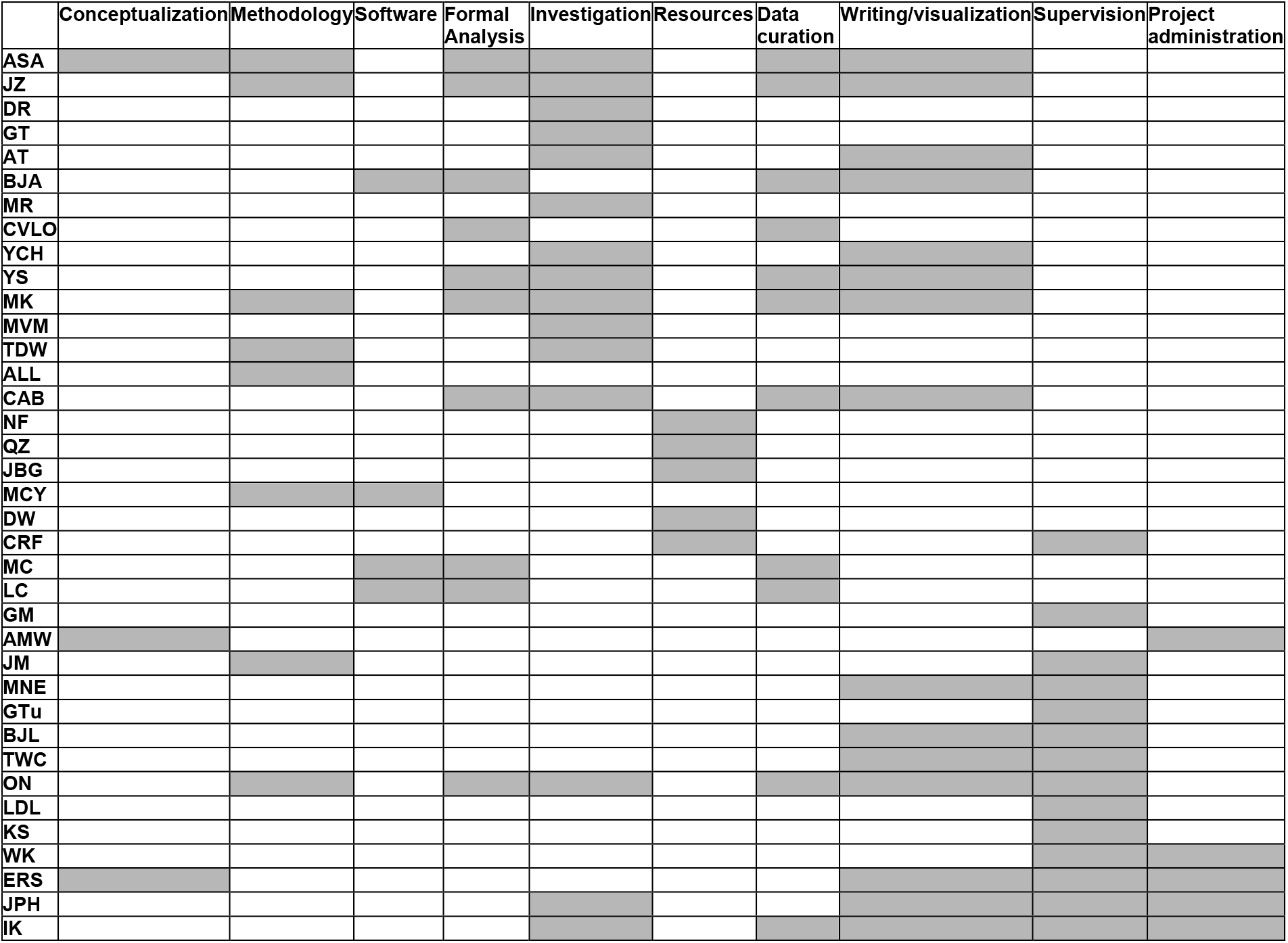

## Acknowledgements

We thank the Janelia cell culture, viviarium, jET, Molecular biology, and Virus Production teams for their assistance. We thank S. Picard, K. Aswath, S. Shrestha for help with DeepSeq. ON, MR, CVLO acknowledge funding from Charles University Grant – PRIMUS/19/MED/003.

## Competing interests

A.S.A., L.D.L., and E.R.S. have filed for a patent on chemigenetic voltage indicators. IK and CRF are co-inventors on a patent describing pipette cleaning that is licensed by Sensapex. MC and LC have performed consulting services for Sensapex.

---

**Supplementary Figure 1:**
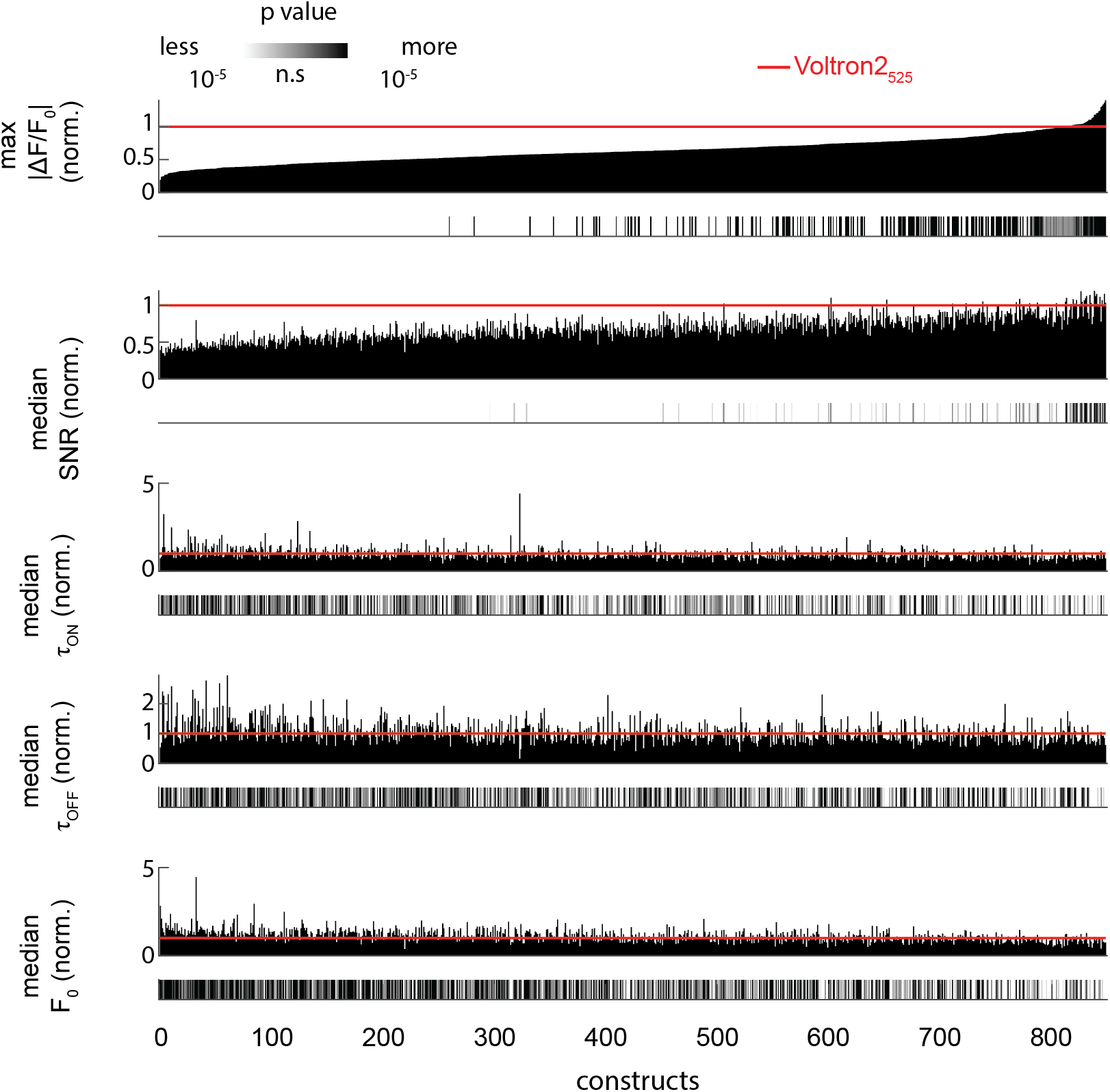
Results of combo mutation screen, ranked by maximum |ΔF/F_0_| for each variant, normalized to in-plate Voltron2_525_ controls. P values indicate significant difference compared to Voltron_525_ in-plate controls.

**Supplementary Figure 2:**
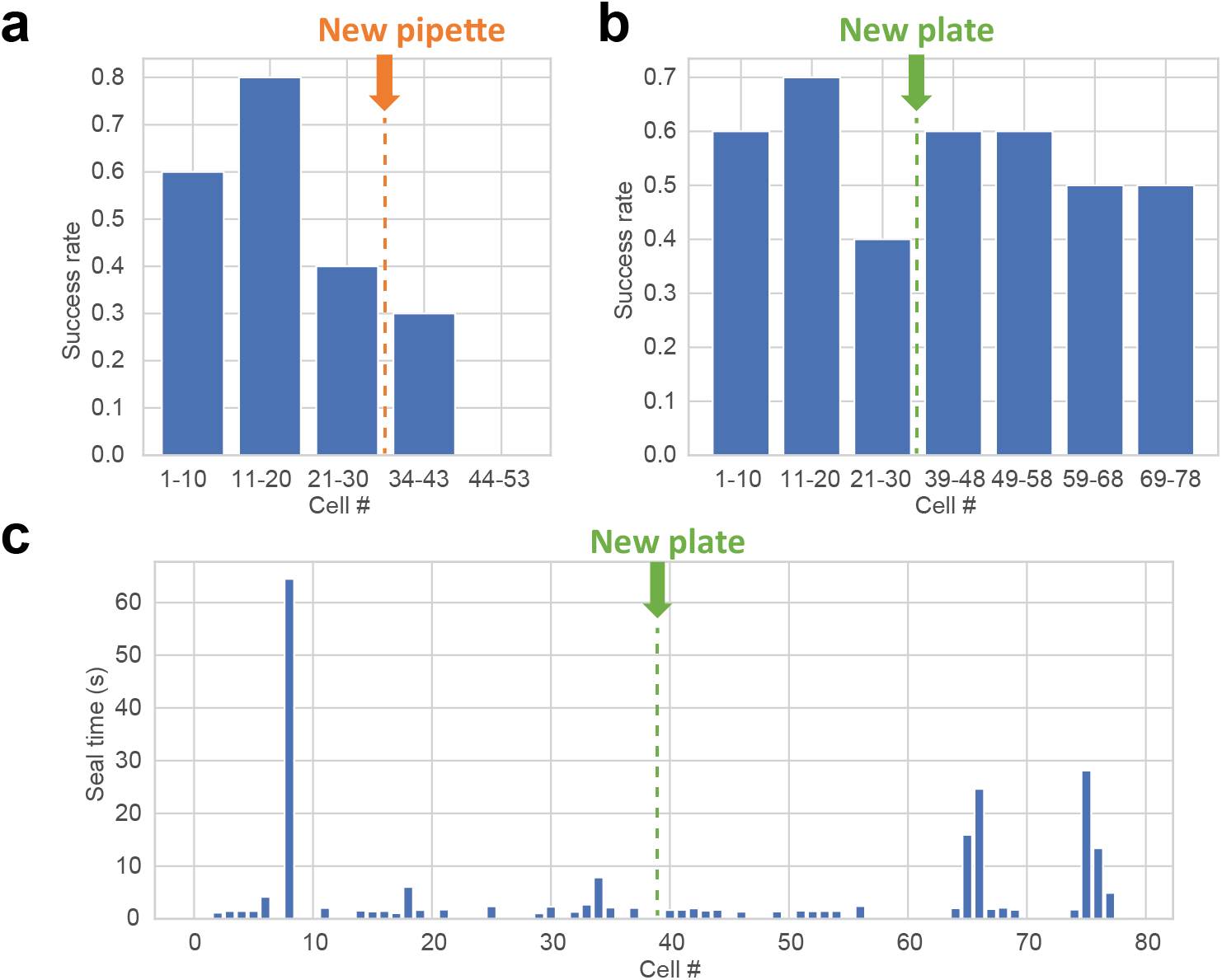
Pipette cleaning with the uM Workstation. a. Whole-cell recording success rate with pipette cleaning after every recording. The pipette was replaced after 30 recordings, but the success rate did not improve. b. Whole-cell recording success rate with a single reused pipette. A new plate of neurons was used after the 30^th^ cell, causing the success rate to improve. c. Time to form a gigaseal over multiple cells using a single pipette. A blank entry indicates that the gigaseal was unsuccessful. A new plate of neurons was used after the 38^th^ cell.

**Supplementary Figure 3:**
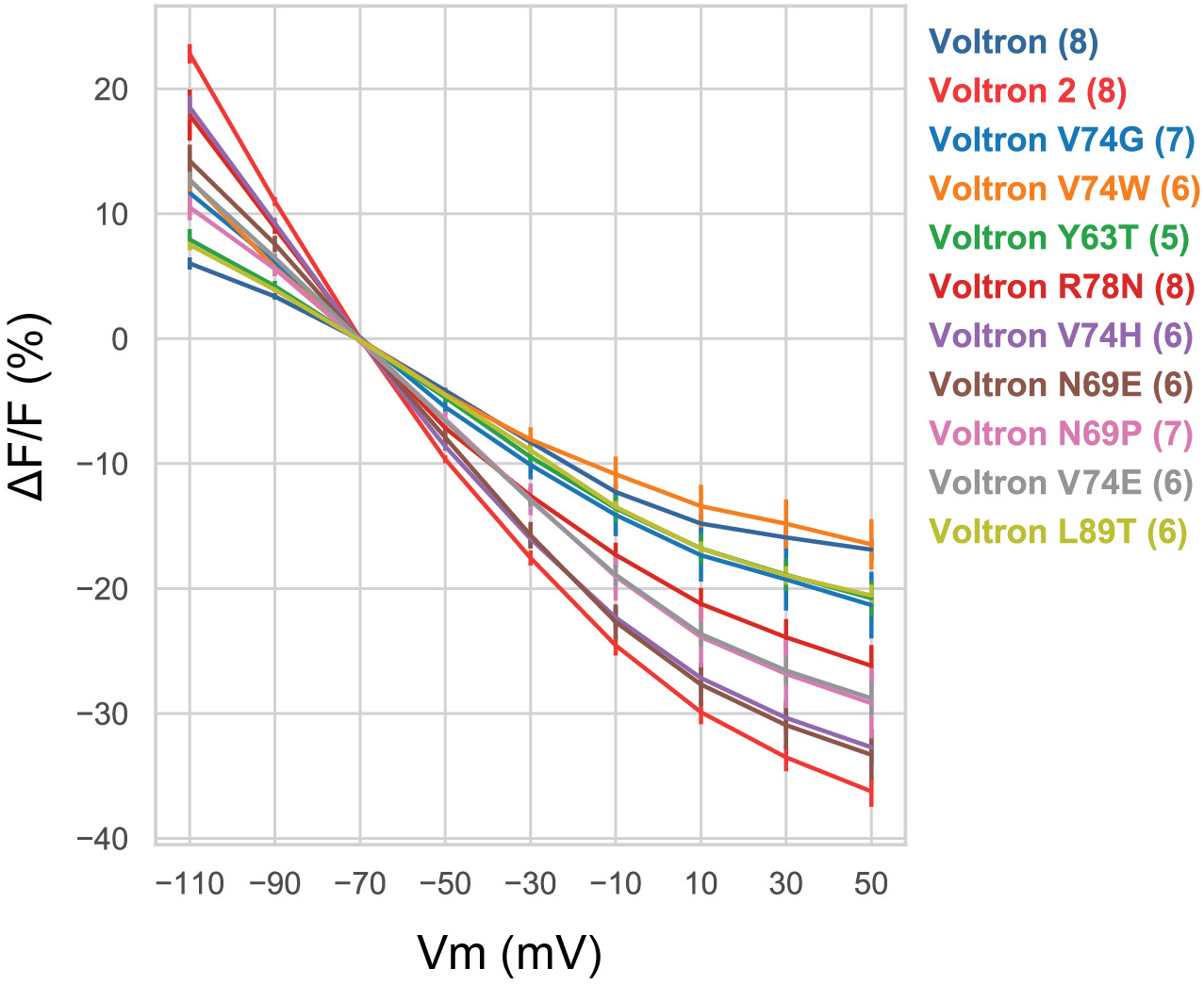
Peak fluorescence response to voltage steps from −70 mV of the top variants from the field stimulation assay, with Voltron and Voltron2 traces (reproduced from Fig. 2) superimposed for reference. Number in parentheses indicates the number of neurons assayed. All sensor mutants were conjugated to JF_525_ dyes for these experiments (Voltron_525_).

**Supplementary Figure 4:**
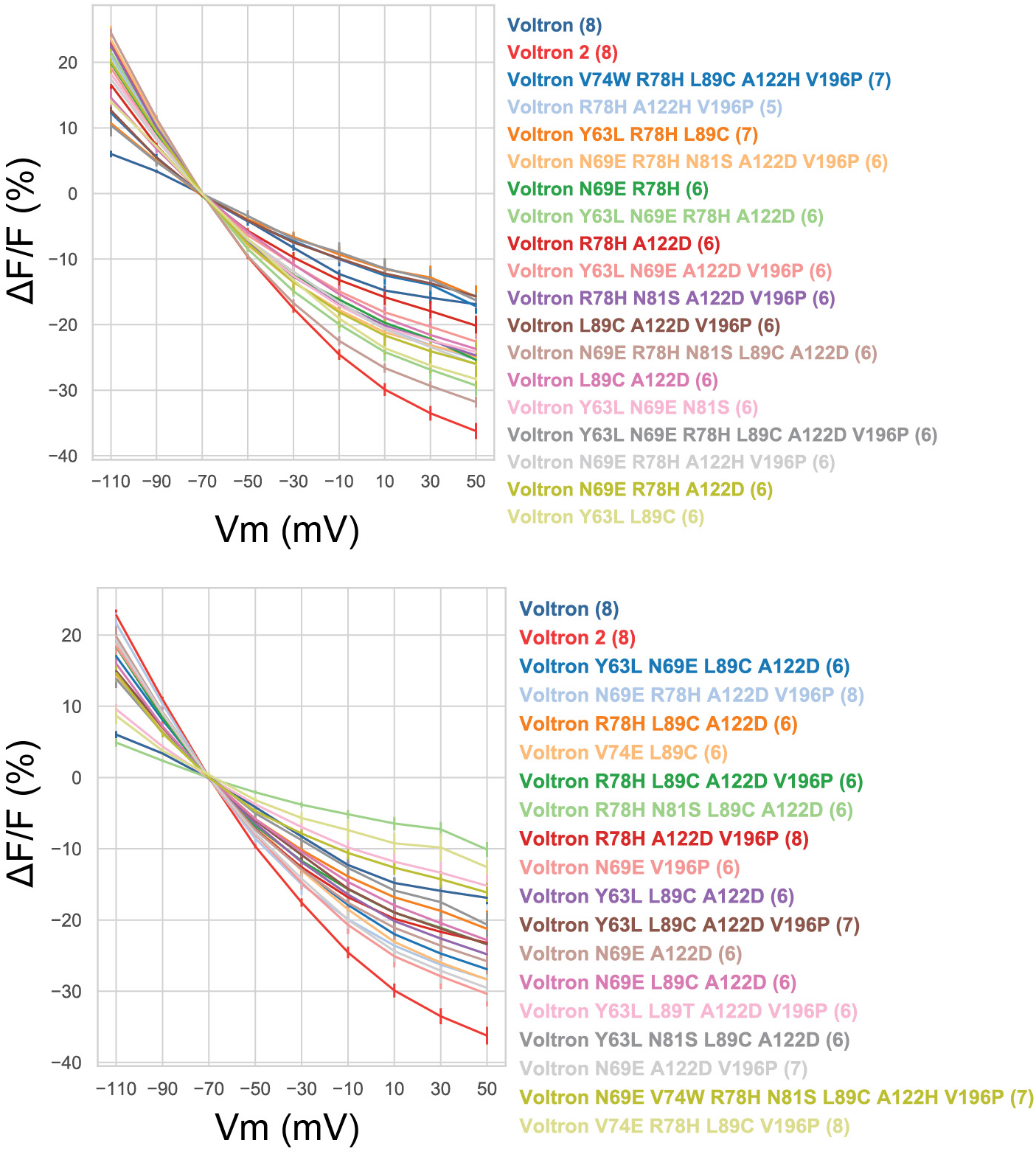
Peak fluorescence response to voltage steps of Voltron, Voltron2 (reproduced from Fig. 2) and the top-performing combo variants from the field stimulation assay. For clarity, variants are divided into two panels. Number in parentheses indicates the number of neurons assayed. All sensor mutants were conjugated to JF_525_ dyes for these experiments (Voltron_525_).

**Supplementary Figure 5:**
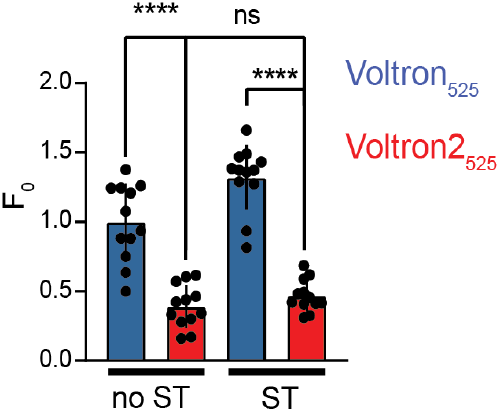
Baseline fluorescence of non soma-tagged (no ST) and soma-tagged (ST) Voltron_525_ and Voltron2_525_ (n=12 neurons for each, from single transfection; error bars: s.d.; ****: P<0.0001; n.s.: P=0.80; one-way ANOVA followed by Tukey’s multiple comparison test). See Supplementary Figs. 6, 7 for cell images.

**Supplementary Figure 6.**
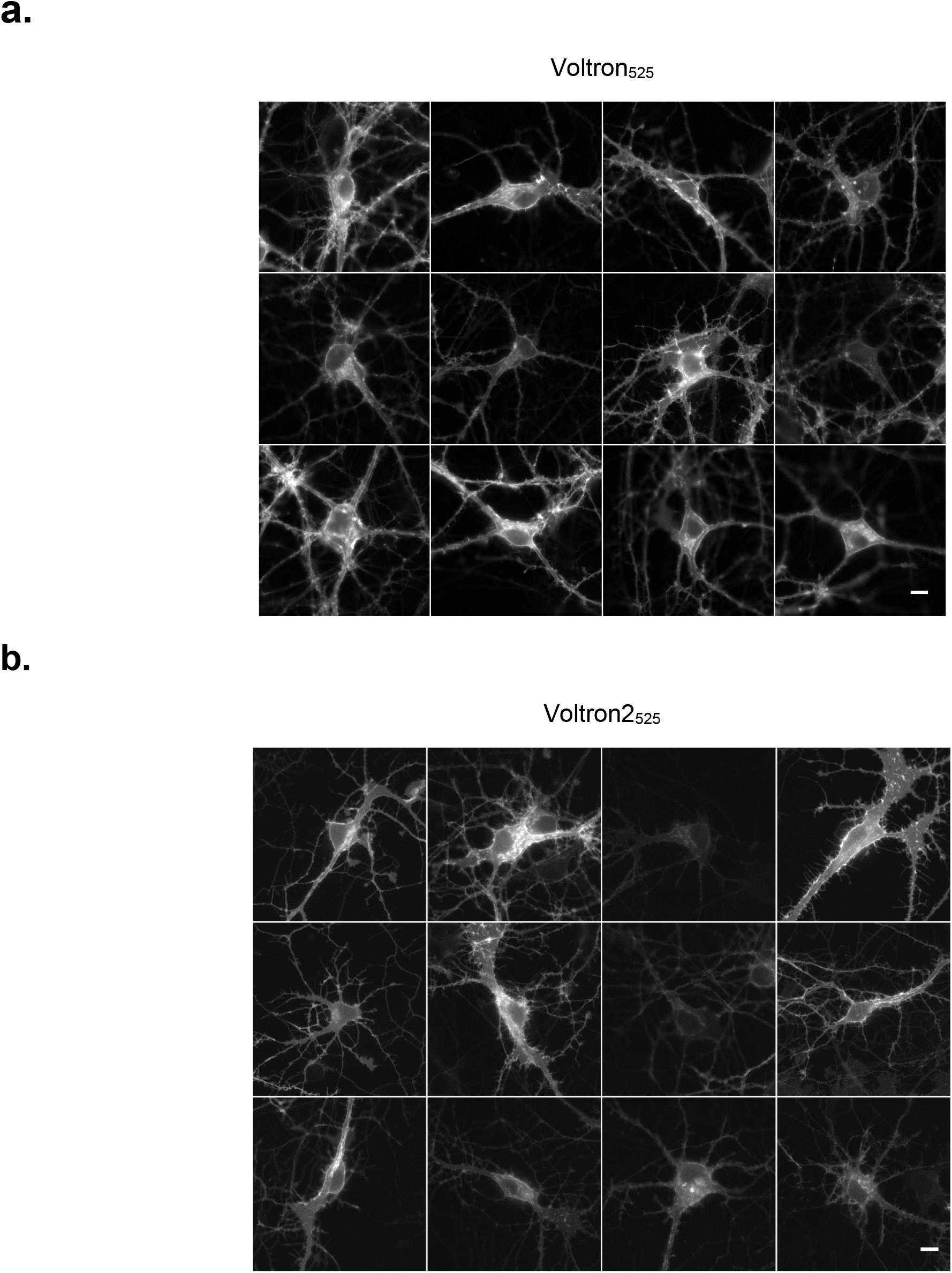
Representative fluorescent images of cultured hippocampal neurons expressing (non-soma tagged) Voltron_525_ and Voltron2_525_. Images were taken with 15 mW/mm^2^ light power and 30 ms exposure time. Dynamic ranges of images were rescaled for clarity. For quantitative comparison of fluorescence intensities, see Fig. 2h. Scale bar: 10 μm

**Supplementary Figure 7.**
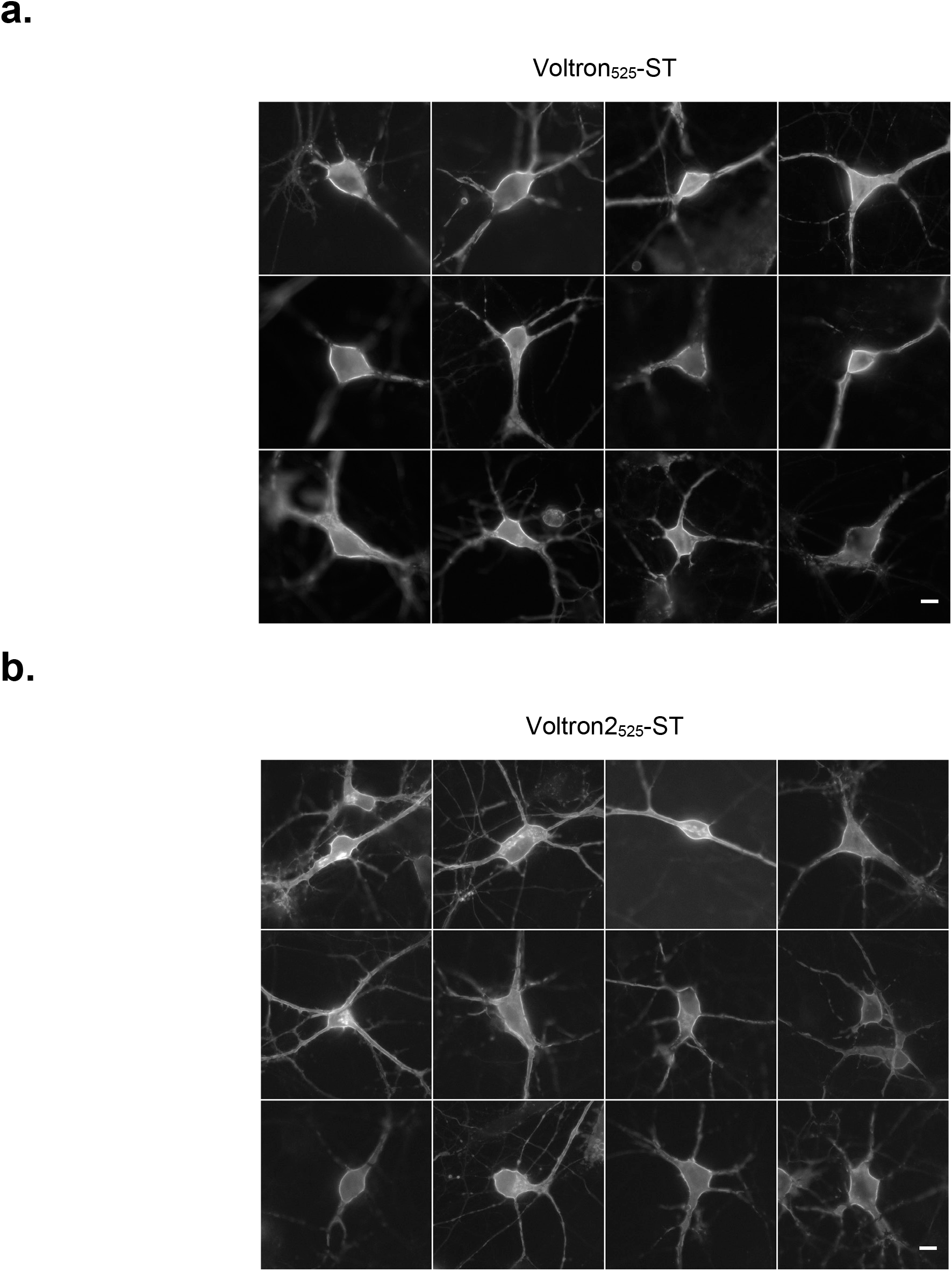
Representative fluorescent images of cultured hippocampal neurons expressing soma-tagged Voltron_525_ and Voltron2_525_. Images were taken with 15 mW/mm^2^ light power and 30 ms exposure time. Dynamic ranges of images were rescaled for clarity. For quantitative comparison of fluorescence intensities, see Fig. 2h. Scale bar: 10 μm

**Supplementary Figure 8:**
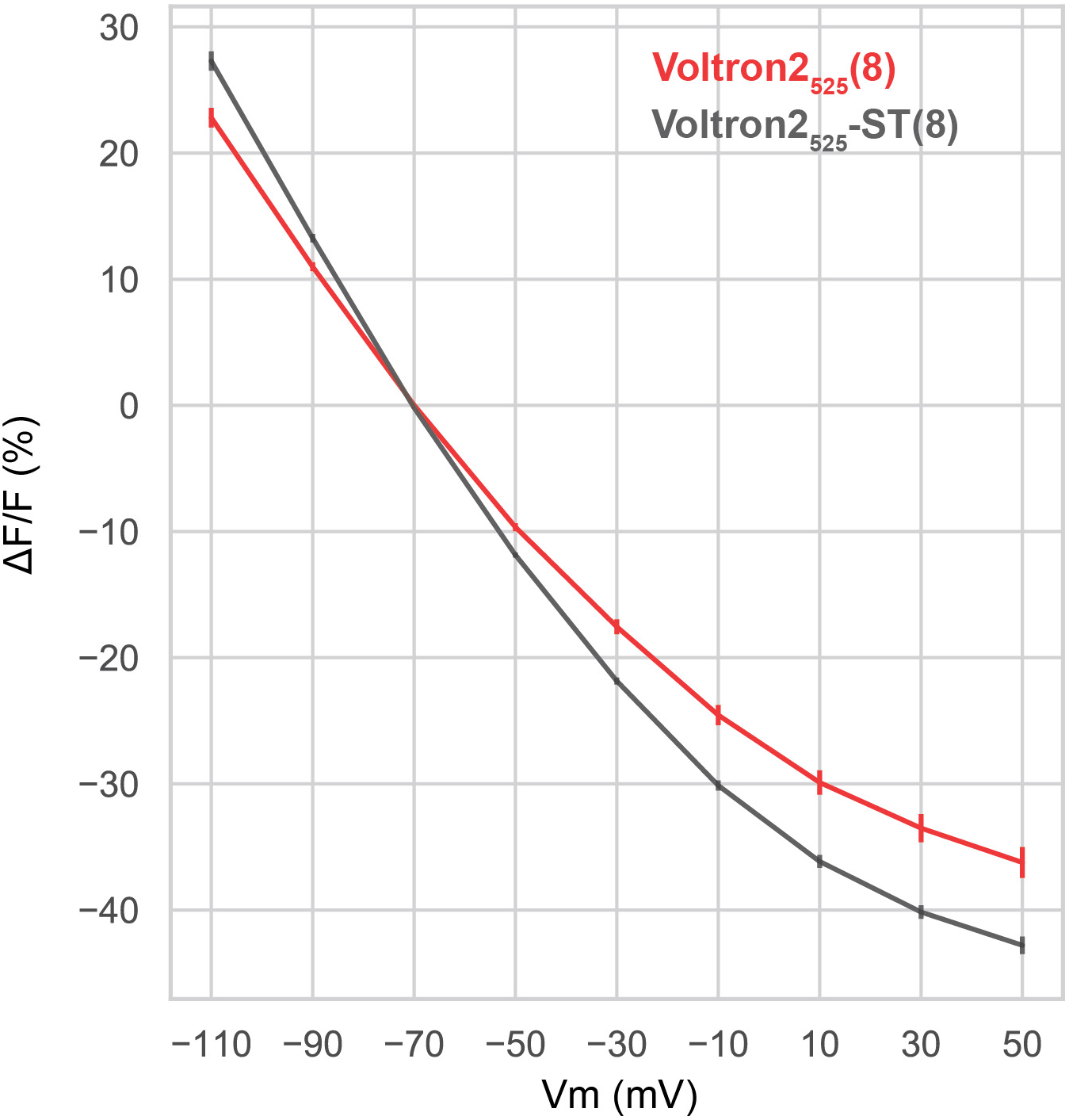
Fluorescence response to voltage steps of Voltron2_525_-ST, compared to Voltron2_525_ (reproduced from Fig. 2). Number in parentheses indicates the number of neurons assayed.

**Supplementary Figure 9:**
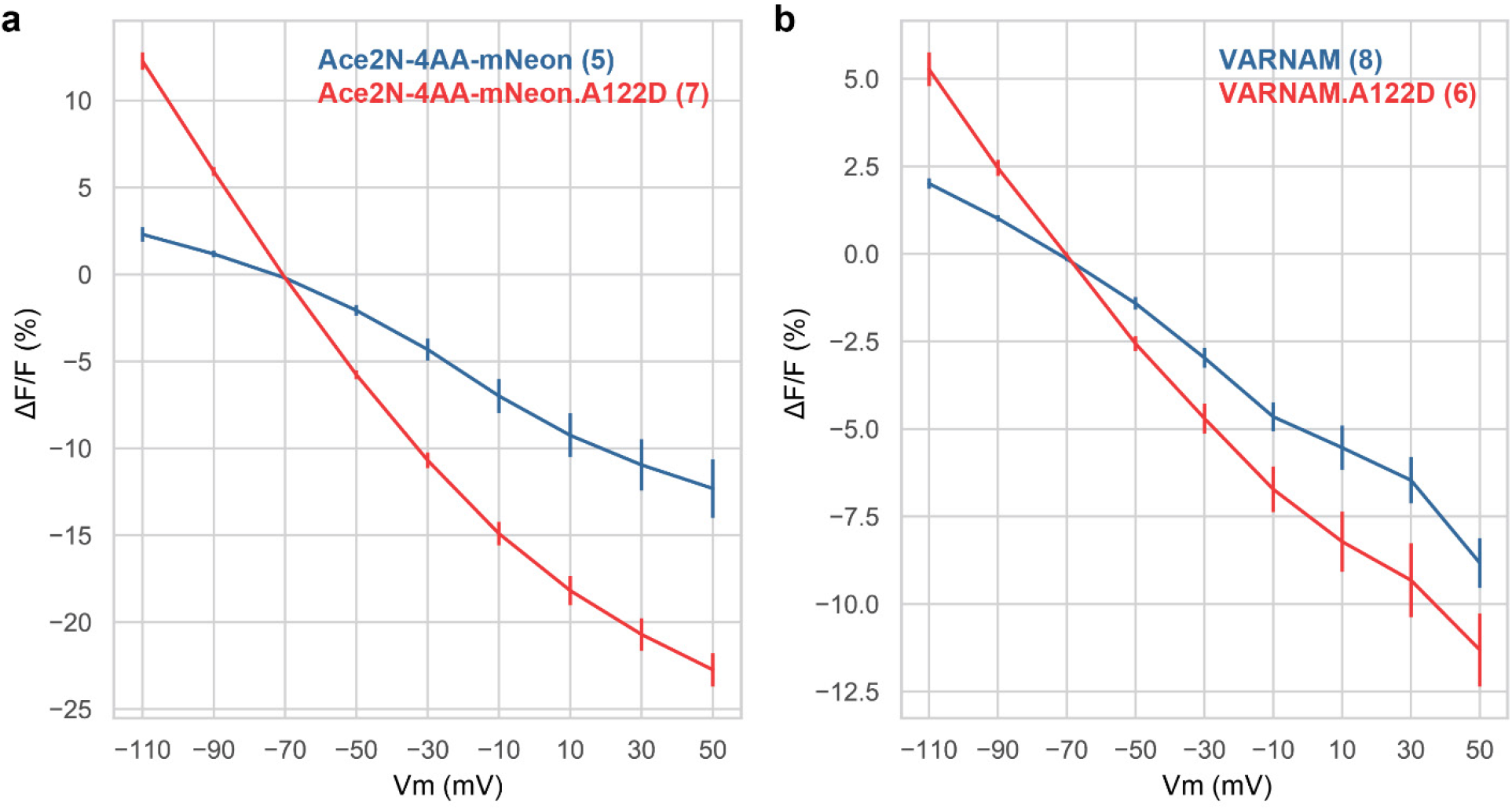
Increased voltage sensitivities of Ace-based FRET opsin indicators. a. Peak fluorescence response to voltage steps from −70 mV of Ace-4AA-mNeon.A122D, with Ace-4AA-mNeon as control. b. Peak fluorescence response to voltage steps from −70 mV of VARNAM.A122D, with VARNAM as control. Number in parentheses indicates the number of neurons assayed.

**Supplementary Figure 10:**
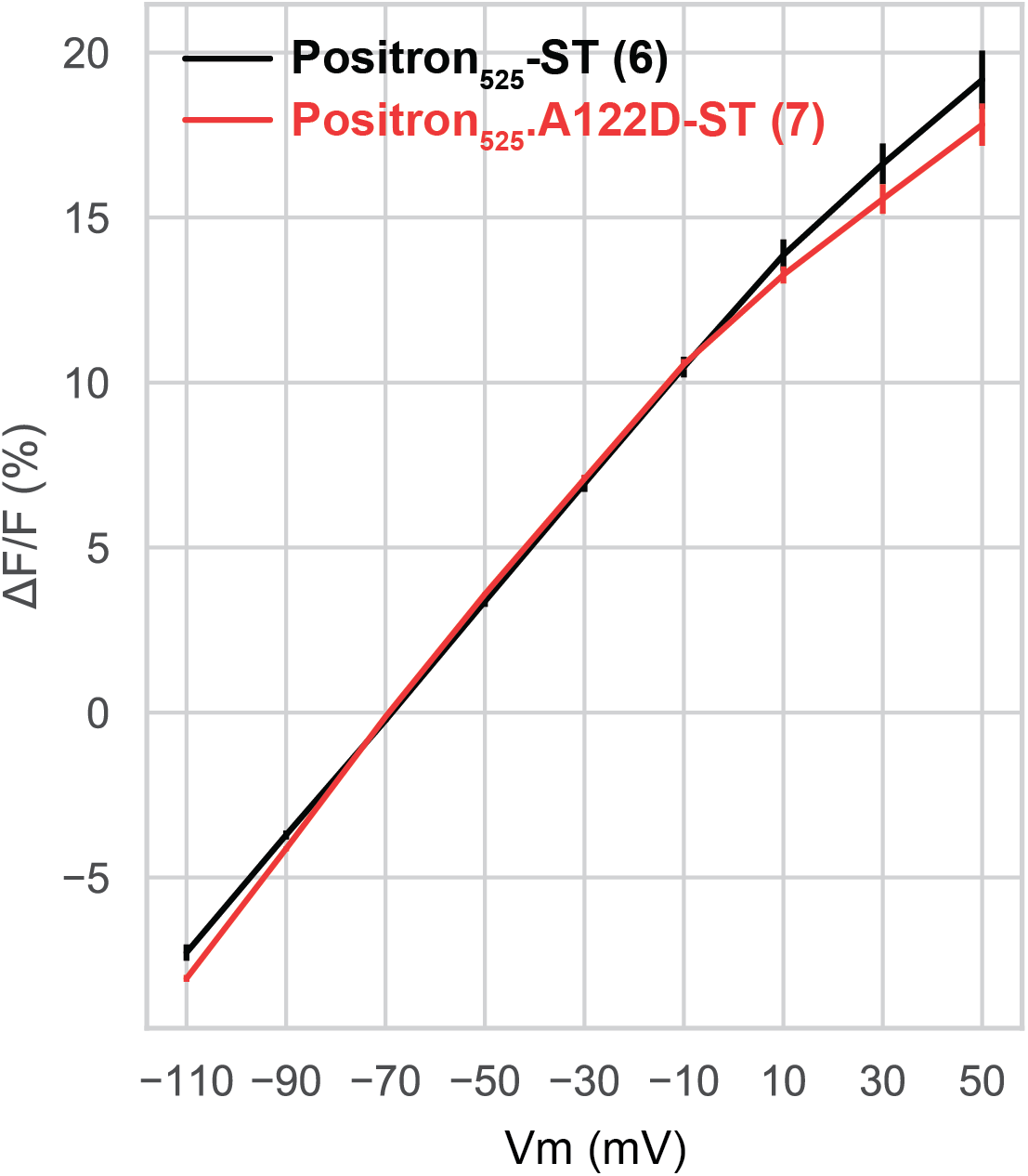
Fluorescence response to voltage steps of Positron_525_-ST, compared to Positron_525_.A122D-ST. Number in parentheses indicates the number of neurons assayed.

**Supplementary Figure 11:**
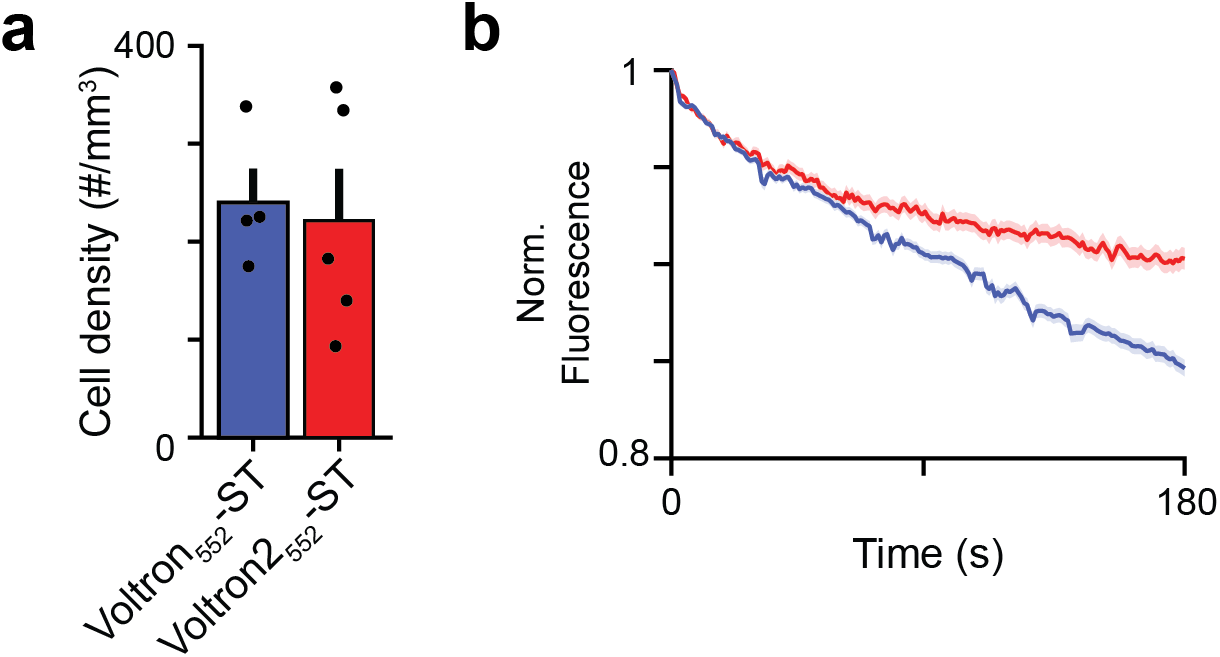
a. Density of visually identifiable neurons in mouse CA1. b. Photobleaching comparison of Voltron_552_ and Voltron2_552_ in mouse CA1 (solid color: mean, shading: SEM).

## Tables

**Supplementary Table 1**: Screening results of field stimulation assay on Voltron point mutants. Normalization performed to in-plate Voltron controls.

**Supplementary Table 2**: Screening results of field stimulation assay on Voltron combo mutants. Normalization performed to in-plate Voltron controls.

**Supplementary Table 3**: Combo variants containing the A122D mutation, arranged by the number of mutations. Data aggregated from Supplementary Table 2.

**Supplementary Table 4**: Custom primers used for library tagmentation and NextSeq sequencing

