## Supplementary Table 1 for "Sensitivity optimization of a rhodopsin-based fluorescent voltage indicator"

| mutation | number of plates | ΔF/F0 (norm.) | ΔF/F0 | ΔF/F0 p value | SNR (norm.) | SNR p value | τ_ON (norm.) | τ_ON p value | τ_OFF (norm.) | τ_OFF p value | F0 (norm.) | F0 p value |
|---|---|---|---|---|---|---|---|---|---|---|---|---|
| V74G | 4 | 2.281662711 | 0.094325 | 2.42E-288 | 1.255663131 | 0.13904313 | 1.017718924 | 0.00130094 | 0.976170287 | 0.004550585 | 0.522177868 | 3.22E-159 |
| V74W | 3 | 2.212789329 | 0.101369 | 1.25E-189 | 1.505708746 | 1.17E-54 | 0.767194753 | 1.28E-11 | 0.799445171 | 4.71E-20 | 0.558297083 | 3.61E-60 |
| A122D | 2 | 2.178792922 | 0.116971 | 1.29E-135 | 1.67525836 | 3.56E-18 | 1.173881953 | 1.09E-10 | 0.989130414 | 0.191438673 | 0.768030879 | 1.71E-15 |
| Y63T | 5 | 2.166666682 | 0.087511 | 5.05E-290 | 1.363397844 | 1.32E-16 | 0.862791844 | 6.93E-09 | 0.877426553 | 2.60E-18 | 0.49982621 | 8.04E-193 |
| V74E | 3 | 2.096680021 | 0.107978 | 1.59E-177 | 1.441941157 | 0.001528438 | 1.168394509 | 1.25E-18 | 0.985231637 | 0.117843541 | 0.553743219 | 3.68E-65 |
| R78N | 1 | 2.096419182 | 0.112549 | 1.82E-154 | 1.491174531 | 8.11E-100 | 0.721690737 | 1.27E-30 | 0.580064675 | 1.07E-63 | 0.645140527 | 8.82E-60 |
| N69E | 2 | 2.077516474 | 0.119531 | 2.43E-45 | 1.760987816 | 3.29E-44 | 1.178091231 | 1.03E-08 | 1.007128376 | 0.000787312 | 0.875871948 | 8.21E-31 |
| N69P | 7 | 2.07320852 | 0.118121 | 3.77E-51 | 1.747110185 | 1.17E-07 | 1.018631313 | 0.163663712 | 0.875559343 | 0.000105087 | 0.903349402 | 1.65E-05 |
| L89T | 2 | 2.033093881 | 0.123187 | 2.15E-135 | 1.79936369 | 1.36E-123 | 0.623310416 | 2.07E-53 | 0.628057192 | 3.95E-114 | 0.99225264 | 0.003127228 |
| V74H | 1 | 2.013548026 | 0.083241 | 0 | 1.236237778 | 7.26E-80 | 0.961781628 | 0.000463999 | 0.89507729 | 1.03E-10 | 0.511081815 | 2.75E-162 |
| M68A | 2 | 1.993435748 | 0.123191 | 4.65E-294 | 1.681092847 | 4.14E-22 | 1.06341669 | 0.001076863 | 1.025923833 | 0.077833564 | 0.844429488 | 2.82E-26 |
| W178N | 2 | 1.992631006 | 0.093149 | 6.22E-55 | 1.332586752 | 0.694144856 | 0.615840343 | 3.89E-33 | 0.681635312 | 1.27E-26 | 0.634812241 | 1.32E-42 |
| V74P | 1 | 1.989688471 | 0.091149 | 1.19E-130 | 1.162588803 | 1.07E-12 | 0.843507713 | 0.051727171 | 0.92034659 | 0.00335364 | 0.430989939 | 5.94E-93 |
| F134V | 1 | 1.986932132 | 0.132004 | 1.30E-28 | 1.553301098 | 8.36E-14 | 0.362407307 | 2.39E-20 | 0.7424208 | 0.000634797 | 0.661965144 | 5.00E-11 |
| M68H | 3 | 1.979262855 | 0.122243 | 2.80E-244 | 1.782287382 | 1.77E-94 | 1.052180581 | 0.019203413 | 0.955721024 | 0.052952118 | 0.950785669 | 0.000171188 |
| Y63I | 4 | 1.978895101 | 0.08943 | 3.11E-85 | 1.157947208 | 2.85E-08 | 0.942926319 | 0.001886572 | 0.997428879 | 0.007399062 | 0.458095106 | 4.02E-68 |
| V74F | 3 | 1.973771918 | 0.098315 | 1.17E-145 | 1.314095315 | 2.09E-12 | 1.001091143 | 0.000212733 | 1.165272081 | 5.19E-10 | 0.609907208 | 1.00E-42 |
| V74A | 4 | 1.97220386 | 0.081532 | 2.75E-244 | 1.203466816 | 7.34E-12 | 1.024185922 | 0.005445279 | 0.928209168 | 1.91E-06 | 0.538700882 | 1.64E-121 |
| R72N | 2 | 1.964016131 | 0.097631 | 1.02E-70 | 1.081991065 | 0.014987314 | 0.811251243 | 0.000164937 | 1.125095072 | 0.013523871 | 0.440155902 | 7.92E-87 |
| V74K | 1 | 1.950252067 | 0.089342 | 2.27E-157 | 1.168447042 | 3.61E-20 | 1.092425552 | 7.65E-08 | 1.126249871 | 0.004196855 | 0.4725639 | 4.12E-87 |
| L89A | 6 | 1.936264333 | 0.127973 | 1.73E-122 | 2.021693943 | 7.03E-11 | 0.561451283 | 2.45E-56 | 0.639813686 | 3.58E-99 | 1.066353137 | 7.75E-22 |
| V74Q | 3 | 1.933996037 | 0.0823 | 3.70E-276 | 1.183123441 | 1.91E-21 | 1.135097522 | 1.09E-10 | 1.005729623 | 5.84E-07 | 0.56451326 | 8.83E-162 |
| Q64P | 4 | 1.930694816 | 0.113959 | 9.12E-65 | 1.699691114 | 0.310926504 | 1.084445901 | 1.12E-05 | 0.88487717 | 2.50E-08 | 0.770049075 | 4.90E-41 |
| V196P | 2 | 1.912534078 | 0.111008 | 9.78E-95 | 1.691625804 | 2.95E-09 | 1.057852525 | 0.009586505 | 0.888103341 | 0.010490811 | 0.755837235 | 6.36E-21 |
| Y63V | 3 | 1.911922976 | 0.099151 | 6.71E-118 | 1.332817607 | 0.47547728 | 1.045037389 | 3.22E-08 | 0.915577092 | 5.29E-05 | 0.62886216 | 1.79E-109 |
| R72V | 4 | 1.911328939 | 0.097024 | 6.70E-53 | 1.112694104 | 0.376816854 | 0.964330219 | 0.224926369 | 0.98848609 | 0.133860454 | 0.452080067 | 3.46E-76 |
| Y63M | 4 | 1.901083179 | 0.072558 | 1.31E-92 | 1.272028718 | 0.008726477 | 0.807934679 | 9.96E-08 | 0.867942763 | 0.000283117 | 0.53448162 | 4.90E-63 |
| R72Y | 2 | 1.897153993 | 0.096305 | 5.07E-84 | 1.306980169 | 0.000834114 | 0.885261063 | 0.011872033 | 0.918478664 | 0.398268499 | 0.507361472 | 1.20E-92 |
| V74D | 3 | 1.890096445 | 0.085068 | 7.22E-114 | 1.111385791 | 0.442916439 | 0.977401444 | 3.91E-08 | 1.164938395 | 4.59E-05 | 0.438920967 | 3.63E-100 |
| V74S | 2 | 1.889011938 | 0.086537 | 2.50E-13 | 1.213179182 | 0.537423389 | 1.043995859 | 0.292536539 | 1.257189055 | 0.456149326 | 0.601933101 | 5.05E-18 |
| R72H | 4 | 1.882877795 | 0.093598 | 1.24E-39 | 1.238216447 | 0.607438196 | 0.925957902 | 2.32E-10 | 1.007441554 | 4.27E-09 | 0.530464433 | 1.36E-44 |
| A55T | 3 | 1.88227318 | 0.091866 | 6.80E-32 | 1.745512468 | 1.14E-21 | 0.937907458 | 0.000726726 | 0.76804406 | 1.02E-15 | 1.197070896 | 4.73E-06 |
| R72G | 9 | 1.87952388 | 0.093431 | 2.47E-88 | 1.195391873 | 0.329764022 | 0.713525367 | 6.92E-15 | 0.78752755 | 1.35E-12 | 0.455700342 | 2.70E-74 |
| M68S | 2 | 1.868416756 | 0.115397 | 691904177745e | 1.957661321 | 0 | 1.045036487 | 0.220519837 | 0.951870182 | 0.429777569 | 1.13314863 | 0.176246141 |
| M68T | 4 | 1.857979136 | 0.103537 | 2.99E-62 | 1.611166333 | 2.05E-12 | 1.07627614 | 0.001932307 | 0.954999466 | 0.020837191 | 1.058304525 | 0.223929372 |
| Y63K | 1 | 1.84721992 | 0.070502 | 7.43E-143 | 1.267739176 | 1.03E-68 | 0.784657599 | 3.59E-18 | 0.675121502 | 3.70E-42 | 0.534559899 | 1.91E-68 |
| M68R | 5 | 1.839434788 | 0.113426 | 5.43E-180 | 1.823235095 | 4.31E-17 | 1.065774378 | 0.003456629 | 0.954259754 | 0.053218764 | 1.058696983 | 0.000191024 |
| M68E | 1 | 1.83703559 | 0.113459 | 0 | 1.455331177 | 4.51E-268 | 1.038709782 | 0.108375163 | 0.990355722 | 0.453727029 | 0.813598582 | 4.42E-45 |
| Y63L | 8 | 1.835920447 | 0.079694 | 1.78E-175 | 1.243897368 | 0.261148697 | 0.874779885 | 1.42E-15 | 0.896602559 | 1.54E-11 | 0.510295846 | 8.76E-97 |
| Y63P | 4 | 1.834234329 | 0.085054 | 4.59E-239 | 1.357444687 | 4.23E-13 | 0.878380154 | 5.90E-10 | 0.897584447 | 1.07E-08 | 0.518470402 | 4.10E-67 |
| Y63D | 1 | 1.81963732 | 0.069449 | 3.45E-138 | 1.234837569 | 1.75E-56 | 0.725966443 | 4.10E-26 | 0.692746922 | 1.56E-29 | 0.500875535 | 2.03E-81 |
| R78H | 4 | 1.815423027 | 0.097463 | 5.08E-124 | 1.357901622 | 0.002363214 | 1.036830999 | 1.12E-05 | 1.183023304 | 9.89E-08 | 0.741310571 | 3.26E-26 |
| L93V | 5 | 1.808195962 | 0.087719 | 2.21E-14 | 1.017034673 | 0.782801311 | 0.942346567 | 1.69E-08 | 1.116352122 | 0.322725527 | 0.767640506 | 5.05E-11 |
| L89G | 6 | 1.805913471 | 0.116545 | 2.63E-74 | 1.751866683 | 5.19E-15 | 0.445827941 | 5.66E-50 | 0.619904609 | 3.77E-55 | 0.802213609 | 2.40E-20 |
| R72K | 1 | 1.79979589 | 0.089468 | 9.40E-289 | 1.16293655 | 2.39E-38 | 0.848844432 | 1.58E-14 | 0.825857925 | 1.01E-21 | 0.454813285 | 5.33E-224 |
| R72D | 2 | 1.793259575 | 0.089143 | 5.39E-104 | 1.032719082 | 0.115763137 | 0.881176123 | 0.000593102 | 0.891002821 | 7.31E-05 | 0.420238054 | 1.53E-102 |
| Y63H | 2 | 1.788949071 | 0.0697 | 1.11E-128 | 1.293981237 | 2.38E-73 | 0.811988752 | 6.97E-17 | 0.801305628 | 1.28E-22 | 0.598202579 | 1.69E-49 |
| Q64Y | 6 | 1.781407366 | 0.095293 | 1.98E-63 | 1.291272321 | 0.746372486 | 0.946908524 | 1.78E-05 | 1.069636065 | 6.83E-05 | 0.694357746 | 1.89E-68 |
| Y63C | 3 | 1.771501082 | 0.067612 | 2.52E-109 | 1.194612882 | 0.971735564 | 0.763845166 | 1.59E-08 | 0.800721729 | 0.004260341 | 0.522417013 | 4.80E-63 |
| Y63S | 4 | 1.76366487 | 0.088616 | 2.64E-121 | 1.270602637 | 0.003623248 | 1.010825065 | 0.041264191 | 0.865189867 | 7.03E-17 | 0.561714743 | 5.66E-63 |
| Y63R | 2 | 1.75246746 | 0.108604 | 1.55E-59 | 1.19570492 | 0.174102224 | 1.121530612 | 1.25E-05 | 0.874041958 | 1.32E-10 | 0.513344005 | 4.30E-80 |
| L174I | 5 | 1.749537338 | 0.09943 | 2.52E-60 | 1.539988393 | 3.49E-11 | 0.567393591 | 1.70E-38 | 0.579722393 | 1.52E-64 | 0.83135039 | 2.57E-08 |
| S189D | 2 | 1.748328348 | 0.088201 | 1.16E-60 | 1.314684616 | 2.72E-18 | 1.187404764 | 2.65E-13 | 0.861920436 | 2.00E-12 | 0.779658998 | 8.26E-32 |
| R72T | 1 | 1.72936881 | 0.085967 | 1.36E-254 | 1.11158986 | 4.11E-18 | 0.975652083 | 0.274992953 | 1.1378463 | 2.40E-08 | 0.490601917 | 1.10E-192 |
| A55E | 1 | 1.724089696 | 0.099197 | 0.017721442 | 1.529333267 | 8.26E-14 | 1.377842844 | 1.15E-07 | 1.391018031 | 3.71E-08 | 1.256186582 | 0.376589528 |
| V74L | 8 | 1.718453914 | 0.092136 | 1.43E-62 | 1.172166457 | 0.865744792 | 1.191271263 | 1.44E-10 | 1.10334709 | 8.16E-05 | 0.656443368 | 2.69E-50 |
| M68G | 2 | 1.718249998 | 0.106123 | 0 | 1.525499405 | 2.59E-214 | 1.029212218 | 0.001935045 | 0.908418464 | 0.038002188 | 0.868753089 | 1.18E-06 |
| R72A | 1 | 1.713286895 | 0.085168 | 4.82E-225 | 1.075694802 | 2.75E-10 | 0.929336811 | 0.018527773 | 1.228240342 | 3.24E-10 | 0.468311387 | 5.82E-205 |
| V74Y | 2 | 1.712644314 | 0.082243 | 1.20E-89 | 1.201510248 | 4.34E-08 | 0.865188813 | 0.007373032 | 1.026872659 | 0.49167799 | 0.548817765 | 5.63E-80 |
| Q64I | 2 | 1.705043327 | 0.074715 | 4.27E-117 | 1.130386174 | 6.16E-12 | 1.059936774 | 0.023921181 | 1.187554589 | 7.66E-07 | 0.538605028 | 2.72E-83 |
| N69H | 4 | 1.701080237 | 0.096919 | 1.07E-117 | 1.585541727 | 4.05E-23 | 1.094603555 | 0.000159317 | 0.928138575 | 0.001198578 | 0.891598347 | 8.40E-06 |
| Y63Q | 4 | 1.69770539 | 0.076722 | 824255071099e | 1.115000777 | 9.09E-18 | 0.953417252 | 6.19E-06 | 0.888173501 | 1.90E-07 | 0.536099326 | 4.53E-230 |
| R72P | 2 | 1.68900384 | 0.085738 | 7.48E-164 | 1.264096751 | 8.91E-43 | 0.849276841 | 0.125898975 | 0.804108849 | 1.08E-08 | 0.644006319 | 2.76E-67 |
| Q64F | 5 | 1.6877532 | 0.095803 | 2.10E-59 | 1.265661029 | 0.414647496 | 1.107785973 | 1.78E-07 | 1.015467321 | 9.94E-08 | 0.695207169 | 2.16E-60 |
| R72M | 3 | 1.686048775 | 0.085588 | 1.43E-75 | 1.065448784 | 0.503509711 | 0.815991356 | 1.46E-05 | 0.843331057 | 0.001174092 | 0.459685345 | 4.18E-73 |
| R72S | 1 | 1.681424351 | 0.083584 | 2.03E-215 | 1.117081104 | 6.53E-23 | 0.865306421 | 9.00E-09 | 1.05778518 | 0.004065483 | 0.486720154 | 7.08E-190 |
| L89C | 4 | 1.678757391 | 0.102512 | 2.32E-51 | 1.895956739 | 4.91E-09 | 0.602105287 | 1.59E-78 | 0.674787643 | 4.61E-65 | 1.231987361 | 8.58E-58 |
| N69Q | 3 | 1.67372949 | 0.095361 | 2.76E-25 | 1.634093007 | 6.88E-12 | 1.177437583 | 1.90E-05 | 1.002906407 | 0.005807551 | 1.108891785 | 0.026507433 |
| Y63N | 2 | 1.663205732 | 0.072197 | 1.89E-187 | 1.320708744 | 1.38E-14 | 0.940937707 | 0.483251211 | 0.980914364 | 0.00943919 | 0.654017653 | 8.26E-44 |
| V74M | 2 | 1.660752744 | 0.078304 | 1.43E-09 | 1.103767269 | 2.79E-06 | 1.105490283 | 3.33E-06 | 1.118274135 | 3.53E-07 | 0.607857272 | 1.51E-44 |
| M68Q | 1 | 1.658964892 | 0.102461 | 0 | 1.529877319 | 0 | 1.064185616 | 0.000191691 | 0.963401941 | 0.069299331 | 1.00768508 | 0.97281745 |
| Q64G | 2 | 1.65007323 | 0.097395 | 2.07E-30 | 1.225667684 | 4.82E-06 | 0.994737519 | 2.43E-13 | 1.019457967 | 4.22E-08 | 0.883767336 | 0.017102961 |
| V196T | 3 | 1.620609518 | 0.082599 | 6.95E-48 | 1.536311192 | 0.988434226 | 1.080440346 | 0.051970031 | 0.990272818 | 0.151271312 | 1.105439262 | 0.000904794 |
| A122C | 1 | 1.616539862 | 0.073647 | 3.93E-10 | 1.200847558 | 2.14E-06 | 1.112453297 | 0.171032342 | 1.088807166 | 0.290222887 | 0.873608989 | 0.172617713 |
| N81S | 3 | 1.614923595 | 0.073573 | 3.03E-26 | 1.574426883 | 0.000113833 | 1.202103947 | 0.041599544 | 0.99254556 | 0.51120255 | 1.036257033 | 0.095271512 |
| D65P | 2 | 1.614896374 | 0.098409 | 4.87E-60 | 1.436210557 | 0.085390472 | 1.076608162 | 8.84E-05 | 0.919749705 | 5.93E-06 | 0.823666355 | 3.07E-18 |
| Y63A | 5 | 1.608889311 | 0.061406 | 1.19E-39 | 1.275725893 | 7.34E-06 | 0.92706624 | 9.14E-06 | 0.991684134 | 0.003315678 | 0.775762878 | 1.74E-12 |
| V74R | 2 | 1.603031862 | 0.076979 | 2.84E-49 | 0.996646367 | 0.252019926 | 1.042864698 | 0.55829353 | 1.058368436 | 0.274334131 | 0.418363563 | 7.08E-61 |
| R72W | 1 | 1.575777087 | 0.078332 | 2.07E-80 | 0.965997181 | 0.211447067 | 0.915078352 | 0.695651784 | 1.205332452 | 3.05E-05 | 0.481188179 | 6.69E-90 |
| Y63G | 3 | 1.573385096 | 0.072976 | 0 | 1.25035293 | 1.98E-79 | 0.958554998 | 4.77E-07 | 0.952711482 | 3.80E-05 | 0.690654472 | 4.51E-120 |
| S189C | 4 | 1.567942146 | 0.080675 | 7.35E-109 | 1.020082645 | 0.059902008 | 1.293798483 | 1.22E-19 | 1.089943683 | 0.004249941 | 0.56462011 | 4.82E-96 |

| | | | | | | | | | | | | |
|---|---|---|---|---|---|---|---|---|---|---|---|---|
| N69G | 1 | 1.567333391 | 0.099517 | 2.09E-12 | 1.319325222 | 1.49E-26 | 1.081903954 | 0.115314951 | 0.969205621 | 0.01452269 | 0.873939435 | 0.000893502 |
| V74T | 4 | 1.559783123 | 0.074902 | 5.56E-66 | 1.091739143 | 0.000654881 | 0.976812681 | 1.15E-06 | 1.251255322 | 1.50E-07 | 0.554168102 | 1.14E-29 |
| A122H | 4 | 1.542990521 | 0.070296 | 1.78E-13 | 1.247899167 | 0.125281139 | 1.138180211 | 0.042943259 | 1.319336441 | 0.001637182 | 0.927154584 | 0.093116777 |
| L88G | 1 | 1.532177562 | 0.117928 | 0.000188703 | 1.17276382 | 0.000168931 | 0.74275241 | 0.039918128 | 1.055067915 | 1.17E-15 | 0.681166532 | 3.56E-15 |
| L174G | 3 | 1.522202538 | 0.078723 | 0.01693375 | 1.275555845 | 0.483101606 | 0.556818008 | 9.05E-20 | 0.644552496 | 5.29E-21 | 1.082647636 | 0.000360574 |
| V74V | 1 | 1.518331616 | 0.062768 | 9.57E-136 | 1.193077967 | 1.31E-46 | 0.786435255 | 6.58E-29 | 0.854953375 | 2.63E-15 | 0.706850516 | 7.91E-36 |
| W178Q | 2 | 1.517439865 | 0.094635 | 4.23E-06 | 1.254355054 | 0.08686103 | 0.618605569 | 4.22E-12 | 0.791297265 | 1.41E-08 | 0.850288141 | 0.025923295 |
| A55C | 2 | 1.51079815 | 0.078027 | 7.93E-54 | 1.49349831 | 1.16E-05 | 0.887459271 | 9.84E-12 | 0.757344693 | 6.51E-36 | 1.083698932 | 0.016135662 |
| A122L | 1 | 1.510403154 | 0.116252 | 0.1066363 | 1.203961716 | 9.50E-32 | 0.972385121 | 8.05E-15 | 1.166296067 | 3.34E-28 | 0.68416869 | 1.25E-26 |
| V196I | 1 | 1.506778359 | 0.079807 | 3.35E-55 | 1.363321911 | 9.99E-27 | 1.081603903 | 0.000682607 | 0.897465092 | 0.00119252 | 1.023122383 | 0.07660244 |
| Q64V | 5 | 1.505646957 | 0.089787 | 3.40E-24 | 1.131347119 | 0.000640124 | 1.013065456 | 5.56E-09 | 1.085237758 | 0.006033046 | 0.645207205 | 1.53E-39 |
| V196L | 1 | 1.50481553 | 0.09119 | 1.12E-62 | 1.483070908 | 4.41E-30 | 0.986769648 | 0.091876977 | 0.834698425 | 0.000125522 | 0.940702339 | 0.00040623 |
| W178C | 2 | 1.503228079 | 0.07648 | 0.000376839 | 1.129885906 | 0.362331015 | 0.489223136 | 4.91E-11 | 0.794111757 | 0.004621173 | 0.683226481 | 1.68E-06 |
| S189E | 2 | 1.496951383 | 0.084113 | 2.76E-75 | 1.406435886 | 4.57E-98 | 1.207211931 | 5.96E-07 | 1.002903816 | 0.00100883 | 1.182064468 | 2.42E-22 |
| G70P | 2 | 1.495905874 | 0.107577 | 2.69E-27 | 1.132399247 | 0.855006653 | 1.167204583 | 0.165764219 | 0.968009043 | 0.064795427 | 0.556486087 | 2.28E-40 |
| E71R | 6 | 1.487740587 | 0.096919 | 4.57E-18 | 1.211918886 | 0.310680527 | 1.065319699 | 1.07E-24 | 1.28932628 | 1.07E-17 | 0.728847607 | 3.07E-33 |
| A122Q | 1 | 1.468116268 | 0.066885 | 0.001258786 | 1.355791709 | 0.436436228 | 1.288426158 | 0.003239759 | 1.721290666 | 1.89E-07 | 1.02749093 | 0.768868064 |
| C218X | 1 | 1.464737795 | 0.078253 | 1.38E-12 | 1.47991418 | 1.09E-14 | 0.36980612 | 8.35E-14 | 0.472150322 | 2.00E-22 | 0.856648184 | 0.047312844 |
| V61R | 1 | 1.461146038 | 0.076242 | 2.60E-51 | 1.8822922 | 6.93E-57 | 1.104740574 | 0.450500583 | 0.9346872 | 5.42E-05 | 1.434281369 | 6.53E-05 |
| R72Q | 1 | 1.460761711 | 0.081283 | 7.56E-104 | 0.93439135 | 4.01E-05 | 1.503198083 | 2.70E-38 | 1.265375396 | 9.92E-14 | 0.461299753 | 6.25E-143 |
| E71I | 2 | 1.456308158 | 0.088108 | 2.74E-10 | 1.022966581 | 0.070733474 | 1.037145935 | 3.11E-05 | 1.120060587 | 1.18E-06 | 0.677217767 | 2.46E-76 |
| D65G | 8 | 1.449171368 | 0.08831 | 1.76E-10 | 1.13284094 | 0.74615461 | 1.058156576 | 3.86E-05 | 1.033578079 | 0.00106425 | 0.926067468 | 2.12E-10 |
| Q64A | 4 | 1.447698713 | 0.087574 | 2.10E-07 | 1.166787529 | 0.049367108 | 1.094914541 | 2.10E-08 | 1.010207011 | 0.013422527 | 0.810114627 | 3.40E-16 |
| Q141G | 1 | 1.44465143 | 0.082781 | 3.29E-08 | 1.483506175 | 2.46E-18 | 0.223601461 | 9.09E-32 | 0.60542443 | 2.70E-07 | 1.175061581 | 0.000983863 |
| V74N | 4 | 1.434073732 | 0.059285 | 2.31E-09 | 1.028303199 | 0.703581117 | 0.943734677 | 0.009633423 | 0.974839696 | 0.548738663 | 0.950248223 | 1.78E-07 |
| T67P | 1 | 1.430919283 | 0.092492 | 7.70E-68 | 1.133357752 | 0.001137072 | 0.871230727 | 0.000203912 | 0.90377799 | 0.010673035 | 1.00300215 | 5.53E-50 |
| M68V | 1 | 1.430466915 | 0.088401 | 1.04E-24 | 1.253831352 | 4.01E-12 | 1.187736354 | 2.44E-10 | 1.06208813 | 0.255337457 | 0.869784335 | 6.31E-09 |
| P182N | 1 | 1.427893686 | 0.107546 | 1.65E-54 | 1.025266648 | 0.332304308 | 1.026191626 | 0.452531432 | 0.986639874 | 0.52139163 | 0.60578929 | 7.11E-73 |
| W178S | 2 | 1.427563413 | 0.059649 | 0.001414795 | 1.166352917 | 0.016137075 | 0.674466403 | 4.20E-08 | 0.776788945 | 1.60E-05 | 0.981500163 | 0.318247334 |
| E71V | 4 | 1.422906162 | 0.086871 | 2.03E-30 | 1.14971418 | 0.413559477 | 0.985929636 | 4.00E-12 | 1.084982746 | 2.33E-05 | 0.746878346 | 5.50E-41 |
| Q141E | 2 | 1.419126118 | 0.064881 | 0.076517633 | 1.563653278 | 1.50E-12 | 0.17306365 | 1.55E-33 | 0.534193609 | 1.64E-12 | 1.163292324 | 0.017277365 |
| L89X | 1 | 1.416312646 | 0.080756 | 5.44E-07 | 1.451520875 | 2.30E-72 | 0.638336671 | 7.36E-34 | 0.610920416 | 1.07E-81 | 1.263408512 | 0.341259496 |
| S189R | 6 | 1.413465685 | 0.091408 | 0.015006591 | 0.918037159 | 0.000354958 | 1.30168699 | 0.000141178 | 1.119156429 | 0.025721965 | 0.500100849 | 2.67E-54 |
| R72E | 1 | 1.409164025 | 0.071533 | 2.99E-18 | 1.031141909 | 0.292301153 | 1.046115472 | 0.856742235 | 1.238831377 | 0.016335445 | 0.591548388 | 2.60E-15 |
| L93L | 1 | 1.407790066 | 0.071015 | 2.00E-28 | 1.182290131 | 0.56646302 | 0.917658874 | 7.32E-05 | 0.851501457 | 2.05E-20 | 0.797506902 | 2.33E-17 |
| Q64S | 3 | 1.406843673 | 0.090976 | 3.24E-06 | 1.176458484 | 0.780448557 | 1.067816755 | 0.002213057 | 0.884020903 | 1.68E-06 | 0.921507976 | 0.000136458 |
| E71X | 1 | 1.40392584 | 0.081843 | 1.23E-22 | 1.078151425 | 5.51E-12 | 1.067584961 | 0.018680051 | 1.302350883 | 2.91E-11 | 0.677181746 | 7.31E-59 |
| R72X | 1 | 1.403526665 | 0.078098 | 2.48E-58 | 1.000789824 | 0.155664999 | 0.845099478 | 0.001189787 | 1.032097707 | 0.363601064 | 0.492970438 | 7.39E-96 |
| V74X | 1 | 1.402325626 | 0.067341 | 2.24E-30 | 0.864439336 | 5.37E-11 | 1.238594085 | 1.74E-06 | 1.322297866 | 1.88E-09 | 0.424876628 | 2.02E-66 |
| Q64L | 5 | 1.402140472 | 0.059088 | 2.21E-06 | 0.989033183 | 0.530922195 | 1.204281285 | 8.92E-05 | 1.143226054 | 0.010863414 | 0.674257648 | 4.50E-17 |
| E71L | 9 | 1.402025865 | 0.093407 | 2.75E-10 | 1.004582251 | 0.44257173 | 0.960303121 | 4.96E-05 | 1.125993439 | 6.21E-05 | 0.74159416 | 3.67E-38 |
| R72C | 1 | 1.398702094 | 0.06953 | 9.36E-12 | 0.991851853 | 0.289961939 | 0.855911196 | 0.32566729 | 1.045737926 | 0.711691011 | 0.459808613 | 8.51E-27 |
| W178X | 1 | 1.396295233 | 0.08099 | 8.16E-20 | 1.131199559 | 0.000655026 | 1.210458914 | 7.98E-10 | 1.06537624 | 0.002424504 | 0.839368049 | 2.01E-24 |
| E71T | 2 | 1.39379713 | 0.077582 | 2.06E-06 | 1.154611348 | 0.035299159 | 1.058479054 | 0.004139014 | 1.062313259 | 7.06E-17 | 0.954779923 | 0.000170795 |
| N81X | 1 | 1.39088346 | 0.063366 | 1.55E-23 | 1.375687122 | 5.90E-11 | 1.223571433 | 0.072286858 | 1.020448144 | 0.333307539 | 1.075888123 | 0.308448178 |
| N69X | 1 | 1.390649889 | 0.088299 | 8.48E-12 | 1.180694405 | 2.25E-64 | 1.217038293 | 0.103396892 | 1.193616237 | 1.25E-06 | 0.825173191 | 9.44E-15 |
| Q141C | 2 | 1.390644846 | 0.063579 | 0.004564614 | 1.571133902 | 5.63E-05 | 0.269803982 | 8.08E-15 | 0.488745946 | 3.23E-10 | 1.755542335 | 8.47E-12 |
| R78X | 1 | 1.389657358 | 0.091132 | 8.72E-18 | 0.96218421 | 0.027768546 | 0.723248946 | 1.93E-17 | 0.757556864 | 7.84E-19 | 0.391406133 | 1.27E-123 |
| V196X | 1 | 1.383659936 | 0.064983 | 1.23E-39 | 1.226886102 | 6.96E-07 | 1.172523852 | 2.18E-06 | 1.07368167 | 0.029301721 | 1.023605345 | 5.06E-11 |
| G70X | 1 | 1.380872648 | 0.094674 | 3.19E-10 | 1.031401164 | 0.868517394 | 1.038750586 | 3.50E-08 | 0.962162311 | 0.041200954 | 0.976582706 | 2.03E-05 |
| R72X | 1 | 1.377926759 | 0.076674 | 7.91E-74 | 0.896684544 | 4.89E-08 | 1.343383428 | 7.17E-25 | 1.634928254 | 2.67E-24 | 0.474004701 | 5.42E-120 |
| A55X | 1 | 1.371422706 | 0.07499 | 1.80E-27 | 1.089609547 | 0.308411125 | 1.182118675 | 3.96E-07 | 1.371353164 | 1.98E-18 | 0.942994805 | 0.04073312 |
| M68X | 1 | 1.368656745 | 0.062296 | 5.40E-49 | 1.247272038 | 1.56E-47 | 1.156437573 | 1.19E-08 | 0.965462032 | 0.82025493 | 0.932687484 | 0.000316464 |
| Q64X | 1 | 1.368333808 | 0.082024 | 1.46E-44 | 1.136666141 | 1.69E-09 | 1.058345227 | 0.139493589 | 0.860355685 | 1.62E-12 | 0.827556521 | 8.05E-09 |
| A210X | 1 | 1.367980913 | 0.090397 | 8.40E-85 | 1.391032583 | 9.27E-24 | 0.820377357 | 3.59E-15 | 0.762530086 | 4.32E-13 | 1.043248558 | 0.001267491 |
| W178X | 1 | 1.367376727 | 0.081077 | 3.67E-27 | 1.15336227 | 9.14E-06 | 0.878148674 | 8.22E-05 | 0.934060596 | 0.084908498 | 0.745290828 | 1.05E-12 |
| E71X | 1 | 1.366157188 | 0.094863 | 4.99E-20 | 1.049106343 | 0.214113608 | 1.39291185 | 0.47079863 | 1.625087057 | 0.457844993 | 0.897529169 | 0.004488492 |
| A55X | 1 | 1.365337629 | 0.076959 | 4.05E-21 | 1.384926327 | 3.17E-15 | 0.986416048 | 0.162168005 | 0.759446109 | 4.26E-08 | 1.009225979 | 0.074269512 |
| Q64X | 1 | 1.364738086 | 0.070186 | 6.18E-24 | 0.982325449 | 0.948090812 | 1.321135538 | 2.24E-32 | 1.196218701 | 3.32E-12 | 0.999214553 | 0.004681036 |
| V74X | 1 | 1.3643171 | 0.056401 | 1.27E-63 | 1.041478469 | 0.089019758 | 0.883729549 | 9.06E-09 | 0.829819194 | 3.58E-11 | 0.629046663 | 3.51E-50 |
| R72X | 1 | 1.363268561 | 0.079747 | 7.63E-24 | 1.05036882 | 0.189576951 | 0.785541127 | 0.01864294 | 0.861291529 | 0.64199775 | 0.506215033 | 2.70E-25 |
| Q64X | 1 | 1.360516516 | 0.059561 | 2.52E-77 | 1.020250339 | 0.079615807 | 1.192501898 | 2.56E-08 | 0.98330959 | 0.727932918 | 0.620032214 | 1.56E-66 |
| L89S | 4 | 1.358717264 | 0.083283 | 3.01E-07 | 1.518900408 | 9.47E-22 | 0.510689964 | 8.87E-49 | 0.555781333 | 3.94E-62 | 1.276073897 | 1.69E-10 |
| V61X | 1 | 1.35847678 | 0.083952 | 0.124659851 | 1.356861553 | 0.007960285 | 1.111085642 | 0.018194462 | 1.048072092 | 0.343504431 | 1.12237611 | 0.021170774 |
| L89M | 3 | 1.356613285 | 0.099793 | 5.84E-35 | 1.582252477 | 1.66E-50 | 0.641191368 | 5.06E-34 | 0.686066027 | 7.44E-41 | 1.168252525 | 0.010409031 |
| E71X | 1 | 1.355667542 | 0.083348 | 0.000700146 | 1.099805708 | 7.01E-08 | 1.137822919 | 0.033286558 | 1.062312233 | 0.483936519 | 0.942728325 | 4.08E-10 |
| N69X | 1 | 1.352867346 | 0.066709 | 2.10E-07 | 1.175132646 | 1.13E-14 | 1.105133703 | 0.059812348 | 1.038218556 | 0.00141402 | 1.001265795 | 1.07E-24 |
| S189N | 4 | 1.352183501 | 0.065088 | 2.09E-18 | 0.931121174 | 1.65E-10 | 1.204236909 | 2.16E-12 | 1.089705302 | 0.013993743 | 0.607661269 | 9.88E-52 |
| S189Q | 3 | 1.351654117 | 0.065748 | 2.62E-12 | 1.088353444 | 0.000788431 | 1.320664465 | 1.56E-18 | 0.980419006 | 2.56E-07 | 0.906367811 | 4.57E-07 |
| W178X | 1 | 1.349757352 | 0.080033 | 0.001741354 | 1.206870852 | 0.000245838 | 0.448019772 | 7.17E-33 | 0.680055429 | 2.01E-12 | 0.821952956 | 4.40E-06 |
| V61X | 1 | 1.348349761 | 0.073094 | 2.12E-38 | 1.520330629 | 9.86E-09 | 1.005029473 | 0.000105881 | 1.054715658 | 5.56E-06 | 0.854540761 | 2.29E-52 |
| A210X | 1 | 1.346301981 | 0.084944 | 4.04E-17 | 1.389624581 | 1.67E-16 | 0.869630025 | 1.17E-12 | 0.825078181 | 9.38E-09 | 1.054501382 | 0.012829799 |
| F226X | 1 | 1.345543441 | 0.111485 | 1.90E-25 | 1.329898505 | 3.43E-40 | 0.62104842 | 2.45E-20 | 0.809926584 | 1.29E-06 | 0.999500596 | 0.834195461 |
| N69X | 1 | 1.344009574 | 0.076575 | 9.81E-69 | 1.297253374 | 4.92E-08 | 1.154659389 | 8.17E-12 | 1.073231168 | 0.032176105 | 1.119839786 | 1.24E-09 |
| A60X | 1 | 1.34228926 | 0.059681 | 0.045849865 | 1.327974633 | 0.263320553 | 1.094334798 | 0.521135405 | 1.043129225 | 0.326743788 | 1.125952913 | 2.05E-05 |
| S189P | 5 | 1.341778872 | 0.0793 | 3.11E-24 | 1.134357269 | 0.834026065 | 1.122323193 | 0.000976345 | 0.928664325 | 0.239675775 | 0.728337244 | 2.36E-56 |
| T67X | 1 | 1.33970052 | 0.077083 | 2.51E-06 | 1.311630697 | 5.49E-26 | 0.937979539 | 7.05E-07 | 0.905596212 | 0.052092448 | 1.059077161 | 1.67E-14 |
| V61X | 1 | 1.338778079 | 0.082734 | 2.43E-27 | 1.313943095 | 1.71E-28 | 1.176924003 | 7.29E-18 | 1.00320107 | 0.311400524 | 1.020037758 | 5.56E-06 |
| T145X | 1 | 1.338411631 | 0.077009 | 1.42E-21 | 1.317896449 | 6.69E-80 | 1.10056525 | 0.017534875 | 0.912973681 | 0.032567961 | 1.217327394 | 7.91E-09 |
| M68X | 1 | 1.337366019 | 0.082599 | 1.32E-132 | 1.314354903 | 2.72E-168 | 1.165877168 | 9.02E-23 | 1.238369986 | 2.76E-31 | 1.116321394 | 8.87E-08 |
| L89H | 2 | 1.335144652 | 0.077365 | 0.16594213 | 1.333021379 | 0.00355306 | 0.673389531 | 0.000114201 | 0.634305679 | 3.78E-17 | 1.028939688 | 0.112957704 |
| E71X | 1 | 1.334839708 | 0.077081 | 0.226973272 | 1.023051249 | 0.072005444 | 0.959249298 | 0.599542335 | 0.953116219 | 0.267819197 | 0.718918892 | 9.42E-10 |
| D65F | 6 | 1.334310336 | 0.078233 | 0.016949645 | 1.107765208 | 0.776604856 | 1.022093157 | 0.003177723 | 1.069248676 | 0.003955268 | 0.767727426 | 2.20E-13 |

| | | | | | | | | | | |
|---|---|---|---|---|---|---|---|---|---|---|
| S189X | 1 | 1.33014828 | 0.090686 | 2.80E-25 | 0.979924245 | 0.588123294 | 0.938217077 | 0.000429796 | 1.065918393 | 0.019770877 | 0.674587271 | 2.47E-30 |
| V196X | 1 | 1.329018109 | 0.067141 | 1.35E-26 | 1.012413906 | 0.734853303 | 1.202645321 | 0.000371807 | 1.394898307 | 7.86E-06 | 0.78462506 | 6.61E-12 |
| D65X | 1 | 1.328452398 | 0.057164 | 3.88E-16 | 1.202287735 | 4.17E-12 | 0.325782568 | 5.97E-32 | 0.781654256 | 1.24E-06 | 0.899599461 | 0.042365931 |
| Y63X | 1 | 1.327687547 | 0.06158 | 2.15E-114 | 1.032473829 | 7.31E-05 | 1.094818967 | 1.49E-08 | 0.992150417 | 0.533793949 | 0.657418353 | 2.55E-108 |
| E71S | 6 | 1.326310076 | 0.080496 | 5.84E-05 | 1.30009524 | 0.907005217 | 1.074513869 | 0.000353748 | 1.043372842 | 0.000271664 | 0.98382171 | 0.062192028 |
| Q64X | 1 | 1.326003597 | 0.058051 | 3.46E-49 | 1.03054263 | 0.028981408 | 1.245563116 | 3.68E-09 | 0.992902554 | 0.828485454 | 0.650269183 | 6.23E-42 |
| R72L | 1 | 1.325063804 | 0.067264 | 6.90E-27 | 1.024178023 | 0.011145131 | 0.849469402 | 2.82E-06 | 0.846328908 | 4.40E-05 | 0.646326711 | 4.85E-25 |
| Q64H | 4 | 1.325007599 | 0.078282 | 2.82E-11 | 1.154557153 | 0.054364436 | 1.035570104 | 0.004264499 | 0.912743353 | 8.52E-09 | 0.919568591 | 0.000170934 |
| A122X | 1 | 1.324971707 | 0.060363 | 3.53E-11 | 1.214998033 | 1.90E-07 | 1.197097503 | 0.357308775 | 1.376773958 | 7.51E-05 | 1.131151963 | 0.37428428 |
| V61X | 1 | 1.324596115 | 0.078932 | 1.26E-68 | 1.465371998 | 1.41E-11 | 0.964954419 | 3.99E-05 | 0.967349925 | 1.34E-13 | 0.893114388 | 3.23E-22 |
| R72X | 1 | 1.323693212 | 0.077432 | 2.31E-12 | 1.03097136 | 0.227651496 | 0.808596375 | 0.000586636 | 0.788136433 | 0.007285818 | 0.545542867 | 4.85E-11 |
| Y63F | 2 | 1.322859032 | 0.059782 | 5.87E-06 | 1.130004234 | 0.276223232 | 0.840176324 | 0.279730088 | 0.916104278 | 0.016944541 | 0.742853857 | 0.020762946 |
| S189L | 2 | 1.322011433 | 0.066694 | 0.007432154 | 0.765041927 | 1.44E-18 | 1.256004224 | 0.312670256 | 1.724946679 | 1.37E-06 | 0.469237818 | 1.21E-18 |
| K227Y | 5 | 1.322007425 | 0.106405 | 4.46E-09 | 1.029764734 | 0.151927911 | 1.037347212 | 0.005526511 | 0.958054708 | 0.488457974 | 0.792244051 | 1.53E-09 |
| Y181F | 4 | 1.320270957 | 0.063722 | 0.007854837 | 1.080344064 | 0.539067452 | 0.700788931 | 2.11E-12 | 1.07268297 | 0.021593914 | 0.747580996 | 1.37E-08 |
| N81X | 1 | 1.320237941 | 0.060148 | 1.25E-16 | 1.282288505 | 7.34E-13 | 1.196882896 | 0.018088955 | 1.037293568 | 0.548122277 | 1.150227931 | 0.011579808 |
| W178X | 1 | 1.318211339 | 0.081107 | 0.015410968 | 1.699965524 | 0.119306806 | 0.485262933 | 5.90E-14 | 0.542115917 | 5.63E-30 | 1.289639412 | 0.002586402 |
| F117X | 1 | 1.317226464 | 0.0545 | 2.77E-20 | 1.143640299 | 7.67E-10 | 1.151936115 | 0.089922347 | 1.041826054 | 0.388028896 | 0.974531705 | 0.65165557 |
| L174X | 1 | 1.31529628 | 0.071119 | 0.000846959 | 1.035409639 | 0.034418655 | 1.260315152 | 9.50E-11 | 1.077875438 | 0.347559764 | 0.83469668 | 4.56E-05 |
| W178X | 1 | 1.314936042 | 0.072514 | 9.92E-08 | 1.168121425 | 0.430449721 | 0.585367262 | 0.001327797 | 0.871423547 | 0.273304656 | 0.685271682 | 2.32E-16 |
| M68X | 1 | 1.311571254 | 0.081005 | 2.10E-116 | 1.268250901 | 3.71E-157 | 1.049646686 | 0.037396633 | 1.017239308 | 0.09122048 | 1.092442465 | 4.99E-05 |
| L89X | 1 | 1.311539221 | 0.069955 | 8.06E-10 | 1.544206089 | 1.03E-15 | 0.760777476 | 0.001153109 | 0.739250114 | 0.004389935 | 1.284965455 | 1.27E-05 |
| N69X | 1 | 1.311529702 | 0.074724 | 3.77E-71 | 1.380455455 | 2.16E-17 | 1.112011862 | 2.54E-08 | 1.032018925 | 0.112840734 | 1.229209632 | 3.21E-11 |
| N69X | 1 | 1.309054888 | 0.087027 | 3.68E-22 | 1.016024095 | 0.015487249 | 1.215179054 | 0.071522663 | 1.215986418 | 2.10E-05 | 0.596722256 | 5.38E-88 |
| D65STOP | 1 | 1.306978593 | 0.083403 | 1.53E-05 | 0.938483902 | 5.33E-08 | 1.003651547 | 0.650125352 | 1.034296843 | 8.24E-05 | 0.624952045 | 8.23E-67 |
| N69X | 1 | 1.306875556 | 0.074459 | 2.01E-20 | 1.279915391 | 0.693854426 | 1.148023923 | 0.000627 | 1.088441197 | 0.446478775 | 0.964183823 | 3.21E-05 |
| A122X | 1 | 1.305873312 | 0.070107 | 2.40E-75 | 1.152914434 | 9.19E-24 | 1.265333193 | 5.17E-22 | 1.698282556 | 7.31E-53 | 0.886823258 | 5.79E-10 |
| F226L | 5 | 1.300587699 | 0.087756 | 0.012444963 | 1.044893443 | 0.52725794 | 1.153442945 | 0.000116917 | 1.087261623 | 4.30E-08 | 0.89157138 | 5.53E-11 |
| Q64M | 1 | 1.299238694 | 0.089098 | 3.72E-30 | 1.056380262 | 0.011337552 | 0.923689834 | 0.000831849 | 1.042169297 | 2.41E-13 | 0.781945274 | 2.85E-19 |
| L174X | 1 | 1.295016218 | 0.070022 | 4.10E-15 | 1.058490213 | 0.051711074 | 1.067563446 | 0.032496767 | 1.015346177 | 0.035085749 | 0.940787043 | 1.69E-15 |
| M68X | 1 | 1.292024562 | 0.079798 | 4.60E-104 | 1.349085512 | 1.24E-219 | 1.089113909 | 9.04E-08 | 1.06028529 | 0.00021772 | 1.252298043 | 1.46E-26 |
| C218T | 2 | 1.282783918 | 0.053823 | 2.91E-15 | 1.190289301 | 1.31E-05 | 0.520937633 | 5.75E-37 | 0.72555874 | 2.29E-14 | 0.781737317 | 2.92E-09 |
| E71X | 1 | 1.282770284 | 0.072271 | 0.065573018 | 0.99909014 | 0.531224844 | 1.025482936 | 0.396459273 | 1.185364804 | 8.63E-17 | 0.802878133 | 0.000918677 |
| S189X | 1 | 1.282292065 | 0.06469 | 3.10E-18 | 1.035842775 | 0.757458614 | 1.009452728 | 0.307401323 | 0.846724851 | 0.000226085 | 0.627479427 | 7.00E-24 |
| V61X | 1 | 1.281619374 | 0.058334 | 6.70E-42 | 1.407042718 | 4.84E-97 | 1.217043001 | 1.56E-19 | 0.948023822 | 0.027367519 | 1.302975695 | 2.17E-20 |
| L66G | 10 | 1.277740231 | 0.086912 | 9.78E-06 | 1.257578587 | 0.509325998 | 1.039089088 | 3.27E-09 | 0.944601079 | 1.13E-09 | 1.064323704 | 2.17E-09 |
| Q64X | 1 | 1.275358643 | 0.053745 | 1.30E-13 | 0.901635723 | 2.61E-06 | 1.016387272 | 0.365411449 | 0.982896775 | 0.507135511 | 0.481982618 | 3.96E-29 |
| I228N | 3 | 1.275207907 | 0.075455 | 0.000436434 | 1.078778852 | 0.047188825 | 1.054957927 | 0.083883639 | 1.005712204 | 0.523947718 | 1.194355714 | 9.14E-22 |
| V196X | 1 | 1.274520552 | 0.051175 | 2.57E-18 | 1.042608532 | 0.124832807 | 1.189503344 | 0.00287134 | 0.985731582 | 0.034559777 | 1.095051165 | 2.11E-07 |
| G118X | 1 | 1.274123251 | 0.057199 | 2.66E-14 | 1.050951101 | 0.15829558 | 0.979489859 | 1.32E-06 | 0.940579497 | 0.011404268 | 1.145136736 | 8.61E-17 |
| L66M | 1 | 1.269179777 | 0.076341 | 7.50E-27 | 1.247692312 | 0.106615197 | 0.886472894 | 3.12E-19 | 0.901935438 | 1.63E-21 | 0.971191873 | 1.10E-11 |
| A122X | 1 | 1.26876366 | 0.068115 | 3.70E-06 | 1.070438669 | 0.004824389 | 1.142150979 | 4.96E-05 | 1.434346507 | 1.87E-10 | 0.94446242 | 0.007485024 |
| R78X | 1 | 1.268073564 | 0.068078 | 1.01E-10 | 1.101215986 | 0.001963994 | 0.800099803 | 0.001644616 | 0.942790505 | 0.106046187 | 0.737144899 | 1.89E-08 |
| V61X | 1 | 1.265954222 | 0.078234 | 1.06E-18 | 1.150027806 | 2.02E-11 | 1.114003258 | 0.005616406 | 0.853877416 | 2.91E-10 | 0.934124752 | 0.020725068 |
| T67X | 1 | 1.263658687 | 0.072708 | 1.44E-16 | 1.231225586 | 2.04E-18 | 1.059293363 | 0.361885239 | 0.96327355 | 0.054535257 | 0.992097662 | 0.031783474 |
| M114A | 5 | 1.263484472 | 0.076784 | 0.195965867 | 1.41661521 | 0.360009884 | 0.297406019 | 8.78E-34 | 0.524947321 | 6.62E-22 | 1.100581318 | 0.052210059 |
| L66X | 1 | 1.259389925 | 0.067604 | 1.04E-12 | 1.161567269 | 0.010178771 | 1.047548656 | 1.54E-05 | 0.935363519 | 0.000925457 | 1.056819342 | 5.85E-10 |
| A55X | 1 | 1.259388973 | 0.062806 | 0.000613458 | 1.102264173 | 0.064314303 | 0.86578865 | 4.42E-08 | 0.777289616 | 5.47E-17 | 1.006142897 | 1.36E-08 |
| N69X | 1 | 1.256134369 | 0.071568 | 1.28E-05 | 1.369320309 | 8.51E-07 | 1.099645548 | 0.001433718 | 1.04418478 | 0.090091734 | 1.311265741 | 1.27E-16 |
| Q64X | 1 | 1.256024044 | 0.0635 | 3.46E-27 | 0.920376177 | 1.63E-09 | 0.979929875 | 0.016769986 | 0.988893591 | 0.817003117 | 0.50918856 | 3.31E-72 |
| I228E | 2 | 1.255312624 | 0.074117 | 1.55E-09 | 1.068542659 | 0.026251828 | 1.00970765 | 0.210119133 | 0.929967286 | 0.243754898 | 1.075746118 | 3.03E-05 |
| M114T | 2 | 1.25332163 | 0.076167 | 1.46E-12 | 1.31966979 | 5.35E-10 | 0.313170107 | 1.25E-41 | 0.611273566 | 2.10E-34 | 1.426525959 | 8.73E-11 |
| I228A | 7 | 1.251548895 | 0.088055 | 1.74E-05 | 1.161961543 | 0.690660219 | 1.102601609 | 0.001500369 | 0.925424781 | 0.000245148 | 0.932245107 | 1.46E-08 |
| V74C | 1 | 1.245595637 | 0.062044 | 0.000735937 | 0.799314256 | 1.46E-12 | 1.615335855 | 6.14E-09 | 1.562224183 | 4.57E-14 | 0.417490847 | 6.39E-24 |
| G70X | 1 | 1.245495388 | 0.094492 | 2.88E-59 | 1.108487516 | 2.25E-20 | 0.972858143 | 0.061468278 | 0.919217681 | 4.09E-05 | 0.831124108 | 1.57E-24 |
| E71X | 1 | 1.244019701 | 0.079515 | 2.08E-16 | 1.197125626 | 0.382241557 | 0.867413378 | 7.09E-05 | 0.885891945 | 0.0007253 | 0.956830508 | 0.000941458 |
| V61X | 1 | 1.243929912 | 0.076873 | 2.09E-25 | 1.147610595 | 0.361949225 | 1.10880783 | 0.000378377 | 1.059680861 | 0.39999291 | 1.230874105 | 0.299530102 |
| Q64X | 1 | 1.2422838 | 0.073384 | 2.19E-05 | 1.090356811 | 5.49E-05 | 0.792781478 | 3.78E-10 | 0.950107963 | 0.02104412 | 0.949854199 | 0.007144041 |
| S189M | 5 | 1.240734021 | 0.062594 | 1.52E-05 | 0.8684186 | 3.15E-10 | 1.170749969 | 0.002635174 | 1.313522147 | 2.42E-07 | 0.508640447 | 3.50E-49 |
| E71X | 1 | 1.240679872 | 0.091808 | 1.27E-56 | 1.042881671 | 8.84E-07 | 0.943547946 | 0.156301405 | 0.998061293 | 0.237785274 | 0.696275984 | 2.50E-60 |
| F226Y | 3 | 1.239705174 | 0.075855 | 1.09E-10 | 1.128693137 | 0.421628894 | 1.068440809 | 0.000866041 | 0.979441179 | 4.32E-07 | 1.004329919 | 5.04E-05 |
| N69X | 1 | 1.239238619 | 0.070606 | 1.26E-21 | 1.234607696 | 5.07E-23 | 1.200139564 | 2.89E-09 | 1.165902841 | 0.002337078 | 1.084286465 | 0.04732287 |
| L66X | 1 | 1.238772757 | 0.073538 | 9.32E-07 | 1.174051692 | 7.52E-17 | 0.900749879 | 1.81E-41 | 0.946666638 | 3.34E-16 | 1.050429256 | 0.089092493 |
| F226S | 14 | 1.238179293 | 0.10259 | 3.51E-06 | 1.345346643 | 0.197734935 | 1.032234154 | 0.012093569 | 1.021386773 | 0.007515897 | 0.944209663 | 0.001796333 |
| L66S | 4 | 1.23621043 | 0.073172 | 8.24E-06 | 1.252527248 | 0.967793133 | 0.996584749 | 2.41E-09 | 0.94464309 | 1.03E-07 | 1.105412459 | 5.55E-08 |
| L174V | 3 | 1.233573592 | 0.073487 | 5.94E-12 | 1.459824231 | 0.628947457 | 0.673251081 | 3.96E-24 | 0.72118702 | 1.82E-34 | 0.83952948 | 7.26E-07 |
| F226X | 1 | 1.232729869 | 0.102138 | 0.000116608 | 1.118868329 | 1.15E-05 | 0.716182901 | 4.83E-07 | 0.84631948 | 0.010185009 | 0.908266425 | 0.202183084 |
| G229L | 6 | 1.23190619 | 0.076201 | 2.75E-10 | 1.084348348 | 0.694174059 | 1.020352771 | 0.00023645 | 0.821324535 | 2.84E-14 | 1.311820537 | 3.02E-11 |
| V196X | 1 | 1.231522725 | 0.056495 | 0.000322001 | 1.129788774 | 0.00719517 | 1.097127858 | 0.001586851 | 1.005853238 | 3.43E-05 | 1.286331663 | 1.45E-26 |
| Y63W | 7 | 1.230209668 | 0.080676 | 4.73E-14 | 1.173808058 | 0.639199835 | 0.964841604 | 0.004714428 | 1.073482041 | 3.45E-10 | 0.934726512 | 5.59E-19 |
| W178X | 1 | 1.228174884 | 0.080584 | 0.062312769 | 1.166260776 | 0.855499868 | 0.402214098 | 1.07E-30 | 0.864279733 | 0.000145511 | 0.822605901 | 9.04E-10 |
| W178X | 1 | 1.227772786 | 0.065094 | 0.00011735 | 1.267636603 | 0.000553538 | 0.527877064 | 2.98E-15 | 0.819118277 | 0.007881519 | 0.962058483 | 0.575017572 |
| V61X | 1 | 1.22750841 | 0.075858 | 2.84E-46 | 1.195638403 | 1.91E-60 | 1.062696241 | 0.00644686 | 0.931565566 | 0.013937581 | 1.12543299 | 4.87E-07 |
| Q64X | 1 | 1.226205497 | 0.058177 | 5.20E-39 | 1.12253151 | 4.77E-21 | 1.03352293 | 0.685173688 | 1.124313291 | 1.47E-07 | 0.868031818 | 2.27E-14 |
| S189A | 4 | 1.225386244 | 0.061819 | 5.25E-05 | 1.246036408 | 0.213133707 | 1.285788875 | 5.39E-05 | 1.012958145 | 0.13205317 | 1.011255222 | 1.52E-10 |
| L93X | 1 | 1.225317027 | 0.059443 | 5.61E-18 | 1.082361529 | 0.001374587 | 1.084832346 | 0.001225259 | 0.954941654 | 0.001595524 | 1.170958823 | 3.69E-13 |
| E71F | 2 | 1.22513444 | 0.078022 | 9.03E-11 | 1.074532872 | 9.32E-07 | 1.178166431 | 3.05E-06 | 1.395789005 | 9.06E-16 | 0.849584335 | 2.32E-10 |
| V61X | 1 | 1.225077901 | 0.066391 | 1.18E-12 | 1.641464899 | 0.818360913 | 0.990062283 | 0.056393648 | 1.007507015 | 0.88337123 | 0.890382166 | 1.11E-18 |
| A122X | 1 | 1.22449006 | 0.094246 | 1.81E-19 | 1.340918796 | 1.18E-13 | 0.707987031 | 0.007980341 | 0.751260141 | 0.304692713 | 1.250727795 | 3.72E-09 |
| N69X | 1 | 1.223837435 | 0.094195 | 1.24E-06 | 1.054789058 | 5.71E-13 | 1.214395163 | 0.103059595 | 0.997496522 | 0.588199589 | 0.800749038 | 0.151060994 |
| Y63X | 1 | 1.223085023 | 0.072192 | 0.165357507 | 1.173712184 | 2.36E-11 | 0.801945751 | 1.16E-11 | 1.094504972 | 6.41E-10 | 1.149088249 | 1.16E-07 |
| I228V | 4 | 1.221633055 | 0.098326 | 0.087256254 | 1.124832827 | 0.124017213 | 0.873120915 | 2.94E-07 | 0.980974352 | 4.82E-05 | 0.914006797 | 2.48E-09 |
| L66Y | 3 | 1.220360615 | 0.062685 | 0.116763067 | 1.319756672 | 0.001873988 | 1.12778241 | 2.39E-09 | 1.124384101 | 2.29E-08 | 1.061732733 | 1.20E-12 |

| | | | | | | | | | | | | |
|---|---|---|---|---|---|---|---|---|---|---|---|---|
| D207X | 1 | 1.219775996 | 0.080603 | 7.61E-07 | 1.020460154 | 0.406821897 | 0.70510631 | 3.62E-08 | 0.742050278 | 0.02077715 | 0.694099286 | 1.53E-10 |
| I228S | 9 | 1.219742748 | 0.083633 | 2.10E-05 | 1.106758674 | 0.826388425 | 1.079674206 | 1.86E-07 | 0.929795828 | 0.002945088 | 1.104330799 | 6.46E-10 |
| Q64X | 1 | 1.219526839 | 0.053389 | 3.11E-27 | 0.956531989 | 0.005701353 | 1.111828696 | 0.069865093 | 0.907172387 | 0.038910867 | 0.680738925 | 7.66E-36 |
| W178X | 1 | 1.218725988 | 0.072263 | 0.002366588 | 1.060267552 | 0.14269553 | 0.515393569 | 3.51E-08 | 0.575206961 | 1.68E-10 | 0.747767813 | 4.46E-08 |
| D92X | 1 | 1.218356041 | 0.091553 | 1.44E-44 | 1.17730859 | 4.27E-45 | 1.272983587 | 4.30E-28 | 0.852111842 | 2.79E-29 | 0.984416535 | 0.095699896 |
| L66C | 4 | 1.217392327 | 0.064843 | 5.41E-10 | 1.217538738 | 0.217971852 | 0.968025233 | 0.001013697 | 0.994638388 | 4.28E-06 | 1.268382588 | 9.00E-23 |
| Q64X | 1 | 1.217380032 | 0.051302 | 3.43E-07 | 0.836061503 | 1.88E-09 | 1.060164916 | 0.1013114 | 0.91049259 | 0.271753567 | 0.430313503 | 5.27E-25 |
| C138S | 7 | 1.215950357 | 0.101158 | 0.03202211 | 0.96599989 | 0.001950478 | 1.134202999 | 3.01E-05 | 1.075884861 | 0.000528455 | 0.697608024 | 5.62E-28 |
| V61X | 1 | 1.215090897 | 0.093522 | 0.245564746 | 1.056231957 | 1.89E-16 | 1.337026481 | 5.92E-26 | 1.13251883 | 4.23E-09 | 0.823116351 | 3.68E-08 |
| N69X | 1 | 1.214976174 | 0.069905 | 4.60E-11 | 1.203260884 | 1.82E-18 | 1.095689881 | 0.084003914 | 0.905853406 | 0.129284555 | 1.25158842 | 0.000661474 |
| V74X | 1 | 1.213840109 | 0.050181 | 2.63E-12 | 0.940360544 | 0.004667144 | 0.726536017 | 4.89E-14 | 0.886731708 | 0.004091333 | 0.67101663 | 3.99E-25 |
| D65X | 1 | 1.207098306 | 0.068104 | 3.79E-10 | 1.005959656 | 0.55198794 | 1.282817988 | 1.68E-13 | 1.013013444 | 0.035384564 | 0.95139985 | 3.91E-06 |
| M68X | 1 | 1.205663784 | 0.074464 | 3.31E-61 | 1.349133538 | 1.52E-196 | 1.054728057 | 0.004548852 | 1.010756356 | 0.24508519 | 1.37377561 | 8.75E-53 |
| E71X | 1 | 1.205467419 | 0.078207 | 1.26E-15 | 1.115930601 | 0.290393291 | 1.221046148 | 1.37E-05 | 1.205244497 | 0.010908918 | 0.879939387 | 0.003790497 |
| E71X | 1 | 1.205373977 | 0.079611 | 0.482685896 | 0.96201282 | 9.05E-05 | 1.045356207 | 7.19E-05 | 1.100837223 | 0.023779738 | 0.8349441 | 0.058002994 |
| N69X | 1 | 1.2049675 | 0.068653 | 2.94E-08 | 1.294934976 | 4.85E-26 | 1.069569929 | 0.275933215 | 1.107030182 | 0.203531361 | 1.225271635 | 7.53E-11 |
| K227A | 4 | 1.203028008 | 0.097378 | 0.004207979 | 1.043900302 | 0.054609115 | 1.043542315 | 0.057613485 | 0.957509724 | 0.024228842 | 0.870020652 | 3.33E-07 |
| T145X | 1 | 1.20202459 | 0.077696 | 2.14E-22 | 0.98833771 | 0.949028911 | 0.731038404 | 0.007417303 | 0.774392117 | 0.060346825 | 1.216802963 | 2.79E-22 |
| G229X | 1 | 1.2004979 | 0.061772 | 1.86E-14 | 0.994417864 | 0.546296269 | 0.96530276 | 2.04E-06 | 0.884889196 | 0.002280445 | 0.967079043 | 2.16E-10 |
| T145X | 1 | 1.200469144 | 0.077596 | 4.22E-42 | 1.129086801 | 8.14E-38 | 1.124561721 | 0.001359259 | 0.98781786 | 0.182468793 | 1.177576912 | 7.84E-06 |
| I228K | 4 | 1.199937923 | 0.087085 | 0.007170766 | 1.191807917 | 0.039983369 | 1.362065351 | 2.02E-15 | 1.036785398 | 0.000209359 | 1.002491358 | 8.66E-05 |
| S189H | 3 | 1.199924034 | 0.070503 | 8.34E-21 | 0.910095928 | 6.23E-17 | 1.127994439 | 4.38E-09 | 1.255318306 | 7.61E-22 | 0.631822588 | 2.49E-79 |
| V61X | 1 | 1.198551069 | 0.073143 | 1.51E-35 | 1.216445291 | 1.34E-30 | 0.945814555 | 0.038345397 | 0.921505234 | 5.30E-13 | 1.023802495 | 0.010116871 |
| E71X | 1 | 1.198147812 | 0.078299 | 6.07E-09 | 1.092228266 | 0.006384361 | 1.086414835 | 0.236782918 | 1.118064687 | 0.00811989 | 0.843483301 | 2.89E-05 |
| V196X | 1 | 1.196582122 | 0.054743 | 5.13E-10 | 1.157919697 | 2.24E-05 | 1.157098466 | 0.001719167 | 0.930919311 | 0.076690823 | 1.24923628 | 0.413849966 |
| V61X | 1 | 1.193932019 | 0.080909 | 3.22E-40 | 1.184312452 | 1.97E-08 | 1.042834411 | 0.250941941 | 0.953797005 | 0.00012807 | 1.070063482 | 4.44E-07 |
| C138X | 1 | 1.193889106 | 0.081329 | 1.70E-12 | 1.274537085 | 0.424484385 | 0.799601108 | 0.064815564 | 0.91746561 | 0.005339465 | 1.405603939 | 0.020129325 |
| F226X | 1 | 1.193322568 | 0.064286 | 2.66E-53 | 1.073275346 | 6.15E-15 | 0.87345744 | 1.00E-24 | 0.846784707 | 1.56E-25 | 0.842315185 | 1.34E-16 |
| W178X | 1 | 1.192671829 | 0.07151 | 2.85E-12 | 0.987110373 | 0.061437007 | 1.041287339 | 0.159118246 | 1.167421085 | 0.000816753 | 0.733789647 | 7.56E-25 |
| V74X | 1 | 1.191311396 | 0.077562 | 1.31E-10 | 1.115002375 | 6.66E-09 | 0.95230777 | 0.062326165 | 0.933441447 | 0.013181935 | 0.947789271 | 0.256303577 |
| Q64X | 1 | 1.191143262 | 0.052147 | 5.50E-15 | 0.939469958 | 7.13E-05 | 1.229391514 | 6.91E-05 | 1.227111367 | 4.85E-06 | 0.690013448 | 1.83E-30 |
| W178X | 1 | 1.191048115 | 0.068051 | 7.60E-06 | 1.222590358 | 0.058066562 | 0.416699643 | 6.79E-31 | 0.745252782 | 2.75E-11 | 1.085166637 | 0.092248287 |
| L66E | 3 | 1.190122427 | 0.0739 | 1.52E-17 | 1.240951697 | 0.02900819 | 1.040877544 | 4.48E-10 | 0.992047784 | 4.23E-07 | 1.138551902 | 1.65E-17 |
| K227P | 7 | 1.18844251 | 0.079306 | 0.001508462 | 0.961436242 | 0.011455305 | 1.09015462 | 0.003740501 | 1.074149916 | 0.005746663 | 0.771434076 | 1.74E-18 |
| G70X | 1 | 1.188257419 | 0.084545 | 1.10E-06 | 1.038135985 | 0.016926652 | 1.166906589 | 0.012810201 | 1.053023739 | 0.007298064 | 0.908884419 | 0.00036164 |
| K227V | 6 | 1.188034934 | 0.081182 | 0.034240419 | 0.997620282 | 0.347952898 | 1.122801944 | 2.44E-12 | 1.101250745 | 1.23E-05 | 0.833023547 | 4.11E-10 |
| Y146X | 1 | 1.188032457 | 0.04997 | 0.005952545 | 1.103077264 | 5.45E-13 | 0.992088698 | 0.396985309 | 0.904642045 | 0.002721266 | 1.06854824 | 2.98E-11 |
| V61X | 1 | 1.187146891 | 0.073364 | 5.45E-23 | 1.104046344 | 1.73E-19 | 1.118590961 | 6.67E-08 | 0.979479859 | 0.667294205 | 0.94427366 | 0.137318136 |
| K227L | 8 | 1.186212768 | 0.089769 | 5.92E-07 | 1.231710243 | 0.800888334 | 1.023352296 | 0.007503852 | 0.956822267 | 6.84E-05 | 0.988867212 | 0.000262173 |
| A60X | 1 | 1.185913624 | 0.060567 | 1.12E-05 | 1.195270046 | 5.63E-14 | 1.0223191 | 0.066002991 | 1.038618975 | 0.27720894 | 1.081527229 | 0.001395706 |
| Q64X | 1 | 1.184364764 | 0.05185 | 8.09E-18 | 0.965363185 | 0.042671789 | 1.437675798 | 3.54E-21 | 1.21913744 | 5.43E-10 | 0.676970712 | 1.70E-28 |
| D65W | 3 | 1.182618085 | 0.082396 | 2.49E-07 | 1.137996951 | 0.839090484 | 1.028431387 | 0.06474565 | 1.115380229 | 0.002336002 | 0.883041124 | 1.29E-05 |
| A122X | 1 | 1.182070824 | 0.090981 | 1.33E-21 | 1.035667066 | 0.513030283 | 0.936509495 | 0.002607097 | 1.12137717 | 0.001304879 | 1.097767707 | 3.85E-16 |
| A122T | 1 | 1.181980906 | 0.053849 | 0.317400675 | 1.072992344 | 0.007055156 | 1.084227867 | 0.692817579 | 0.983873178 | 0.645028116 | 1.175546412 | 0.088533468 |
| G229F | 2 | 1.181425304 | 0.080546 | 1.65E-11 | 0.907705111 | 0.014607545 | 2.026733202 | 0.291041398 | 1.236454746 | 0.342739585 | 1.3268806 | 8.15E-21 |
| P182X | 1 | 1.179888546 | 0.088662 | 2.34E-21 | 1.038737574 | 2.38E-08 | 1.037087382 | 0.015818622 | 0.97723662 | 0.002894989 | 0.945843879 | 0.000238143 |
| E71P | 2 | 1.179355374 | 0.078477 | 0.000628092 | 1.012040019 | 0.108658666 | 0.973666653 | 0.005065402 | 1.08482863 | 2.57E-10 | 0.833648909 | 1.14E-09 |
| T67X | 1 | 1.179039319 | 0.067839 | 3.13E-05 | 1.143902734 | 5.00E-09 | 1.18170438 | 0.001313869 | 1.008478639 | 0.116779317 | 1.269944235 | 0.012527885 |
| L66X | 1 | 1.177927603 | 0.064064 | 0.008996654 | 1.159623965 | 1.29E-05 | 1.099585033 | 0.000312377 | 0.907783826 | 2.35E-06 | 1.26464728 | 1.32E-14 |
| A60X | 1 | 1.177335818 | 0.072228 | 3.26E-10 | 1.094638497 | 0.049918199 | 1.002355881 | 0.51197713 | 0.967838275 | 0.037843975 | 0.880966133 | 0.357670693 |
| E71X | 1 | 1.177158312 | 0.081792 | 0.05651152 | 0.992312244 | 0.297333935 | 1.067506041 | 2.16E-11 | 1.036264047 | 0.261364138 | 0.886246935 | 0.363177764 |
| E71X | 1 | 1.176954035 | 0.081725 | 4.47E-17 | 1.040351584 | 0.00011516 | 1.143915481 | 0.018076115 | 1.152726122 | 0.192181521 | 0.731893588 | 5.43E-29 |
| V74X | 1 | 1.176347394 | 0.058595 | 4.86E-30 | 0.986039675 | 0.309796275 | 1.025129697 | 0.530247463 | 1.055529846 | 0.002826463 | 0.792630855 | 1.33E-26 |
| F226I | 1 | 1.17572394 | 0.07194 | 3.26E-07 | 1.061540719 | 0.250377633 | 0.91200724 | 2.45E-07 | 0.951524456 | 0.000910544 | 1.059735021 | 0.010460149 |
| G70F | 3 | 1.175011796 | 0.089145 | 0.00020784 | 1.009512176 | 0.076062345 | 1.095271163 | 6.89E-06 | 0.967051061 | 0.003748332 | 0.753307423 | 6.77E-24 |
| G229R | 6 | 1.174643913 | 0.081501 | 0.00599528 | 1.167490224 | 0.059245629 | 1.28652192 | 4.35E-06 | 0.974979335 | 0.005834752 | 1.177860753 | 2.55E-07 |
| E71X | 1 | 1.173856061 | 0.076727 | 2.68E-14 | 1.180957693 | 0.000228638 | 1.114573055 | 0.166306579 | 1.111850514 | 0.383685542 | 0.978220612 | 0.119274019 |
| D65L | 3 | 1.173801796 | 0.045666 | 0.007152047 | 0.997319251 | 0.991385034 | 0.904388498 | 0.034172403 | 0.970648501 | 0.035397178 | 0.869543006 | 0.000781698 |
| D65Y | 2 | 1.171311675 | 0.071378 | 1.32E-12 | 1.156823701 | 0.068813531 | 1.075527471 | 0.000222116 | 0.99410974 | 0.017825446 | 0.947098205 | 0.000240426 |
| E71H | 1 | 1.171281195 | 0.085286 | 4.39E-05 | 1.000942846 | 0.446390055 | 1.115492478 | 0.024249188 | 1.136610251 | 2.23E-06 | 0.899009116 | 0.274796464 |
| L66X | 1 | 1.170896958 | 0.06631 | 3.77E-08 | 1.083373883 | 0.019686178 | 0.959319074 | 0.168281732 | 0.965318336 | 2.62E-08 | 1.147722329 | 6.67E-06 |
| V196X | 1 | 1.170364886 | 0.055087 | 5.38E-07 | 1.245645901 | 1.14E-13 | 1.019639941 | 0.838031426 | 0.807079235 | 1.04E-06 | 1.335790756 | 1.33E-58 |
| V74X | 1 | 1.16998855 | 0.066018 | 0.007058488 | 1.031362139 | 0.205254748 | 0.886585538 | 1.17E-05 | 1.001334615 | 0.277246438 | 0.978738438 | 0.012544765 |
| G70L | 3 | 1.16881261 | 0.080759 | 0.000835121 | 1.005161508 | 0.838105258 | 1.060876168 | 0.191239068 | 0.997447242 | 0.077949871 | 0.855137504 | 6.85E-05 |
| G229M | 5 | 1.168739779 | 0.062995 | 1.01E-19 | 1.098186088 | 0.602090903 | 1.037313411 | 0.023419036 | 0.805355523 | 3.57E-27 | 1.153465543 | 0.018998143 |
| A122X | 1 | 1.168702631 | 0.062743 | 2.25E-24 | 1.114916384 | 5.19E-12 | 1.296385341 | 9.74E-22 | 1.132511989 | 0.016078187 | 0.968943178 | 0.041615916 |
| D207X | 1 | 1.168340611 | 0.054065 | 9.14E-10 | 1.237613355 | 0.000111485 | 1.116367127 | 0.0001343 | 0.895726201 | 0.001280108 | 1.157077125 | 8.51E-06 |
| L174X | 1 | 1.168022324 | 0.060406 | 3.18E-11 | 0.980864187 | 0.198884696 | 1.047838861 | 0.598206875 | 0.791164216 | 4.09E-07 | 0.841695886 | 8.48E-07 |
| G229V | 5 | 1.167868793 | 0.07196 | 7.95E-16 | 1.061959582 | 4.86E-05 | 1.137304671 | 0.000629384 | 1.035669842 | 0.011574344 | 1.145510231 | 0.005750789 |
| K227M | 3 | 1.166754626 | 0.058632 | 0.023779784 | 1.096321311 | 0.368177301 | 0.939086257 | 0.003215489 | 1.073754425 | 0.01471843 | 0.952214067 | 0.130258666 |
| F226W | 1 | 1.166444571 | 0.093884 | 3.24E-12 | 0.985192118 | 0.895655426 | 0.935826304 | 2.69E-07 | 0.957675093 | 0.950031452 | 0.866923035 | 2.36E-07 |
| W178X | 1 | 1.166275289 | 0.067283 | 0.038604576 | 1.167949479 | 4.07E-06 | 0.421005536 | 1.92E-42 | 0.639230072 | 1.10E-35 | 0.924263838 | 0.010204538 |
| L66I | 1 | 1.165622951 | 0.072385 | 0.064641958 | 1.050742684 | 2.02E-06 | 1.06479245 | 0.112458927 | 1.026114506 | 0.536806288 | 0.824773123 | 0.00077757 |
| E71X | 1 | 1.165591592 | 0.071662 | 0.234140175 | 0.960522638 | 0.029927775 | 1.082619739 | 0.01984227 | 1.193852886 | 0.195243055 | 0.856531167 | 0.006060323 |
| E71X | 1 | 1.165314562 | 0.094109 | 0.08605806 | 1.0771331 | 9.89E-07 | 0.770775827 | 1.63E-09 | 0.984227054 | 0.063088354 | 0.861204908 | 2.93E-06 |
| V61X | 1 | 1.164581878 | 0.071446 | 2.86E-06 | 1.068367123 | 0.364619514 | 1.046928054 | 0.00999737 | 1.017464558 | 0.072999857 | 0.933861454 | 0.02298184 |
| T67X | 1 | 1.163961192 | 0.066971 | 0.013878012 | 1.187714652 | 0.05191769 | 0.943889668 | 0.010820543 | 1.0140103 | 0.720105294 | 1.149220373 | 8.02E-06 |
| G229C | 1 | 1.163544273 | 0.059871 | 7.01E-08 | 1.027626166 | 0.168745409 | 0.84730397 | 0.238978347 | 0.778272935 | 6.24E-12 | 1.175202441 | 0.033549244 |
| L93X | 1 | 1.162855485 | 0.056413 | 4.25E-08 | 1.130769337 | 0.007047109 | 0.775254406 | 4.03E-11 | 1.042432667 | 0.016941017 | 1.252155164 | 5.21E-17 |
| E71G | 4 | 1.162441889 | 0.079257 | 7.69E-12 | 1.176977681 | 0.596942411 | 1.001080695 | 0.001192295 | 0.977708956 | 0.091420676 | 1.045369324 | 2.82E-09 |
| M68X | 1 | 1.162123923 | 0.071817 | 1.55E-28 | 1.248098456 | 8.09E-88 | 1.099952809 | 7.76E-08 | 0.957124478 | 0.029721954 | 1.308834233 | 2.98E-25 |
| C218I | 1 | 1.158901402 | 0.048626 | 1.02E-06 | 1.155502651 | 6.84E-06 | 0.479004751 | 3.88E-39 | 0.783562254 | 7.87E-08 | 0.92663961 | 0.021576097 |
| G70X | 1 | 1.15867817 | 0.079078 | 0.0126595 | 1.106025781 | 5.74E-05 | 1.110676514 | 5.45E-06 | 1.06194821 | 6.95E-11 | 0.909629462 | 0.273010355 |

| | | | | | | | | | | | | |
|---|---|---|---|---|---|---|---|---|---|---|---|---|
| I228X | 1 | 1.157641586 | 0.079375 | 0.263018857 | 1.035507876 | 0.176609378 | 1.113043256 | 0.477373828 | 0.903712877 | 0.422196652 | 0.946897548 | 0.027870227 |
| N69X | 1 | 1.157293852 | 0.066586 | 7.48E-09 | 1.182459879 | 7.28E-12 | 1.060448703 | 0.213478136 | 0.874100596 | 0.000563378 | 1.309095792 | 2.55E-05 |
| L66X | 1 | 1.155928587 | 0.075755 | 8.47E-15 | 1.225008912 | 0.303854935 | 1.019863228 | 0.062706797 | 0.890547915 | 2.38E-11 | 1.048264081 | 4.73E-08 |
| K227T | 4 | 1.155092766 | 0.095706 | 0.014967914 | 1.136917092 | 3.50E-05 | 1.009093584 | 2.78E-05 | 0.994935365 | 0.11927456 | 0.859416614 | 0.000460918 |
| Y146X | 1 | 1.154412887 | 0.069956 | 0.008620792 | 1.348940295 | 0.657580353 | 1.080162231 | 0.1862535 | 1.033765931 | 0.264598748 | 0.968819876 | 1.76E-12 |
| Q64X | 1 | 1.15370347 | 0.064535 | 0.045842408 | 0.979246193 | 0.00882346 | 0.827293506 | 9.59E-06 | 1.200338091 | 0.001568712 | 0.853485344 | 2.86E-15 |
| T67X | 1 | 1.153586617 | 0.066375 | 6.05E-05 | 1.225578303 | 0.501333988 | 1.047724465 | 0.183172613 | 0.925714405 | 0.033610844 | 1.252004655 | 7.75E-31 |
| G70H | 1 | 1.152879013 | 0.083056 | 8.96E-12 | 1.013144112 | 0.406553073 | 1.480618865 | 1.87E-11 | 1.164972812 | 2.56E-05 | 0.893885639 | 2.01E-06 |
| I228X | 1 | 1.152401588 | 0.068188 | 0.003311132 | 1.081340835 | 0.727312715 | 0.942147246 | 0.155204465 | 0.900136958 | 0.005295605 | 1.048589003 | 0.018547589 |
| L89X | 1 | 1.151373174 | 0.06565 | 0.001118896 | 1.0774928 | 1.92E-10 | 0.966475716 | 0.235216949 | 1.031488421 | 0.00114504 | 0.9775195 | 2.02E-21 |
| W178X | 1 | 1.149878808 | 0.097016 | 0.479504372 | 1.599729774 | 0.005252974 | 0.316186749 | 2.14E-07 | 0.702841819 | 0.073624885 | 1.434421113 | 0.000535105 |
| D65M | 1 | 1.149263484 | 0.044711 | 3.28E-05 | 1.068705071 | 0.000298432 | 0.922178851 | 0.018741984 | 0.996883732 | 0.573712389 | 0.998013074 | 0.585824459 |
| V61X | 1 | 1.148493396 | 0.070459 | 2.20E-23 | 1.210130861 | 4.28E-05 | 1.006571543 | 0.093858691 | 0.886363962 | 9.75E-08 | 1.258369456 | 2.37E-08 |
| W178X | 1 | 1.146800266 | 0.047918 | 0.000272206 | 1.041973079 | 0.172075566 | 0.647316367 | 1.14E-05 | 1.10458052 | 0.011012611 | 0.997763028 | 0.007030158 |
| Y146X | 1 | 1.146432661 | 0.052202 | 5.15E-28 | 1.123350278 | 9.83E-25 | 0.974933205 | 0.075832353 | 0.879480836 | 1.55E-06 | 1.012141444 | 0.336845871 |
| E71X | 1 | 1.144833088 | 0.082422 | 0.238593128 | 0.968763169 | 0.005569303 | 0.712852065 | 0.024333526 | 0.907044207 | 0.226318943 | 0.82961909 | 0.166109401 |
| V74X | 1 | 1.144487125 | 0.074513 | 8.95E-15 | 1.055765699 | 0.000837397 | 1.266351135 | 2.86E-18 | 1.091637779 | 9.06E-08 | 0.809010547 | 2.76E-14 |
| W178X | 1 | 1.144453335 | 0.044479 | 0.014927358 | 1.199620016 | 2.90E-06 | 0.220609936 | 3.54E-22 | 0.704776919 | 4.43E-10 | 1.248624826 | 0.010100564 |
| K227X | 1 | 1.144336435 | 0.059451 | 1.67E-18 | 0.925914749 | 5.93E-13 | 0.996484945 | 0.519502904 | 1.286773577 | 4.54E-27 | 0.616763627 | 1.32E-68 |
| K227X | 1 | 1.143745076 | 0.045956 | 2.74E-09 | 0.886858198 | 2.37E-11 | 1.238296654 | 2.50E-06 | 0.859773883 | 2.11E-06 | 0.61286106 | 1.50E-28 |
| E71A | 1 | 1.142342975 | 0.073674 | 8.96E-05 | 1.050113037 | 0.031395577 | 0.99713112 | 0.443103611 | 0.914415435 | 0.008752873 | 0.888901995 | 0.000407753 |
| F226R | 6 | 1.142229081 | 0.077605 | 0.003595844 | 1.025669384 | 0.539808237 | 1.053787566 | 2.70E-05 | 1.083540175 | 0.002671282 | 0.935316574 | 0.025009207 |
| E71X | 1 | 1.14095429 | 0.079225 | 0.325060467 | 0.945099853 | 2.49E-06 | 1.258271503 | 0.291666356 | 1.326682537 | 0.451331741 | 0.787777607 | 0.00025965 |
| G118X | 1 | 1.140889262 | 0.051218 | 4.49E-08 | 1.229661831 | 7.73E-32 | 0.822343135 | 1.78E-13 | 0.789131742 | 3.08E-18 | 1.181580529 | 0.000262229 |
| G70X | 1 | 1.140716322 | 0.078818 | 3.11E-06 | 1.183902645 | 0.188595941 | 1.054188619 | 6.51E-27 | 0.870398608 | 0.001254988 | 1.153362191 | 6.05E-15 |
| L174Q | 1 | 1.140703648 | 0.061678 | 4.17E-16 | 1.037514334 | 0.610183406 | 1.13625982 | 0.00036277 | 1.011093391 | 1.77E-09 | 1.342411694 | 1.41E-13 |
| K227S | 4 | 1.140351172 | 0.078767 | 0.010384737 | 1.13173695 | 0.804296578 | 1.046413909 | 0.114860972 | 0.983664532 | 0.001591199 | 0.929596774 | 1.36E-12 |
| W178X | 1 | 1.139334627 | 0.068119 | 3.01E-08 | 1.097562377 | 0.336491071 | 0.682619453 | 4.32E-32 | 0.781054887 | 6.63E-12 | 0.912233849 | 3.85E-06 |
| L66F | 4 | 1.138212307 | 0.074823 | 6.08E-08 | 1.130507659 | 0.990838757 | 1.039760936 | 0.001919507 | 1.065980956 | 0.000602405 | 0.933387762 | 0.000111352 |
| K227X | 1 | 1.137319127 | 0.0594 | 0.244793698 | 1.184077392 | 1.26E-08 | 0.984537197 | 4.49E-06 | 0.937490875 | 1.33E-13 | 1.088823432 | 0.374569158 |
| K227G | 5 | 1.137048571 | 0.091518 | 0.053618724 | 1.02386546 | 0.433140327 | 0.935598543 | 3.33E-06 | 1.061439966 | 0.004639976 | 0.886867227 | 0.0005938 |
| F226A | 6 | 1.13677555 | 0.082566 | 1.41E-07 | 1.056122467 | 0.005469157 | 0.997811913 | 0.218236109 | 1.044394982 | 0.000742255 | 1.057948508 | 0.000206254 |
| K227D | 3 | 1.13604916 | 0.090172 | 0.010932491 | 0.917220805 | 8.98E-11 | 0.939594624 | 2.88E-09 | 1.098942229 | 4.91E-05 | 0.685977896 | 2.05E-29 |
| E71X | 1 | 1.135470338 | 0.080392 | 3.40E-13 | 1.046633309 | 0.005457266 | 1.119099221 | 0.288294035 | 1.142515586 | 0.005934171 | 0.843167641 | 4.49E-07 |
| L66R | 4 | 1.134724221 | 0.070967 | 1.00E-10 | 1.135705672 | 0.9048184 | 1.045146514 | 0.026923364 | 0.961936255 | 0.001290242 | 1.169759147 | 1.03E-06 |
| C138T | 5 | 1.134553438 | 0.0821 | 0.000256823 | 1.095228034 | 0.615191581 | 0.701406547 | 4.52E-24 | 0.838095618 | 1.23E-06 | 1.221741201 | 4.34E-08 |
| F226V | 5 | 1.131671804 | 0.084216 | 5.16E-11 | 1.047621322 | 0.755528333 | 0.912729091 | 0.543958129 | 1.06192283 | 0.082434355 | 1.009134895 | 9.14E-06 |
| F226X | 1 | 1.131591441 | 0.055632 | 1.86E-14 | 0.903092159 | 7.98E-18 | 0.757254564 | 1.17E-22 | 1.038873169 | 0.294436132 | 0.682602996 | 6.77E-68 |
| E71X | 1 | 1.131345176 | 0.078558 | 4.61E-08 | 0.944333617 | 2.33E-08 | 1.474759106 | 0.014872228 | 1.31402092 | 0.099110487 | 0.835029596 | 0.259834328 |
| R72X | 1 | 1.128144077 | 0.062775 | 1.58E-05 | 1.05579773 | 0.744219304 | 1.04194556 | 0.550318128 | 0.789830217 | 0.002214952 | 0.772235866 | 1.50E-09 |
| T67X | 1 | 1.128058072 | 0.052432 | 1.15E-21 | 1.174464545 | 2.84E-41 | 1.041626757 | 0.215079814 | 0.909622252 | 0.000142842 | 1.052043436 | 0.000791481 |
| V196X | 1 | 1.127437378 | 0.065439 | 4.36E-09 | 0.923839477 | 3.71E-05 | 0.990708398 | 0.985578482 | 1.030622726 | 0.873920152 | 0.65879357 | 1.78E-30 |
| Y146X | 1 | 1.127201143 | 0.08494 | 9.47E-15 | 1.009884467 | 0.145272933 | 1.223229582 | 1.98E-15 | 1.07770524 | 0.001288013 | 0.922942752 | 6.36E-18 |
| L174S | 9 | 1.126833212 | 0.067556 | 1.27E-12 | 1.013193967 | 0.777419353 | 1.054944159 | 0.00018049 | 0.938449211 | 0.032654786 | 1.283756071 | 1.16E-07 |
| L66D | 1 | 1.125699311 | 0.064851 | 0.516778152 | 1.14472937 | 0.031352027 | 1.236161322 | 8.18E-23 | 0.902898928 | 2.19E-17 | 1.105954274 | 0.004829324 |
| N69X | 1 | 1.12569224 | 0.064136 | 9.75E-11 | 1.247892412 | 3.23E-19 | 1.044898038 | 0.140840106 | 0.961402248 | 0.07897616 | 1.181949943 | 0.000443418 |
| Y63X | 1 | 1.125222607 | 0.048844 | 5.90E-15 | 1.144794995 | 2.19E-35 | 0.936048028 | 0.006879925 | 1.048046169 | 0.43758193 | 1.006721043 | 0.251789681 |
| V61X | 1 | 1.124513886 | 0.069493 | 9.96E-17 | 1.333704451 | 0.001850344 | 1.041084747 | 0.078119913 | 0.902679842 | 6.69E-05 | 1.44422058 | 1.03E-51 |
| N69X | 1 | 1.123237164 | 0.063996 | 0.264309797 | 1.21421734 | 1.84E-10 | 1.138335673 | 2.61E-05 | 1.06805193 | 0.272471169 | 1.186538539 | 4.47E-10 |
| L144X | 1 | 1.122633373 | 0.063218 | 1.34E-14 | 1.012747691 | 0.468046375 | 1.006991126 | 4.68E-06 | 1.064907813 | 7.68E-13 | 0.825485083 | 2.80E-39 |
| E71X | 1 | 1.11875256 | 0.077684 | 3.76E-05 | 1.126139151 | 0.090151198 | 1.220505506 | 0.020303449 | 0.980981157 | 1.12E-08 | 1.159113173 | 0.038398701 |
| L174L | 1 | 1.118489615 | 0.065444 | 0.000191603 | 0.913201939 | 5.85E-05 | 1.14332315 | 0.097626548 | 0.848539359 | 0.000595795 | 0.977929627 | 0.296175623 |
| T67X | 1 | 1.118356767 | 0.072288 | 5.37E-12 | 0.969648818 | 0.037308999 | 1.023080713 | 0.941796719 | 0.91090515 | 0.0010865 | 0.758775939 | 2.60E-44 |
| F226X | 1 | 1.118333434 | 0.090012 | 1.73E-05 | 1.012407368 | 0.181791628 | 0.784530873 | 3.30E-07 | 0.937697648 | 0.999620939 | 0.846598084 | 0.000118725 |
| C138G | 4 | 1.117899726 | 0.086782 | 1.31E-19 | 1.034388772 | 0.545945102 | 0.763745766 | 8.81E-06 | 0.855018381 | 2.98E-06 | 1.084389884 | 3.11E-05 |
| M114V | 4 | 1.117233192 | 0.070323 | 0.155981646 | 1.055904379 | 0.312345911 | 0.501049831 | 3.54E-15 | 0.707514097 | 4.92E-12 | 0.902734238 | 0.011409909 |
| N69X | 1 | 1.116807535 | 0.08441 | 1.53E-19 | 1.030190407 | 2.96E-05 | 1.305874819 | 0.000452192 | 1.090401052 | 0.462618197 | 0.808189576 | 4.73E-11 |
| E71X | 1 | 1.116134964 | 0.067864 | 0.004869314 | 1.085626877 | 0.000280832 | 0.994791863 | 0.257411349 | 0.945985975 | 0.355999899 | 0.969432884 | 0.139376922 |
| Q64X | 1 | 1.114888576 | 0.056365 | 4.75E-11 | 1.083081387 | 1.13E-08 | 0.718566252 | 4.12E-35 | 0.794022346 | 7.06E-14 | 0.848795789 | 1.33E-07 |
| L174R | 1 | 1.114607824 | 0.061213 | 0.047826916 | 1.312214601 | 9.12E-08 | 0.646588634 | 4.75E-10 | 0.62059153 | 2.33E-13 | 0.875986039 | 0.867883848 |
| R78X | 1 | 1.114604945 | 0.07864 | 0.122193419 | 1.14735281 | 1.90E-29 | 0.98421936 | 0.00593679 | 0.991530564 | 0.000698077 | 1.250818318 | 1.38E-14 |
| A122X | 1 | 1.114468364 | 0.059832 | 0.33824872 | 0.970888344 | 0.060198368 | 1.498205499 | 4.90E-22 | 1.361198937 | 3.55E-06 | 0.907770837 | 0.18826154 |
| F226N | 3 | 1.112881958 | 0.092224 | 6.12E-18 | 0.991016572 | 0.110013071 | 1.198580787 | 4.13E-10 | 1.084370815 | 4.51E-09 | 1.139540831 | 8.04E-09 |
| C138L | 5 | 1.112754974 | 0.091007 | 0.000231603 | 1.334931917 | 0.439515589 | 0.416985873 | 2.53E-34 | 0.59161614 | 1.44E-15 | 1.72254877 | 2.77E-18 |
| V61X | 1 | 1.112219168 | 0.068233 | 7.33E-12 | 1.198744854 | 0.033415197 | 0.950568805 | 0.018976548 | 0.98780702 | 0.012896473 | 1.181603626 | 5.35E-13 |
| D65R | 3 | 1.112005107 | 0.046629 | 0.062907025 | 1.098963159 | 0.543240652 | 1.037076947 | 0.072781812 | 0.971543273 | 0.246423792 | 1.020076686 | 0.690751403 |
| G70S | 5 | 1.110930463 | 0.080034 | 0.00880932 | 1.301832946 | 0.041928906 | 1.097940197 | 0.000569692 | 1.156738997 | 8.37E-12 | 0.882403053 | 6.78E-10 |
| I228L | 8 | 1.110555804 | 0.08914 | 4.42E-05 | 1.126434175 | 0.756191429 | 1.13048593 | 0.000310571 | 0.972177804 | 0.12754054 | 1.011211118 | 1.75E-10 |
| E71Q | 1 | 1.109439349 | 0.074798 | 9.76E-07 | 0.964431874 | 0.026169049 | 1.034179572 | 3.00E-08 | 1.140244184 | 6.86E-06 | 0.923096544 | 4.78E-07 |
| P182Q | 1 | 1.107833413 | 0.083248 | 2.18E-05 | 0.855111593 | 4.47E-19 | 1.112072066 | 0.012670666 | 0.988171152 | 0.49250935 | 0.62638923 | 1.86E-42 |
| K227Q | 4 | 1.106489156 | 0.081245 | 0.009621858 | 1.026419818 | 0.439045798 | 1.001142777 | 0.148385851 | 1.033035433 | 0.068063137 | 0.95170423 | 0.001265373 |
| K227R | 4 | 1.105848116 | 0.091121 | 0.000363347 | 1.18303693 | 0.183690391 | 1.009668376 | 0.003714358 | 0.960549156 | 0.060089888 | 0.964495629 | 5.41E-10 |
| L144C | 4 | 1.105717028 | 0.057503 | 2.47E-23 | 1.351769605 | 4.68E-09 | 0.54762325 | 7.38E-155 | 0.74071878 | 5.99E-112 | 1.495137665 | 2.75E-112 |
| N69X | 1 | 1.105373126 | 0.062979 | 5.72E-07 | 1.201232182 | 2.02E-07 | 1.094127032 | 0.001546771 | 1.193858549 | 0.009390348 | 1.153439053 | 0.0290755 |
| L174M | 3 | 1.104629941 | 0.073126 | 6.49E-09 | 1.048514911 | 0.044479079 | 1.266094345 | 3.77E-08 | 1.083739895 | 0.0017659 | 1.072143097 | 0.010880531 |
| L66X | 1 | 1.103061395 | 0.054985 | 4.18E-14 | 1.081362212 | 5.95E-30 | 1.005086236 | 0.662060584 | 0.873234459 | 9.39E-12 | 1.127697017 | 1.46E-07 |
| Q64X | 1 | 1.102470171 | 0.055737 | 0.134312687 | 0.764683467 | 1.31E-21 | 0.71774211 | 1.94E-05 | 1.051417366 | 0.626439669 | 0.511895191 | 4.53E-27 |
| L144X | 1 | 1.10238975 | 0.061568 | 1.40E-15 | 1.115999421 | 4.88E-36 | 0.890698809 | 8.91E-11 | 1.094928151 | 4.88E-06 | 1.003693409 | 0.962283701 |
| C218V | 3 | 1.100497988 | 0.056078 | 1.60E-06 | 1.062601814 | 0.173971798 | 0.84823683 | 2.46E-10 | 0.968513058 | 0.096603473 | 0.903216432 | 0.000131426 |
| V61X | 1 | 1.100277642 | 0.067501 | 8.09E-12 | 1.17223024 | 0.377573046 | 1.021057206 | 0.118178107 | 0.929144683 | 0.014884638 | 1.052808061 | 1.29E-25 |
| V61X | 1 | 1.100095183 | 0.061365 | 0.438197785 | 1.387320958 | 7.78E-39 | 1.060938455 | 0.063399454 | 0.912309666 | 0.044757951 | 1.415636871 | 1.49E-22 |
| L93X | 1 | 1.099452453 | 0.055461 | 3.97E-08 | 1.107044647 | 0.421963624 | 1.058417961 | 0.13540147 | 1.036171489 | 0.134033126 | 1.115516181 | 0.262656191 |
| V61X | 1 | 1.098843118 | 0.067413 | 4.38E-11 | 1.074873941 | 0.225718133 | 1.061686002 | 2.33E-07 | 1.096549694 | 5.48E-07 | 0.992043862 | 0.156389362 |

| | | | | | | | | | | | | |
|---|---|---|---|---|---|---|---|---|---|---|---|---|
| A122X | 1 | 1.09651944 | 0.084396 | 0.00709223 | 1.131694908 | 4.24E-14 | 0.783910638 | 1.24E-21 | 0.805767885 | 0.005991371 | 1.16995116 | 4.24E-05 |
| L89X | 1 | 1.095973607 | 0.074903 | 0.436038033 | 1.079743762 | 0.738002895 | 1.044991862 | 0.499712407 | 1.172937573 | 5.12E-08 | 1.042282424 | 0.007040164 |
| T67X | 1 | 1.095054262 | 0.050898 | 1.21E-14 | 1.196219585 | 3.06E-43 | 0.98439864 | 0.34027941 | 0.925648945 | 0.001527439 | 1.134444342 | 3.06E-08 |
| C138X | 1 | 1.094992411 | 0.091095 | 3.52E-07 | 1.127693077 | 2.62E-09 | 1.224277406 | 6.78E-05 | 1.00310243 | 9.84E-06 | 0.868057403 | 1.67E-40 |
| S189V | 1 | 1.094648413 | 0.052691 | 0.013816905 | 0.963178048 | 0.757895344 | 1.0962989 | 0.019898846 | 1.196739022 | 4.23E-05 | 0.929116777 | 0.040599 |
| T145X | 1 | 1.093323933 | 0.07067 | 1.36E-10 | 1.309243703 | 9.54E-66 | 0.359027115 | 2.35E-97 | 0.779904174 | 3.57E-19 | 1.276387616 | 2.05E-21 |
| I228P | 6 | 1.092188101 | 0.070088 | 2.23E-08 | 0.963855329 | 0.002658877 | 1.24922281 | 0.000364578 | 1.136910769 | 0.002704283 | 0.899917217 | 7.29E-05 |
| A210X | 1 | 1.091872011 | 0.072151 | 6.17E-07 | 1.411333453 | 0.394759434 | 0.910363399 | 0.005990092 | 0.873713446 | 0.244415711 | 1.401233286 | 0.000436194 |
| I228R | 7 | 1.091407366 | 0.085074 | 7.16E-05 | 1.232293566 | 0.647646247 | 1.162591317 | 1.31E-09 | 0.946135422 | 0.236368622 | 1.012734903 | 9.61E-09 |
| G229I | 4 | 1.0912567 | 0.089488 | 0.000129336 | 1.475002025 | 0.642640783 | 0.939625897 | 8.25E-11 | 0.966671032 | 0.053067796 | 1.262335275 | 8.79E-22 |
| Q64R | 2 | 1.08827185 | 0.051632 | 3.54E-07 | 1.132387946 | 0.145027295 | 1.030762205 | 0.33563761 | 1.018773849 | 0.624437088 | 0.964183235 | 0.001007571 |
| R72F | 1 | 1.087467053 | 0.063613 | 0.000380244 | 0.980449046 | 0.191208507 | 0.782784576 | 0.014936872 | 1.444052973 | 7.74E-15 | 0.749257949 | 4.37E-12 |
| K227E | 5 | 1.087456206 | 0.069001 | 0.002382692 | 1.021322616 | 0.03518045 | 1.123575278 | 0.028379802 | 0.979693003 | 0.129778229 | 0.828182535 | 1.73E-09 |
| Y146X | 1 | 1.087252926 | 0.045838 | 0.244105835 | 1.035132986 | 0.591482025 | 1.106410484 | 0.001870857 | 1.019781918 | 0.390946018 | 0.971036842 | 0.110426842 |
| I48C | 1 | 1.086772263 | 0.071814 | 0.00014644 | 1.053717674 | 0.105982436 | 1.066100931 | 0.334247879 | 0.982131789 | 0.040273489 | 1.0008836 | 0.019531603 |
| D207X | 1 | 1.086704158 | 0.062918 | 0.024352445 | 1.400109926 | 0.000761185 | 0.959000854 | 0.000406858 | 0.948979093 | 0.001075463 | 1.135504417 | 3.59E-07 |
| G70X | 1 | 1.086349789 | 0.082376 | 9.86E-05 | 1.01961233 | 0.357296085 | 1.155732478 | 9.02E-10 | 1.157930399 | 1.24E-23 | 0.99401846 | 4.12E-08 |
| F134I | 1 | 1.084082695 | 0.063662 | 0.044842856 | 0.992085062 | 0.057364927 | 0.590945882 | 2.55E-06 | 0.931004932 | 0.056099077 | 0.681883829 | 5.24E-09 |
| T67X | 1 | 1.083829284 | 0.062361 | 1.26E-05 | 1.18606984 | 3.40E-06 | 0.819326908 | 0.163264953 | 0.989476718 | 0.07453965 | 1.287710802 | 0.25854121 |
| K227N | 6 | 1.082855478 | 0.082047 | 0.01690893 | 0.994930069 | 0.887305302 | 1.069084603 | 0.000432376 | 1.066036126 | 4.96E-05 | 0.829759111 | 4.44E-11 |
| T145X | 1 | 1.082644171 | 0.054426 | 5.60E-08 | 1.223327143 | 2.88E-19 | 1.092438956 | 1.05E-05 | 0.912849014 | 0.00192904 | 1.511510652 | 7.11E-89 |
| L66K | 1 | 1.082585777 | 0.057948 | 2.66E-10 | 1.26120229 | 2.67E-09 | 1.143415257 | 1.85E-06 | 0.913819583 | 3.42E-08 | 1.434750507 | 5.75E-49 |
| Y181X | 1 | 1.082151949 | 0.054924 | 1.26E-12 | 1.065837098 | 4.11E-17 | 0.981437758 | 0.36333042 | 1.12560657 | 6.68E-08 | 1.068107483 | 0.763026229 |
| L66X | 1 | 1.080871779 | 0.070836 | 0.011578495 | 1.102307454 | 0.027109179 | 1.169588399 | 9.17E-10 | 1.048392681 | 0.031387674 | 1.033254178 | 5.23E-06 |
| L66X | 1 | 1.078881876 | 0.06111 | 8.62E-09 | 1.148547313 | 0.000254368 | 0.985327798 | 8.66E-13 | 1.016878193 | 9.68E-15 | 1.211301915 | 9.26E-18 |
| A55X | 1 | 1.078658679 | 0.059119 | 4.88E-07 | 1.174329905 | 0.000597754 | 0.978812885 | 0.130067707 | 0.826228981 | 7.81E-17 | 1.349063094 | 3.92E-13 |
| S189X | 1 | 1.078591604 | 0.06662 | 1.03E-09 | 0.88986964 | 2.46E-32 | 1.236133533 | 5.44E-13 | 1.371510346 | 1.28E-45 | 0.676108346 | 1.35E-100 |
| Y146X | 1 | 1.078164335 | 0.058035 | 0.128497748 | 1.197182634 | 1.22E-06 | 1.154379282 | 1.36E-05 | 1.063071731 | 0.438586907 | 1.049360891 | 0.005527672 |
| L174X | 1 | 1.077553434 | 0.055727 | 0.289767601 | 0.951495008 | 0.094621977 | 0.98245407 | 0.136781804 | 0.778203852 | 1.69E-08 | 0.964075525 | 0.286061335 |
| E71M | 1 | 1.077399485 | 0.071538 | 0.004101925 | 0.982122615 | 0.005289983 | 1.089865599 | 2.97E-07 | 1.027608123 | 0.00013002 | 0.794378774 | 4.11E-10 |
| Q64X | 1 | 1.076724843 | 0.051085 | 4.48E-08 | 0.965176326 | 0.018553783 | 0.890218179 | 0.008361621 | 1.026902096 | 0.812928404 | 0.81284126 | 8.93E-16 |
| Y146X | 1 | 1.076184382 | 0.057517 | 0.061376333 | 1.234964371 | 5.83E-09 | 1.22649351 | 9.16E-08 | 1.034684433 | 0.202289478 | 1.388565589 | 0.00044409 |
| C218L | 1 | 1.075196042 | 0.057442 | 0.06685525 | 0.833128241 | 1.68E-11 | 0.614461802 | 0.000251732 | 2.872071254 | 4.01E-19 | 0.542634915 | 1.55E-23 |
| D65X | 1 | 1.074888831 | 0.080795 | 0.000954494 | 0.98279373 | 0.027721727 | 1.050670646 | 0.060990933 | 1.047345376 | 4.39E-06 | 0.954904557 | 0.013356587 |
| D65K | 1 | 1.072642591 | 0.044979 | 7.65E-05 | 1.040204189 | 0.006122919 | 1.000168467 | 0.789479542 | 1.127814182 | 8.70E-09 | 1.032453803 | 0.487281421 |
| F226X | 1 | 1.071244865 | 0.05771 | 1.49E-06 | 0.855901776 | 3.21E-52 | 1.052918731 | 0.523930055 | 0.996951165 | 0.928061132 | 0.595381032 | 2.81E-101 |
| L66H | 2 | 1.070964476 | 0.069047 | 0.003040407 | 1.2346774 | 5.55E-06 | 0.96564196 | 0.008940688 | 0.950078144 | 1.41E-05 | 1.416500588 | 3.76E-30 |
| G70A | 4 | 1.070272562 | 0.081199 | 0.000152652 | 0.979019223 | 0.016223954 | 1.210438076 | 2.62E-07 | 1.059823858 | 0.031487535 | 1.024106849 | 0.007231477 |
| E71X | 1 | 1.07007713 | 0.071839 | 0.162975046 | 1.012644457 | 0.222919918 | 1.034834002 | 3.93E-08 | 1.19083199 | 1.47E-09 | 0.947085498 | 0.009911924 |
| F134Y | 6 | 1.069588947 | 0.063087 | 1.92E-07 | 1.042774565 | 0.975728287 | 1.092870016 | 1.95E-08 | 1.207270145 | 6.51E-21 | 1.014310108 | 1.53E-34 |
| N69X | 1 | 1.068908678 | 0.071492 | 0.000571845 | 1.046356517 | 1.66E-05 | 1.007286784 | 0.008244527 | 0.929355968 | 0.072076249 | 1.044976324 | 0.010886478 |
| V61X | 1 | 1.068127779 | 0.066009 | 2.65E-07 | 1.129369584 | 0.008749593 | 1.039013999 | 0.011919729 | 0.902656779 | 0.061462932 | 1.401472472 | 1.02E-20 |
| G229A | 3 | 1.067111245 | 0.072753 | 2.68E-17 | 0.992777102 | 0.575285795 | 1.27732067 | 4.69E-07 | 1.068275304 | 1.39E-11 | 1.346682995 | 1.59E-31 |
| P182X | 1 | 1.066284073 | 0.05491 | 0.000521251 | 1.312903458 | 1.06E-20 | 0.990251026 | 0.180190952 | 0.849868207 | 1.69E-06 | 1.331285845 | 1.74E-33 |
| N69X | 1 | 1.066280021 | 0.060414 | 0.001571974 | 1.115639628 | 0.001409354 | 1.258602408 | 1.20E-07 | 1.221723117 | 0.001145876 | 1.127019798 | 0.004779589 |
| V61X | 1 | 1.066256105 | 0.065413 | 0.258663377 | 1.075372365 | 0.000564241 | 0.7885262 | 0.061518318 | 0.87033514 | 0.065760684 | 0.801370977 | 0.492678409 |
| T145X | 1 | 1.066089621 | 0.06891 | 0.00567063 | 0.956301401 | 0.000465244 | 1.057864677 | 0.085788078 | 0.874651787 | 0.002120101 | 1.241382029 | 4.96E-22 |
| I228H | 3 | 1.066056708 | 0.088344 | 0.000577487 | 1.019118579 | 0.349407795 | 1.217512738 | 0.000269725 | 0.969295224 | 0.003100685 | 1.047472413 | 2.30E-06 |
| C218C | 1 | 1.065204485 | 0.065177 | 0.188078536 | 0.997623379 | 0.569332837 | 0.908325319 | 0.018319718 | 0.875836441 | 0.097694179 | 1.039418944 | 0.129102517 |
| G70V | 4 | 1.063651196 | 0.073493 | 0.010560468 | 0.975342437 | 0.000853846 | 1.212379688 | 0.00489952 | 1.162641146 | 0.152751994 | 0.767478514 | 1.34E-14 |
| Y146X | 1 | 1.06280976 | 0.064405 | 0.004361907 | 1.256048911 | 1.18E-28 | 1.114100294 | 0.232069114 | 0.931404767 | 0.410717409 | 1.426936448 | 3.40E-28 |
| V74X | 1 | 1.062310155 | 0.051013 | 0.117319636 | 0.933705374 | 0.004172306 | 1.006708212 | 0.54796087 | 0.942319758 | 0.165558735 | 0.772823635 | 1.56E-05 |
| S189T | 4 | 1.061015546 | 0.062707 | 0.006711744 | 1.086460495 | 0.270689077 | 1.096015667 | 0.001626429 | 0.994752658 | 4.32E-09 | 1.347224647 | 4.64E-35 |
| C218X | 1 | 1.060757967 | 0.054053 | 0.003564978 | 1.041719201 | 0.115366051 | 0.752043407 | 2.48E-11 | 1.0044856 | 0.20615925 | 0.880610307 | 3.70E-05 |
| I228X | 1 | 1.060509003 | 0.062111 | 0.010110672 | 0.854479543 | 2.56E-10 | 1.170980978 | 0.186074347 | 1.07586795 | 0.157948225 | 0.749242104 | 5.57E-11 |
| N69X | 1 | 1.059713189 | 0.055307 | 0.012728795 | 1.171049479 | 1.48E-35 | 1.236246277 | 8.30E-12 | 1.112060867 | 0.309646192 | 1.308912422 | 9.72E-24 |
| Y146X | 1 | 1.058835997 | 0.062586 | 0.026750544 | 1.226330793 | 1.52E-14 | 1.164948727 | 9.12E-05 | 0.97416673 | 0.390714601 | 1.28610238 | 2.50E-07 |
| G70K | 2 | 1.05645309 | 0.072996 | 0.000831658 | 1.044882994 | 1.83E-05 | 1.008927478 | 2.80E-06 | 1.012282309 | 0.607489151 | 1.170120146 | 4.88E-17 |
| S189X | 1 | 1.056294897 | 0.062428 | 5.80E-10 | 0.888409267 | 3.77E-62 | 1.333635719 | 6.05E-83 | 1.070336024 | 8.49E-10 | 0.697235988 | 1.82E-154 |
| W178X | 1 | 1.05553728 | 0.063435 | 0.111532777 | 1.158326796 | 0.222332328 | 0.520145529 | 1.47E-21 | 0.710629155 | 2.14E-15 | 1.036002734 | 0.072620625 |
| G70E | 2 | 1.055205974 | 0.078695 | 0.32258766 | 1.145289065 | 3.75E-07 | 1.246467075 | 4.06E-06 | 1.103652239 | 0.007967537 | 0.959097033 | 2.16E-08 |
| W178X | 1 | 1.054628871 | 0.061146 | 0.08035787 | 1.021741969 | 0.128264468 | 0.495972021 | 1.47E-12 | 0.788426668 | 1.40E-09 | 0.96291257 | 0.057273839 |
| I228F | 3 | 1.051539734 | 0.066342 | 0.001715828 | 1.074206341 | 0.368243815 | 0.984663644 | 2.64E-07 | 1.02266256 | 0.340772664 | 1.070881886 | 0.002678152 |
| D65X | 1 | 1.051296705 | 0.044084 | 0.017407732 | 1.067567978 | 0.00171128 | 0.796447116 | 2.27E-09 | 0.995983043 | 0.71878645 | 1.050326073 | 0.611510427 |
| Q141F | 2 | 1.05032401 | 0.060185 | 0.451199961 | 1.000015189 | 0.909500361 | 0.935055408 | 0.490779754 | 1.112677239 | 9.22E-05 | 1.652611923 | 0.176524871 |
| L89Q | 1 | 1.049474459 | 0.064989 | 0.067373291 | 1.269833155 | 5.18E-28 | 0.785032558 | 3.67E-09 | 0.641323024 | 1.49E-48 | 1.33913134 | 1.19E-05 |
| A60X | 1 | 1.048416346 | 0.064319 | 0.327462891 | 0.986778986 | 0.125663303 | 0.962902928 | 0.006557757 | 1.011332331 | 0.007802875 | 0.869177228 | 0.029731686 |
| Q64X | 1 | 1.048100689 | 0.049727 | 0.001195987 | 1.035409472 | 0.036680328 | 0.923622863 | 0.000402333 | 1.004117616 | 0.623794236 | 0.960893466 | 0.006488177 |
| L93X | 1 | 1.047968566 | 0.055157 | 0.002496477 | 1.057669549 | 0.001939362 | 1.093716464 | 3.42E-06 | 1.088388704 | 0.00300515 | 1.047875979 | 0.009874091 |
| Y181X | 1 | 1.04649962 | 0.053543 | 0.000142668 | 1.118422289 | 6.34E-39 | 0.926242453 | 1.13E-07 | 1.003540951 | 0.572063128 | 1.049533406 | 1.34E-05 |
| C218X | 1 | 1.046166171 | 0.055891 | 0.073715478 | 0.936205556 | 0.098516411 | 1.018594055 | 0.281937206 | 1.086596294 | 0.184137272 | 0.769813235 | 7.87E-06 |
| A210X | 1 | 1.046047973 | 0.060495 | 0.203272402 | 1.042006109 | 0.009643595 | 0.982072695 | 0.328081678 | 0.90340324 | 0.199932199 | 1.026071801 | 1.84E-06 |
| C138V | 7 | 1.045843832 | 0.086215 | 1.13E-22 | 1.16061604 | 0.722026102 | 0.533146628 | 4.14E-36 | 0.871474952 | 1.07E-07 | 1.652261626 | 4.45E-36 |
| E71X | 1 | 1.045198436 | 0.066962 | 0.412563993 | 1.012228339 | 0.476175945 | 1.068595505 | 0.000179388 | 1.050382393 | 7.47E-15 | 0.988468023 | 0.451964862 |
| C218X | 1 | 1.044706652 | 0.043834 | 0.073059971 | 1.069818741 | 0.002811483 | 0.735130613 | 7.19E-26 | 0.883001436 | 7.39E-05 | 1.023597893 | 0.648466082 |
| F226M | 3 | 1.044601554 | 0.056131 | 0.011312951 | 1.02855906 | 0.490081 | 0.918821889 | 3.61E-10 | 0.987835573 | 0.112002572 | 0.963948568 | 0.008745891 |
| Q64X | 1 | 1.044366943 | 0.071458 | 0.180656555 | 1.040851382 | 0.048609095 | 0.821319678 | 2.76E-06 | 0.970733558 | 2.35E-10 | 1.015615891 | 0.001780967 |
| D65X | 1 | 1.043948587 | 0.062969 | 0.002902687 | 1.078609403 | 1.26E-18 | 1.19554585 | 0.001379068 | 0.99582727 | 1.05E-07 | 1.323326282 | 3.51E-16 |
| L93X | 1 | 1.043196705 | 0.052624 | 0.33167683 | 1.113916562 | 0.012845597 | 0.84671595 | 0.005118987 | 1.110602311 | 0.106304807 | 1.010455645 | 0.001098926 |
| G118X | 1 | 1.04156719 | 0.057948 | 0.003581151 | 1.029186764 | 0.004168902 | 1.122773917 | 5.50E-05 | 0.968842852 | 0.107925401 | 1.284148792 | 0.000549723 |
| D65T | 1 | 1.041100633 | 0.040503 | 0.114622801 | 1.038057662 | 0.23814618 | 0.897662644 | 0.002126323 | 1.040911237 | 0.675528587 | 1.116530845 | 0.406906119 |
| Y146X | 1 | 1.040947265 | 0.058065 | 0.111456425 | 1.283753388 | 1.82E-06 | 1.042845304 | 0.139933235 | 0.979217951 | 0.46325344 | 1.433246261 | 2.70E-05 |
| L66V | 8 | 1.040895068 | 0.067747 | 0.012560975 | 1.08499552 | 0.00680871 | 1.050560154 | 0.000448262 | 0.9818301 | 7.12E-07 | 1.193146148 | 7.12E-13 |

| | | | | | | | | | | | | |
|---|---|---|---|---|---|---|---|---|---|---|---|---|
| G229X | 1 | 1.039869438 | 0.073329 | 0.129579875 | 1.032800351 | 0.000816433 | 1.420084943 | 2.04E-49 | 1.051531165 | 0.007099103 | 1.136778198 | 2.33E-10 |
| G229X | 1 | 1.039483942 | 0.070869 | 0.085483657 | 0.90001007 | 4.35E-05 | 1.023884804 | 0.072864426 | 0.996346349 | 0.25143133 | 0.956975262 | 0.298403839 |
| G70W | 3 | 1.037857695 | 0.078739 | 0.56317696 | 0.859791484 | 9.99E-17 | 1.13754944 | 0.073272378 | 1.244195405 | 2.95E-06 | 0.611149316 | 3.66E-31 |
| Y146X | 1 | 1.037522044 | 0.078183 | 0.091652843 | 0.925029794 | 4.23E-07 | 1.163231375 | 1.18E-06 | 1.044154889 | 0.181159026 | 0.954449172 | 0.134318906 |
| T145X | 1 | 1.037329189 | 0.067051 | 0.003522022 | 1.036044255 | 0.001359569 | 1.709666918 | 3.80E-14 | 1.713957127 | 0.001894793 | 1.40467815 | 0.434160321 |
| Y146S | 1 | 1.036900149 | 0.078097 | 0.04324888 | 1.066167167 | 0.456684939 | 0.991798882 | 0.256231034 | 0.996375111 | 0.450258707 | 0.866945275 | 0.000817452 |
| G70X | 1 | 1.035911772 | 0.074629 | 0.061021593 | 0.937134001 | 1.25E-05 | 2.197761518 | 3.99E-39 | 1.747747924 | 5.95E-18 | 0.832176952 | 7.89E-09 |
| F226C | 2 | 1.035320432 | 0.070368 | 0.016107706 | 1.036546891 | 0.5052549 | 1.179047205 | 3.90E-07 | 1.025087261 | 0.485473067 | 0.996457452 | 5.06E-05 |
| L66P | 1 | 1.034533441 | 0.061414 | 4.35E-09 | 1.035624741 | 0.001660023 | 1.041133302 | 0.000490167 | 0.93659307 | 7.39E-05 | 1.011170101 | 0.415193858 |
| T145X | 1 | 1.034470199 | 0.066866 | 0.000641272 | 1.192608116 | 1.73E-87 | 1.05896994 | 0.035517223 | 1.03275862 | 0.032546853 | 1.283712383 | 7.36E-53 |
| A55X | 1 | 1.034246577 | 0.068343 | 0.222780045 | 1.069033062 | 0.003315019 | 0.947842576 | 0.216344839 | 0.87436757 | 0.387634781 | 1.068712272 | 0.02677501 |
| L89D | 2 | 1.034090689 | 0.076467 | 0.299590101 | 1.229879415 | 1.85E-05 | 0.377726791 | 6.08E-18 | 0.98384949 | 6.41E-10 | 0.964781718 | 0.000112482 |
| C138A | 1 | 1.03353961 | 0.085983 | 0.012242937 | 0.952054205 | 0.001610822 | 0.742859863 | 0.001413273 | 1.059529376 | 0.019160396 | 0.883063836 | 0.10048815 |
| G229T | 10 | 1.032700648 | 0.074809 | 6.17E-09 | 1.131557955 | 0.334999449 | 1.238472981 | 2.68E-05 | 1.003667115 | 0.065994918 | 1.272966213 | 9.33E-23 |
| V196X | 1 | 1.032368682 | 0.051517 | 0.010629183 | 1.054653799 | 0.894393835 | 1.098795577 | 0.116166523 | 0.996039882 | 0.219282542 | 1.08486674 | 0.041503935 |
| L174X | 1 | 1.031764511 | 0.053359 | 0.457965761 | 0.891927951 | 0.000149712 | 1.055402574 | 0.712414355 | 0.849856104 | 0.001877055 | 0.820894838 | 0.001023645 |
| G229H | 3 | 1.031327848 | 0.058303 | 1.59E-05 | 1.05210598 | 0.317227992 | 1.014589661 | 0.020817623 | 0.798870872 | 7.08E-27 | 1.194553913 | 5.43E-09 |
| V61I | 1 | 1.031224216 | 0.061404 | 0.160134648 | 1.064341507 | 2.30E-09 | 1.07900036 | 0.36959267 | 1.116752681 | 1.69E-06 | 0.884795894 | 2.57E-07 |
| T145X | 1 | 1.031026284 | 0.066643 | 0.674616613 | 1.12057035 | 5.28E-21 | 0.631413879 | 1.51E-46 | 0.805326106 | 7.05E-21 | 1.245359219 | 7.90E-26 |
| E71X | 1 | 1.030794634 | 0.072604 | 0.332044531 | 1.052191953 | 0.097266145 | 1.072778859 | 0.252509886 | 1.017605912 | 0.323884997 | 0.941240354 | 0.363363639 |
| L144X | 1 | 1.029842577 | 0.059156 | 0.000129533 | 1.261542832 | 0.030793875 | 1.055827481 | 3.09E-05 | 0.87630749 | 8.86E-12 | 1.52350704 | 4.34E-32 |
| V74X | 1 | 1.029827673 | 0.042574 | 0.927348265 | 0.95069648 | 0.000184621 | 0.942943562 | 0.000192436 | 0.862504978 | 3.79E-08 | 0.888281328 | 0.000215166 |
| G70T | 1 | 1.029355662 | 0.078094 | 0.063231056 | 1.098879439 | 6.73E-27 | 1.270386564 | 1.66E-23 | 0.944597466 | 9.23E-06 | 1.219853836 | 1.13E-28 |
| I228X | 1 | 1.028692893 | 0.070533 | 0.171967244 | 1.065094219 | 0.325135004 | 0.938250064 | 0.13775372 | 0.8732665 | 0.000324873 | 1.148410723 | 0.124246006 |
| L144T | 3 | 1.026579472 | 0.068867 | 0.041401872 | 1.221036169 | 1.10E-101 | 0.613876116 | 2.93E-173 | 0.848302965 | 1.90E-58 | 1.40912866 | 3.71E-83 |
| A55X | 1 | 1.02652698 | 0.038392 | 4.79E-22 | 0.928196538 | 0.004098652 | 1.299160548 | 4.46E-07 | 1.490919008 | 1.27E-06 | 1.298920463 | 3.39E-15 |
| L144X | 1 | 1.02600486 | 0.057302 | 0.132163119 | 1.108790669 | 3.98E-46 | 1.001522347 | 0.949364007 | 1.042117245 | 0.003501676 | 1.194136372 | 5.49E-25 |
| Y146X | 1 | 1.02573985 | 0.077257 | 0.010954121 | 0.94089902 | 2.32E-09 | 1.158984737 | 0.003940147 | 1.067230145 | 0.344650494 | 0.989666341 | 9.21E-14 |
| I228W | 3 | 1.025691256 | 0.073478 | 0.013975228 | 0.914284401 | 9.22E-08 | 1.082541274 | 0.000309956 | 0.993813459 | 0.051464085 | 0.951918529 | 0.010384261 |
| F226X | 1 | 1.025681398 | 0.045861 | 0.10897952 | 0.966872987 | 0.000866981 | 0.916964265 | 0.000117023 | 1.009762454 | 0.564931171 | 0.906273459 | 0.001040531 |
| Y63X | 1 | 1.02512465 | 0.046327 | 0.007299237 | 1.107575758 | 9.97E-17 | 1.007903868 | 0.998757979 | 1.195436699 | 1.18E-17 | 1.175182344 | 1.22E-08 |
| Q64X | 1 | 1.025028329 | 0.048632 | 0.340371409 | 0.894416338 | 4.12E-09 | 0.846751883 | 0.001186273 | 1.05318893 | 0.999796739 | 0.764778059 | 2.61E-10 |
| Q141X | 1 | 1.024389705 | 0.073361 | 0.005944987 | 1.067443036 | 0.011138504 | 0.909352043 | 0.016713182 | 0.904313068 | 3.63E-05 | 1.231605309 | 0.465104899 |
| T145X | 1 | 1.023432905 | 0.078362 | 0.381704992 | 0.986344401 | 0.450676635 | 0.932667572 | 0.001022355 | 1.020857582 | 0.02671607 | 1.086925117 | 0.002755944 |
| G70D | 2 | 1.023212749 | 0.060742 | 0.000666603 | 1.066733997 | 4.06E-10 | 0.766650331 | 5.88E-46 | 0.817530218 | 4.76E-31 | 1.071479005 | 4.06E-07 |
| D65I | 2 | 1.020482002 | 0.042791 | 0.448704785 | 0.873733596 | 6.79E-10 | 1.080568499 | 0.413890966 | 1.152375969 | 0.043417251 | 0.766701515 | 2.05E-06 |
| T145X | 1 | 1.019642258 | 0.065908 | 0.899141364 | 1.016107687 | 0.016031476 | 1.12783087 | 1.63E-22 | 1.197184383 | 8.43E-15 | 1.122049815 | 2.00E-06 |
| R78X | 1 | 1.019337161 | 0.079052 | 0.496179109 | 1.063882886 | 7.04E-07 | 1.257961015 | 2.40E-29 | 0.988821172 | 0.255517142 | 1.177599903 | 2.29E-09 |
| T67X | 1 | 1.018808513 | 0.05862 | 0.232891876 | 1.12285521 | 2.15E-12 | 1.081813082 | 0.233683289 | 1.093573414 | 0.245444109 | 1.175182538 | 9.26E-11 |
| E71X | 1 | 1.018162286 | 0.070699 | 0.281116241 | 0.964662185 | 0.506525929 | 1.205741908 | 3.74E-06 | 1.112585034 | 0.022214161 | 0.91921966 | 0.467643836 |
| V196X | 1 | 1.016574064 | 0.07639 | 0.016333933 | 1.069537503 | 1.62E-07 | 1.037945627 | 0.27062059 | 0.980027507 | 0.184071729 | 1.024670332 | 0.917645953 |
| L144X | 1 | 1.016302796 | 0.052419 | 0.74871054 | 0.913079237 | 2.10E-15 | 0.859016087 | 1.82E-05 | 0.931866025 | 0.060428784 | 0.856049835 | 5.56E-14 |
| F226T | 8 | 1.016024944 | 0.084183 | 0.043732342 | 1.087663314 | 8.26E-05 | 1.035690701 | 0.002599593 | 1.137691047 | 0.001749082 | 0.809982275 | 1.71E-13 |
| K227X | 1 | 1.015983513 | 0.040822 | 0.082516281 | 1.034091968 | 0.002700015 | 1.170938348 | 1.32E-10 | 0.948796001 | 0.013901619 | 1.046387167 | 0.433662522 |
| G70X | 1 | 1.015977667 | 0.073193 | 0.936076336 | 0.900661579 | 0.023045773 | 1.161742114 | 0.163807778 | 1.327803467 | 0.001221556 | 0.936539021 | 0.332034542 |
| D65C | 2 | 1.013934489 | 0.042517 | 0.12395996 | 0.911604956 | 5.26E-07 | 1.044086515 | 0.501873656 | 1.06069813 | 0.089210586 | 0.918272592 | 0.106012717 |
| L66X | 1 | 1.013445562 | 0.06153 | 0.00020967 | 1.136700162 | 0.487640657 | 1.045610335 | 1.90E-11 | 1.034725975 | 0.000339158 | 1.398000721 | 4.91E-59 |
| P182X | 1 | 1.013193766 | 0.059252 | 0.547319197 | 1.239099514 | 1.06E-33 | 1.061243534 | 0.37975656 | 0.873714627 | 0.00344049 | 1.476249769 | 5.80E-36 |
| L66T | 2 | 1.012026978 | 0.062417 | 3.13E-05 | 1.139401251 | 0.010697254 | 1.108363763 | 5.49E-11 | 0.963806746 | 0.000186429 | 1.315288555 | 3.69E-10 |
| D65X | 1 | 1.011180697 | 0.039339 | 0.312222972 | 1.005481507 | 0.709737536 | 0.916790047 | 0.037156019 | 1.078287001 | 0.267183535 | 1.014436937 | 0.864970646 |
| L89X | 1 | 1.010739612 | 0.059657 | 0.000631876 | 1.25965508 | 0.049335879 | 0.711224799 | 0.400008686 | 0.567014607 | 1.30E-20 | 1.369455699 | 1.37E-07 |
| F226P | 5 | 1.010124483 | 0.077091 | 7.68E-05 | 0.912559362 | 1.04E-13 | 1.019480979 | 0.016509029 | 0.994036922 | 0.09826813 | 0.814363263 | 7.22E-15 |
| L89X | 1 | 1.009437408 | 0.061372 | 0.347348073 | 1.167730924 | 1.28E-06 | 0.81962134 | 0.323475444 | 0.910227701 | 0.004125191 | 1.418397793 | 0.011840559 |
| S189G | 2 | 1.00934513 | 0.06661 | 0.301584524 | 0.831514222 | 6.47E-26 | 1.312344031 | 0.000432858 | 1.143792813 | 0.01226996 | 0.655997115 | 9.78E-24 |
| K227F | 2 | 1.009078226 | 0.050709 | 0.406603918 | 1.084922769 | 2.48E-16 | 1.047216473 | 5.60E-14 | 1.047504162 | 0.068059631 | 1.003496924 | 0.008368538 |
| T145X | 1 | 1.008064735 | 0.065159 | 0.665180447 | 1.107889155 | 0.687044064 | 1.003618582 | 0.18786549 | 0.985372245 | 0.558250502 | 0.955170486 | 9.31E-05 |
| D65X | 1 | 1.007962278 | 0.075088 | 0.071108746 | 1.10789533 | 2.22E-05 | 1.130023908 | 4.68E-08 | 0.957403015 | 6.31E-05 | 1.348120701 | 3.45E-15 |
| K227C | 4 | 1.007940054 | 0.064991 | 0.111537539 | 1.014987423 | 0.146192198 | 1.067616211 | 0.001345579 | 0.998076228 | 0.023434505 | 0.97908049 | 6.69E-15 |
| K227H | 3 | 1.006685369 | 0.087144 | 0.086799976 | 1.057845066 | 0.47963472 | 0.8660006 | 5.98E-06 | 0.876723771 | 5.15E-07 | 1.004689088 | 0.019733598 |
| G70X | 1 | 1.006532426 | 0.072513 | 0.038806379 | 0.907288038 | 3.21E-07 | 1.572438564 | 2.02E-07 | 1.103309377 | 0.022133052 | 0.786606292 | 9.55E-13 |
| I228T | 7 | 1.005463952 | 0.074742 | 4.82E-07 | 1.087654176 | 0.704076665 | 1.107052077 | 6.07E-05 | 0.971166659 | 0.013550178 | 1.163729843 | 2.11E-11 |
| L66X | 1 | 1.004906554 | 0.058253 | 0.388093244 | 1.100432885 | 9.27E-06 | 1.138674425 | 0.134061478 | 1.181673704 | 0.182986159 | 1.365752476 | 2.93E-15 |
| T85X | 1 | 1.004341564 | 0.065977 | 0.227639309 | 1.043937743 | 1.22E-06 | 0.997373634 | 0.098091937 | 0.935165268 | 0.018133518 | 1.238765831 | 1.39E-07 |
| A122X | 1 | 1.003570334 | 0.077242 | 0.471203544 | 1.024439472 | 0.590147947 | 1.054300995 | 0.148588803 | 0.992901254 | 0.078102596 | 1.088802717 | 0.000498588 |
| F226X | 1 | 1.00199424 | 0.055039 | 0.214090968 | 1.063601678 | 1.58E-23 | 0.996456295 | 0.58255758 | 0.969903171 | 0.011644067 | 1.184368696 | 4.78E-31 |
| L144X | 1 | 1.001706016 | 0.056483 | 0.256369419 | 1.132885858 | 2.92E-09 | 1.171086111 | 0.000147044 | 1.056263171 | 0.001178007 | 1.016001921 | 5.96E-05 |
| I228C | 3 | 1.001604046 | 0.06718 | 7.71E-06 | 0.989134809 | 0.20325399 | 1.010114408 | 1.09E-09 | 1.040622725 | 0.000123187 | 1.032016449 | 0.034367937 |
| G70X | 1 | 1.001497923 | 0.075981 | 0.523381986 | 0.849335086 | 1.32E-34 | 1.084968228 | 0.39578809 | 1.101545345 | 0.021115969 | 0.72554513 | 4.67E-31 |
| L66X | 1 | 1.001076147 | 0.059427 | 0.000136377 | 1.090091576 | 4.44E-13 | 0.979637989 | 0.03688416 | 1.029327541 | 7.42E-12 | 1.177650875 | 5.38E-10 |
| L144V | 3 | 1.001041396 | 0.063698 | 0.319975851 | 1.191610915 | 1.18E-08 | 0.74323806 | 4.73E-24 | 0.765957529 | 1.44E-25 | 1.333422668 | 2.47E-66 |
| P182X | 1 | 1.000822089 | 0.059552 | 0.820821517 | 1.370359163 | 2.91E-25 | 1.004098696 | 0.078862091 | 0.853254924 | 5.85E-05 | 1.606864429 | 4.68E-34 |
| T145X | 1 | 1.000640809 | 0.075366 | 0.306911922 | 0.944901299 | 0.002215609 | 0.97898972 | 0.278683871 | 1.020577413 | 0.418130438 | 1.071828847 | 1.26E-10 |
| K227W | 2 | 1.000084938 | 0.051957 | 0.003913574 | 1.140340665 | 7.64E-06 | 0.983784842 | 0.29174166 | 0.973809014 | 0.533909825 | 0.918929785 | 1.97E-09 |
| L93X | 1 | 1.000053299 | 0.052636 | 0.085836638 | 0.936759575 | 1.62E-07 | 0.930181725 | 0.039741805 | 1.117722954 | 0.000125877 | 0.898921338 | 0.001063474 |
| S189X | 1 | 0.999671037 | 0.048119 | 0.672700251 | 0.961190443 | 0.379599746 | 1.191536404 | 6.80E-06 | 1.002190736 | 0.871575862 | 1.017984911 | 0.937831483 |
| G229X | 1 | 0.998997149 | 0.05278 | 0.488688112 | 1.165351323 | 1.40E-06 | 0.989556407 | 0.000881275 | 1.055735436 | 0.006224027 | 1.754316355 | 3.48E-20 |
| P182X | 1 | 0.998028983 | 0.056961 | 0.513636883 | 1.266165391 | 1.90E-10 | 1.031624012 | 0.050220316 | 0.897486535 | 0.021252543 | 1.472401835 | 5.64E-13 |
| G70Y | 3 | 0.997701898 | 0.075654 | 0.063276587 | 0.774920257 | 8.15E-29 | 1.160081659 | 2.05E-06 | 1.416426607 | 2.12E-19 | 0.577692341 | 1.11E-48 |
| L144X | 1 | 0.997345856 | 0.055702 | 0.192258279 | 1.008046546 | 0.372972508 | 0.997865793 | 2.33E-05 | 1.001653304 | 0.000898285 | 0.95754437 | 0.329791883 |
| Y146X | 1 | 0.996823283 | 0.075116 | 0.534391216 | 0.990542463 | 0.442304089 | 1.076407881 | 0.105210839 | 1.025795654 | 0.349912784 | 0.875836946 | 1.22E-10 |
| Y146V | 1 | 0.996044368 | 0.07502 | 0.131419248 | 1.101415979 | 3.20E-14 | 1.09518088 | 0.079261764 | 0.982872126 | 0.007937392 | 1.033807084 | 0.000205575 |
| I228G | 4 | 0.995197449 | 0.082472 | 6.05E-07 | 0.979808347 | 0.15407763 | 1.12873717 | 0.007742299 | 0.978546374 | 1.24E-06 | 1.093164554 | 4.96E-11 |
| L93I | 1 | 0.99511844 | 0.052376 | 0.956841705 | 0.846744679 | 2.71E-23 | 1.273359318 | 5.46E-09 | 1.059613009 | 0.220618926 | 0.688359367 | 6.35E-21 |

| | | | | | | | | | | | | |
|---|---|---|---|---|---|---|---|---|---|---|---|---|
| E71X | 1 | 0.994937108 | 0.06822 | 0.60569434 | 0.930858815 | 1.04E-09 | 1.442647504 | 6.64E-06 | 1.289550637 | 0.113797509 | 0.862951958 | 9.79E-07 |
| T67X | 1 | 0.994117696 | 0.050259 | 0.339721775 | 1.084388834 | 0.0004142 | 1.039122278 | 0.23362596 | 1.031606279 | 0.223011878 | 1.165084485 | 5.09E-06 |
| G70X | 1 | 0.994087584 | 0.068687 | 0.445653218 | 0.85436566 | 2.30E-07 | 1.928827364 | 1.86E-10 | 1.41223966 | 0.002412627 | 0.579446778 | 1.84E-12 |
| F226X | 1 | 0.993805993 | 0.082356 | 0.296100012 | 0.855898713 | 4.34E-14 | 1.185373119 | 4.83E-05 | 1.08503361 | 0.138627118 | 0.911384055 | 5.65E-06 |
| Y181X | 1 | 0.993762988 | 0.050845 | 0.596334136 | 1.02534387 | 0.000689628 | 0.881401003 | 1.20E-15 | 0.889887514 | 3.12E-16 | 1.147083906 | 1.33E-05 |
| L66A | 3 | 0.993366076 | 0.049517 | 5.89E-08 | 1.197711666 | 8.42E-09 | 0.997655719 | 0.024014855 | 0.960244323 | 0.005674677 | 1.386677607 | 2.38E-39 |
| Y181X | 1 | 0.993350192 | 0.047944 | 0.766687788 | 1.033160681 | 2.84E-05 | 1.08831762 | 6.30E-06 | 1.111908739 | 3.52E-09 | 1.063524249 | 0.017735172 |
| T145X | 1 | 0.992636771 | 0.057114 | 0.326543521 | 1.166196197 | 9.84E-06 | 1.05593779 | 0.004124212 | 0.938618614 | 0.001049337 | 1.407384821 | 7.36E-45 |
| T145X | 1 | 0.9916466 | 0.064098 | 0.367050969 | 0.985095089 | 0.297894344 | 0.958265286 | 0.009937778 | 0.95592664 | 0.286472172 | 0.979446436 | 0.002600937 |
| L66X | 1 | 0.990468576 | 0.06561 | 0.040441225 | 1.212255577 | 1.30E-15 | 0.916402749 | 1.12E-09 | 0.902196444 | 4.12E-05 | 1.290367546 | 1.39E-32 |
| G118X | 1 | 0.990219205 | 0.058175 | 0.514725943 | 0.919741506 | 0.00024233 | 1.092738713 | 0.495800589 | 0.928323097 | 0.051111816 | 0.848897013 | 0.001125664 |
| S189X | 1 | 0.989306733 | 0.086374 | 0.334613147 | 0.750086885 | 1.18E-07 | 1.942408751 | 0.000269818 | 1.40314896 | 0.001259578 | 0.55673449 | 4.61E-12 |
| S189X | 1 | 0.989046224 | 0.049896 | 0.017959615 | 0.935804311 | 9.53E-05 | 1.195408314 | 2.35E-17 | 1.037346556 | 0.000668141 | 0.946945805 | 0.868835141 |
| R72X | 1 | 0.989009878 | 0.049164 | 0.09431619 | 1.050398957 | 3.03E-10 | 1.138133603 | 6.28E-16 | 0.988513487 | 0.845615819 | 1.152026112 | 6.36E-14 |
| G70X | 1 | 0.988347304 | 0.074983 | 0.011022792 | 0.907411527 | 4.99E-13 | 1.353052389 | 1.13E-15 | 0.991789583 | 0.340981537 | 0.860104579 | 3.96E-07 |
| I228X | 1 | 0.987512313 | 0.065628 | 0.102610637 | 0.971872864 | 0.031607464 | 1.09929528 | 0.336198429 | 1.023085009 | 0.431546218 | 0.977729797 | 4.50E-07 |
| K227X | 1 | 0.987259752 | 0.039668 | 0.023604107 | 0.969142812 | 0.03370883 | 1.085156898 | 0.022329855 | 0.92660743 | 0.001615072 | 1.132738749 | 0.125554988 |
| C218X | 1 | 0.987135827 | 0.049566 | 0.622849658 | 1.09415815 | 1.42E-07 | 0.747623547 | 8.64E-22 | 0.896808922 | 5.65E-05 | 1.111968639 | 0.000530357 |
| Q64W | 1 | 0.986893347 | 0.082138 | 0.045455983 | 0.827670534 | 4.41E-50 | 1.351374447 | 4.06E-26 | 1.367247068 | 1.75E-36 | 0.768025391 | 3.42E-56 |
| C138F | 3 | 0.986712178 | 0.082087 | 8.04E-08 | 1.048688946 | 0.037952801 | 0.728708611 | 3.69E-05 | 0.943377368 | 0.00494322 | 1.239203824 | 0.000446016 |
| D65X | 1 | 0.985917913 | 0.038356 | 0.737398914 | 1.149083343 | 1.65E-09 | 0.958478008 | 0.059060656 | 1.074020464 | 0.101836715 | 1.365825681 | 2.92E-07 |
| T145S | 1 | 0.98526435 | 0.063685 | 0.414225813 | 1.035241646 | 0.003402527 | 0.710368732 | 4.10E-07 | 0.817757071 | 0.24644699 | 1.562908985 | 3.02E-12 |
| Q141X | 1 | 0.984942264 | 0.059171 | 0.834111907 | 0.957270638 | 0.144439647 | 0.414250463 | 3.36E-25 | 0.69446819 | 4.87E-07 | 0.863647858 | 0.003663843 |
| T67X | 1 | 0.983082781 | 0.048633 | 0.02185848 | 1.104206788 | 0.817732072 | 1.093922679 | 0.20830658 | 1.000876697 | 0.049567018 | 1.267381001 | 1.60E-13 |
| G70X | 1 | 0.982859507 | 0.070807 | 0.803832167 | 0.942164792 | 5.20E-05 | 1.327225689 | 4.33E-06 | 1.094224093 | 0.010177557 | 0.95951193 | 0.132304618 |
| F226G | 5 | 0.981832183 | 0.069306 | 1.19E-05 | 0.867707531 | 1.27E-10 | 0.97564544 | 0.008165731 | 0.994739929 | 0.283194448 | 0.854423041 | 7.48E-18 |
| L144I | 2 | 0.981318043 | 0.057449 | 0.000765561 | 1.138368007 | 3.44E-55 | 0.626543596 | 8.16E-149 | 0.760393326 | 9.70E-82 | 1.364287177 | 8.48E-49 |
| K227X | 1 | 0.980234609 | 0.049259 | 0.185172452 | 0.90174468 | 2.53E-28 | 1.010097191 | 0.233469754 | 1.132182615 | 9.83E-07 | 0.84126752 | 2.46E-13 |
| E71X | 1 | 0.979734945 | 0.070595 | 0.235817624 | 0.887174909 | 1.29E-32 | 1.291879815 | 0.003733244 | 1.196388387 | 0.000943694 | 0.860998672 | 1.23E-07 |
| L66X | 1 | 0.979137289 | 0.060222 | 0.232145385 | 1.225778499 | 5.13E-10 | 1.072226451 | 0.032108789 | 1.058480646 | 0.001511752 | 1.500572842 | 1.62E-20 |
| D65A | 3 | 0.978631384 | 0.041037 | 0.069065852 | 1.012144663 | 0.66474803 | 0.943247076 | 0.003640345 | 1.00800958 | 0.131233087 | 1.122554679 | 0.013309772 |
| G70M | 1 | 0.977644969 | 0.070432 | 0.809132904 | 0.858784224 | 9.97E-14 | 1.306501337 | 0.009717151 | 1.134543978 | 0.007941146 | 0.808535397 | 7.01E-09 |
| Y181X | 1 | 0.977587672 | 0.049617 | 0.458832967 | 0.961296712 | 0.007199818 | 0.937585258 | 0.955013736 | 1.266595666 | 1.10E-09 | 0.824616112 | 1.75E-13 |
| Y181X | 1 | 0.976497825 | 0.049961 | 0.12420143 | 0.809652803 | 1.34E-59 | 1.077010192 | 0.052818836 | 1.201296037 | 4.67E-14 | 0.655233217 | 2.75E-33 |
| R72X | 1 | 0.976194767 | 0.049554 | 0.261685561 | 1.132831403 | 3.25E-36 | 0.930308373 | 3.72E-06 | 0.687421282 | 6.72E-53 | 1.324983658 | 1.45E-21 |
| D92X | 1 | 0.975776986 | 0.073325 | 0.448876421 | 1.061013217 | 1.82E-07 | 1.369760567 | 9.91E-44 | 1.315283313 | 1.42E-43 | 1.253487981 | 2.07E-24 |
| T145X | 1 | 0.974908563 | 0.074647 | 0.031546116 | 1.011289927 | 0.032281098 | 1.032373339 | 0.000903029 | 0.958688456 | 0.474801885 | 1.031126252 | 0.016877299 |
| F226X | 1 | 0.972460944 | 0.047809 | 0.003177388 | 0.988670941 | 0.052883509 | 1.080225906 | 0.002497838 | 0.990386938 | 0.753369978 | 1.060185343 | 0.023315276 |
| W178X | 1 | 0.97020525 | 0.081857 | 0.001077534 | 1.092187041 | 0.645748654 | 0.461840171 | 0.000114942 | 1.273930188 | 0.118712765 | 1.033553459 | 0.107827563 |
| Y181X | 1 | 0.969422635 | 0.049599 | 6.17E-05 | 1.016343618 | 0.043570518 | 0.798038884 | 3.08E-31 | 1.076684424 | 2.76E-06 | 1.133010202 | 1.11E-07 |
| G70R | 2 | 0.968393801 | 0.066911 | 0.021380246 | 0.987430763 | 0.164559001 | 1.082502388 | 0.042820272 | 1.046816124 | 0.493208109 | 0.865934962 | 0.165495714 |
| G70I | 2 | 0.968008841 | 0.069737 | 0.163630832 | 0.801369256 | 6.86E-25 | 1.892971697 | 5.71E-08 | 1.469368805 | 0.02201465 | 0.648879861 | 6.17E-14 |
| L66X | 1 | 0.967750637 | 0.05843 | 0.032397554 | 1.134607059 | 0.000712473 | 0.989361224 | 0.11185726 | 0.936270778 | 1.59E-05 | 1.3771054 | 1.48E-19 |
| K227X | 1 | 0.967001968 | 0.077832 | 0.132294198 | 0.922298286 | 1.58E-06 | 1.285990954 | 0.907602909 | 1.065248679 | 0.002582223 | 1.029494675 | 0.877219338 |
| D65X | 1 | 0.966408279 | 0.042153 | 0.247974296 | 1.068771951 | 2.09E-07 | 0.963113038 | 0.18287807 | 0.995541294 | 0.48046194 | 1.164076539 | 7.48E-05 |
| D65S | 4 | 0.962519766 | 0.037446 | 5.61E-10 | 1.149891194 | 0.000155135 | 1.003624808 | 0.117197972 | 0.976251045 | 0.100464392 | 1.415885983 | 7.80E-13 |
| I228Y | 2 | 0.962275643 | 0.0849 | 1.18E-09 | 1.038648739 | 0.672457045 | 1.134381458 | 0.00348065 | 0.98084773 | 0.008091462 | 1.153629865 | 0.000110337 |
| I48X | 1 | 0.961995281 | 0.057984 | 0.001591233 | 1.193577069 | 4.84E-11 | 1.126003362 | 0.295457391 | 1.037431846 | 1.59E-06 | 1.320412891 | 0.147827049 |
| L174C | 2 | 0.961791675 | 0.049741 | 7.26E-14 | 1.020834827 | 0.288879807 | 1.32508184 | 3.16E-05 | 0.857129545 | 2.39E-05 | 1.21830496 | 2.66E-05 |
| L66Q | 1 | 0.961786924 | 0.048555 | 0.037303739 | 1.077415674 | 4.37E-13 | 1.252702819 | 4.12E-22 | 1.095817618 | 0.000121987 | 1.284902041 | 1.21E-20 |
| C138N | 3 | 0.961587076 | 0.081152 | 0.000297516 | 0.866399438 | 1.81E-10 | 0.929492914 | 0.008063073 | 1.1441005 | 0.140354652 | 0.703497048 | 7.94E-09 |
| Y146X | 1 | 0.960946503 | 0.043756 | 0.025487036 | 1.121318668 | 0.137663268 | 1.058061097 | 0.368813834 | 0.989001369 | 0.638768707 | 1.421327253 | 0.116023407 |
| L144X | 1 | 0.960851671 | 0.049559 | 0.005985536 | 0.990781687 | 0.225576735 | 1.161101098 | 0.062888045 | 1.062551361 | 3.51E-05 | 1.021621831 | 0.016402718 |
| R72X | 1 | 0.960647898 | 0.053454 | 0.867624313 | 0.999024222 | 0.640213746 | 0.984374369 | 0.976714404 | 0.845388384 | 3.54E-07 | 0.989444786 | 0.012271148 |
| E199X | 1 | 0.960135562 | 0.067706 | 0.165188909 | 0.908964288 | 4.41E-07 | 1.08637305 | 0.356490251 | 1.00187623 | 0.454772636 | 0.936877181 | 3.13E-05 |
| G229N | 5 | 0.958893802 | 0.078633 | 2.24E-14 | 1.044784523 | 0.255491456 | 1.301771728 | 1.05E-07 | 1.185013399 | 0.000253757 | 1.475619959 | 8.54E-21 |
| D65X | 1 | 0.958650198 | 0.037296 | 0.209419943 | 0.888730179 | 2.84E-06 | 1.250489204 | 0.008977161 | 1.302497397 | 0.000450102 | 0.978122517 | 0.524907033 |
| T145X | 1 | 0.958536541 | 0.061958 | 0.002413464 | 1.067016764 | 5.32E-12 | 0.910972259 | 1.86E-05 | 0.939977079 | 0.00354935 | 1.314146955 | 1.11E-28 |
| F134F | 1 | 0.957476892 | 0.056284 | 0.002750082 | 0.9465924 | 4.89E-18 | 1.208731594 | 9.81E-16 | 1.249085611 | 8.10E-37 | 0.935818524 | 2.96E-06 |
| G229P | 9 | 0.956800593 | 0.063029 | 1.25E-07 | 0.940416527 | 0.432596499 | 1.030023599 | 0.004724844 | 0.945893951 | 0.024970341 | 1.202481264 | 0.017607771 |
| M114S | 3 | 0.954423663 | 0.058002 | 9.63E-09 | 1.041149267 | 0.095449543 | 0.622906877 | 8.77E-18 | 0.859758587 | 0.011555084 | 1.181198152 | 0.000368257 |
| N69X | 1 | 0.953119113 | 0.054838 | 0.154255457 | 1.061331614 | 0.00057224 | 0.970275889 | 0.914378305 | 0.754279701 | 8.60E-07 | 1.327395608 | 8.22E-08 |
| Y146X | 1 | 0.953063952 | 0.071818 | 0.008871407 | 1.029286839 | 0.055258455 | 1.211987955 | 0.238137201 | 1.051508058 | 0.248416397 | 1.034310019 | 0.414602123 |
| L174A | 1 | 0.949568067 | 0.051344 | 0.031926686 | 1.099235312 | 0.458668421 | 0.727190812 | 0.000165088 | 0.881953988 | 0.00783906 | 1.5772397 | 0.125425779 |
| Q141S | 1 | 0.948244767 | 0.054336 | 0.055284584 | 1.01312675 | 0.986950418 | 0.598185996 | 5.23E-06 | 0.833462586 | 0.023926105 | 1.119718227 | 0.087414225 |
| Q64X | 1 | 0.947895126 | 0.044972 | 0.031736704 | 0.921488942 | 6.54E-08 | 1.108264759 | 0.003170885 | 0.886968118 | 0.000154243 | 0.908065599 | 0.003218965 |
| I228X | 1 | 0.947753016 | 0.060461 | 0.144104431 | 0.87195407 | 9.92E-20 | 1.157415864 | 0.057890551 | 0.997546495 | 0.116427929 | 0.904697525 | 0.005049913 |
| Q64X | 1 | 0.946721917 | 0.041446 | 0.006961914 | 1.030024838 | 0.00199406 | 1.244855497 | 1.40E-06 | 0.871486426 | 0.000567239 | 1.247829976 | 2.65E-11 |
| Y181X | 1 | 0.945997816 | 0.048014 | 0.00468979 | 1.083608336 | 1.21E-20 | 1.132369201 | 3.94E-11 | 1.095183422 | 9.63E-06 | 1.274450344 | 2.36E-19 |
| G229X | 1 | 0.944805205 | 0.082489 | 0.155497 | 1.04710657 | 0.000380252 | 1.182837425 | 0.394032466 | 0.866102041 | 0.00031867 | 1.270010573 | 1.37E-11 |
| S189Y | 1 | 0.94317555 | 0.047582 | 0.15220492 | 0.723264795 | 1.17E-17 | 1.166213193 | 0.031259612 | 1.345045378 | 0.000608774 | 0.573491167 | 6.26E-10 |
| V61X | 1 | 0.942636783 | 0.042454 | 0.000190579 | 1.121776248 | 1.01E-21 | 0.932277415 | 0.003600438 | 0.916100222 | 9.59E-05 | 1.3148022 | 1.39E-49 |
| F226H | 1 | 0.941629935 | 0.051723 | 3.09E-10 | 0.970462192 | 0.02787094 | 0.91956189 | 0.00308544 | 0.918220528 | 5.83E-07 | 1.023266589 | 0.001024229 |
| G70X | 1 | 0.941204915 | 0.07137 | 0.001439635 | 0.881018184 | 2.96E-28 | 1.099043285 | 7.12E-06 | 1.115013796 | 1.54E-06 | 0.835920228 | 7.22E-14 |
| G229S | 8 | 0.941142533 | 0.066367 | 9.47E-20 | 1.222439601 | 0.83093574 | 0.955644403 | 0.003231235 | 0.894224332 | 6.77E-08 | 1.383099742 | 2.37E-32 |
| C138D | 1 | 0.941106265 | 0.077581 | 0.249788839 | 0.640470157 | 9.28E-43 | 1.196046468 | 0.017172713 | 1.095605788 | 0.137521079 | 0.490656858 | 1.16E-32 |
| C138X | 1 | 0.938468819 | 0.077363 | 1.35E-10 | 1.078623337 | 0.0497472 | 0.979375289 | 0.000245517 | 0.896029903 | 0.002894439 | 1.225220052 | 2.65E-06 |
| I228M | 3 | 0.938080657 | 0.077725 | 2.41E-09 | 1.179951508 | 0.018477125 | 1.101541879 | 6.28E-09 | 0.997694429 | 0.000107077 | 1.403427489 | 3.60E-12 |
| F226X | 1 | 0.936913612 | 0.041892 | 2.03E-06 | 0.958286193 | 0.000104325 | 0.973416373 | 0.371014062 | 0.967303856 | 0.079245397 | 1.040225204 | 0.012576729 |
| D65X | 1 | 0.936782043 | 0.036445 | 0.178197496 | 1.072867445 | 0.000119451 | 1.01668903 | 0.243238141 | 1.371074282 | 6.80E-06 | 1.497621586 | 3.39E-12 |
| L174X | 1 | 0.936652661 | 0.056832 | 0.006730278 | 0.982420999 | 0.294761866 | 0.988413236 | 0.01791041 | 1.073859357 | 0.502644839 | 0.879694328 | 0.485211996 |
| D207X | 1 | 0.936289409 | 0.070357 | 3.36E-05 | 0.913880232 | 1.89E-09 | 1.234637016 | 1.53E-13 | 0.926093834 | 0.000439938 | 0.959778006 | 0.032496885 |
| I228D | 3 | 0.93597825 | 0.077551 | 2.78E-07 | 1.054704723 | 3.82E-07 | 1.321587273 | 0.066673936 | 1.170819475 | 1.20E-06 | 1.102917668 | 0.040583483 |

| | | | | | | | | | | | | |
|---|---|---|---|---|---|---|---|---|---|---|---|---|
| I115V | 1 | 0.935237895 | 0.070143 | 1.51E-06 | 1.113876124 | 4.70E-21 | 1.065012648 | 0.314856846 | 0.950522111 | 0.023716426 | 1.188994441 | 0.002539005 |
| D65X | 1 | 0.935205723 | 0.040792 | 0.173473361 | 1.075281867 | 1.12E-07 | 0.690618803 | 1.99E-21 | 0.858705018 | 4.72E-06 | 1.17686802 | 2.24E-06 |
| C138X | 1 | 0.935063423 | 0.068196 | 0.047325066 | 0.701721196 | 2.73E-25 | 1.659781996 | 1.45E-07 | 1.81485842 | 4.41E-06 | 0.566604618 | 1.34E-15 |
| S189X | 1 | 0.934624972 | 0.044988 | 0.011033835 | 0.987863156 | 0.877631915 | 1.106968911 | 0.005350249 | 0.922473934 | 0.01429801 | 1.214474985 | 1.82E-05 |
| Y181X | 1 | 0.934268365 | 0.050519 | 4.89E-16 | 0.978916313 | 0.002151866 | 1.008350173 | 0.06346582 | 0.937997276 | 0.005230905 | 1.39606637 | 7.92E-19 |
| L66X | 1 | 0.933822306 | 0.058402 | 1.38E-10 | 1.154389929 | 0.42772488 | 1.014582325 | 0.008594478 | 1.049797323 | 0.008660122 | 1.32517686 | 4.44E-27 |
| C218X | 1 | 0.933580216 | 0.049876 | 0.07610485 | 0.877260187 | 4.70E-13 | 1.028839587 | 0.126911181 | 1.755686472 | 5.81E-21 | 0.730618424 | 5.95E-12 |
| T145X | 1 | 0.931582494 | 0.070165 | 0.025381194 | 0.948124462 | 0.000711767 | 1.075889148 | 0.009404346 | 1.157498565 | 2.31E-13 | 0.882694048 | 0.001281122 |
| L93X | 1 | 0.931167284 | 0.056233 | 6.88E-12 | 1.020872672 | 0.924811123 | 0.94684145 | 0.000522021 | 0.965542493 | 0.045136349 | 1.217009842 | 7.66E-10 |
| F226X | 1 | 0.930965814 | 0.041626 | 3.01E-07 | 0.924869005 | 5.98E-06 | 1.040537578 | 0.026957484 | 0.980158407 | 0.419866581 | 1.214901344 | 0.232447039 |
| N69X | 1 | 0.930962643 | 0.071654 | 7.29E-11 | 1.101764246 | 0.251042906 | 1.084807953 | 9.38E-05 | 0.937137209 | 0.009494965 | 1.379896329 | 3.29E-11 |
| L144X | 1 | 0.929777547 | 0.047956 | 4.11E-07 | 0.983659855 | 0.000875346 | 0.590373463 | 5.36E-64 | 0.886439912 | 1.62E-06 | 1.098969701 | 0.02205958 |
| T67X | 1 | 0.929047387 | 0.046489 | 5.14E-05 | 1.112573086 | 7.83E-06 | 1.009560238 | 0.006396681 | 0.958824724 | 0.027755649 | 1.40312193 | 1.09E-20 |
| D207L | 1 | 0.928864859 | 0.069799 | 4.66E-06 | 0.995922038 | 0.834353648 | 1.268708036 | 3.37E-23 | 1.049537514 | 0.000750016 | 1.11319703 | 7.76E-07 |
| L144X | 1 | 0.928461836 | 0.047888 | 1.11E-07 | 1.079383233 | 2.87E-09 | 0.665602015 | 5.74E-75 | 0.799278394 | 1.70E-23 | 1.325370405 | 5.95E-22 |
| W178X | 1 | 0.928143682 | 0.055034 | 0.004718701 | 1.046639004 | 0.371537737 | 0.696654458 | 0.309712213 | 0.8493854 | 0.002321492 | 1.157920223 | 0.009700937 |
| P182X | 1 | 0.927692678 | 0.069711 | 0.000342738 | 0.876990992 | 1.25E-15 | 1.035173399 | 0.724959874 | 0.905670546 | 1.50E-05 | 0.82440531 | 8.30E-09 |
| K227X | 1 | 0.927316826 | 0.037259 | 6.17E-05 | 0.856931252 | 1.85E-24 | 1.218083619 | 2.64E-07 | 0.963229284 | 0.221510012 | 0.839880708 | 8.50E-07 |
| A210X | 1 | 0.925860584 | 0.069574 | 2.19E-05 | 0.943903746 | 4.26E-08 | 1.279024191 | 5.94E-28 | 1.037998461 | 0.134117231 | 1.002120637 | 0.098648596 |
| L89X | 1 | 0.925059848 | 0.060227 | 4.60E-05 | 1.015740609 | 0.458122922 | 1.09747698 | 0.043278717 | 0.892782359 | 0.14188324 | 1.219247779 | 0.308411127 |
| L89X | 1 | 0.925000508 | 0.058171 | 2.89E-06 | 1.086785632 | 3.24E-10 | 1.058846865 | 2.59E-12 | 0.89341515 | 0.017954371 | 1.276669337 | 0.187634274 |
| S189X | 1 | 0.924629124 | 0.054646 | 5.88E-15 | 0.93174021 | 8.15E-31 | 1.200604243 | 1.98E-33 | 1.05104215 | 0.004837445 | 0.950805607 | 0.151551468 |
| T145X | 1 | 0.92442133 | 0.070781 | 0.000817887 | 1.174581751 | 0.523010779 | 0.68485477 | 0.034836295 | 0.781504255 | 0.080353165 | 1.453452195 | 7.45E-32 |
| R72X | 1 | 0.922898337 | 0.051354 | 0.060732115 | 0.991382364 | 0.944232503 | 1.012520221 | 0.296420577 | 0.773550595 | 2.32E-16 | 1.064083576 | 0.674065794 |
| L93X | 1 | 0.922586375 | 0.055715 | 0.000114736 | 0.935059815 | 0.000158248 | 0.9052057 | 0.064969255 | 0.958808048 | 0.119240982 | 1.061358959 | 0.222667276 |
| Y146X | 1 | 0.921762989 | 0.069426 | 6.19E-05 | 1.007948742 | 0.113269355 | 1.080372371 | 0.001584447 | 1.002652077 | 0.616752588 | 1.082887415 | 0.000317656 |
| F226Q | 2 | 0.920882266 | 0.049609 | 2.98E-08 | 0.871638853 | 2.85E-50 | 0.936764204 | 0.060302013 | 1.00773042 | 0.032013851 | 0.84868452 | 0.013610715 |
| G70X | 1 | 0.920642769 | 0.063612 | 0.014800499 | 0.949538755 | 0.000358612 | 1.307981077 | 0.000334059 | 1.192760706 | 0.004276648 | 0.936865227 | 0.132057071 |
| F226K | 2 | 0.919444952 | 0.050504 | 2.09E-11 | 0.955836818 | 0.057894337 | 1.001272123 | 0.072245015 | 1.07570741 | 0.37434086 | 1.08506537 | 0.078056191 |
| G118X | 1 | 0.918482778 | 0.041234 | 8.43E-06 | 1.057702207 | 0.024827798 | 1.085018615 | 0.227689669 | 0.957976634 | 0.185344582 | 1.189502224 | 4.24E-10 |
| L144F | 4 | 0.917703951 | 0.047333 | 1.31E-133 | 0.855214085 | 4.38E-53 | 1.808070892 | 1.21E-235 | 2.187763842 | 5.29E-249 | 1.088869429 | 2.16E-14 |
| L174X | 1 | 0.917641824 | 0.050396 | 0.000516891 | 0.974650293 | 0.121779173 | 0.960545434 | 4.99E-06 | 1.029858123 | 0.014647138 | 1.185004843 | 0.472638181 |
| L174X | 1 | 0.91728788 | 0.05072 | 5.77E-08 | 1.052805795 | 0.2569381 | 0.940475863 | 0.118949028 | 0.982236285 | 0.115648678 | 1.15967076 | 5.65E-06 |
| A210X | 1 | 0.913637063 | 0.068655 | 0.000314681 | 0.804245047 | 1.31E-25 | 1.07062949 | 0.967586732 | 1.067466054 | 0.052529688 | 0.853875886 | 2.07E-08 |
| D65X | 1 | 0.91227871 | 0.038254 | 1.14E-05 | 0.907843892 | 1.27E-10 | 1.117030104 | 0.01278789 | 1.058494886 | 0.001643686 | 1.004922085 | 0.536675101 |
| I228X | 1 | 0.911317919 | 0.075508 | 0.000891585 | 1.034354159 | 0.010111921 | 1.29825884 | 1.77E-08 | 1.310056 | 4.23E-14 | 1.165565763 | 1.34E-06 |
| K227X | 1 | 0.910673764 | 0.047312 | 1.90E-05 | 0.999330098 | 0.489561009 | 0.776450207 | 1.01E-31 | 0.872593662 | 1.43E-06 | 1.119548013 | 0.000102414 |
| F226X | 1 | 0.908459192 | 0.040619 | 2.15E-09 | 0.901597405 | 3.97E-23 | 1.084321132 | 0.006350674 | 1.059537235 | 0.002180777 | 1.009621274 | 0.06587706 |
| M114L | 4 | 0.908230074 | 0.051478 | 0.001788836 | 1.147857842 | 0.347118281 | 0.759736571 | 0.001376951 | 0.829633914 | 0.011715131 | 1.085221095 | 0.000301607 |
| W178X | 1 | 0.907660552 | 0.042788 | 0.000514426 | 0.851050751 | 7.69E-06 | 1.009711787 | 1.85E-06 | 1.49749554 | 0.44668768 | 1.814383295 | 0.302384289 |
| D65Q | 2 | 0.905226663 | 0.039484 | 0.035586983 | 0.944906222 | 0.014052469 | 0.959795142 | 0.000737294 | 1.025697256 | 0.411746222 | 1.272304762 | 0.112862816 |
| G229W | 2 | 0.904039279 | 0.045641 | 0.018889396 | 1.073129005 | 0.082522529 | 1.312421916 | 0.004742357 | 1.000677933 | 0.338725481 | 1.250353137 | 0.38969884 |
| W178X | 1 | 0.90132352 | 0.066952 | 2.13E-16 | 1.034608414 | 0.004731355 | 1.165164575 | 4.46E-23 | 1.113889196 | 5.41E-06 | 1.170165945 | 4.74E-17 |
| C138X | 1 | 0.900766962 | 0.073508 | 1.21E-08 | 0.861346439 | 1.18E-28 | 0.959609872 | 0.130709866 | 0.935919538 | 0.255498239 | 0.89909522 | 9.49E-06 |
| D65X | 1 | 0.900464483 | 0.035032 | 0.038055858 | 0.999917155 | 0.673593445 | 0.907849204 | 0.013267875 | 1.162488076 | 0.002878574 | 1.319189328 | 1.89E-05 |
| Y146X | 1 | 0.900304837 | 0.049165 | 1.64E-12 | 1.148340456 | 0.08314711 | 1.075487411 | 0.305358611 | 0.996726794 | 0.167280126 | 1.559514763 | 1.08E-21 |
| T67X | 1 | 0.899356682 | 0.044352 | 2.43E-06 | 1.101562642 | 0.005427258 | 0.989738143 | 0.005850378 | 1.018541822 | 0.357632853 | 1.485344199 | 3.17E-24 |
| G70C | 1 | 0.897281405 | 0.068039 | 0.000142602 | 0.766245556 | 9.21E-51 | 1.030866248 | 0.069093419 | 1.464093233 | 7.76E-25 | 0.679325943 | 1.31E-31 |
| S189X | 1 | 0.895553665 | 0.078189 | 0.023020529 | 1.050811896 | 0.004055589 | 1.206989244 | 0.124327817 | 0.870842406 | 0.002583077 | 1.281002853 | 5.50E-11 |
| S189X | 1 | 0.894145846 | 0.045109 | 2.06E-14 | 0.914819915 | 2.10E-15 | 1.292726974 | 9.29E-30 | 1.214810943 | 3.25E-21 | 1.043940922 | 0.093720331 |
| G70X | 1 | 0.893817871 | 0.067776 | 0.016989393 | 0.662482484 | 5.91E-34 | 1.069691316 | 0.031199236 | 2.062758628 | 1.75E-19 | 0.578942875 | 2.46E-19 |
| W178X | 1 | 0.893762264 | 0.037345 | 0.023979972 | 1.012052503 | 0.418222875 | 0.399942246 | 2.28E-28 | 0.842358786 | 0.00018085 | 1.27499724 | 0.186862728 |
| L144X | 1 | 0.89372225 | 0.046096 | 9.50E-19 | 1.018279472 | 0.000271961 | 1.029658001 | 0.113172406 | 0.981969527 | 0.100114707 | 1.340454462 | 9.81E-51 |
| C218A | 1 | 0.893331831 | 0.045522 | 7.27E-05 | 1.015423675 | 0.219259366 | 0.665147996 | 7.01E-09 | 0.864130548 | 0.002377869 | 1.034760797 | 0.32586506 |
| D65E | 1 | 0.892662838 | 0.038411 | 0.013832404 | 0.879431383 | 2.81E-10 | 1.067611793 | 0.064425296 | 0.974571268 | 0.60591338 | 0.999344649 | 0.192983856 |
| F226X | 1 | 0.890121014 | 0.048894 | 6.43E-31 | 0.941754562 | 9.08E-12 | 0.952692663 | 0.029923698 | 1.013205072 | 0.55029721 | 1.085476171 | 5.00E-06 |
| G229K | 2 | 0.889101188 | 0.077626 | 0.004388548 | 1.05904076 | 0.037358811 | 1.47515914 | 0.006821831 | 1.018874826 | 0.479179571 | 1.557095933 | 0.001096604 |
| S189X | 1 | 0.887890306 | 0.054841 | 2.53E-28 | 1.03522065 | 0.00905806 | 0.97791587 | 0.286801921 | 1.062134867 | 0.002161261 | 1.34679532 | 8.28E-64 |
| L174X | 1 | 0.887798057 | 0.061289 | 1.46E-09 | 1.103295945 | 0.511646724 | 0.849274222 | 0.000374692 | 0.905080422 | 0.285378776 | 1.284050172 | 0.050861003 |
| D207X | 1 | 0.887727755 | 0.066708 | 5.57E-13 | 1.065273323 | 1.55E-06 | 1.140200139 | 4.05E-07 | 0.921815667 | 5.21E-07 | 1.348650977 | 8.46E-42 |
| L144M | 2 | 0.887411246 | 0.050232 | 7.13E-35 | 1.054545873 | 0.160133302 | 1.056724099 | 0.011703778 | 0.911351286 | 3.34E-07 | 1.500065162 | 4.01E-31 |
| F226X | 1 | 0.884899806 | 0.043504 | 1.34E-31 | 0.957722665 | 3.70E-08 | 1.107830675 | 1.54E-05 | 1.082954468 | 3.75E-06 | 1.143521435 | 2.34E-08 |
| G229D | 3 | 0.884825779 | 0.072559 | 2.60E-20 | 1.017621367 | 0.429153141 | 1.170733976 | 0.000495292 | 1.233004297 | 3.36E-05 | 1.286384428 | 1.72E-05 |
| D207X | 1 | 0.88446018 | 0.066463 | 3.56E-07 | 0.905242461 | 2.22E-14 | 1.302825724 | 2.19E-28 | 1.128886985 | 1.92E-05 | 1.011435901 | 0.378622506 |
| F226X | 1 | 0.883862989 | 0.047615 | 2.30E-18 | 0.853410229 | 2.28E-69 | 1.086372145 | 0.000610163 | 0.969543181 | 0.635819172 | 0.90075671 | 2.97E-18 |
| K227X | 1 | 0.882964028 | 0.036424 | 1.56E-14 | 0.911366214 | 4.80E-10 | 1.007003831 | 0.576166737 | 0.974385428 | 0.171581864 | 1.051476684 | 0.468471488 |
| W178X | 1 | 0.880801585 | 0.074314 | 1.61E-05 | 1.093842036 | 0.000126954 | 0.422496656 | 8.57E-05 | 0.742597209 | 0.20136006 | 1.202099717 | 0.157237276 |
| F134L | 4 | 0.880093011 | 0.05521 | 0.004604771 | 0.727566042 | 1.78E-10 | 1.172026842 | 0.007792977 | 1.447910341 | 0.000167845 | 0.905917351 | 0.01384322 |
| D207P | 1 | 0.880004777 | 0.066128 | 3.56E-16 | 1.055079799 | 2.55E-05 | 1.139501943 | 0.002439796 | 1.017943304 | 0.967587896 | 1.274093902 | 2.98E-32 |
| Q64X | 1 | 0.879437432 | 0.044461 | 5.53E-14 | 0.943011931 | 3.28E-08 | 0.996088033 | 0.022852574 | 0.902161856 | 0.000540639 | 1.035684533 | 0.090680362 |
| L174N | 3 | 0.879212851 | 0.060696 | 1.19E-06 | 1.025239156 | 0.853608382 | 1.081498444 | 0.071443838 | 1.259862449 | 3.26E-09 | 1.082678544 | 0.000406927 |
| G229E | 2 | 0.878543931 | 0.056986 | 2.55E-06 | 1.284630463 | 8.33E-06 | 1.099203693 | 2.94E-08 | 0.934056498 | 0.158005547 | 2.114162513 | 1.03E-56 |
| G70X | 1 | 0.87772087 | 0.063233 | 0.145693211 | 0.820262445 | 4.80E-08 | 1.259835333 | 0.330080574 | 1.033930981 | 0.544744582 | 0.887938984 | 0.00733454 |
| L89N | 2 | 0.877674126 | 0.05359 | 1.99E-08 | 1.080176458 | 0.285543421 | 1.031437644 | 0.366186555 | 0.876131698 | 0.000400859 | 1.272563978 | 0.16385987 |
| A55X | 1 | 0.877462346 | 0.053509 | 1.22E-18 | 0.987079651 | 0.437368144 | 0.985053611 | 0.029900878 | 0.90261433 | 0.06092349 | 1.208660498 | 0.001199115 |
| F226X | 1 | 0.871713359 | 0.038976 | 2.26E-22 | 0.888503499 | 5.65E-24 | 1.11636423 | 5.62E-06 | 1.000880566 | 0.537975853 | 1.065551399 | 0.001258834 |
| I228X | 1 | 0.868840367 | 0.058275 | 1.04E-19 | 1.064780986 | 1.75E-06 | 1.347827483 | 0.004650092 | 1.063419479 | 0.024234164 | 1.379104083 | 9.70E-10 |
| Q64X | 1 | 0.868373007 | 0.043902 | 1.78E-12 | 0.996653537 | 0.83433399 | 0.729854805 | 1.03E-27 | 0.70534614 | 2.83E-24 | 1.152967285 | 3.16E-06 |
| C218X | 1 | 0.868318718 | 0.044247 | 1.82E-12 | 0.935032941 | 9.15E-07 | 0.844231414 | 2.82E-06 | 1.123778154 | 0.001950934 | 0.982687768 | 0.522237157 |
| Y181X | 1 | 0.868226523 | 0.048497 | 2.00E-09 | 1.04486964 | 3.41E-06 | 1.022791344 | 0.08150264 | 1.045534644 | 0.068507203 | 1.341190402 | 8.19E-20 |
| D65H | 1 | 0.868216268 | 0.036407 | 9.39E-11 | 0.932756471 | 3.30E-05 | 0.930566963 | 0.001615789 | 1.031024761 | 0.236162464 | 1.198459064 | 1.85E-05 |
| Q64X | 1 | 0.867496014 | 0.043858 | 7.43E-16 | 1.024170761 | 0.249047111 | 0.784163667 | 1.02E-28 | 0.698164405 | 2.94E-37 | 1.152536888 | 1.59E-09 |
| Y181X | 1 | 0.86684167 | 0.043996 | 3.71E-20 | 0.977135128 | 0.890058147 | 1.209163568 | 3.65E-21 | 1.115309513 | 3.84E-06 | 1.215161877 | 7.33E-14 |

| | | | | | | | | | | | | |
|---|---|---|---|---|---|---|---|---|---|---|---|---|
| L93X | 1 | 0.866132161 | 0.049568 | 1.87E-12 | 0.937867597 | 4.10E-06 | 1.044427437 | 0.978843573 | 0.91532861 | 0.349324547 | 0.988655744 | 0.537834466 |
| W178X | 1 | 0.865374551 | 0.055095 | 2.11E-19 | 1.081204804 | 0.016331096 | 0.525151143 | 1.00E-22 | 0.6794778 | 1.87E-09 | 1.830449991 | 1.21E-11 |
| D65X | 1 | 0.86418232 | 0.037186 | 0.000240194 | 0.860592875 | 3.31E-12 | 1.061784011 | 0.102244032 | 1.065360525 | 0.002402554 | 1.018056528 | 0.089271726 |
| L66X | 1 | 0.864111117 | 0.042607 | 6.32E-24 | 1.019591373 | 0.114398001 | 0.907449866 | 1.25E-06 | 0.914789436 | 1.70E-08 | 1.306798801 | 4.21E-37 |
| D65V | 3 | 0.863954231 | 0.037477 | 1.44E-09 | 0.972026186 | 0.014447652 | 0.985572276 | 0.365492136 | 1.030372402 | 0.710137494 | 1.232488295 | 0.000121002 |
| R72X | 1 | 0.863876006 | 0.050534 | 2.84E-06 | 0.960414424 | 0.00240722 | 0.982943705 | 0.302805124 | 0.880338917 | 0.023393184 | 1.051113045 | 0.017434553 |
| W178X | 1 | 0.862514507 | 0.076092 | 2.05E-06 | 1.257839947 | 0.25154219 | 0.277155468 | 7.70E-40 | 0.692805324 | 9.31E-09 | 1.487361147 | 7.22E-12 |
| V61X | 1 | 0.85961299 | 0.051252 | 2.66E-11 | 1.08017583 | 5.31E-10 | 0.971109315 | 0.445536644 | 0.988186556 | 0.097866186 | 1.191998577 | 0.023872251 |
| D65X | 1 | 0.858723688 | 0.037456 | 1.55E-06 | 0.941633305 | 0.000116977 | 0.980963096 | 0.440916749 | 0.973583273 | 0.558969895 | 1.051987316 | 0.253413656 |
| L93X | 1 | 0.858630891 | 0.049138 | 2.62E-21 | 0.993405637 | 0.626158101 | 1.075106728 | 0.006782965 | 0.974992213 | 0.17652214 | 1.309940939 | 4.69E-27 |
| L174X | 1 | 0.857611606 | 0.052036 | 7.38E-14 | 0.928464032 | 0.000853749 | 0.987144389 | 0.313490043 | 1.180391493 | 0.009938573 | 1.083916664 | 0.020335335 |
| R72X | 1 | 0.85657253 | 0.050107 | 0.00250126 | 1.022260406 | 0.958496845 | 0.761192838 | 7.86E-07 | 0.72536381 | 0.000219224 | 0.974624457 | 0.79143988 |
| W178X | 1 | 0.855162767 | 0.05625 | 8.82E-07 | 1.063899444 | 0.92353401 | 0.606389177 | 1.23E-09 | 0.852419655 | 0.00077338 | 1.30488085 | 0.000895394 |
| L144A | 4 | 0.853229016 | 0.056215 | 1.06E-43 | 1.078637834 | 0.62094849 | 0.577882456 | 4.47E-99 | 0.750731001 | 5.45E-43 | 1.407226081 | 1.63E-48 |
| L93X | 1 | 0.853100336 | 0.051518 | 3.00E-06 | 0.829929313 | 2.92E-08 | 1.309677312 | 0.002705098 | 1.051393073 | 0.150634265 | 0.834020573 | 0.008429549 |
| T145X | 1 | 0.851915536 | 0.055066 | 3.55E-17 | 0.921568151 | 2.06E-06 | 0.867974505 | 0.294728299 | 0.838647634 | 0.005463077 | 1.636231763 | 5.96E-13 |
| S189K | 1 | 0.847131896 | 0.052324 | 0.000192321 | 0.736475311 | 2.14E-16 | 1.563847721 | 1.48E-09 | 1.602014683 | 8.86E-08 | 0.626766869 | 8.54E-12 |
| L144E | 1 | 0.843682928 | 0.04712 | 1.57E-09 | 0.710367495 | 1.37E-63 | 1.03703852 | 0.461370267 | 1.118242923 | 0.005524716 | 0.703192398 | 1.15E-19 |
| C138X | 1 | 0.843612288 | 0.069544 | 7.35E-23 | 1.194774848 | 0.006063333 | 0.45565109 | 1.29E-39 | 0.65526589 | 6.02E-30 | 2.010211975 | 1.44E-72 |
| F134X | 1 | 0.842765636 | 0.052868 | 1.21E-25 | 0.94952146 | 5.37E-07 | 1.213149228 | 4.61E-20 | 1.101035253 | 1.46E-12 | 1.080272755 | 0.000364606 |
| W178X | 1 | 0.841507307 | 0.055213 | 0.001100743 | 0.956068162 | 0.067833708 | 0.381647382 | 6.40E-14 | 0.787107283 | 0.033965685 | 1.197418139 | 0.007629452 |
| C138I | 2 | 0.841428655 | 0.07 | 2.86E-13 | 1.125493584 | 0.213610142 | 0.429356748 | 2.04E-62 | 0.659029572 | 2.05E-14 | 1.516881702 | 2.76E-22 |
| Q64X | 1 | 0.839449174 | 0.04244 | 1.17E-20 | 1.008694281 | 0.586453363 | 0.889843229 | 2.67E-09 | 0.749898788 | 5.34E-22 | 1.299127475 | 2.33E-17 |
| T145X | 1 | 0.83942043 | 0.063224 | 2.74E-18 | 0.910645106 | 3.68E-07 | 1.501794031 | 4.41E-62 | 1.041971926 | 0.005831521 | 1.150636628 | 8.63E-08 |
| Y146X | 1 | 0.836348942 | 0.063023 | 6.07E-23 | 0.924039077 | 1.80E-11 | 1.193747802 | 0.393888377 | 1.005806828 | 0.062424306 | 1.151590198 | 0.052447869 |
| F134X | 1 | 0.834298584 | 0.048994 | 3.23E-11 | 0.778995208 | 5.78E-40 | 0.854926554 | 0.178320879 | 1.305669316 | 4.52E-05 | 0.785478973 | 6.39E-10 |
| L144S | 6 | 0.829810113 | 0.046791 | 2.07E-57 | 1.043259743 | 0.056894094 | 0.721627099 | 3.90E-51 | 0.828363306 | 1.03E-31 | 1.50788552 | 2.61E-77 |
| C138M | 1 | 0.827925101 | 0.064271 | 1.45E-12 | 1.039797125 | 0.006011984 | 0.498585382 | 9.86E-68 | 0.718123448 | 2.08E-17 | 1.382044135 | 6.19E-27 |
| W178X | 1 | 0.823193905 | 0.047033 | 0.000171405 | 1.250804179 | 6.71E-06 | 0.63192505 | 0.000850042 | 0.869628849 | 0.005205726 | 1.332510594 | 0.00362493 |
| L66X | 1 | 0.821364013 | 0.045087 | 5.33E-46 | 1.06551171 | 1.95E-12 | 0.960474511 | 0.194428429 | 0.916604744 | 2.08E-05 | 1.681106816 | 1.47E-119 |
| E71D | 1 | 0.821348694 | 0.062937 | 5.31E-25 | 0.963728259 | 0.001019882 | 1.047872883 | 0.058958907 | 0.937037047 | 0.237645628 | 1.2751613 | 7.74E-12 |
| T145X | 1 | 0.821054015 | 0.06184 | 1.69E-16 | 0.838954208 | 5.04E-17 | 1.196300666 | 0.403552899 | 1.13623118 | 0.023039389 | 1.077821581 | 0.481323292 |
| L144X | 1 | 0.821026882 | 0.056755 | 7.58E-72 | 1.454723796 | 0.001917032 | 1.054902815 | 1.80E-24 | 0.919418585 | 1.88E-07 | 1.95595953 | 3.16E-114 |
| L144H | 1 | 0.820450525 | 0.042317 | 8.61E-08 | 0.805611998 | 4.73E-20 | 0.671141645 | 8.57E-08 | 1.027118305 | 0.784992583 | 0.908401801 | 0.002868618 |
| Y146X | 1 | 0.819923267 | 0.061755 | 3.50E-23 | 0.869617442 | 4.36E-15 | 1.395807296 | 6.14E-22 | 1.238496036 | 4.22E-17 | 0.853231772 | 0.001831388 |
| L144X | 1 | 0.819901089 | 0.045791 | 4.79E-69 | 1.009685435 | 0.124077839 | 1.12004083 | 2.27E-23 | 0.999152975 | 0.477755305 | 1.431973633 | 2.02E-75 |
| T67X | 1 | 0.816610326 | 0.037956 | 2.20E-17 | 1.122780007 | 1.14E-07 | 0.878908698 | 0.007771298 | 0.955096128 | 0.196857701 | 1.683493443 | 6.19E-39 |
| W178X | 1 | 0.814848202 | 0.038413 | 3.85E-10 | 1.017030076 | 0.765167404 | 1.011536482 | 0.189601989 | 1.530430572 | 8.22E-14 | 1.444703391 | 4.22E-05 |
| P182X | 1 | 0.812826196 | 0.061221 | 2.30E-36 | 0.949303422 | 2.60E-05 | 1.21874593 | 1.11E-22 | 1.111965185 | 3.17E-07 | 1.334322699 | 7.84E-32 |
| L144X | 1 | 0.812728119 | 0.054521 | 3.04E-76 | 1.035924858 | 0.000468307 | 0.649773303 | 4.36E-132 | 0.76456904 | 9.22E-78 | 1.413082448 | 1.18E-92 |
| T145X | 1 | 0.810516696 | 0.061047 | 1.43E-28 | 0.679076547 | 1.57E-100 | 2.70508456 | 2.44E-109 | 3.346895089 | 3.41E-143 | 0.752189135 | 2.92E-26 |
| S189X | 1 | 0.810299267 | 0.050049 | 3.21E-48 | 0.871684895 | 1.54E-36 | 1.180451532 | 3.95E-08 | 1.174241217 | 8.66E-11 | 1.109982375 | 4.10E-05 |
| L89E | 2 | 0.805742879 | 0.051998 | 4.83E-07 | 1.123786775 | 0.277678071 | 0.846433452 | 0.046272758 | 0.667347507 | 0.000208973 | 1.783677483 | 1.31E-11 |
| D65X | 1 | 0.799251099 | 0.034862 | 8.68E-12 | 0.959828077 | 0.015607154 | 1.012891783 | 0.237842698 | 0.888640401 | 0.008824139 | 1.230465629 | 1.32E-05 |
| P182X | 1 | 0.79352994 | 0.059767 | 1.84E-39 | 0.963981942 | 0.016908919 | 1.217922657 | 4.69E-16 | 1.119057785 | 8.81E-08 | 1.282715565 | 6.24E-27 |
| W178X | 1 | 0.792353294 | 0.062615 | 8.75E-19 | 1.003067991 | 0.933349847 | 0.607810862 | 0.043862571 | 0.896298188 | 1.99E-05 | 1.495602185 | 5.16E-19 |
| D65N | 2 | 0.787026027 | 0.033866 | 6.10E-12 | 0.898945351 | 3.29E-08 | 0.884129487 | 0.038419825 | 1.068869221 | 0.133289363 | 1.320140336 | 7.65E-06 |
| Q141X | 1 | 0.786534924 | 0.04507 | 1.12E-06 | 1.110496836 | 0.018175538 | 0.210932403 | 3.55E-09 | 1.009766745 | 0.657949752 | 2.269160052 | 3.48E-15 |
| F226E | 1 | 0.785988879 | 0.038642 | 4.91E-38 | 0.854158925 | 1.75E-30 | 1.01475309 | 0.976366251 | 1.071075727 | 0.037461254 | 1.145711044 | 0.06211374 |
| L144X | 1 | 0.785365002 | 0.043863 | 1.07E-24 | 0.817788566 | 1.63E-37 | 0.708494549 | 5.79E-12 | 1.08285158 | 0.341801224 | 0.997301726 | 0.160647428 |
| F134M | 1 | 0.784465664 | 0.052117 | 7.42E-13 | 0.841708914 | 1.85E-16 | 0.831803076 | 0.000305667 | 1.056258486 | 0.085376534 | 1.093038097 | 0.429448855 |
| A210X | 1 | 0.783694561 | 0.058891 | 1.99E-58 | 0.942435435 | 1.08E-09 | 1.486362792 | 1.01E-64 | 1.222220607 | 5.68E-23 | 1.355979541 | 1.34E-61 |
| D65D | 1 | 0.781060945 | 0.033609 | 3.24E-15 | 0.951684385 | 0.000421049 | 1.196248226 | 4.60E-06 | 1.003351576 | 0.123534304 | 1.454540781 | 2.01E-09 |
| L174X | 1 | 0.778178417 | 0.044741 | 1.53E-10 | 0.871735218 | 9.39E-06 | 1.180996015 | 0.409695388 | 1.149141159 | 0.345805326 | 1.060552412 | 0.495734666 |
| L174T | 2 | 0.776898988 | 0.053633 | 7.95E-31 | 0.877172785 | 8.98E-15 | 1.039345255 | 0.096828235 | 1.045802395 | 0.005034396 | 1.156749739 | 2.62E-05 |
| L144X | 1 | 0.776477964 | 0.052089 | 4.49E-45 | 0.762185893 | 2.99E-115 | 1.097046584 | 0.002248904 | 1.114904583 | 4.85E-06 | 0.819929373 | 4.41E-13 |
| W178X | 1 | 0.772617978 | 0.039309 | 4.54E-12 | 0.857808569 | 4.45E-12 | 0.563906045 | 2.01E-16 | 0.926517391 | 0.315916439 | 0.962598986 | 0.652196895 |
| L144Q | 1 | 0.766247047 | 0.051403 | 4.46E-58 | 0.94657447 | 2.72E-12 | 0.761546271 | 1.90E-32 | 0.793095934 | 1.10E-29 | 1.164841626 | 1.72E-15 |
| W178X | 1 | 0.763852361 | 0.062501 | 3.53E-14 | 1.009248282 | 0.173084394 | 0.461325894 | 1.08E-17 | 0.966945173 | 0.245831883 | 1.267145204 | 3.20E-05 |
| D65X | 1 | 0.74945945 | 0.031427 | 7.83E-21 | 0.947896814 | 0.037651379 | 0.912607017 | 0.014020327 | 1.087842905 | 0.009815292 | 1.43413513 | 6.79E-11 |
| A210X | 1 | 0.749205325 | 0.049508 | 2.67E-10 | 0.751563763 | 5.08E-18 | 0.80952496 | 0.000921776 | 1.055156297 | 0.12272952 | 1.030252837 | 0.449762 |
| D65X | 1 | 0.745578166 | 0.031264 | 1.28E-15 | 0.956294342 | 0.038847107 | 1.032641457 | 0.460577202 | 0.913686832 | 0.037414618 | 1.413550636 | 1.35E-08 |
| S189X | 1 | 0.739100736 | 0.064529 | 2.05E-10 | 0.841232518 | 1.25E-10 | 1.699300831 | 2.07E-07 | 1.040789864 | 0.886436182 | 1.039531659 | 0.007252489 |
| L144X | 1 | 0.737210755 | 0.038024 | 6.00E-29 | 0.89696329 | 4.40E-10 | 0.953455876 | 0.798553123 | 1.267538393 | 3.26E-07 | 1.352788171 | 1.53E-07 |
| A55X | 1 | 0.728955565 | 0.045057 | 1.90E-49 | 0.839699156 | 8.89E-16 | 1.169368578 | 1.87E-06 | 1.187544507 | 0.005897133 | 1.384192725 | 3.85E-10 |
| W178X | 1 | 0.718566992 | 0.036559 | 7.52E-16 | 1.082864278 | 0.080914657 | 0.489550833 | 0.006777161 | 1.665914008 | 0.000360822 | 1.715707211 | 1.37E-17 |
| M114M | 1 | 0.703668552 | 0.042763 | 1.42E-92 | 0.941982664 | 7.94E-12 | 0.76670235 | 1.58E-14 | 0.936820317 | 0.171546276 | 1.813400655 | 5.75E-85 |
| V61X | 1 | 0.701855177 | 0.043374 | 2.99E-65 | 0.92236177 | 2.14E-10 | 1.235153139 | 5.16E-17 | 1.058500448 | 0.20325199 | 1.67450513 | 3.86E-54 |
| W178X | 1 | 0.691024535 | 0.058302 | 2.73E-14 | 0.892727409 | 0.000490642 | 0.862359484 | 0.343540713 | 0.911693135 | 0.01530532 | 1.206910071 | 0.000463205 |
| M114X | 1 | 0.675170098 | 0.037741 | 7.26E-45 | 1.004900753 | 0.391591839 | 0.635998219 | 8.58E-17 | 0.873764779 | 0.000808912 | 1.539915469 | 3.56E-24 |
| W178X | 1 | 0.667460241 | 0.039577 | 2.92E-25 | 0.924784673 | 0.015311989 | 0.868136957 | 0.042466053 | 1.144461425 | 0.032562245 | 1.459191982 | 4.31E-17 |
| W178X | 1 | 0.634006661 | 0.029888 | 5.21E-28 | 0.844357854 | 6.01E-11 | 0.575006436 | 7.92E-12 | 0.933605889 | 0.3521162 | 1.558536792 | 3.14E-09 |
| F134W | 1 | 0.632209512 | 0.042002 | 2.49E-86 | 0.832323421 | 4.08E-39 | 1.274741529 | 4.59E-11 | 1.406117523 | 2.82E-15 | 1.435695124 | 3.33E-28 |
| Q141X | 1 | 0.624106996 | 0.037494 | 6.67E-58 | 0.771434437 | 5.77E-31 | 0.982154913 | 0.785473796 | 0.996334454 | 0.778629812 | 1.131290037 | 0.037728057 |
| N81X | 1 | 0.612351421 | 0.027898 | 1.27E-36 | 0.975794778 | 0.5872264 | 1.242483968 | 0.002795955 | 1.206886749 | 0.029074971 | 2.669311922 | 4.81E-41 |
| L144G | 3 | 0.611452076 | 0.031537 | 2.71E-85 | 0.782861694 | 6.05E-55 | 1.039830203 | 0.01976363 | 1.477372483 | 2.23E-23 | 1.277403499 | 1.95E-10 |
| G70X | 1 | 0.609416255 | 0.043904 | 2.50E-15 | 0.895970368 | 0.018469478 | 2.04310676 | 3.14E-13 | 1.38721334 | 0.000117847 | 0.738821042 | 2.38E-09 |
| G229X | 1 | 0.559134735 | 0.028228 | 6.96E-25 | 0.781527116 | 2.30E-13 | 1.565115318 | 5.95E-06 | 1.281368871 | 0.007297556 | 1.152887973 | 0.01057226 |
| L174X | 1 | 0.517599786 | 0.035732 | 3.10E-28 | 0.685129705 | 5.38E-28 | 1.42984257 | 4.10E-06 | 1.07111734 | 0.172308176 | 1.102195519 | 0.0581559 |
| L144W | 2 | 0.47597029 | 0.026839 | 1.58E-232 | 0.560615746 | 5.43E-304 | 3.150282155 | 2.44E-176 | 3.947149935 | 8.44E-180 | 1.369106227 | 1.78E-08 |
| G229X | 1 | 0.462091591 | 0.023329 | 1.09E-38 | 0.89117171 | 5.08E-06 | 1.973246195 | 3.65E-09 | 1.445716349 | 2.01E-11 | 1.705772138 | 3.29E-36 |
| Q141X | 1 | 0.44306635 | 0.025388 | 3.68E-50 | 0.803475705 | 1.66E-22 | 1.824922154 | 9.43E-13 | 2.024773919 | 1.92E-20 | 2.050597291 | 1.63E-27 |
| L174W | 2 | 0.442189597 | 0.025702 | 1.51E-63 | 0.63774938 | 4.70E-56 | 2.386454951 | 8.53E-28 | 2.345329198 | 6.99E-24 | 1.860087153 | 6.03E-21 |

| | | | | | | | | | | | | |
|---|---|---|---|---|---|---|---|---|---|---|---|---|
| L174F | 1 | 0.426213445 | 0.029424 | 5.40E-26 | 0.549037624 | 1.29E-40 | 2.969566769 | 6.21E-28 | 2.397824676 | 1.15E-17 | 1.129899282 | 0.016387202 |
