## Supplementary Table 2 for "Sensitivity optimization of a rhodopsin-based fluorescent voltage indicator"

| mutation | number of plates | ΔF/F0 (norm.) | ΔF/F0 | ΔF/F0 p value | SNR (norm.) | SNR p value) | τ_ON (norm.) | τ_ON p value | τ_OFF (norm.) | τ_OFF p value | F0 (norm.) | F0 p value |
|---|---|---|---|---|---|---|---|---|---|---|---|---|
| Y63L N69E L89C A122D | 1 | 1.400394 | 0.119656 | 4.75E-122 | 1.034546 | 0.030383 | 0.724406 | 3.05E-78 | 0.615754 | 1.11E-202 | 0.748288 | 2.23E-48 |
| N69E R78H A122D V196P | 1 | 1.363739 | 0.117594 | 0 | 1.158031 | 5.06E-103 | 0.949387 | 5.94E-32 | 0.849834 | 1.47E-126 | 0.743315 | 1.61E-176 |
| R78H L89C A122D | 1 | 1.340723 | 0.114557 | 3.57E-130 | 0.994041 | 0.007851 | 0.986795 | 0.006965 | 0.929625 | 1.90E-06 | 0.785408 | 2.93E-36 |
| V74E L89C | 1 | 1.317506 | 0.129118 | 3.57E-97 | 1.070194 | 0.000153 | 0.699228 | 251667299 | 0.927285 | 9.69E-30 | 0.957425 | 5.19E-07 |
| R78H A122D | 1 | 1.298843 | 0.088718 | 5.16E-105 | 0.77821 | 5.82E-55 | 1.151587 | 7.09E-13 | 1.056086 | 0.014619 | 0.391597 | 5.87E-199 |
| R78H L89C A122D V196P | 1 | 1.244361 | 0.130071 | 1.85E-119 | 1.114812 | 4.31E-40 | 0.706522 | 1.75E-193 | 0.612595 | 0 | 1.021059 | 0.569827903 |
| R78H N81S L89C A122D | 1 | 1.234064 | 0.140012 | 3.02E-142 | 0.980819 | 0.000452 | 0.900165 | 3.67E-14 | 0.817163 | 8.57E-61 | 0.996237 | 0.293434984 |
| R78H A122D V196P | 1 | 1.210048 | 0.100904 | 1.74E-110 | 1.018632 | 0.08261 | 0.934847 | 0.000233 | 0.961053 | 1.26E-05 | 0.684806 | 8.83E-83 |
| N69E V196P | 1 | 1.205604 | 0.103958 | 2.61E-212 | 1.154624 | 1.34E-88 | 0.898615 | 1.61E-92 | 0.722051 | 0 | 0.955946 | 3.83E-09 |
| Y63L L89C A122D | 1 | 1.183029 | 0.101083 | 6.62E-32 | 1.125133 | 3.08E-05 | 0.612449 | 4.41E-157 | 0.599039 | 3.53E-201 | 0.85768 | 1.14E-13 |
| Y63L L89C A122D V196P | 1 | 1.182272 | 0.117974 | 2.45E-58 | 1.195757 | 1.42E-72 | 0.747788 | 4.41E-86 | 0.766772 | 7.51E-29 | 1.059917 | 5.04E-07 |
| Y63L R78H L89C | 1 | 1.143691 | 0.097722 | 3.23E-05 | 0.842773 | 1.02E-28 | 0.723599 | 7.22E-25 | 0.716444 | 4.82E-36 | 0.705428 | 3.02E-26 |
| N69E R78H N81S A122D V196P | 1 | 1.130005 | 0.091943 | 6.82E-19 | 0.743939 | 1.46E-72 | 0.952367 | 0.05621 | 0.746956 | 2.40E-41 | 0.448253 | 3.76E-158 |
| N69E A122D | 1 | 1.106093 | 0.095377 | 6.20E-70 | 1.032264 | 2.46E-06 | 0.756723 | 007889932 | 0.846678 | 1.63E-104 | 0.793757 | 5.02E-101 |
| N69E L89C A122D | 1 | 1.105754 | 0.094481 | 1.17E-08 | 1.141084 | 6.13E-07 | 0.762349 | 4.28E-66 | 0.653628 | 2.26E-167 | 0.911098 | 8.95E-05 |
| Y63L N69E R78H A122D | 1 | 1.104923 | 0.099115 | 1.07E-05 | 0.815734 | 2.32E-11 | 0.772426 | 1.33E-06 | 0.996705 | 0.585153 | 0.485159 | 1.70E-34 |
| V74E R78H L89C V196P | 1 | 1.085856 | 0.082851 | 6.92E-09 | 0.898502 | 2.78E-06 | 0.821859 | 1.62E-08 | 1.286838 | 1.37E-10 | 0.679678 | 3.34E-22 |
| Y63L L89T A122D V196P | 1 | 1.073899 | 0.11473 | 3.88E-10 | 1.125742 | 1.24E-41 | 0.644917 | 9.37E-223 | 0.631749 | 685049248 | 1.27916 | 2.46E-57 |
| N69E A122D V196P | 1 | 1.06332 | 0.088668 | 2.80E-21 | 1.095655 | 2.51E-19 | 0.778408 | 1.29E-131 | 0.712489 | 1.98E-185 | 0.993914 | 0.907292122 |
| N69E R78H A122D | 1 | 1.058286 | 0.106503 | 8.30E-16 | 1.06627 | 8.09E-08 | 1.097203 | 3.08E-09 | 1.032515 | 0.01048 | 0.894019 | 1.91E-14 |
| R78H N81S A122D V196P | 1 | 1.045193 | 0.085042 | 4.04E-09 | 0.846839 | 9.08E-35 | 0.971346 | 0.287465 | 1.093144 | 5.46E-08 | 0.669482 | 1.63E-127 |
| N69E V74W R78H N81S L89C A122H V196P | 1 | 1.04128 | 0.057434 | 8.15E-05 | 1.048359 | 0.023188 | 0.680422 | 7.52E-50 | 1.224852 | 8.67E-10 | 0.837954 | 1.16E-22 |
| L89C A122D V196P | 1 | 1.039568 | 0.103734 | 1.41E-08 | 1.190961 | 3.99E-75 | 0.771544 | 5.20E-61 | 0.831676 | 5.87E-15 | 1.226516 | 7.10E-44 |
| V74W R78H L89C A122H V196P | 1 | 1.038873 | 0.063749 | 7.22E-05 | 1.016822 | 0.769376 | 0.620448 | 1.78E-114 | 0.775707 | 2.31E-66 | 0.994721 | 0.03166405 |
| Y63L N81S L89C A122D | 1 | 1.035985 | 0.143584 | 2.10E-09 | 0.836455 | 4.18E-95 | 0.764011 | 348022840 | 0.736834 | 4.93E-307 | 0.792572 | 3.07E-70 |
| N69E R78H N81S L89C A122D | 1 | 1.034793 | 0.098573 | 0.016588 | 0.813389 | 2.43E-44 | 1.012577 | 0.067677 | 0.733254 | 2.59E-22 | 0.645204 | 7.62E-52 |
| L89C A122D | 1 | 1.032703 | 0.10444 | 9.99E-13 | 1.072703 | 2.49E-24 | 0.651141 | 0 | 0.871042 | 2.82E-62 | 1.116849 | 8.79E-18 |
| N69E R78H A122H V196P | 1 | 1.030751 | 0.08888 | 0.000483 | 1.021719 | 0.011447 | 0.972939 | 1.10E-08 | 1.078824 | 3.99E-18 | 0.980762 | 0.048711196 |
| N69E R78H | 1 | 1.029205 | 0.092323 | 0.000334 | 1.022218 | 0.001361 | 1.015109 | 0.207593 | 1.230523 | 8.86E-53 | 0.947855 | 0.157585922 |
| Y63L L89C | 1 | 1.02749 | 0.103913 | 0.021235 | 1.083646 | 8.52E-09 | 0.535497 | 0 | 0.636653 | 0 | 1.087456 | 7.84E-06 |
| Y63L N69E R78H L89C A122D V196P | 1 | 1.019571 | 0.107465 | 0.495617 | 0.818142 | 1.69E-36 | 0.761988 | 7.05E-18 | 0.989391 | 0.863619 | 0.775642 | 1.06E-17 |
| Y63L N69E A122D V196P | 1 | 1.010975 | 0.087175 | 0.737677 | 1.02074 | 0.003618 | 0.925734 | 6.68E-24 | 0.827228 | 1.71E-128 | 1.039246 | 6.39E-05 |
| R78H A122H V196P | 1 | 1.002203 | 0.100623 | 0.808531 | 0.939774 | 0.000288 | 1.229619 | 3.15E-31 | 1.408653 | 9.73E-56 | 0.971521 | 0.222404832 |
| Y63L N69E N81S | 1 | 1.001762 | 0.115356 | 0.066055 | 0.841426 | 1.85E-75 | 1.028833 | 0.213396 | 1.246649 | 1.16E-35 | 0.850827 | 7.99E-16 |
| N69E R78H L89C | 1 | 0.997271 | 0.091509 | 0.978397 | 0.737825 | 3.54E-204 | 0.854603 | 8.61E-42 | 1.749287 | 0 | 0.587918 | 4.80E-289 |
| Y63L L89C A122H | 1 | 0.994943 | 0.091887 | 0.583093 | 1.101347 | 8.29E-11 | 0.718513 | 1.89E-146 | 0.722182 | 4.47E-84 | 1.233063 | 3.77E-29 |
| Y63L L89G A122D V196P | 1 | 0.994078 | 0.067157 | 0.489407 | 1.046659 | 0.043738 | 0.687103 | 1.79E-21 | 0.633852 | 8.79E-22 | 1.205166 | 0.012638153 |
| V74E V196P | 1 | 0.991191 | 0.067704 | 0.468258 | 0.797755 | 2.90E-45 | 1.115214 | 0.003148 | 1.076265 | 0.034437 | 0.536922 | 6.62E-71 |
| N69E V74E | 1 | 0.988972 | 0.096075 | 0.001957 | 0.917356 | 1.17E-19 | 1.023539 | 0.000172 | 1.11061 | 3.70E-17 | 0.929547 | 2.91E-08 |
| Y63L R78H L89C A122D | 1 | 0.988076 | 0.096833 | 0.32413 | 0.668311 | 1.62E-229 | 0.759982 | 1.46E-84 | 1.032156 | 0.428752 | 0.433545 | 2.91657523744909e-317 |
| Y63L N69E L89C A122H | 1 | 0.985519 | 0.099668 | 0.902729 | 0.848397 | 1.08E-70 | 0.817749 | 1.62E-88 | 1.508343 | 3.24E-307 | 0.655984 | 3.00E-230 |
| Y63L N69E L89C A122D V196P | 1 | 0.984917 | 0.064293 | 0.15398 | 0.871369 | 1.04E-11 | 0.88798 | 1.49E-06 | 0.855211 | 1.39E-07 | 0.803957 | 4.25E-17 |
| V74E R78H N81S V196P | 1 | 0.984578 | 0.111505 | 0.006454 | 0.917044 | 2.78E-18 | 0.964561 | 1.10E-08 | 0.993827 | 2.20E-09 | 0.956105 | 0.00703316 |
| R78H N81S L89C A122D V196P | 1 | 0.981448 | 0.062161 | 0.071859 | 0.85244 | 3.08E-68 | 0.71012 | 7.87E-102 | 0.798033 | 7.00E-47 | 0.637003 | 3.49E-237 |
| Y63L L89C V196P | 1 | 0.98069 | 0.07164 | 0.001266 | 1.028751 | 0.979434 | 0.433697 | 4.27E-187 | 0.711468 | 3.90E-75 | 1.034175 | 0.410516661 |
| Y63L V74E L89C V196P | 1 | 0.97866 | 0.074672 | 0.282539 | 0.766549 | 4.63E-43 | 0.929854 | 0.229037 | 1.214791 | 0.001275 | 0.586806 | 1.02E-33 |

| | | | | | | | | | | | | |
|---|---|---|---|---|---|---|---|---|---|---|---|---|
| Y63L R78H A122D V196P | 1 | 0.976019 | 0.151814 | 0.009227 | 0.773881 | 6.45E-90 | 0.886374 | 8.11E-05 | 0.903989 | 0.494086 | 0.572813 | 1.64E-135 |
| N69E R78H L89G A122D V196P | 1 | 0.974496 | 0.090024 | 0.004431 | 0.709387 | 1.51E-33 | 0.716993 | 1.73E-15 | 0.69502 | 5.03E-15 | 0.491718 | 1.97E-20 |
| V74E L89C V196P | 1 | 0.970716 | 0.059567 | 2.47E-05 | 0.972106 | 0.000124 | 0.673179 | 5.47E-92 | 0.84972 | 4.73E-28 | 1.043786 | 0.492637078 |
| N69E V74W R78H V196P | 1 | 0.969191 | 0.103087 | 0.00062 | 0.92118 | 8.15E-12 | 0.846881 | 1.30E-22 | 0.884991 | 3.67E-10 | 0.874118 | 7.68E-09 |
| V74W R78H N81S L89C A122H V196P | 1 | 0.96893 | 0.053443 | 0.123106 | 0.970553 | 0.023771 | 0.68994 | 4.61E-47 | 0.950495 | 0.969426 | 0.855533 | 4.08E-20 |
| Y63L N69E L89G A122H | 1 | 0.966973 | 0.086382 | 0.33803 | 0.832692 | 7.93E-24 | 0.571046 | 6.03E-54 | 0.678086 | 6.89E-29 | 0.640286 | 5.59E-22 |
| Y63L N81S L89C V196P | 1 | 0.965972 | 0.092017 | 0.329877 | 0.74375 | 8.50E-20 | 0.663494 | 1.34E-05 | 0.650767 | 7.81E-07 | 0.59665 | 6.91E-15 |
| N69E L89G A122D | 1 | 0.965969 | 0.086293 | 0.000402 | 0.938371 | 3.47E-12 | 0.655903 | 4.99E-118 | 0.6108 | 7.12E-147 | 0.922265 | 3.55E-07 |
| V74W R78H N81S L89C A122D V196P | 1 | 0.961697 | 0.053044 | 0.033573 | 0.856591 | 4.81E-10 | 0.678943 | 1.45E-15 | 0.857459 | 0.450175 | 0.636486 | 1.68E-29 |
| V74W R78H L89C V196P | 1 | 0.960649 | 0.058949 | 0.002039 | 0.811566 | 3.61E-35 | 0.877673 | 1.35E-06 | 1.03711 | 0.394627 | 0.670465 | 8.77E-41 |
| Y63L N69E L89G A122D | 1 | 0.959276 | 0.085695 | 0.000383 | 0.839704 | 2.76E-26 | 0.583237 | 1.38E-101 | 0.445286 | 1.44E-160 | 0.716798 | 6.34E-14 |
| N69E N81S | 1 | 0.953544 | 0.109804 | 3.16E-09 | 0.833266 | 6.02E-85 | 1.231109 | 4.09E-68 | 1.231479 | 1.44E-62 | 0.922834 | 1.26E-06 |
| N69E R78H L89C A122D | 1 | 0.951163 | 0.093216 | 3.41E-09 | 0.795766 | 2.24E-107 | 0.754614 | 3.26E-115 | 0.885874 | 2.94E-32 | 0.749851 | 3.24E-86 |
| V74E A122D | 1 | 0.950374 | 0.073353 | 2.31E-06 | 0.820806 | 2.93E-40 | 0.867578 | 1.64E-13 | 0.78908 | 1.63E-28 | 0.765781 | 2.80E-29 |
| N69E N81S L89T | 1 | 0.949179 | 0.098335 | 2.08E-17 | 1.00802 | 0.401677 | 0.633647 | 0 | 0.593217 | 0 | 1.005644 | 0.654407038 |
| Y63L N69E L89T A122D | 1 | 0.948997 | 0.101386 | 8.09E-13 | 1.036986 | 4.40E-05 | 0.512883 | 7.30E-257 | 0.490741 | 0 | 1.178572 | 1.36E-18 |
| Y63L A122H | 1 | 0.947497 | 0.086438 | 2.74E-14 | 0.827724 | 6.43E-70 | 1.079613 | 1.21E-13 | 1.026927 | 0.000755 | 0.719393 | 2.74E-71 |
| R78H N81S V196P | 1 | 0.941406 | 0.076598 | 1.18E-10 | 0.948387 | 4.53E-07 | 0.903221 | 2.17E-14 | 1.123519 | 1.79E-14 | 1.008831 | 0.069251579 |
| V74E L89G | 1 | 0.939776 | 0.110426 | 2.11E-06 | 0.888902 | 1.03E-23 | 0.653883 | 3.21E-73 | 0.486344 | 3.97E-192 | 1.037956 | 0.986645587 |
| R78H V196P | 1 | 0.935076 | 0.063871 | 1.76E-05 | 0.914108 | 2.00E-08 | 0.941695 | 0.000614 | 0.946719 | 0.045206 | 0.90582 | 3.22E-05 |
| R78H N81S A122D | 1 | 0.934064 | 0.083443 | 2.81E-31 | 0.868752 | 1.40E-44 | 1.51437 | 7.17E-270 | 1.532489 | 4.30E-230 | 0.904976 | 1.15E-05 |
| Y63L L89G A122H | 1 | 0.932028 | 0.083261 | 2.87E-07 | 0.953956 | 6.24E-08 | 0.800104 | 5.27E-32 | 0.69707 | 6.68E-54 | 0.839499 | 1.89E-12 |
| V74E N81S L89T V196P | 1 | 0.93155 | 0.064302 | 3.07E-10 | 1.028814 | 0.370923 | 0.35307 | 6.98E-270 | 0.792205 | 8.73E-22 | 1.051387 | 0.575671161 |
| Y63L N69E L89C A122H V196P | 1 | 0.930881 | 0.057123 | 1.73E-05 | 0.918646 | 7.39E-09 | 0.783853 | 4.52E-18 | 0.735854 | 1.02E-27 | 0.919896 | 1.33E-05 |
| V74W R78H L89C A122D | 1 | 0.929709 | 0.085862 | 1.87E-15 | 0.956411 | 6.69E-12 | 0.997421 | 0.175992 | 1.535552 | 4.87E-149 | 1.006271 | 0.720721476 |
| V74E N81S L89C | 1 | 0.927389 | 0.067687 | 5.46E-43 | 0.887514 | 5.00E-61 | 0.772445 | 3.08E-155 | 0.923958 | 2.14E-16 | 0.787646 | 4.35E-174 |
| N69E V74W R78H L89C A122H V196P | 1 | 0.920587 | 0.056491 | 8.94E-09 | 0.874186 | 1.64E-14 | 0.746591 | 4.07E-20 | 0.94056 | 9.41E-05 | 0.877871 | 2.36E-08 |
| Y63L N69E N81S L89C A122D | 1 | 0.919785 | 0.068603 | 7.12E-38 | 0.9904 | 0.130086 | 0.768167 | 2.41E-169 | 0.853289 | 3.38E-53 | 1.119116 | 2.27E-27 |
| V74E R78H V196P | 1 | 0.918706 | 0.081498 | 6.44E-14 | 0.999232 | 0.766974 | 0.899322 | 2.09E-11 | 0.756751 | 7.35E-40 | 1.110265 | 4.40E-08 |
| R78H L89C V196P | 1 | 0.917086 | 0.066993 | 3.63E-08 | 0.796727 | 3.24E-44 | 0.828851 | 1.64E-17 | 1.043295 | 0.369302 | 0.702273 | 2.52E-24 |
| Y63L N69E N81S V196P | 1 | 0.914358 | 0.085207 | 2.35E-24 | 0.940223 | 4.04E-10 | 1.137401 | 2.09E-27 | 1.071135 | 5.79E-06 | 1.029754 | 0.296923754 |
| N81S L89G | 1 | 0.913884 | 0.051123 | 1.68E-13 | 1.086321 | 0.00086 | 0.694764 | 3.94E-31 | 0.614626 | 7.54E-48 | 1.456131 | 3.30E-93 |
| Y63L N81S L89G A122H V196P | 1 | 0.912933 | 0.069657 | 0.085443 | 0.775006 | 2.24E-12 | 0.776545 | 3.48E-05 | 0.712851 | 0.000147 | 0.591035 | 8.86E-09 |
| Y63L N81S | 1 | 0.912373 | 0.107729 | 1.13E-44 | 0.805534 | 3.63E-119 | 0.945723 | 8.28E-12 | 0.813087 | 2.81E-74 | 0.821162 | 7.11E-51 |
| Y63L N69E L89G A122D V196P | 1 | 0.910354 | 0.084098 | 3.25E-26 | 1.044077 | 0.075784 | 0.613239 | 5.37E-308 | 0.61133 | 4.52E-237 | 1.126553 | 4.11E-09 |
| V74E R78H N81S | 1 | 0.909464 | 0.094507 | 5.28E-33 | 0.89447 | 3.65E-44 | 1.206329 | 2.28E-31 | 0.859123 | 1.05E-14 | 0.91561 | 2.54E-09 |
| Y63L N69E R78H N81S A122D | 1 | 0.908722 | 0.081179 | 1.76E-26 | 0.749036 | 1.15E-164 | 1.506171 | 3.59E-165 | 1.257018 | 2.19E-39 | 0.642159 | 2.03E-90 |
| Y63L N69E R78H | 1 | 0.90828 | 0.088236 | 8.98E-11 | 0.740083 | 5.94E-57 | 0.747686 | 7.46E-19 | 0.602684 | 1.29E-55 | 0.686016 | 6.72E-29 |
| N69E R78H L89C A122D V196P | 1 | 0.904424 | 0.059039 | 0.003235 | 0.875096 | 1.19E-13 | 0.95032 | 0.000577 | 1.066235 | 0.002179 | 0.785107 | 2.64E-22 |
| N69E V74W R78H N81S V196P | 1 | 0.903036 | 0.073995 | 2.12E-18 | 0.956015 | 0.008633 | 0.896363 | 0.000111 | 0.718875 | 2.52E-44 | 1.024899 | 0.102689408 |
| V74W R78H V196P | 1 | 0.902994 | 0.077864 | 8.33E-51 | 0.890993 | 4.78E-30 | 0.890134 | 4.00E-44 | 0.768445 | 2.36E-177 | 0.888616 | 9.91E-19 |
| Y63L R78H N81S L89C A122D V196P | 1 | 0.90239 | 0.113495 | 1.01E-46 | 0.894209 | 3.32E-60 | 0.889044 | 2.27E-24 | 0.717609 | 3.30E-114 | 1.088434 | 2.80E-09 |
| Y63L N81S L89C | 1 | 0.902337 | 0.06145 | 9.27E-21 | 0.918858 | 3.00E-10 | 0.934689 | 3.54E-09 | 0.957633 | 1.43E-06 | 0.91736 | 0.000140301 |
| V74W R78H A122H V196P | 1 | 0.902149 | 0.075228 | 1.54E-49 | 0.847692 | 1.38E-78 | 1.0255 | 0.03716 | 1.045519 | 0.003529 | 0.81695 | 5.89E-43 |
| N69E R78H N81S A122D | 1 | 0.901792 | 0.081804 | 1.16E-49 | 0.877002 | 3.88E-40 | 1.408917 | 4.96E-260 | 1.993117 | 0 | 0.751089 | 1.40E-95 |
| Y63L V74W N81S | 1 | 0.895598 | 0.101428 | 1.76E-51 | 0.756196 | 7.39E-86 | 1.15158 | 6.00E-07 | 1.358314 | 7.12E-41 | 0.727338 | 7.18E-18 |
| R78H N81S A122H V196P | 1 | 0.893898 | 0.068204 | 1.42E-08 | 0.737385 | 2.91E-28 | 0.947538 | 0.064345 | 1.145546 | 0.005827 | 0.692364 | 9.23E-08 |
| N69E L89T A122D | 1 | 0.892195 | 0.095317 | 5.05E-52 | 0.92577 | 1.99E-19 | 0.774393 | 4.71E-91 | 0.677926 | 3.12E-198 | 1.211721 | 1.16E-42 |

| | | | | | | | | | | | | |
|---|---|---|---|---|---|---|---|---|---|---|---|---|
| N81S L89C A122D | 1 | 0.891462 | 0.094819 | 1.20E-39 | 0.83903 | 3.32E-66 | 0.886966 | 1.26E-13 | 1.085182 | 2.63E-11 | 0.979845 | 0.01437813 |
| N69E V74E R78H | 1 | 0.889885 | 0.082207 | 6.22E-32 | 0.805064 | 1.19E-81 | 1.073858 | 7.67E-05 | 0.998764 | 0.775303 | 0.631619 | 6.49E-46 |
| Y63L L89G A122D | 1 | 0.889871 | 0.082925 | 7.43E-45 | 0.983888 | 4.41E-06 | 0.596978 | 3.94E-255 | 0.601619 | 4.86E-232 | 1.089723 | 0.017662857 |
| R78H N81S L89T A122D | 1 | 0.885633 | 0.091752 | 1.63E-91 | 0.984772 | 1.71E-06 | 0.853805 | 2.40E-117 | 0.937306 | 6.94E-14 | 1.06376 | 0.000302217 |
| N69E N81S L89C | 1 | 0.884176 | 0.111204 | 3.34E-46 | 0.908305 | 1.72E-26 | 0.892618 | 1.72E-16 | 0.936557 | 2.71E-05 | 1.322873 | 4.34E-66 |
| Y63L R78H A122D | 1 | 0.881492 | 0.068036 | 4.33E-31 | 0.791392 | 8.89E-71 | 0.971504 | 0.042472 | 0.73014 | 5.87E-44 | 0.818133 | 2.35E-27 |
| Y63L N69E R78H N81S A122D V196P | 1 | 0.879282 | 0.080343 | 1.45E-16 | 0.651424 | 2.10E-103 | 1.348643 | 2.15E-47 | 1.274949 | 1.27E-20 | 0.588237 | 6.62E-78 |
| Y63L N69E L89A | 1 | 0.876098 | 0.079473 | 6.03E-62 | 0.857975 | 1.24E-32 | 0.642714 | 6.17E-197 | 1.154284 | 2.32E-11 | 0.820736 | 1.15E-18 |
| V74W R78H | 1 | 0.875375 | 0.08504 | 1.36E-54 | 0.900245 | 8.59E-30 | 0.900463 | 1.58E-09 | 0.834717 | 2.78E-35 | 1.002037 | 0.779617434 |
| Y63L V74E N81S L89C V196P | 1 | 0.874125 | 0.065198 | 4.71E-61 | 0.786428 | 1.37E-103 | 0.846425 | 5.03E-25 | 0.938518 | 3.16E-10 | 0.733702 | 2.44E-64 |
| N69E R78H L89C A122H V196P | 1 | 0.87108 | 0.053453 | 5.26E-19 | 0.83055 | 2.04E-36 | 0.731649 | 2.09E-26 | 0.945866 | 0.021222 | 0.779162 | 2.84E-15 |
| Y63L A122D V196P | 1 | 0.868473 | 0.119839 | 7.42E-46 | 0.773971 | 3.72E-143 | 0.897562 | 4.08E-08 | 0.905643 | 0.005817 | 1.013959 | 0.245801024 |
| N69E R78H V196P | 1 | 0.865504 | 0.059119 | 2.03E-22 | 0.989596 | 0.737502 | 0.931051 | 0.000449 | 1.097484 | 0.000768 | 1.191188 | 1.33E-29 |
| Y63L R78H N81S L89C A122D | 1 | 0.861211 | 0.085334 | 4.00E-69 | 0.731626 | 1.03E-122 | 1.201394 | 1.13E-61 | 1.195104 | 1.56E-44 | 0.727908 | 5.66E-74 |
| N69E R78H N81S L89C A122D V196P | 1 | 0.860702 | 0.06282 | 3.50E-54 | 0.843136 | 4.94E-52 | 0.798869 | 1.11E-47 | 0.95452 | 0.00022 | 0.853867 | 2.19E-55 |
| N69E L89C A122D V196P | 1 | 0.859304 | 0.057236 | 1.40E-35 | 0.902899 | 2.12E-12 | 0.936622 | 0.00581 | 0.88169 | 1.19E-11 | 1.088918 | 0.000423404 |
| N69E V74E L89C A122D V196P | 1 | 0.858002 | 0.085616 | 4.71E-30 | 1.053459 | 0.038607 | 0.629606 | 4.39E-116 | 1.079869 | 1.50E-14 | 1.228041 | 1.56E-28 |
| V74W R78H A122H | 1 | 0.857132 | 0.079182 | 2.94E-73 | 0.903319 | 1.42E-21 | 1.011594 | 0.450513 | 0.900185 | 2.30E-18 | 0.977158 | 0.2489542 |
| V74W R78H N81S A122D V196P | 1 | 0.856572 | 0.059126 | 1.83E-17 | 0.767549 | 1.15E-62 | 1.038934 | 0.846449 | 0.961637 | 0.961125 | 0.682781 | 7.49E-48 |
| N81S L89T | 1 | 0.85538 | 0.088618 | 2.38E-189 | 0.959408 | 3.75E-10 | 0.626009 | 0 | 0.612566 | 0 | 1.138925 | 2.50E-28 |
| N69E L89C | 1 | 0.853172 | 0.066378 | 1.00E-96 | 0.926319 | 2.29E-19 | 0.988326 | 0.509052 | 1.340489 | 9.89E-101 | 1.151211 | 1.72E-23 |
| V74W L89G A122D V196P | 1 | 0.852859 | 0.077804 | 1.67E-56 | 0.896674 | 3.29E-27 | 0.5276 | 6.12E-250 | 0.573942 | 7.15E-216 | 1.104956 | 0.000817095 |
| V74W R78H A122D V196P | 1 | 0.85162 | 0.071015 | 3.67E-77 | 0.867484 | 1.06E-46 | 0.94702 | 0.001936 | 0.888493 | 1.12E-13 | 0.96243 | 0.010550039 |
| Y63L N69E N81S L89T | 1 | 0.85145 | 0.066244 | 2.17E-51 | 0.885623 | 5.48E-25 | 0.835464 | 2.54E-28 | 0.889296 | 1.27E-06 | 0.925547 | 3.47E-09 |
| Y63L R78H N81S A122D V196P | 1 | 0.849053 | 0.097809 | 1.39E-66 | 0.766025 | 6.63E-151 | 1.1918 | 5.78E-25 | 1.382131 | 2.21E-93 | 0.807976 | 4.14E-22 |
| V74E R78H | 1 | 0.847673 | 0.078308 | 1.49E-83 | 0.877008 | 1.03E-66 | 1.050944 | 1.16E-05 | 1.045803 | 8.06E-06 | 0.835699 | 1.24E-22 |
| L89G V196P | 1 | 0.845956 | 0.097414 | 2.48E-59 | 0.84759 | 3.91E-51 | 0.868812 | 4.07E-10 | 0.843478 | 1.13E-28 | 1.097527 | 1.17E-05 |
| Y63L N81S L89T V196P | 1 | 0.844617 | 0.05706 | 1.75E-10 | 0.947574 | 0.234772 | 0.559542 | 6.74E-30 | 0.788961 | 1.12E-06 | 1.346384 | 9.16E-08 |
| N69E V74W R78H N81S L89C A122H | 1 | 0.844431 | 0.057507 | 3.74E-58 | 0.820681 | 1.12E-51 | 1.12445 | 0.00029 | 1.189947 | 1.10E-19 | 0.895821 | 2.72E-13 |
| N69E V74W R78H A122H | 1 | 0.84277 | 0.077855 | 5.96E-68 | 0.929075 | 3.14E-17 | 1.033421 | 0.290829 | 0.980882 | 0.249194 | 0.947995 | 0.033788201 |
| Y63L L89A A122D V196P | 1 | 0.839293 | 0.077013 | 4.35E-91 | 0.738299 | 1.05E-153 | 0.686983 | 1.59E-181 | 0.899552 | 5.24E-22 | 0.821405 | 3.40E-37 |
| L89G A122D V196P | 1 | 0.834463 | 0.081065 | 6.67E-94 | 1.02959 | 0.848035 | 0.629613 | 4.81E-161 | 0.553653 | 1.39E-291 | 1.419246 | 4.21E-34 |
| Y63L L89T A122D | 1 | 0.834171 | 0.062874 | 1.65E-117 | 0.917178 | 4.47E-30 | 0.752038 | 2.40E-211 | 0.595975 | 0 | 1.448155 | 3.12E-171 |
| Y63L N69E L89T V196P | 1 | 0.830939 | 0.061498 | 3.50E-94 | 0.829472 | 1.01E-107 | 0.71904 | 2.92E-162 | 1.286691 | 1.29E-67 | 0.988729 | 0.888255658 |
| L89T V196P | 1 | 0.830016 | 0.06143 | 1.27E-97 | 0.884256 | 1.33E-35 | 0.71439 | 1.82E-177 | 0.960225 | 4.89E-05 | 1.13023 | 4.35E-25 |
| N69E V74W | 1 | 0.829901 | 0.07571 | 9.22E-115 | 0.813931 | 5.83E-99 | 0.950462 | 0.00103 | 0.99764 | 0.146397 | 0.936735 | 2.18E-09 |
| N69E V74E R78H V196P | 1 | 0.828146 | 0.063919 | 3.40E-49 | 0.888285 | 1.26E-19 | 0.880833 | 2.16E-14 | 0.552532 | 9.71E-156 | 1.187519 | 4.39E-08 |
| N69E R78H N81S L89C V196P | 1 | 0.827349 | 0.060385 | 1.82E-144 | 0.80097 | 1.14E-105 | 0.963856 | 0.105687 | 1.333433 | 3.74E-114 | 1.00309 | 0.980207872 |
| N69E V74W L89G V196P | 1 | 0.827237 | 0.082547 | 1.29E-26 | 0.942138 | 3.78E-05 | 0.895876 | 6.20E-09 | 1.172992 | 1.55E-10 | 1.152012 | 7.82E-08 |
| N69E V74W R78H L89C A122D V196P | 1 | 0.826239 | 0.108507 | 1.59E-120 | 0.856644 | 5.09E-43 | 0.84491 | 2.17E-47 | 0.949458 | 6.76E-11 | 1.019158 | 0.083674567 |
| N69E V74W R78H N81S | 1 | 0.826058 | 0.086346 | 2.51E-28 | 0.909645 | 3.07E-09 | 0.958662 | 0.018604 | 0.977761 | 0.410629 | 0.873117 | 5.87E-08 |
| N69E N81S V196P | 1 | 0.824688 | 0.096903 | 2.26E-46 | 0.690245 | 5.79E-138 | 0.90876 | 0.005542 | 0.873143 | 0.00036 | 0.874617 | 1.09E-07 |
| V74E R78H N81S A122D | 1 | 0.82386 | 0.073095 | 1.26E-13 | 0.660379 | 2.20E-40 | 1.274891 | 4.83E-10 | 1.077675 | 0.42078 | 0.622979 | 3.90E-21 |
| N69E L89G A122D V196P | 1 | 0.822474 | 0.079901 | 9.65E-113 | 1.024251 | 0.224204 | 0.617439 | 1.07E-270 | 0.743039 | 2.74E-93 | 1.282294 | 3.82E-26 |
| Y63L V74W R78H N81S L89C A122H V196P | 1 | 0.821959 | 0.045337 | 2.28E-22 | 0.841769 | 6.51E-17 | 0.583678 | 7.80E-30 | 0.922919 | 0.933074 | 0.840509 | 4.12E-11 |
| Y63L R78H L89C V196P | 1 | 0.820495 | 0.050349 | 6.72E-12 | 0.836132 | 7.69E-11 | 0.558938 | 3.41E-29 | 0.586663 | 5.35E-18 | 0.892795 | 0.005966154 |
| Y63L V74E L89C | 1 | 0.819506 | 0.080313 | 2.92E-136 | 0.745057 | 9.96E-153 | 0.785214 | 9.70E-115 | 1.173257 | 1.08E-33 | 0.907746 | 1.91E-18 |
| N69E V74W R78H N81S A122H V196P | 1 | 0.819337 | 0.08636 | 2.93E-63 | 0.841239 | 3.06E-37 | 1.065267 | 0.251677 | 1.257238 | 9.91E-32 | 0.854301 | 3.15E-18 |

| | | | | | | | | | | | | |
|---|---|---|---|---|---|---|---|---|---|---|---|---|
| N69E V74E R78H N81S V196P | 1 | 0.818847 | 0.067097 | 4.45E-57 | 0.908754 | 3.63E-12 | 0.983609 | 0.87601 | 0.900641 | 2.97E-06 | 1.052407 | 0.042775579 |
| V74W R78H L89C A122D V196P | 1 | 0.817602 | 0.053372 | 6.39E-23 | 0.839983 | 7.75E-19 | 0.986035 | 0.000254 | 1.045245 | 0.005543 | 0.90951 | 0.010205136 |
| V74W V196P | 1 | 0.817466 | 0.068167 | 7.58E-148 | 0.914843 | 3.00E-25 | 0.816693 | 2.91E-43 | 0.755838 | 7.32E-95 | 1.11749 | 3.46E-14 |
| Y63L N69E R78H L89C | 1 | 0.816407 | 0.074913 | 3.84E-82 | 0.698914 | 2.94E-198 | 0.702934 | 9.49E-101 | 1.152826 | 1.63E-13 | 0.773043 | 5.63E-46 |
| N69E N81S L89A | 1 | 0.815302 | 0.059924 | 1.84E-166 | 0.910323 | 1.61E-24 | 0.674448 | 1.84E-215 | 0.723613 | 1.75E-157 | 1.255478 | 6.37E-65 |
| Y63L N69E R78H N81S L89C | 1 | 0.812531 | 0.112614 | 1.73E-42 | 0.668056 | 3.00E-232 | 0.764742 | 4.48E-33 | 0.865528 | 7.92E-05 | 0.833409 | 0.015409578 |
| Y63L N69E R78H A122D V196P | 1 | 0.812299 | 0.055484 | 3.37E-33 | 0.909102 | 5.55E-09 | 0.805487 | 2.00E-06 | 0.953696 | 0.017712 | 1.109175 | 0.000411377 |
| Y63L L89A A122D | 1 | 0.811949 | 0.082615 | 4.61E-250 | 0.938426 | 1.25E-17 | 0.655093 | 0 | 0.711921 | 7.47E-190 | 1.150825 | 4.00E-18 |
| N69E V74W V196P | 1 | 0.811371 | 0.067659 | 7.97E-190 | 0.864554 | 9.97E-61 | 0.89145 | 4.03E-11 | 0.981942 | 0.004222 | 1.187305 | 2.75E-34 |
| V74E N81S V196P | 1 | 0.809934 | 0.074006 | 2.39E-113 | 0.742902 | 1.99E-145 | 1.156778 | 3.27E-28 | 1.316187 | 3.53E-87 | 0.750511 | 1.52E-94 |
| N69E V74E N81S L89C V196P | 1 | 0.806567 | 0.07155 | 7.60E-42 | 0.774634 | 5.07E-62 | 1.116247 | 4.10E-07 | 0.784429 | 1.10E-17 | 0.810494 | 1.08E-17 |
| N69E V74W R78H L89C A122H | 1 | 0.8065 | 0.081563 | 6.29E-47 | 0.805997 | 1.70E-33 | 0.77747 | 1.02E-46 | 0.792219 | 5.57E-44 | 0.900183 | 7.72E-05 |
| Y63L A122D | 1 | 0.805976 | 0.094705 | 1.08E-94 | 0.767362 | 5.12E-150 | 1.245091 | 3.33E-42 | 0.755517 | 4.88E-34 | 0.830838 | 5.63E-39 |
| N69E V74W R78H A122H V196P | 1 | 0.805362 | 0.101292 | 1.60E-73 | 0.842763 | 2.25E-58 | 0.863422 | 4.40E-17 | 0.821763 | 3.32E-18 | 1.174595 | 6.68E-15 |
| Y63L V74E N81S V196P | 1 | 0.805047 | 0.07356 | 2.12E-49 | 0.621484 | 3.90E-139 | 1.096583 | 1.23E-07 | 1.123103 | 6.73E-07 | 0.580703 | 1.16E-82 |
| N69E V74E L89C | 1 | 0.804475 | 0.063439 | 4.32E-78 | 0.838946 | 9.97E-38 | 0.886915 | 1.15E-15 | 1.087177 | 1.82E-06 | 1.057554 | 0.000996302 |
| V74E N81S | 1 | 0.803967 | 0.094002 | 3.47E-128 | 0.878235 | 2.37E-40 | 1.138294 | 1.28E-20 | 1.110247 | 3.49E-25 | 1.104689 | 0.341635874 |
| N69E V74E L89T | 1 | 0.802807 | 0.06051 | 4.13E-146 | 0.879606 | 9.22E-52 | 0.56534 | 0 | 0.765583 | 4.45E-75 | 1.414455 | 3.34E-146 |
| N81S V196P | 1 | 0.802549 | 0.063287 | 7.62E-139 | 0.76858 | 1.66E-108 | 1.45106 | 3.85E-164 | 1.294897 | 2.41E-78 | 0.903863 | 5.10E-08 |
| N69E R78H L89T A122D | 1 | 0.800219 | 0.085491 | 2.34E-65 | 0.757494 | 2.14E-99 | 0.762984 | 2.84E-25 | 0.780074 | 4.36E-22 | 0.929039 | 0.001175802 |
| Y63L N69E V74E L89T | 1 | 0.799951 | 0.054641 | 1.46E-07 | 0.823627 | 1.14E-05 | 0.218188 | 7.97E-12 | 1.121705 | 0.780919 | 1.227233 | 0.09077162 |
| Y63L N69E V74E N81S L89C A122D | 1 | 0.799713 | 0.070942 | 4.96E-33 | 0.709439 | 2.05E-52 | 1.260813 | 1.97E-12 | 1.206932 | 4.61E-06 | 0.780288 | 3.19E-11 |
| N69E R78H L89C A122H | 1 | 0.799582 | 0.063962 | 1.37E-86 | 0.670643 | 2.35E-152 | 1.512305 | 1.21E-103 | 1.534393 | 6.05E-101 | 0.672959 | 1.31E-96 |
| Y63L L89A A122H | 1 | 0.799289 | 0.09052 | 1.26E-102 | 0.821436 | 4.54E-76 | 0.774778 | 1.72E-67 | 0.689437 | 8.10E-98 | 1.154361 | 4.81E-05 |
| V74W R78H N81S V196P | 1 | 0.798501 | 0.063819 | 3.71E-104 | 0.861952 | 1.14E-28 | 0.891107 | 8.78E-10 | 0.602666 | 1.69E-87 | 1.068291 | 0.138906658 |
| Y63L N69E L89T | 1 | 0.797502 | 0.100303 | 2.96E-107 | 0.8594 | 1.41E-70 | 0.789987 | 5.16E-53 | 0.733263 | 8.09E-49 | 1.226062 | 1.02E-23 |
| Y63L N69E R78H L89C A122D | 1 | 0.796909 | 0.078099 | 1.16E-106 | 0.766134 | 2.89E-118 | 0.662412 | 1.92E-241 | 0.613715 | 3.44E-232 | 0.742669 | 7.59E-44 |
| V74W R78H N81S A122H V196P | 1 | 0.794297 | 0.093787 | 6.14E-209 | 0.774051 | 7.21E-149 | 0.921187 | 2.74E-14 | 0.835911 | 4.07E-48 | 0.992425 | 0.002020771 |
| N69E V74W R78H A122D V196P | 1 | 0.793747 | 0.061264 | 5.22E-157 | 0.853775 | 1.51E-54 | 1.2395 | 2.81E-40 | 1.137113 | 2.42E-19 | 1.156549 | 1.24E-21 |
| N69E A122H | 1 | 0.79285 | 0.07233 | 2.30E-259 | 0.780284 | 2.42E-173 | 1.276541 | 2.97E-135 | 1.586626 | 0 | 1.020486 | 0.086934063 |
| L89A A122D | 1 | 0.792712 | 0.081101 | 2.93E-305 | 1.007256 | 0.043248 | 0.661302 | 1.50E-254 | 0.491264 | 0 | 1.650974 | 5.30E-236 |
| V74W R78H N81S L89C A122D | 1 | 0.792645 | 0.109858 | 1.50E-79 | 0.631961 | 0 | 0.913202 | 1.01E-30 | 0.798682 | 6.57E-89 | 0.667411 | 1.91E-97 |
| V74W R78H L89C A122H | 1 | 0.790923 | 0.077512 | 4.75E-127 | 0.755013 | 1.16E-123 | 0.746362 | 2.70E-121 | 0.738351 | 1.16E-122 | 0.956086 | 0.00041043 |
| N69E V74W R78H N81S L89C V196P | 1 | 0.790582 | 0.07531 | 2.07E-50 | 0.841301 | 5.71E-24 | 1.13859 | 2.27E-05 | 0.98851 | 0.949726 | 1.044667 | 0.523989443 |
| N69E V74E R78H L89C A122D | 1 | 0.790243 | 0.079919 | 1.37E-17 | 0.669718 | 2.95E-56 | 0.603473 | 1.66E-40 | 0.835024 | 2.87E-11 | 0.603816 | 5.78E-37 |
| Y63L N69E N81S L89T V196P | 1 | 0.789727 | 0.054512 | 1.19E-57 | 0.853054 | 2.91E-32 | 0.617951 | 1.53E-55 | 0.758609 | 1.15E-18 | 0.979611 | 0.022172849 |
| V74E R78H N81S L89C | 1 | 0.785538 | 0.05859 | 2.00E-164 | 0.784738 | 3.58E-103 | 0.979862 | 0.03936 | 1.030628 | 0.008185 | 0.949011 | 1.48E-05 |
| Y63L N69E V74E L89C A122D | 1 | 0.78511 | 0.072508 | 4.22E-58 | 0.833811 | 2.30E-24 | 0.94438 | 0.000242 | 1.402467 | 9.21E-44 | 1.057595 | 0.027792571 |
| Y63L V74E N81S | 1 | 0.782307 | 0.108425 | 2.73E-70 | 0.702795 | 7.96E-198 | 1.262495 | 1.86E-67 | 1.478894 | 4.64E-66 | 0.948597 | 0.001408062 |
| Y63L V74W A122H | 1 | 0.781967 | 0.072238 | 2.70E-27 | 0.695043 | 5.98E-47 | 0.894317 | 6.95E-05 | 0.94417 | 0.546053 | 0.610529 | 1.41E-15 |
| N69E V74W R78H N81S A122D V196P | 1 | 0.78169 | 0.062475 | 9.44E-32 | 0.722681 | 5.31E-47 | 1.403676 | 8.60E-28 | 1.05843 | 0.000668 | 0.705747 | 6.37E-24 |
| Y63L N69E N81S A122H | 1 | 0.7815 | 0.077983 | 1.14E-22 | 0.779452 | 6.32E-33 | 1.46207 | 8.90E-11 | 1.460947 | 3.92E-17 | 0.97885 | 0.589020671 |
| V74E L89G A122D | 1 | 0.780472 | 0.077874 | 4.48E-31 | 0.841024 | 5.66E-06 | 0.844176 | 0.001138 | 0.660881 | 9.17E-23 | 1.037245 | 0.076350487 |
| N69E V74E L89C V196P | 1 | 0.779594 | 0.086056 | 5.94E-168 | 0.785801 | 5.14E-108 | 0.726955 | 3.64E-166 | 0.833687 | 2.68E-28 | 0.959727 | 0.016318901 |
| Y63L R78H L89C A122D V196P | 1 | 0.779021 | 0.051888 | 1.66E-26 | 0.861492 | 1.53E-10 | 0.819831 | 6.10E-11 | 0.763275 | 5.13E-14 | 0.870704 | 2.16E-05 |
| R78H L89C | 1 | 0.778358 | 0.066506 | 3.57E-96 | 0.90477 | 7.94E-19 | 0.818279 | 1.09E-21 | 1.174186 | 2.40E-14 | 1.089996 | 0.000995305 |
| Y63L N69E V74E L89C A122H V196P | 1 | 0.77816 | 0.047751 | 3.81E-12 | 0.751782 | 2.24E-22 | 0.454585 | 8.06E-25 | 0.774535 | 0.000287 | 0.907638 | 0.000517389 |
| Y63L N69E N81S L89C | 1 | 0.778156 | 0.077104 | 8.46E-198 | 0.764355 | 2.89E-110 | 0.90715 | 2.95E-12 | 0.892198 | 2.31E-14 | 0.948747 | 0.000246451 |

| | | | | | | | | | | | | |
|---|---|---|---|---|---|---|---|---|---|---|---|---|
| N69E V74E L89C A122H | 1 | 0.777077 | 0.066397 | 2.82E-45 | 0.691317 | 4.60E-72 | 0.852712 | 3.17E-08 | 1.043783 | 0.083073 | 0.932575 | 0.027417885 |
| V74E R78H N81S L89C A122D V196P | 1 | 0.776366 | 0.042822 | 4.33E-18 | 0.8058 | 4.17E-14 | 0.793215 | 3.74E-09 | 0.977793 | 0.490782 | 0.800376 | 1.74E-09 |
| N69E V74W R78H | 1 | 0.776013 | 0.091184 | 1.93E-139 | 0.772422 | 5.27E-134 | 0.936422 | 7.89E-05 | 0.756885 | 2.02E-18 | 0.840924 | 1.32E-28 |
| Y63L L89A | 1 | 0.775371 | 0.061144 | 1.97E-49 | 0.74172 | 1.65E-59 | 0.910113 | 1.49E-05 | 1.106375 | 0.006091 | 0.86526 | 1.87E-13 |
| N69E L89T | 1 | 0.775267 | 0.058434 | 5.08E-229 | 0.862018 | 1.62E-67 | 0.786384 | 4.17E-135 | 0.820827 | 4.79E-60 | 1.563165 | 1.27E-260 |
| Y63L N81S L89T | 1 | 0.774732 | 0.09361 | 5.21E-279 | 0.847977 | 2.57E-81 | 0.726322 | 3.90E-245 | 0.788783 | 7.45E-84 | 1.253879 | 1.91E-81 |
| N81S L89T V196P | 1 | 0.774303 | 0.077924 | 1.02E-93 | 1.077951 | 0.002561 | 0.513955 | 1.02E-128 | 0.781534 | 3.19E-17 | 1.496918 | 4.10E-65 |
| Y63L L89A V196P | 1 | 0.77239 | 0.061787 | 9.37E-165 | 0.82387 | 1.12E-48 | 0.763628 | 3.51E-55 | 0.828849 | 1.15E-32 | 1.151977 | 1.55E-09 |
| Y63L N81S A122D V196P | 1 | 0.772363 | 0.06173 | 4.93E-52 | 0.688744 | 8.48E-95 | 1.118985 | 3.51E-06 | 0.845911 | 2.87E-07 | 0.76797 | 1.91E-29 |
| L89G A122D | 1 | 0.771358 | 0.071881 | 7.27E-231 | 0.971177 | 1.66E-06 | 0.797692 | 1.65E-66 | 0.838204 | 2.60E-35 | 1.377952 | 8.39E-72 |
| Y63L N69E A122H | 1 | 0.7701 | 0.090489 | 5.00E-116 | 0.708714 | 1.77E-185 | 1.305841 | 7.43E-41 | 0.97227 | 0.130138 | 0.750034 | 1.82E-57 |
| Y63L V74E N81S L89C A122D V196P | 1 | 0.769722 | 0.043059 | 1.63E-19 | 0.738872 | 4.68E-24 | 1.144547 | 0.176147 | 1.419758 | 0.001811 | 0.858345 | 0.000121281 |
| N69E R78H A122H | 1 | 0.767226 | 0.068822 | 1.49E-155 | 0.906941 | 4.87E-13 | 1.014585 | 0.15577 | 1.297936 | 5.57E-62 | 1.285485 | 2.44E-34 |
| Y63L N69E N81S L89C A122H | 1 | 0.766571 | 0.052204 | 4.84E-115 | 0.801435 | 6.95E-65 | 0.918715 | 5.55E-05 | 1.267083 | 3.55E-27 | 0.971387 | 0.327801327 |
| R78H N81S L89C V196P | 1 | 0.766296 | 0.052186 | 5.58E-149 | 0.79706 | 1.01E-78 | 1.120967 | 3.09E-12 | 1.742565 | 3.04E-135 | 1.032151 | 6.94E-06 |
| R78H N81S L89A A122D V196P | 1 | 0.765752 | 0.055428 | 8.25E-100 | 0.95409 | 5.09E-06 | 0.508879 | 2.70E-125 | 0.671562 | 3.26E-46 | 1.666989 | 2.24E-85 |
| N69E L89G A122H | 1 | 0.765671 | 0.0684 | 3.33E-44 | 0.734962 | 3.06E-58 | 1.079322 | 0.00691 | 0.783753 | 2.97E-11 | 0.790925 | 2.26E-09 |
| Y63L N69E N81S L89C V196P | 1 | 0.764546 | 0.055802 | 3.76E-248 | 0.720418 | 1.40E-266 | 0.779214 | 3.57E-70 | 1.051834 | 2.42E-05 | 0.722481 | 2.57E-250 |
| V74W R78H N81S L89C A122H | 1 | 0.762213 | 0.051907 | 6.47E-155 | 0.795255 | 6.49E-87 | 1.056855 | 0.002591 | 1.225462 | 3.66E-25 | 1.068884 | 4.21E-05 |
| N81S L89A | 1 | 0.762008 | 0.056007 | 0 | 0.985654 | 1.88E-07 | 0.818379 | 2.07E-138 | 0.702608 | 1.60E-277 | 1.693602 | 0 |
| Y63L N69E R78H N81S L89C V196P | 1 | 0.761917 | 0.059278 | 3.72E-111 | 0.720122 | 4.63E-157 | 1.128439 | 4.66E-05 | 1.731161 | 4.93E-103 | 0.770258 | 2.70E-56 |
| N69E V74E A122D | 1 | 0.761745 | 0.068331 | 9.19E-17 | 0.585175 | 3.04E-35 | 1.769694 | 3.27E-15 | 0.957243 | 0.958246 | 0.576116 | 8.51E-11 |
| Y63L N81S L89G | 1 | 0.761537 | 0.042601 | 3.67E-44 | 0.862849 | 5.93E-22 | 1.138272 | 0.21156 | 0.796074 | 5.09E-06 | 1.115268 | 0.00042298 |
| Y63L N69E V74W R78H N81S A122H V196P | 1 | 0.759865 | 0.058732 | 6.91E-27 | 0.598766 | 3.36E-86 | 1.513185 | 9.78E-39 | 1.14755 | 4.37E-07 | 0.48219 | 8.99E-30 |
| Y63L N69E V74W L89C A122H V196P | 1 | 0.759732 | 0.04662 | 9.04E-18 | 0.798079 | 6.36E-17 | 0.765662 | 2.48E-07 | 0.746077 | 2.38E-08 | 0.973264 | 0.052973315 |
| Y63L N69E N81S L89G | 1 | 0.759542 | 0.041894 | 9.61E-57 | 0.823642 | 4.67E-29 | 0.681592 | 5.79E-33 | 0.907245 | 0.400679 | 0.935958 | 0.043624443 |
| N69E V74E L89G V196P | 1 | 0.759379 | 0.087445 | 1.02E-62 | 0.820597 | 1.24E-33 | 0.686398 | 1.04E-29 | 1.056582 | 0.667105 | 1.165154 | 8.49E-05 |
| Y63L N69E V74E N81S L89C V196P | 1 | 0.759288 | 0.056633 | 6.00E-213 | 0.680182 | 7.46E-249 | 1.048176 | 0.11555 | 1.030868 | 0.096813 | 0.690122 | 2.57E-91 |
| Y63L N69E V196P | 1 | 0.759259 | 0.076528 | 3.25E-176 | 0.792475 | 5.53E-85 | 0.794253 | 6.89E-71 | 0.743623 | 1.51E-36 | 0.873005 | 9.23E-13 |
| N69E N81S L89G | 1 | 0.758071 | 0.095344 | 7.02E-125 | 0.867249 | 9.64E-30 | 0.665868 | 2.47E-76 | 0.761575 | 1.36E-32 | 1.162742 | 8.82E-13 |
| R78H L89C A122H V196P | 1 | 0.757517 | 0.055337 | 7.40E-171 | 0.954271 | 7.80E-07 | 0.655468 | 4.94E-121 | 1.165119 | 6.98E-17 | 1.464186 | 2.56E-67 |
| V74E N81S L89C V196P | 1 | 0.756782 | 0.055235 | 0 | 0.797013 | 1.98E-198 | 1.030406 | 0.000101 | 1.26534 | 1.45E-139 | 1.158147 | 1.88E-72 |
| N69E R78H N81S L89G A122D | 1 | 0.756543 | 0.078326 | 2.59E-172 | 0.768999 | 2.70E-107 | 0.661906 | 5.47E-85 | 0.931605 | 1.00E-06 | 1.105209 | 5.50E-10 |
| N69E V74W A122H V196P | 1 | 0.756318 | 0.058375 | 7.16E-117 | 0.830567 | 1.04E-40 | 0.95206 | 0.003096 | 0.852696 | 1.95E-12 | 1.151561 | 1.66E-10 |
| V74E R78H L89C A122H V196P | 1 | 0.754086 | 0.046274 | 1.48E-21 | 0.702638 | 6.41E-33 | 0.641557 | 1.54E-09 | 1.151636 | 0.043452 | 0.921355 | 0.029313374 |
| V74W L89C V196P | 1 | 0.75361 | 0.055051 | 1.30E-61 | 0.86808 | 2.97E-19 | 0.654233 | 5.40E-49 | 0.908543 | 0.0001 | 1.169133 | 0.000218488 |
| V74E L89A | 1 | 0.752747 | 0.077985 | 0 | 0.854683 | 3.30E-120 | 0.696506 | 5.53E-235 | 0.761775 | 9.94E-140 | 1.21277 | 4.11E-44 |
| Y63L N81S L89C A122D V196P | 1 | 0.751989 | 0.047628 | 3.80E-56 | 0.764183 | 4.16E-51 | 0.823542 | 1.07E-06 | 0.830773 | 8.01E-12 | 0.847802 | 4.70E-15 |
| N69E V74W L89C V196P | 1 | 0.751773 | 0.075016 | 3.82E-17 | 0.897573 | 2.11E-05 | 0.75551 | 2.48E-09 | 0.776997 | 0.035207 | 0.998553 | 0.369328527 |
| Y63L N81S A122D | 1 | 0.751128 | 0.069996 | 2.73E-85 | 0.74048 | 8.04E-72 | 1.087634 | 0.000528 | 0.936171 | 0.041364 | 0.881799 | 1.41E-11 |
| N69E R78H N81S L89A A122D | 1 | 0.751032 | 0.094459 | 1.18E-77 | 0.910662 | 6.02E-19 | 0.727739 | 3.29E-33 | 1.045979 | 0.010205 | 1.37223 | 1.72E-25 |
| Y63L N69E V74W R78H N81S A122H | 1 | 0.750971 | 0.079154 | 2.73E-29 | 0.645519 | 1.18E-59 | 1.046509 | 0.496451 | 1.574257 | 3.11E-14 | 0.705371 | 9.42E-18 |
| Y63L N69E V74E A122D | 1 | 0.750521 | 0.089865 | 3.73E-17 | 0.652208 | 2.79E-33 | 1.924689 | 1.05E-15 | 1.144234 | 0.314586 | 0.845698 | 0.052712309 |
| Y63L N69E L89A A122D | 1 | 0.749626 | 0.073092 | 8.14E-129 | 0.839689 | 2.28E-49 | 0.757679 | 1.10E-45 | 0.752361 | 2.26E-42 | 1.139597 | 9.94E-11 |
| N69E R78H N81S L89A A122D V196P | 1 | 0.749118 | 0.04912 | 2.49E-141 | 0.783365 | 1.61E-72 | 0.798823 | 1.70E-39 | 0.785421 | 1.86E-28 | 1.006338 | 0.68775301 |
| R78H L89G A122D V196P | 1 | 0.749088 | 0.069201 | 1.43E-51 | 0.746157 | 1.95E-53 | 1.048499 | 0.800059 | 1.057881 | 0.245611 | 0.762412 | 1.76E-06 |
| V74E R78H A122H V196P | 1 | 0.747689 | 0.051071 | 8.71E-41 | 0.743773 | 2.19E-40 | 1.05702 | 0.124579 | 0.808703 | 8.23E-07 | 0.926554 | 0.008335337 |
| N69E V74E R78H L89C A122D V196P | 1 | 0.747337 | 0.078118 | 8.60E-40 | 0.710582 | 7.45E-72 | 0.970553 | 0.106429 | 1.482894 | 6.86E-23 | 0.863157 | 0.001685041 |

| | | | | | | | | | | | | |
|---|---|---|---|---|---|---|---|---|---|---|---|---|
| Y63L N69E N81S L89G V196P | 1 | 0.746109 | 0.085917 | 6.20E-41 | 0.698965 | 9.01E-91 | 0.820647 | 4.26E-07 | 0.813951 | 1.08E-10 | 0.905668 | 0.000232676 |
| Y63L L89T V196P | 1 | 0.744497 | 0.056115 | 1.33E-269 | 0.804371 | 9.96E-130 | 0.85745 | 7.35E-26 | 0.946872 | 1.69E-06 | 1.290435 | 1.17E-75 |
| V74E R78H N81S L89C V196P | 1 | 0.743547 | 0.055458 | 9.39E-300 | 0.765152 | 3.13E-174 | 0.979218 | 0.236007 | 1.178199 | 9.51E-25 | 1.030262 | 0.429996256 |
| Y63L N69E V74E L89C A122D V196P | 1 | 0.743546 | 0.074195 | 1.22E-46 | 0.825876 | 5.61E-39 | 1.304922 | 1.54E-12 | 1.014314 | 0.823327 | 0.972466 | 0.584116152 |
| N69E V74W R78H N81S L89C A122D V196P | 1 | 0.743536 | 0.059235 | 1.38E-226 | 0.611686 | 0 | 1.132577 | 2.17E-09 | 1.141086 | 1.23E-09 | 0.524408 | 1.99E-283 |
| N69E V74W R78H N81S L89C A122D | 1 | 0.742931 | 0.055413 | 1.84E-254 | 0.807282 | 2.73E-94 | 0.867778 | 4.89E-22 | 1.040628 | 0.597124 | 1.150526 | 4.37E-09 |
| R78H N81S L89G A122D | 1 | 0.742162 | 0.085496 | 5.00E-103 | 0.917239 | 8.33E-14 | 0.666902 | 1.32E-142 | 0.63509 | 5.69E-176 | 1.247967 | 1.81E-13 |
| Y63L N69E A122H V196P | 1 | 0.741616 | 0.063949 | 9.26E-228 | 0.713779 | 1.24E-223 | 1.248341 | 7.05E-59 | 1.197934 | 1.52E-41 | 0.764965 | 1.01E-78 |
| Y63L N81S L89G A122D V196P | 1 | 0.740938 | 0.073935 | 8.05E-12 | 1.104254 | 0.074026 | 0.578869 | 1.57E-13 | 0.611849 | 3.01E-05 | 1.321617 | 0.000295258 |
| Y63L V74W A122H V196P | 1 | 0.740042 | 0.061711 | 1.18E-93 | 0.812161 | 2.38E-42 | 0.844849 | 6.04E-11 | 1.03248 | 0.046277 | 1.000803 | 0.466156692 |
| L89A V196P | 1 | 0.739747 | 0.075682 | 0 | 0.986837 | 0.000348 | 0.744003 | 2.16E-148 | 0.655885 | 1.22E-226 | 1.783839 | 4.82E-306 |
| Y63L N69E V74W L89G A122H | 1 | 0.738442 | 0.068814 | 2.37E-16 | 0.574504 | 1.57E-61 | 1.150124 | 0.004213 | 0.760712 | 0.000423 | 0.451413 | 9.92E-22 |
| Y63L N69E R78H N81S V196P | 1 | 0.737458 | 0.068722 | 7.15E-80 | 0.678216 | 2.61E-114 | 1.049862 | 0.006369 | 1.083882 | 0.010112 | 0.748405 | 1.61E-20 |
| N81S L89C V196P | 1 | 0.736325 | 0.050145 | 2.56E-206 | 0.749532 | 1.48E-118 | 0.9342 | 0.001748 | 1.277535 | 2.86E-31 | 0.994802 | 0.986930724 |
| N69E R78H N81S L89C A122H | 1 | 0.734561 | 0.053613 | 1.26E-307 | 0.710168 | 1.51E-243 | 0.865061 | 3.04E-28 | 1.549823 | 1.34E-156 | 0.781218 | 7.57E-140 |
| Y63L N81S L89C A122H | 1 | 0.732069 | 0.073788 | 894771153 | 0.725834 | 4.24E-279 | 0.891795 | 1.48E-31 | 0.884079 | 3.00E-05 | 0.873282 | 5.61E-26 |
| Y63L N69E V74W R78H N81S L89C A122H V196P | 1 | 0.731255 | 0.040334 | 5.05E-23 | 0.680053 | 6.19E-24 | 0.905164 | 0.023269 | 2.30808 | 8.88E-13 | 0.686092 | 2.17E-14 |
| V74E L89C A122D | 1 | 0.730623 | 0.057615 | 4.41E-198 | 0.733294 | 7.17E-136 | 1.382158 | 1.90E-71 | 1.586029 | 1.89E-93 | 0.914307 | 7.69E-07 |
| Y63L R78H N81S L89C V196P | 1 | 0.730294 | 0.053302 | 8.71E-177 | 0.676214 | 4.48E-197 | 0.905909 | 3.44E-06 | 1.09185 | 7.37E-06 | 0.67363 | 5.00E-191 |
| R78H N81S L89T A122D V196P | 1 | 0.725366 | 0.049003 | 2.26E-11 | 0.869718 | 6.13E-07 | 0.716324 | 0.000237 | 1.005749 | 0.096582 | 1.217318 | 0.097049358 |
| L89T A122D | 1 | 0.724445 | 0.054604 | 420923624 | 0.890879 | 7.38E-60 | 0.889375 | 6.36E-25 | 0.793487 | 3.91E-92 | 1.745232 | 0 |
| V74E L89T | 1 | 0.72364 | 0.068414 | 0 | 0.827097 | 3.73E-105 | 0.696291 | 2.03E-160 | 0.817508 | 3.36E-49 | 1.355582 | 4.63E-141 |
| N69E V74E L89G | 1 | 0.723635 | 0.056299 | 4.55E-115 | 0.735472 | 1.93E-104 | 0.982548 | 0.791859 | 0.843828 | 7.77E-09 | 0.862658 | 5.61E-18 |
| Y63L N69E V74W L89C A122D V196P | 1 | 0.721688 | 0.083105 | 1.69E-137 | 0.82499 | 6.38E-70 | 0.771042 | 1.46E-28 | 1.090943 | 0.012694 | 1.172123 | 2.75E-15 |
| Y63L V74W N81S L89C A122H V196P | 1 | 0.721082 | 0.040338 | 9.07E-65 | 0.840896 | 5.98E-28 | 1.09932 | 0.392394 | 1.064005 | 0.450019 | 1.250897 | 6.06E-16 |
| Y63L N81S V196P | 1 | 0.719631 | 0.084559 | 5.98E-106 | 0.63058 | 6.10E-185 | 1.392387 | 2.33E-50 | 0.60026 | 1.84E-60 | 0.906091 | 5.92E-08 |
| N69E V74W A122D V196P | 1 | 0.719017 | 0.049113 | 1.16E-49 | 0.775058 | 1.01E-25 | 0.807152 | 4.88E-11 | 0.779637 | 2.42E-09 | 1.116919 | 0.000296888 |
| N69E V74W L89G A122D V196P | 1 | 0.718822 | 0.076456 | 2.88E-52 | 0.767476 | 2.26E-32 | 0.786122 | 2.18E-05 | 0.60596 | 2.87E-28 | 1.049394 | 0.387090366 |
| Y63L A122H V196P | 1 | 0.718819 | 0.072452 | 3.78E-248 | 0.71543 | 7.69E-232 | 0.960539 | 6.21E-05 | 0.834523 | 1.74E-12 | 0.85926 | 3.35E-25 |
| V74E R78H L89C | 1 | 0.718701 | 0.066375 | 5.00E-186 | 0.863683 | 1.86E-51 | 0.956278 | 1.95E-07 | 0.895472 | 1.30E-07 | 1.552846 | 6.27E-116 |
| Y63L N69E V74W N81S A122D V196P | 1 | 0.717802 | 0.049547 | 1.94E-06 | 0.647182 | 4.45E-26 | 0.858453 | 0.003666 | 0.886334 | 0.297037 | 0.500466 | 1.75E-13 |
| Y63L N69E L89G A122H V196P | 1 | 0.717336 | 0.069687 | 4.72E-79 | 0.829488 | 4.61E-30 | 0.848391 | 5.89E-09 | 0.675507 | 3.56E-39 | 1.069788 | 0.06134567 |
| L89C A122H V196P | 1 | 0.716142 | 0.043945 | 1.90E-164 | 0.81786 | 3.08E-70 | 0.848827 | 1.18E-14 | 1.071685 | 0.004429 | 1.234009 | 1.66E-24 |
| N69E V74W L89G A122H V196P | 1 | 0.715914 | 0.076147 | 3.42E-18 | 0.592816 | 4.50E-51 | 0.947939 | 0.658295 | 0.708678 | 7.18E-08 | 0.666312 | 5.07E-09 |
| V74E N81S A122D | 1 | 0.714773 | 0.058569 | 6.36E-52 | 0.751086 | 3.00E-35 | 1.06738 | 0.025313 | 1.359329 | 1.47E-11 | 0.966378 | 0.004147003 |
| V74E N81S L89C A122D | 1 | 0.714383 | 0.071725 | 1.01E-131 | 0.892586 | 3.56E-13 | 0.926235 | 0.005212 | 1.071103 | 0.26884 | 1.518454 | 3.04E-29 |
| V74W L89C A122H | 1 | 0.712439 | 0.060874 | 4.31E-32 | 0.770859 | 7.67E-15 | 0.782511 | 3.07E-07 | 0.712218 | 3.02E-08 | 1.00875 | 0.620794763 |
| N69E V74W R78H N81S A122D | 1 | 0.712162 | 0.061335 | 5.59E-275 | 0.764302 | 1.14E-96 | 1.206204 | 2.00E-25 | 1.3536 | 5.25E-67 | 1.0591 | 0.002509734 |
| V74W A122D V196P | 1 | 0.711849 | 0.071639 | 3.38E-69 | 0.860557 | 2.07E-17 | 0.898366 | 0.00059 | 0.975302 | 0.329813 | 1.176471 | 3.44E-08 |
| N69E V74E R78H L89C V196P | 1 | 0.711843 | 0.054314 | 8.17E-39 | 0.682356 | 7.81E-58 | 1.444993 | 8.60E-17 | 1.474204 | 5.68E-09 | 0.864172 | 1.09E-05 |
| N69E N81S L89T V196P | 1 | 0.709529 | 0.048976 | 2.13E-60 | 0.879382 | 1.82E-15 | 0.697078 | 3.99E-44 | 0.578999 | 1.45E-67 | 1.471183 | 4.43E-24 |
| N69E A122H V196P | 1 | 0.709383 | 0.059154 | 1.63E-179 | 0.879286 | 2.90E-26 | 0.655565 | 6.43E-95 | 0.979052 | 0.028733 | 1.5131 | 5.40E-62 |
| N69E V74W R78H L89C A122D | 1 | 0.70897 | 0.0717 | 2.11E-77 | 0.682437 | 3.38E-104 | 0.763849 | 1.49E-30 | 0.795258 | 2.01E-40 | 0.756522 | 2.48E-20 |
| V74W N81S L89C A122H V196P | 1 | 0.708884 | 0.039655 | 5.59E-48 | 0.887469 | 1.46E-08 | 0.952804 | 0.549632 | 0.897774 | 0.002936 | 1.387674 | 4.62E-33 |
| N69E N81S A122D V196P | 1 | 0.708301 | 0.05661 | 9.39E-65 | 0.831959 | 6.58E-18 | 0.82483 | 6.21E-09 | 0.606562 | 8.72E-25 | 1.158247 | 0.024229146 |
| V74W R78H N81S | 1 | 0.708197 | 0.084506 | 4.85E-233 | 0.733322 | 3.95E-167 | 1.060031 | 0.00658 | 0.839817 | 1.59E-42 | 1.045275 | 9.91E-08 |
| Y63L L89G V196P | 1 | 0.707818 | 0.081507 | 2.05E-55 | 0.703828 | 4.15E-79 | 0.888373 | 0.000406 | 0.932003 | 0.041744 | 0.927231 | 0.004949304 |
| N69E L89A | 1 | 0.707324 | 0.086143 | 0 | 0.828805 | 5.53E-97 | 0.756931 | 2.90E-255 | 0.804343 | 2.56E-133 | 1.325486 | 1.06E-106 |

| | | | | | | | | | | | | |
|---|---|---|---|---|---|---|---|---|---|---|---|---|
| Y63L N69E V74E R78H N81S L89C | 1 | 0.707316 | 0.075744 | 2.91E-59 | 0.670923 | 1.83E-57 | 1.087671 | 0.022127 | 0.920099 | 0.039 | 0.781439 | 2.33E-08 |
| Y63L N69E N81S A122D | 1 | 0.70714 | 0.065897 | 1.78E-112 | 0.708429 | 4.23E-102 | 1.197144 | 2.29E-19 | 1.024554 | 0.03912 | 0.835996 | 6.57E-15 |
| Y63L N69E N81S L89T A122D | 1 | 0.707009 | 0.073469 | 1.33E-34 | 0.89307 | 7.64E-07 | 0.602656 | 2.46E-19 | 0.619761 | 9.08E-17 | 1.13742 | 0.003892781 |
| N69E L89G | 1 | 0.706715 | 0.071472 | 3.38E-234 | 0.870324 | 3.98E-46 | 0.625445 | 2.09E-307 | 0.694027 | 1.81E-161 | 1.340645 | 8.07E-59 |
| N69E V74E N81S A122D | 1 | 0.706305 | 0.063096 | 1.18E-41 | 0.610869 | 3.52E-74 | 1.45136 | 4.36E-16 | 1.367947 | 1.89E-11 | 0.643643 | 2.22E-12 |
| N69E N81S L89C A122D V196P | 1 | 0.706103 | 0.048086 | 1.19E-221 | 0.775789 | 2.00E-88 | 0.810512 | 1.04E-21 | 0.94475 | 0.000121 | 1.266557 | 8.15E-51 |
| R78H N81S L89C A122H V196P | 1 | 0.705797 | 0.056228 | 3.75E-300 | 0.647726 | 6.36E-277 | 1.348221 | 2.65E-67 | 1.271791 | 8.43E-37 | 0.675802 | 1.72E-147 |
| Y63L N69E N81S L89G A122D | 1 | 0.70307 | 0.07529 | 4.01E-207 | 0.765203 | 5.28E-109 | 0.657229 | 1.09E-116 | 0.833427 | 4.02E-16 | 1.1623 | 8.72E-13 |
| R78H L89A A122D | 1 | 0.701691 | 0.071396 | 0 | 0.711806 | 6.48E-289 | 0.875514 | 1.87E-27 | 0.951586 | 0.264242 | 0.927258 | 3.55E-06 |
| N69E V74E N81S L89T V196P | 1 | 0.70135 | 0.083977 | 7.21E-74 | 0.929671 | 0.000255 | 0.588434 | 2.60E-34 | 0.646635 | 3.50E-33 | 1.899645 | 2.50E-70 |
| V74E R78H A122D | 1 | 0.701332 | 0.058483 | 1.93E-21 | 0.541241 | 5.84E-75 | 1.113498 | 0.001536 | 1.246055 | 0.001337 | 0.515792 | 3.76E-17 |
| Y63L N81S L89A A122D | 1 | 0.700957 | 0.045962 | 3.73E-75 | 0.716896 | 1.77E-50 | 0.70078 | 2.08E-24 | 0.62102 | 2.09E-28 | 0.940117 | 0.000736864 |
| N81S L89T A122D | 1 | 0.700538 | 0.072576 | 0 | 0.805373 | 4.60E-221 | 0.863843 | 2.92E-43 | 0.719887 | 1.11E-183 | 1.192949 | 6.41E-32 |
| Y63L N69E V74W N81S L89C A122H V196P | 1 | 0.699617 | 0.039137 | 1.53E-59 | 0.819481 | 1.48E-28 | 1.281575 | 1.08E-05 | 1.153855 | 0.096909 | 1.173087 | 1.31E-10 |
| Y63L L89T | 1 | 0.699308 | 0.052709 | 4.14E-212 | 0.753083 | 3.98E-133 | 0.718425 | 6.60E-78 | 0.542291 | 1.89E-223 | 1.350366 | 2.63E-67 |
| Y63L V74E N81S L89C A122D | 1 | 0.699013 | 0.047604 | 1.47E-126 | 0.749621 | 6.01E-73 | 1.0396 | 0.070758 | 1.377814 | 3.37E-29 | 0.972046 | 0.402423707 |
| N69E V74W R78H L89C V196P | 1 | 0.697994 | 0.053949 | 1.61E-68 | 0.625849 | 2.33E-72 | 0.70776 | 4.45E-14 | 1.45573 | 1.70E-18 | 0.864856 | 2.02E-09 |
| N81S A122D | 1 | 0.696766 | 0.06493 | 0 | 0.857236 | 2.37E-49 | 1.12831 | 6.66E-29 | 1.363163 | 6.55E-72 | 1.431891 | 4.32E-64 |
| V74E N81S L89A | 1 | 0.696344 | 0.049564 | 0 | 0.859729 | 5.78E-71 | 0.73948 | 5.74E-108 | 0.894473 | 1.56E-23 | 1.569639 | 1.22E-292 |
| Y63L N69E N81S A122D V196P | 1 | 0.695569 | 0.056595 | 1.04E-62 | 0.731879 | 7.67E-32 | 0.784882 | 3.73E-07 | 0.887914 | 0.000339 | 0.985313 | 0.6882076 |
| Y63L N69E R78H N81S L89C A122D V196P | 1 | 0.695005 | 0.050726 | [illegible]470595911 | 0.725269 | 2.29E-175 | 0.837718 | 4.58E-31 | 1.019632 | 0.918842 | 1.086472 | 3.19E-10 |
| Y63L N81S L89G A122D | 1 | 0.69448 | 0.055554 | 1.16E-111 | 0.703321 | 2.49E-79 | 0.827522 | 1.46E-11 | 0.923391 | 0.000113 | 0.880646 | 2.11E-11 |
| V74E R78H N81S L89C A122H V196P | 1 | 0.693802 | 0.043943 | 2.69E-146 | 0.751662 | 4.05E-86 | 0.852559 | 2.95E-09 | 0.998858 | 0.914671 | 1.103504 | 0.004839217 |
| N69E V74E R78H N81S | 1 | 0.690408 | 0.095688 | 1.25E-117 | 0.643379 | 3.88E-198 | 0.969697 | 0.137085 | 0.806303 | 2.91E-10 | 0.903543 | 0.067362235 |
| Y63L N69E V74E L89C | 1 | 0.690179 | 0.070611 | 0 | 0.763784 | 2.87E-138 | 1.291245 | 7.57E-47 | 1.185702 | 1.31E-33 | 1.244177 | 2.40E-35 |
| N69E V74E L89G A122D V196P | 1 | 0.689278 | 0.062881 | 1.59E-25 | 0.617681 | 3.55E-40 | 0.701268 | 5.47E-10 | 0.752124 | 7.45E-06 | 0.739298 | 2.06E-10 |
| Y63L N69E L89A A122H V196P | 1 | 0.689103 | 0.052578 | 6.27E-15 | 0.80616 | 1.66E-09 | 0.421392 | 5.07E-17 | 0.564392 | 2.03E-12 | 1.194996 | 0.458611645 |
| V74E N81S L89T | 1 | 0.686594 | 0.073088 | 4.06E-282 | 0.716942 | 2.73E-196 | 0.756866 | 2.31E-68 | 0.671942 | 8.00E-149 | 0.921757 | 0.000146878 |
| N69E V74W A122D | 1 | 0.68626 | 0.057226 | 1.79E-92 | 0.798831 | 3.71E-37 | 0.714205 | 6.39E-28 | 0.915857 | 0.000984 | 1.014101 | 0.634069342 |
| Y63L V74W L89C A122D V196P | 1 | 0.686103 | 0.044788 | 6.61E-14 | 0.69131 | 1.35E-17 | 0.731172 | 0.005774 | 0.676052 | 0.00012 | 0.705538 | 2.04E-07 |
| V74W R78H N81S A122H | 1 | 0.685764 | 0.070929 | 2.54E-273 | 0.79289 | 5.45E-123 | 1.171502 | 5.13E-27 | 0.915845 | 7.74E-08 | 1.141942 | 1.50E-15 |
| Y63L V74E R78H | 1 | 0.684244 | 0.081929 | 2.92E-20 | 0.622227 | 2.28E-29 | 1.659617 | 4.65E-07 | 0.728024 | 0.002726 | 0.702913 | 7.86E-07 |
| N69E V74W R78H N81S L89T A122D V196P | 1 | 0.684208 | 0.046223 | 6.73E-14 | 0.826301 | 4.31E-09 | 0.631522 | 1.98E-07 | 0.847934 | 0.060168 | 1.289029 | 0.087095534 |
| Y63L V74W N81S L89C A122H | 1 | 0.6841 | 0.046588 | 1.30E-154 | 0.672273 | 9.77E-125 | 1.052406 | 0.018381 | 1.107846 | 0.000606 | 0.840472 | 8.60E-21 |
| N69E V74E A122D V196P | 1 | 0.683658 | 0.046698 | 6.36E-44 | 0.775315 | 1.11E-21 | 0.840803 | 0.146514 | 0.958096 | 0.035288 | 1.061199 | 0.0373516 |
| V74W R78H A122D | 1 | 0.681589 | 0.058773 | 1.53E-235 | 0.662766 | 1.51E-230 | 1.139386 | 9.11E-17 | 0.905765 | 2.68E-12 | 0.796309 | 1.80E-35 |
| V74E R78H N81S L89C A122D | 1 | 0.681327 | 0.046399 | 3.10E-186 | 0.748777 | 9.23E-94 | 0.988648 | 0.07853 | 1.372306 | 3.05E-51 | 1.025308 | 3.41E-05 |
| N69E L89C A122H | 1 | 0.680828 | 0.069382 | 8.86E-118 | 0.706263 | 1.31E-113 | 1.143167 | 4.17E-08 | 1.194187 | 1.28E-05 | 1.041997 | 0.413144 |
| N69E L89G A122H V196P | 1 | 0.68062 | 0.061054 | 1.84E-32 | 0.882836 | 3.66E-05 | 0.568424 | 1.17E-13 | 0.858499 | 0.016053 | 1.544726 | 4.75E-11 |
| R78H L89T A122D | 1 | 0.680568 | 0.09391 | 9.48E-191 | 0.778672 | 5.18E-102 | 0.756359 | 1.88E-32 | 0.742806 | 2.43E-38 | 1.36612 | 1.33E-48 |
| Y63L V74W N81S L89C V196P | 1 | 0.679161 | 0.068189 | 3.63E-90 | 0.826825 | 3.00E-34 | 0.998815 | 0.440682 | 1.37773 | 1.08E-08 | 1.199299 | 2.00E-07 |
| Y63L V74E L89C A122D | 1 | 0.677543 | 0.053429 | 7.06E-65 | 0.640739 | 1.96E-63 | 0.883442 | 0.005252 | 1.877272 | 3.40E-57 | 0.671568 | 7.96E-31 |
| V74W R78H L89G A122H | 1 | 0.677158 | 0.055487 | 3.94E-35 | 0.904828 | 5.67E-05 | 0.515948 | 0.0003 | 0.39559 | 1.12E-35 | 1.384313 | 5.13E-15 |
| N69E V74W L89T V196P | 1 | 0.676006 | 0.085022 | 9.20E-19 | 0.658329 | 2.69E-25 | 0.611097 | 0.000581 | 0.707449 | 7.03E-05 | 0.927991 | 0.577974706 |
| Y63L N69E V74E L89C V196P | 1 | 0.675069 | 0.079709 | 2.90E-170 | 0.658572 | 1.39E-135 | 0.979376 | 0.03832 | 0.879985 | 3.86E-08 | 0.876103 | 1.97E-15 |
| N69E V74E L89A A122H V196P | 1 | 0.67416 | 0.077632 | 9.21E-52 | 0.733717 | 1.15E-50 | 0.80316 | 2.70E-07 | 0.7975 | 2.82E-13 | 1.326965 | 5.57E-11 |
| Y63L N69E V74W N81S L89C V196P | 1 | 0.673863 | 0.045891 | 6.06E-169 | 0.738123 | 1.46E-86 | 0.801314 | 5.64E-14 | 1.247945 | 2.84E-11 | 1.073705 | 0.001694049 |
| N69E V74E N81S L89C A122H V196P | 1 | 0.673761 | 0.042674 | 9.60E-28 | 0.575348 | 1.36E-57 | 1.010415 | 0.650169 | 1.062455 | 0.95885 | 0.594498 | 5.42E-30 |

| | | | | | | | | | | | | |
|---|---|---|---|---|---|---|---|---|---|---|---|---|
| V74E L89C A122H | 1 | 0.672795 | 0.057487 | 1.31E-76 | 0.714813 | 1.64E-50 | 1.012614 | 0.210986 | 0.816103 | 2.17E-11 | 1.001934 | 0.651432026 |
| N69E N81S L89C V196P | 1 | 0.672417 | 0.045792 | 5.29E-139 | 0.710646 | 7.90E-94 | 0.993976 | 0.904016 | 1.194894 | 8.08E-11 | 0.973266 | 0.018232486 |
| Y63L N81S A122H V196P | 1 | 0.67122 | 0.088178 | 2.15E-55 | 0.600752 | 1.69E-90 | 1.128437 | 0.001023 | 1.150279 | 1.51E-07 | 0.887743 | 0.447878369 |
| N69E L89A A122D | 1 | 0.670723 | 0.068245 | 0 | 0.69517 | 1.01E-223 | 0.776987 | 8.90E-65 | 0.609376 | 2.36E-203 | 0.939288 | 1.05E-05 |
| N69E R78H L89C V196P | 1 | 0.669452 | 0.080889 | 2.33E-235 | 0.623701 | 9.65E-188 | 0.796987 | 6.57E-12 | 1.099488 | 4.20E-06 | 0.97297 | 0.108438552 |
| N69E R78H N81S L89C A122H V196P | 1 | 0.668503 | 0.053257 | 0 | 0.614986 | 0 | 1.304475 | 4.83E-49 | 1.171388 | 3.41E-15 | 0.647246 | 2.15E-173 |
| N69E V74E N81S L89C A122D | 1 | 0.667653 | 0.0636 | 1.53E-60 | 0.804363 | 1.69E-18 | 0.619671 | 2.07E-14 | 0.602848 | 2.46E-19 | 1.197317 | 0.011405479 |
| N69E R78H L89A A122D | 1 | 0.6676 | 0.065094 | 2.12E-52 | 0.750962 | 2.52E-29 | 0.634719 | 1.10E-14 | 1.153451 | 0.000241 | 0.907926 | 0.02504565 |
| R78H N81S | 1 | 0.667201 | 0.066577 | 3.33E-84 | 1.027859 | 0.060405 | 0.882994 | 3.90E-09 | 0.843203 | 0.008345 | 1.457011 | 3.69E-30 |
| Y63L V74W R78H N81S L89C A122H | 1 | 0.666279 | 0.067899 | 2.56E-87 | 0.701145 | 1.95E-71 | 1.053271 | 0.762478 | 0.943804 | 0.712827 | 0.947334 | 0.220744696 |
| N81S L89C | 1 | 0.66617 | 0.045367 | 0 | 0.791057 | 1.03E-98 | 1.043365 | 0.127713 | 0.96659 | 0.014192 | 1.434746 | 4.68E-162 |
| Y63L V74E R78H V196P | 1 | 0.665834 | 0.051391 | 9.08E-28 | 0.593696 | 1.24E-35 | 1.145109 | 0.000465 | 1.093349 | 0.046734 | 0.939798 | 0.186853915 |
| V74W R78H N81S A122D | 1 | 0.665827 | 0.057345 | 8.83E-98 | 0.640973 | 4.18E-77 | 1.108004 | 0.000813 | 1.107428 | 0.06588 | 0.964111 | 0.018573309 |
| Y63L V74W N81S L89T | 1 | 0.665456 | 0.060365 | 2.59E-43 | 0.521889 | 2.86E-73 | 1.033701 | 0.63279 | 1.115004 | 0.009733 | 0.631102 | 3.75E-24 |
| V74E L89C A122D V196P | 1 | 0.664033 | 0.044229 | 8.05E-74 | 0.724023 | 2.03E-48 | 1.127624 | 0.107241 | 1.228085 | 9.46E-09 | 1.113136 | 0.011715063 |
| N69E V74W L89C A122D V196P | 1 | 0.663209 | 0.076371 | 2.97E-261 | 0.790177 | 1.65E-147 | 1.030714 | 0.018257 | 1.014448 | 0.702494 | 1.427974 | 3.99E-66 |
| Y63L N69E V74E N81S L89G | 1 | 0.662211 | 0.041942 | 6.76E-27 | 0.645526 | 4.89E-29 | 0.911242 | 0.076756 | 1.264021 | 0.002022 | 0.753636 | 1.06E-06 |
| R78H L89G A122D | 1 | 0.661732 | 0.060027 | 2.43E-205 | 0.661093 | 2.15E-170 | 0.948435 | 0.00778 | 0.977172 | 0.2853 | 0.828774 | 1.78E-16 |
| Y63L N69E V74W L89G A122D V196P | 1 | 0.661043 | 0.059297 | 6.77E-18 | 0.851213 | 3.97E-05 | 0.499614 | 1.14E-17 | 0.813732 | 0.002494 | 1.173861 | 0.023278713 |
| Y63L V74W L89G A122H | 1 | 0.660666 | 0.061566 | 9.28E-15 | 0.725494 | 2.04E-17 | 1.16185 | 0.006803 | 0.717567 | 4.18E-05 | 0.778464 | 0.001580659 |
| Y63L V74E R78H N81S L89C | 1 | 0.660456 | 0.049261 | 2.35E-169 | 0.625897 | 8.87E-141 | 0.906007 | 0.132411 | 1.016981 | 0.460405 | 0.761317 | 3.29E-16 |
| N81S L89A A122D | 1 | 0.659918 | 0.074872 | 1.26E-210 | 0.785932 | 7.97E-93 | 0.821335 | 1.92E-18 | 0.747737 | 1.05E-28 | 1.337665 | 1.60E-40 |
| Y63L L89T A122H | 1 | 0.659086 | 0.049677 | 2.64E-200 | 0.673232 | 5.20E-181 | 0.835995 | 2.64E-07 | 0.625267 | 1.12E-98 | 1.187096 | 1.02E-20 |
| V74W A122H V196P | 1 | 0.658019 | 0.050788 | 1.65E-170 | 0.716019 | 1.23E-121 | 1.010618 | 0.480898 | 1.334561 | 3.01E-32 | 1.101358 | 0.000680541 |
| R78H N81S L89G A122D V196P | 1 | 0.657831 | 0.068106 | 1.40E-286 | 0.720238 | 9.33E-169 | 0.776792 | 1.25E-30 | 0.911069 | 6.27E-05 | 1.037692 | 0.882534916 |
| N69E V74E R78H A122H V196P | 1 | 0.657541 | 0.050751 | 3.60E-185 | 0.775178 | 1.58E-72 | 1.218682 | 2.18E-12 | 1.47546 | 1.72E-41 | 1.365438 | 1.57E-33 |
| N69E V74W N81S L89C A122H V196P | 1 | 0.657376 | 0.036774 | 5.11E-135 | 0.909053 | 5.88E-13 | 1.026803 | 0.403898 | 0.856613 | 0.000118 | 2.077587 | 2.51E-149 |
| N81S L89G V196P | 1 | 0.65644 | 0.07029 | 4.64E-196 | 0.729068 | 6.01E-105 | 0.714413 | 1.09E-32 | 0.858626 | 2.12E-17 | 1.060148 | 0.461667133 |
| Y63L N69E V74W R78H A122H V196P | 1 | 0.656333 | 0.05473 | 5.77E-78 | 0.657179 | 2.10E-59 | 0.98885 | 0.873967 | 1.152294 | 4.20E-05 | 0.93335 | 0.006735059 |
| Y63L L89G A122H V196P | 1 | 0.655918 | 0.058838 | 1.93E-79 | 0.681358 | 7.25E-49 | 1.103463 | 0.14648 | 0.857019 | 0.000268 | 0.922859 | 0.039100303 |
| R78H L89A A122D V196P | 1 | 0.655892 | 0.060184 | 5.16E-267 | 0.62882 | 5.47E-186 | 0.823708 | 5.91E-31 | 1.36704 | 1.02E-29 | 1.004534 | 0.06261094 |
| V74E L89G V196P | 1 | 0.655009 | 0.043628 | 1.51E-30 | 0.706871 | 1.03E-17 | 0.796774 | 6.73E-05 | 0.912711 | 0.177758 | 1.085477 | 0.085437223 |
| N69E V74W R78H N81S L89T A122H | 1 | 0.654783 | 0.052333 | 7.53E-159 | 0.777095 | 5.48E-38 | 1.142077 | 4.68E-07 | 0.974687 | 0.314693 | 1.249277 | 8.41E-25 |
| V74W L89G A122H V196P | 1 | 0.654651 | 0.078385 | 9.46E-64 | 0.796659 | 5.28E-28 | 0.819848 | 1.50E-06 | 0.600769 | 2.74E-24 | 1.176522 | 5.88E-07 |
| Y63L N69E V74W A122D V196P | 1 | 0.653622 | 0.050449 | 1.07E-40 | 0.641022 | 7.45E-51 | 1.180864 | 0.005847 | 0.857217 | 0.13521 | 0.927166 | 0.16294171 |
| N69E V74W R78H N81S L89C | 1 | 0.653504 | 0.066598 | 9.37E-68 | 0.737096 | 3.98E-28 | 0.773945 | 0.000156 | 0.817431 | 2.19E-06 | 0.97941 | 0.903395042 |
| Y63L N69E L89C V196P | 1 | 0.653425 | 0.078953 | 4.03E-241 | 0.643022 | 2.73E-195 | 0.953162 | 0.049855 | 0.747497 | 3.07E-65 | 0.892794 | 1.25E-13 |
| L89G A122H | 1 | 0.653327 | 0.058364 | 1.71E-157 | 0.795073 | 1.69E-41 | 0.768097 | 7.81E-08 | 0.744908 | 1.14E-22 | 1.391634 | 1.13E-17 |
| Y63L V74W N81S V196P | 1 | 0.652853 | 0.073936 | 4.85E-150 | 0.661355 | 1.77E-193 | 1.095015 | 0.037846 | 1.161019 | 8.95E-10 | 0.945214 | 0.003870871 |
| N69E V74E R78H L89C A122H V196P | 1 | 0.652642 | 0.040049 | 2.89E-18 | 0.650413 | 4.95E-22 | 0.805811 | 0.183564 | 1.133134 | 0.118006 | 0.929017 | 0.453725861 |
| N69E V74E L89C A122D | 1 | 0.652564 | 0.060267 | 1.02E-199 | 0.856831 | 2.74E-37 | 1.060861 | 0.000865 | 0.81364 | 8.09E-33 | 1.45779 | 9.41E-67 |
| V74W L89G A122D | 1 | 0.652188 | 0.05344 | 6.22E-74 | 0.772151 | 1.79E-19 | 0.811898 | 0.247016 | 0.945436 | 0.040038 | 1.27064 | 5.64E-11 |
| Y63L R78H A122H | 1 | 0.651648 | 0.059448 | 5.20E-165 | 0.611221 | 2.19E-184 | 1.208389 | 7.79E-23 | 1.200493 | 5.56E-20 | 0.829578 | 1.16E-15 |
| Y63L N69E N81S L89G A122H | 1 | 0.65048 | 0.036388 | 4.54E-55 | 0.814098 | 4.86E-19 | 1.068716 | 0.723771 | 0.830473 | 0.015855 | 1.402426 | 4.24E-32 |
| N69E V74E R78H N81S L89C A122H V196P | 1 | 0.649646 | 0.073706 | 2.66E-59 | 0.760371 | 1.94E-43 | 0.864518 | 0.001206 | 0.892969 | 0.01069 | 1.187461 | 5.07E-05 |
| V74W R78H N81S L89G A122H | 1 | 0.648306 | 0.041061 | 3.26E-24 | 0.628434 | 2.62E-27 | 0.468523 | 1.80E-10 | 1.05688 | 0.889527 | 0.867915 | 0.000504394 |
| N69E L89G V196P | 1 | 0.647848 | 0.043151 | 3.29E-66 | 0.703373 | 1.00E-42 | 1.057557 | 0.43202 | 1.402395 | 6.26E-17 | 1.114973 | 0.000521574 |
| Y63L R78H N81S L89T A122D V196P | 1 | 0.647115 | 0.069134 | 7.37E-231 | 0.728916 | 1.06E-149 | 1.001844 | 0.731388 | 1.031278 | 0.203976 | 1.199323 | 9.26E-15 |

| | | | | | | | | | | | | |
|---|---|---|---|---|---|---|---|---|---|---|---|---|
| Y63L N69E V74E R78H L89A A122H V196P | 1 | 0.645883 | 0.068077 | 3.71E-67 | 0.892864 | 5.07E-06 | 0.698168 | 1.28E-31 | 0.71001 | 1.52E-15 | 1.228016 | 1.50E-05 |
| Y63L N69E V74E N81S L89C | 1 | 0.644136 | 0.091004 | 0 | 0.565734 | 0 | 0.898942 | 8.59E-28 | 1.074941 | 1.21E-09 | 0.693798 | 2.25E-87 |
| V74W R78H N81S L89T A122H | 1 | 0.644112 | 0.044461 | 1.40E-147 | 0.729037 | 4.88E-84 | 0.976625 | 0.1921 | 1.314405 | 8.23E-15 | 1.05805 | 0.045982836 |
| Y63L N69E V74E N81S L89T A122H V196P | 1 | 0.643988 | 0.067315 | 5.66E-19 | 0.556031 | 1.76E-53 | 0.949517 | 0.910799 | 1.163917 | 0.500823 | 0.659116 | 5.98E-07 |
| V74E R78H L89C A122H | 1 | 0.643175 | 0.063032 | 8.92E-106 | 0.634479 | 3.24E-92 | 0.887898 | 1.31E-08 | 0.697944 | 1.64E-53 | 0.997119 | 0.075284677 |
| N69E L89T V196P | 1 | 0.642222 | 0.047531 | 0 | 0.762381 | 1.29E-209 | 0.83315 | 1.33E-58 | 0.733145 | 1.03E-131 | 1.663707 | 3.26E-264 |
| V74E N81S L89G | 1 | 0.642091 | 0.051153 | 5.02E-159 | 0.68534 | 1.26E-92 | 0.702136 | 4.03E-27 | 0.590111 | 6.93E-65 | 0.87498 | 1.99E-05 |
| V74E L89A V196P | 1 | 0.642 | 0.051356 | 0 | 0.665454 | 1.26E-212 | 0.778396 | 1.32E-29 | 1.214264 | 5.42E-20 | 0.959243 | 0.002853845 |
| Y63L N69E R78H V196P | 1 | 0.641223 | 0.064531 | 4.61E-60 | 0.806744 | 1.38E-25 | 0.971021 | 0.980806 | 0.872228 | 0.000104 | 0.989168 | 0.587780515 |
| N69E R78H L89A A122D V196P | 1 | 0.640694 | 0.065549 | 0 | 0.818704 | 1.88E-66 | 0.92223 | 1.19E-08 | 1.136616 | 5.70E-22 | 1.61695 | 8.91E-130 |
| N69E R78H N81S L89T A122D | 1 | 0.640543 | 0.058105 | 0 | 0.728755 | 2.38E-193 | 0.916001 | 6.55E-24 | 1.301939 | 4.94E-76 | 0.915709 | 8.31E-08 |
| V74W L89T | 1 | 0.640469 | 0.072665 | 1.23E-86 | 0.874009 | 4.06E-14 | 0.572617 | 6.32E-43 | 0.597078 | 3.31E-25 | 1.766876 | 8.20E-50 |
| N69E V74E N81S | 1 | 0.640373 | 0.068414 | 1.82E-137 | 0.733704 | 6.52E-74 | 1.072217 | 0.002793 | 1.034245 | 0.182254 | 0.971158 | 0.068509102 |
| Y63L V74W R78H N81S V196P | 1 | 0.637848 | 0.051899 | 3.46E-84 | 0.707227 | 1.52E-51 | 1.124572 | 0.000162 | 0.876481 | 0.012975 | 1.037904 | 0.456472943 |
| N69E N81S L89T A122D | 1 | 0.637551 | 0.063619 | 7.96E-52 | 0.917455 | 5.43E-05 | 0.696093 | 6.14E-15 | 0.597056 | 4.18E-10 | 1.234537 | 1.87E-07 |
| Y63L V74W R78H N81S L89C A122D V196P | 1 | 0.637202 | 0.050763 | 2.19E-126 | 0.562257 | 1.65E-136 | 1.077158 | 0.4056 | 0.782812 | 3.69E-09 | 0.594142 | 1.30E-52 |
| N69E V74E R78H L89C | 1 | 0.637142 | 0.065185 | 0 | 0.730989 | 4.58E-217 | 1.116865 | 2.01E-09 | 0.972168 | 0.706498 | 1.205566 | 6.57E-16 |
| Y63L N69E V74E | 1 | 0.637112 | 0.058856 | 1.90E-95 | 0.643182 | 8.72E-85 | 1.51932 | 1.15E-51 | 1.508108 | 9.40E-25 | 0.713179 | 0.001349194 |
| Y63L R78H N81S L89C | 1 | 0.637037 | 0.06411 | 6.46E-123 | 0.87519 | 1.69E-14 | 0.734271 | 4.54E-19 | 1.108824 | 0.0005 | 1.308198 | 1.04E-24 |
| Y63L R78H N81S L89G A122D V196P | 1 | 0.63662 | 0.06591 | 7.82E-42 | 0.600014 | 4.44E-41 | 0.702028 | 0.003196 | 0.79294 | 3.46E-05 | 0.781435 | 1.54E-06 |
| N69E R78H L89G A122D | 1 | 0.63661 | 0.052164 | 4.30E-45 | 0.681227 | 3.83E-26 | 0.530035 | 7.10E-31 | 1.664133 | 1.47E-08 | 0.934919 | 0.204018389 |
| Y63L N69E V74E N81S | 1 | 0.635683 | 0.066803 | 3.11E-105 | 0.579928 | 1.82E-250 | 1.652779 | 4.87E-82 | 1.451751 | 3.49E-28 | 0.655674 | 3.73E-07 |
| Y63L N81S A122H | 1 | 0.634999 | 0.065678 | 1.38E-69 | 0.665801 | 2.04E-61 | 1.300908 | 9.89E-18 | 0.965522 | 0.26028 | 0.743364 | 7.67E-15 |
| Y63L N69E N81S L89A A122D V196P | 1 | 0.632683 | 0.041485 | 1.41E-121 | 0.688554 | 6.84E-65 | 0.767805 | 2.55E-17 | 0.553177 | 1.81E-39 | 1.261602 | 4.08E-13 |
| Y63L R78H N81S L89T V196P | 1 | 0.631362 | 0.043581 | 2.76E-28 | 0.680335 | 6.87E-24 | 0.751786 | 2.79E-08 | 0.889755 | 0.515245 | 1.054475 | 0.932345827 |
| Y63L V74W R78H N81S | 1 | 0.631277 | 0.071493 | 7.08E-39 | 0.566224 | 8.80E-57 | 1.186022 | 8.43E-05 | 1.482283 | 5.53E-10 | 0.888003 | 0.078693075 |
| N69E V74W R78H N81S A122H | 1 | 0.630713 | 0.041172 | 2.54E-31 | 0.647549 | 1.28E-36 | 1.211059 | 0.000235 | 1.383804 | 6.97E-06 | 0.934663 | 0.272649884 |
| N69E N81S A122D | 1 | 0.63071 | 0.058774 | 7.28E-164 | 0.755653 | 3.54E-55 | 1.127205 | 0.000288 | 1.014703 | 0.711552 | 1.110961 | 0.007467025 |
| Y63L V74E N81S L89C | 1 | 0.630424 | 0.046013 | 0 | 0.66504 | 0 | 0.982516 | 0.27033 | 0.985052 | 0.977882 | 1.02495 | 3.79E-05 |
| Y63L R78H N81S A122D | 1 | 0.630161 | 0.050409 | 5.12E-205 | 0.60418 | 9.04E-213 | 1.442872 | 7.54E-69 | 1.392269 | 1.86E-37 | 0.758968 | 2.53E-32 |
| Y63L N69E V74W N81S V196P | 1 | 0.630111 | 0.054269 | 6.93E-97 | 0.55487 | 1.24E-126 | 1.701532 | 6.27E-58 | 1.300556 | 7.79E-16 | 0.78091 | 4.97E-14 |
| V74E R78H A122H | 1 | 0.629293 | 0.056449 | 4.94E-122 | 0.75999 | 2.96E-34 | 1.088794 | 0.000615 | 1.232816 | 4.65E-16 | 1.31427 | 3.28E-14 |
| N81S A122D V196P | 1 | 0.628984 | 0.050271 | 3.16E-38 | 0.657596 | 3.75E-36 | 0.905136 | 0.093656 | 0.503985 | 2.21E-12 | 0.841123 | 0.000164951 |
| V74W L89C A122D V196P | 1 | 0.628286 | 0.041848 | 4.92E-80 | 0.706137 | 7.38E-61 | 1.052869 | 0.306924 | 1.06191 | 0.00644 | 1.091482 | 0.040774391 |
| Y63L N69E L89A A122H | 1 | 0.62807 | 0.06933 | 0 | 0.704396 | 4.87E-210 | 0.864425 | 3.92E-28 | 1.032797 | 2.61E-05 | 1.072494 | 0.422598298 |
| Y63L V74W R78H L89C A122H V196P | 1 | 0.628057 | 0.072323 | 2.33E-65 | 0.672512 | 8.24E-69 | 0.925182 | 0.561515 | 1.757887 | 1.44E-26 | 0.911139 | 0.001800554 |
| Y63L N69E V74E N81S L89T | 1 | 0.627992 | 0.071121 | 2.66E-282 | 0.765897 | 2.64E-119 | 0.765806 | 2.91E-30 | 1.007057 | 0.00285 | 1.307757 | 8.50E-25 |
| N69E V74E R78H N81S L89C | 1 | 0.627801 | 0.067229 | 2.08E-274 | 0.718809 | 1.25E-130 | 0.898202 | 0.004841 | 1.05302 | 0.313927 | 1.037068 | 0.561382227 |
| N81S L89C A122D V196P | 1 | 0.627098 | 0.049958 | 2.45E-135 | 0.573778 | 9.60E-114 | 1.105219 | 3.06E-05 | 1.096106 | 0.650225 | 0.706828 | 3.76E-25 |
| Y63L L89G | 1 | 0.626727 | 0.073642 | 2.61E-45 | 0.692574 | 1.82E-36 | 0.833812 | 0.340745 | 0.402179 | 1.63E-34 | 1.209745 | 0.009058429 |
| Y63L N81S L89A A122H | 1 | 0.626632 | 0.043254 | 2.27E-70 | 0.650885 | 2.56E-73 | 0.980121 | 0.414733 | 1.605659 | 2.34E-21 | 0.909757 | 3.36E-06 |
| V74E R78H L89C A122D | 1 | 0.625564 | 0.061306 | 1.04E-150 | 0.566911 | 2.41E-264 | 0.881371 | 8.78E-12 | 0.990645 | 0.50657 | 0.652333 | 6.54E-56 |
| N69E V74W R78H A122D | 1 | 0.624937 | 0.074828 | 6.65E-118 | 0.706743 | 6.93E-77 | 1.494559 | 8.55E-24 | 0.965306 | 0.331757 | 1.159685 | 1.14E-06 |
| V74W R78H N81S L89T A122H V196P | 1 | 0.624105 | 0.092253 | 2.76E-189 | 0.811977 | 4.65E-125 | 0.934884 | 0.228586 | 0.994471 | 0.002097 | 1.612725 | 1.35E-99 |
| Y63L N69E V74W L89C | 1 | 0.622592 | 0.060706 | 3.59E-85 | 0.649781 | 5.45E-72 | 1.186523 | 0.000309 | 1.616316 | 3.17E-23 | 0.803873 | 6.55E-13 |
| Y63L N69E R78H L89C V196P | 1 | 0.622278 | 0.073476 | 6.01E-177 | 0.579337 | 5.79E-187 | 0.918842 | 0.001536 | 0.744093 | 2.24E-26 | 0.819067 | 7.89E-17 |
| N69E V74E R78H N81S A122H V196P | 1 | 0.621858 | 0.064425 | 1.76E-135 | 0.600098 | 1.12E-140 | 0.685444 | 4.99E-54 | 0.892555 | 4.84E-06 | 0.757773 | 2.28E-26 |
| N69E V74E L89G A122H | 1 | 0.621202 | 0.053501 | 8.38E-32 | 0.648484 | 7.85E-36 | 0.869194 | 0.008407 | 0.771025 | 1.31E-06 | 0.937395 | 0.004543142 |

| | | | | | | | | | | | | |
|---|---|---|---|---|---|---|---|---|---|---|---|---|
| N69E R78H N81S A122H V196P | 1 | 0.620873 | 0.07502 | 0 | 0.686227 | 1.24E-270 | 1.045223 | 0.006236 | 1.042096 | 4.20E-05 | 0.998612 | 0.498512628 |
| Y63L V74E A122D V196P | 1 | 0.620192 | 0.053478 | 2.72E-116 | 0.579722 | 3.18E-153 | 1.172191 | 1.83E-07 | 1.155926 | 7.49E-08 | 0.716183 | 5.56E-30 |
| Y63L V74W L89C A122D | 1 | 0.620068 | 0.057266 | 1.82E-54 | 0.729512 | 1.40E-36 | 0.704533 | 1.82E-22 | 0.71397 | 3.49E-12 | 1.218511 | 3.57E-05 |
| Y63L R78H | 1 | 0.618823 | 0.065031 | 4.30E-262 | 0.612934 | 1.15E-236 | 0.849914 | 1.01E-12 | 0.83958 | 1.11E-07 | 0.741471 | 2.37E-35 |
| Y63L L89T A122H V196P | 1 | 0.617954 | 0.070111 | 6.13E-58 | 0.785007 | 8.56E-29 | 0.734958 | 9.38E-12 | 0.589593 | 4.51E-21 | 1.558999 | 1.99E-17 |
| Y63L N69E N81S L89T A122H V196P | 1 | 0.617665 | 0.04032 | 6.88E-21 | 0.64158 | 6.61E-28 | 0.831174 | 0.004872 | 1.179488 | 0.000669 | 0.718923 | 4.33E-05 |
| V74E A122H V196P | 1 | 0.617428 | 0.05324 | 6.84E-168 | 0.644583 | 9.89E-114 | 0.991793 | 0.384966 | 1.389872 | 2.03E-23 | 0.897858 | 3.68E-06 |
| V74W N81S | 1 | 0.617182 | 0.069897 | 2.46E-233 | 0.700476 | 5.24E-176 | 1.082891 | 0.002949 | 0.990607 | 9.08E-15 | 1.11196 | 8.59E-05 |
| Y63L L89A A122H V196P | 1 | 0.616839 | 0.072833 | 0 | 0.606335 | 3.10E-303 | 0.738485 | 5.98E-71 | 0.784302 | 8.41E-31 | 0.98397 | 0.57625171 |
| N69E V74E L89G A122H V196P | 1 | 0.616795 | 0.056979 | 7.68E-102 | 0.676357 | 9.03E-73 | 0.757545 | 7.76E-13 | 0.976801 | 0.607032 | 1.003111 | 0.528383586 |
| N81S L89G A122D | 1 | 0.616615 | 0.066032 | 2.12E-117 | 0.687456 | 1.58E-68 | 0.718979 | 1.24E-30 | 0.417703 | 6.45E-112 | 1.250583 | 4.94E-06 |
| V74W R78H L89C | 1 | 0.61617 | 0.056906 | 1.39E-298 | 0.769548 | 1.04E-100 | 0.928261 | 2.60E-06 | 0.988297 | 0.912077 | 1.441506 | 6.56E-86 |
| N69E N81S L89C A122H | 1 | 0.615855 | 0.064719 | 9.05E-58 | 0.512366 | 9.41E-126 | 1.30553 | 0.11496 | 1.748729 | 5.88E-21 | 0.873539 | 7.30E-19 |
| Y63L V74W L89G A122H V196P | 1 | 0.615377 | 0.056139 | 2.99E-118 | 0.609478 | 1.03E-145 | 0.910264 | 0.004602 | 0.948285 | 0.006132 | 0.865896 | 9.07E-12 |
| V74W R78H N81S L89T A122D V196P | 1 | 0.612813 | 0.063384 | 3.69E-303 | 0.757442 | 1.25E-102 | 0.954412 | 0.842675 | 0.817289 | 3.04E-18 | 1.398266 | 4.07E-64 |
| Y63L V74E N81S L89C A122H V196P | 1 | 0.612798 | 0.038812 | 1.83E-31 | 0.571717 | 1.07E-32 | 1.038958 | 0.887754 | 0.83886 | 0.03499 | 1.019856 | 0.01786144 |
| N69E N81S A122H | 1 | 0.611625 | 0.048926 | 0 | 0.654687 | 8.87E-288 | 1.43919 | 2.28E-122 | 2.287409 | 525324304 | 1.118367 | 5.55E-10 |
| V74W A122D | 1 | 0.611623 | 0.051002 | 3.63E-154 | 0.740344 | 4.07E-62 | 0.848555 | 1.00E-05 | 0.763732 | 2.35E-21 | 1.238096 | 1.05E-08 |
| N69E V74W L89T A122H V196P | 1 | 0.611212 | 0.044923 | 3.50E-189 | 0.737946 | 3.04E-67 | 0.692001 | 1.89E-61 | 0.62965 | 8.40E-57 | 1.371407 | 6.36E-32 |
| Y63L V74W L89G A122D | 1 | 0.610061 | 0.052542 | 2.48E-47 | 0.68057 | 1.10E-33 | 0.84825 | 0.00939 | 0.670454 | 1.04E-08 | 1.089152 | 0.962098841 |
| R78H L89G | 1 | 0.609753 | 0.061666 | 0 | 0.701814 | 1.24E-270 | 0.823209 | 2.42E-46 | 1.561042 | 2.94E-141 | 1.006353 | 0.349473768 |
| N69E V74W A122H | 1 | 0.60799 | 0.059064 | 3.04E-268 | 0.84434 | 4.25E-44 | 0.766441 | 8.42E-21 | 0.780421 | 4.72E-28 | 1.784124 | 3.50E-52 |
| Y63L N69E L89A V196P | 1 | 0.607944 | 0.061858 | 129168249 | 0.654683 | 1.59E-250 | 0.767921 | 1.44E-72 | 0.633993 | 2.47E-88 | 0.848995 | 1.99E-13 |
| Y63L N81S L89T A122D | 1 | 0.607913 | 0.073454 | 0 | 0.713755 | 1.12E-238 | 0.776237 | 3.25E-77 | 0.910762 | 1.10E-07 | 1.177472 | 7.38E-25 |
| Y63L R78H N81S V196P | 1 | 0.607656 | 0.063144 | 1.09E-42 | 0.746491 | 2.04E-31 | 1.10297 | 0.58622 | 1.371957 | 1.45E-05 | 0.816397 | 0.000306851 |
| Y63L V74E R78H N81S | 1 | 0.607652 | 0.084219 | 2.60E-39 | 0.466474 | 4.93E-50 | 1.691953 | 0.027574 | 1.554823 | 3.83E-13 | 0.75236 | 0.006298466 |
| Y63L R78H N81S L89T A122D | 1 | 0.607403 | 0.071719 | 0 | 0.667237 | 2.79E-182 | 0.981482 | 0.19463 | 1.004606 | 0.468938 | 1.143931 | 4.61E-07 |
| Y63L N69E L89G | 1 | 0.606798 | 0.061367 | 4.31E-120 | 0.644696 | 7.11E-120 | 0.730507 | 1.62E-28 | 0.921265 | 0.038032 | 0.829586 | 7.88E-15 |
| N69E L89A A122H | 1 | 0.606142 | 0.072577 | 1.21E-71 | 0.729376 | 1.33E-51 | 0.93565 | 0.037701 | 0.772732 | 2.56E-06 | 1.54471 | 2.64E-24 |
| Y63L R78H V196P | 1 | 0.605781 | 0.061059 | 1.81E-118 | 0.657211 | 1.53E-89 | 0.810057 | 7.07E-12 | 0.579557 | 6.20E-45 | 0.840238 | 1.34E-07 |
| Y63L N69E V74W L89C A122H | 1 | 0.604174 | 0.055798 | 3.70E-110 | 0.74676 | 2.96E-61 | 0.758249 | 1.52E-29 | 0.697462 | 1.89E-24 | 1.514746 | 2.01E-32 |
| Y63L R78H N81S L89A A122D V196P | 1 | 0.604017 | 0.043721 | 2.70E-49 | 0.61829 | 2.54E-35 | 0.684247 | 7.43E-10 | 1.251569 | 0.073791 | 1.147814 | 0.184134063 |
| N69E V74E N81S L89T | 1 | 0.603521 | 0.064245 | 0 | 0.66165 | 7.47E-193 | 0.636481 | 1.25E-101 | 0.82493 | 1.11E-25 | 0.95919 | 0.029538036 |
| Y63L V74W N81S L89T A122H | 1 | 0.603512 | 0.041658 | 9.63E-34 | 0.676518 | 9.78E-25 | 0.778851 | 0.000177 | 1.191355 | 0.008315 | 0.94697 | 0.085574508 |
| Y63L N69E N81S L89A A122D | 1 | 0.603293 | 0.04273 | 3.18E-145 | 0.649813 | 2.90E-98 | 0.867241 | 0.000411 | 0.811442 | 1.12E-06 | 0.959129 | 0.006812152 |
| V74E N81S L89C A122H | 1 | 0.60302 | 0.044977 | 1.07E-296 | 0.672814 | 7.25E-151 | 1.044654 | 0.019936 | 1.167182 | 4.44E-08 | 1.339042 | 3.94E-26 |
| Y63L N69E N81S L89G A122D V196P | 1 | 0.602545 | 0.064519 | 8.67E-167 | 0.673782 | 1.71E-114 | 0.741314 | 3.77E-21 | 0.657057 | 2.94E-58 | 0.960651 | 0.011567842 |
| Y63L N69E V74E R78H N81S L89C A122H | 1 | 0.602086 | 0.044907 | 8.74E-137 | 0.649259 | 3.27E-82 | 0.905698 | 0.001256 | 1.104428 | 0.458945 | 1.151771 | 0.010948242 |
| Y63L V74W A122D V196P | 1 | 0.601669 | 0.083023 | 9.00E-56 | 0.657517 | 1.43E-49 | 0.670927 | 4.18E-07 | 0.774317 | 1.08E-05 | 1.212129 | 0.000758732 |
| V74W R78H N81S L89T A122D | 1 | 0.601244 | 0.062478 | 695875807 | 0.876355 | 3.41E-33 | 0.819921 | 1.32E-14 | 1.01842 | 0.658631 | 1.767827 | 5.08E-144 |
| Y63L V74E N81S L89G A122H | 1 | 0.600334 | 0.033113 | 9.36E-31 | 0.682503 | 1.53E-22 | 0.904905 | 0.025381 | 1.064101 | 0.227096 | 0.937079 | 0.529656897 |
| Y63L V74W R78H A122H V196P | 1 | 0.599404 | 0.051686 | 3.88E-33 | 0.520763 | 1.48E-63 | 1.255437 | 9.92E-05 | 1.391993 | 4.08E-10 | 0.522028 | 1.81E-19 |
| L89A A122D V196P | 1 | 0.599325 | 0.058437 | 0 | 0.84251 | 4.19E-82 | 0.890329 | 5.14E-21 | 1.033409 | 0.010531 | 1.519039 | 6.37E-137 |
| Y63L N81S L89G A122H | 1 | 0.599112 | 0.08267 | 4.59E-193 | 0.761532 | 3.19E-77 | 0.656356 | 7.48E-30 | 0.654013 | 1.40E-39 | 1.26395 | 1.84E-21 |
| Y63L V74W R78H N81S L89C | 1 | 0.598938 | 0.063705 | 9.78E-97 | 0.781861 | 8.28E-26 | 0.849468 | 1.62E-06 | 0.939516 | 0.777751 | 1.347434 | 2.15E-11 |
| Y63L N69E V74W N81S A122H V196P | 1 | 0.598836 | 0.066103 | 3.50E-245 | 0.634167 | 1.03E-188 | 0.887998 | 2.13E-09 | 1.012254 | 0.212205 | 0.784439 | 6.15E-10 |
| L89A A122H | 1 | 0.598706 | 0.061013 | 7.63E-93 | 0.744315 | 2.50E-39 | 0.664397 | 5.67E-14 | 0.845673 | 0.000485 | 1.229576 | 1.92E-08 |
| Y63L V74W A122D | 1 | 0.597411 | 0.051514 | 4.38E-66 | 0.586694 | 1.05E-87 | 0.96168 | 0.795132 | 0.918673 | 0.008803 | 0.781827 | 2.14E-13 |

| | | | | | | | | | | | |
|---|---|---|---|---|---|---|---|---|---|---|---|
| V74E L89A A122H V196P | 1 | 0.597212 | 0.068771 | 1.05E-37 | 0.656958 | 8.78E-39 | 1.087644 | 0.206956 | 0.89663 | 0.04453 | 1.229908 | 0.013207683 |
| V74E N81S L89G V196P | 1 | 0.597085 | 0.06394 | 1.56E-297 | 0.717653 | 3.41E-129 | 0.774698 | 5.01E-42 | 0.665941 | 2.32E-62 | 1.067616 | 0.001818345 |
| Y63L N69E V74W L89C A122D | 1 | 0.597017 | 0.055137 | 2.02E-63 | 0.77694 | 9.13E-32 | 0.880189 | 1.21E-05 | 1.060719 | 0.22718 | 1.483591 | 2.22E-13 |
| V74E R78H A122D V196P | 1 | 0.597015 | 0.05148 | 1.10E-81 | 0.553065 | 3.72E-131 | 1.204972 | 1.55E-09 | 1.048535 | 0.000301 | 0.671965 | 6.55E-25 |
| V74E R78H N81S L89C A122H | 1 | 0.596941 | 0.052954 | 1.99E-156 | 0.701096 | 1.19E-93 | 1.112359 | 0.028923 | 1.142744 | 0.000326 | 1.115269 | 0.020186573 |
| R78H N81S L89A A122D | 1 | 0.594748 | 0.046368 | 0 | 0.662755 | 2.34E-175 | 0.894064 | 1.61E-10 | 0.842239 | 8.79E-07 | 1.169918 | 4.62E-14 |
| N69E L89C A122H V196P | 1 | 0.594371 | 0.043419 | 1.63E-198 | 0.819887 | 4.52E-37 | 0.639096 | 2.12E-67 | 0.785743 | 4.28E-23 | 1.850217 | 4.04E-85 |
| Y63L N69E N81S L89A V196P | 1 | 0.593917 | 0.046303 | 0 | 0.691052 | 2.27E-167 | 0.711389 | 3.81E-54 | 0.759903 | 2.71E-35 | 1.307146 | 3.02E-52 |
| Y63L V74W N81S A122H | 1 | 0.592725 | 0.061306 | 7.87E-107 | 0.690596 | 6.04E-73 | 1.21967 | 3.54E-11 | 1.019025 | 0.715466 | 1.010476 | 0.987325769 |
| Y63L V74E N81S A122D | 1 | 0.592535 | 0.052933 | 5.42E-34 | 0.570531 | 6.57E-56 | 1.74839 | 8.86E-17 | 1.18664 | 0.000619 | 0.719763 | 3.70E-07 |
| V74W L89C | 1 | 0.59253 | 0.058069 | 1.63E-237 | 0.619472 | 1.21E-166 | 0.828089 | 1.79E-31 | 0.769066 | 1.59E-40 | 1.020625 | 0.937368663 |
| Y63L N69E N81S L89T A122H | 1 | 0.591482 | 0.048126 | 1.10E-72 | 0.6328 | 1.67E-58 | 1.115015 | 0.003231 | 0.992998 | 0.524614 | 1.022274 | 0.110601369 |
| Y63L N69E V74W A122H | 1 | 0.590543 | 0.054554 | 6.54E-112 | 0.692883 | 7.49E-53 | 1.294583 | 4.73E-23 | 1.181122 | 1.25E-05 | 1.023627 | 0.135265942 |
| L89C A122H | 1 | 0.590443 | 0.05453 | 0 | 0.781267 | 7.50E-169 | 1.116256 | 1.61E-12 | 1.266377 | 1.27E-61 | 2.049223 | 0 |
| N69E V74E L89C A122H V196P | 1 | 0.59038 | 0.043127 | 1.77E-104 | 0.838609 | 3.00E-26 | 0.547417 | 8.02E-50 | 0.704014 | 8.30E-14 | 1.623513 | 1.01E-27 |
| N69E R78H L89G A122H | 1 | 0.59009 | 0.052354 | 6.55E-69 | 0.608529 | 1.41E-63 | 1.117205 | 0.000723 | 0.901961 | 0.071565 | 0.959344 | 0.025960191 |
| N69E V74E R78H N81S L89C V196P | 1 | 0.588328 | 0.081182 | 1.56E-88 | 0.613816 | 4.60E-65 | 0.825877 | 5.34E-05 | 1.082716 | 0.166572 | 1.087491 | 0.055546964 |
| N69E V74W R78H N81S L89A A122D | 1 | 0.587492 | 0.038522 | 1.05E-40 | 0.620209 | 1.91E-33 | 0.934356 | 0.126134 | 0.749153 | 0.000208 | 1.015114 | 0.674100013 |
| Y63L R78H N81S L89G A122D | 1 | 0.58698 | 0.060771 | 8.07E-65 | 0.689772 | 5.08E-37 | 0.592231 | 6.83E-22 | 0.661378 | 2.81E-17 | 1.147921 | 0.219984973 |
| N81S L89A V196P | 1 | 0.586338 | 0.045713 | 0 | 0.694203 | 9.20E-253 | 0.715426 | 3.81E-77 | 0.798067 | 5.50E-40 | 1.312482 | 2.31E-90 |
| V74W R78H N81S L89G A122H V196P | 1 | 0.586071 | 0.060901 | 3.53E-91 | 0.698206 | 1.74E-73 | 0.949869 | 5.74E-08 | 0.954435 | 1.36E-06 | 1.28373 | 2.46E-07 |
| Y63L N69E V74W R78H N81S | 1 | 0.584761 | 0.067363 | 2.36E-280 | 0.647252 | 1.06E-229 | 1.159786 | 5.01E-09 | 1.285618 | 5.77E-29 | 1.03733 | 0.000273869 |
| R78H L89T | 1 | 0.583971 | 0.044016 | 0 | 0.709912 | 0 | 0.778932 | 3.61E-69 | 1.075949 | 4.20E-06 | 1.492434 | 2.59E-155 |
| Y63L N69E L89T A122D V196P | 1 | 0.583784 | 0.05543 | 0 | 0.675571 | 0 | 0.789621 | 5.32E-111 | 1.173898 | 8.86E-32 | 1.470848 | 2.66E-244 |
| N69E L89A V196P | 1 | 0.583415 | 0.059362 | 0 | 0.67386 | 7.44E-247 | 0.768238 | 1.05E-53 | 0.729677 | 2.56E-58 | 1.007921 | 0.82592253 |
| N81S L89C A122H | 1 | 0.58319 | 0.042565 | 0 | 0.618719 | 0 | 1.029785 | 4.47E-05 | 1.853559 | 1.27E-246 | 1.008311 | 0.600076211 |
| Y63L V74W R78H N81S A122H V196P | 1 | 0.583117 | 0.068852 | 1.45E-306 | 0.560198 | 428737619 | 1.333463 | 2.79E-70 | 0.835987 | 3.47E-14 | 0.766782 | 6.76E-41 |
| Y63L N69E V74E N81S V196P | 1 | 0.582871 | 0.05207 | 1.42E-38 | 0.65552 | 6.15E-41 | 1.438176 | 1.08E-14 | 1.502112 | 2.77E-10 | 1.016367 | 0.732400666 |
| N69E V74W R78H L89C | 1 | 0.581638 | 0.045252 | 4.31E-158 | 0.643414 | 8.61E-114 | 1.336733 | 4.13E-12 | 1.693233 | 1.54E-28 | 1.056119 | 0.76244054 |
| N69E V74W L89A A122D V196P | 1 | 0.581483 | 0.056698 | 0 | 0.684354 | 3.93E-187 | 0.858615 | 1.13E-11 | 1.161478 | 3.39E-13 | 1.255194 | 1.20E-26 |
| Y63L N69E V74W R78H A122H | 1 | 0.581357 | 0.053036 | 1.72E-105 | 0.544098 | 4.47E-137 | 1.472936 | 8.38E-47 | 1.350667 | 2.17E-29 | 0.718916 | 9.50E-15 |
| Y63L N69E V74E N81S L89C A122H V196P | 1 | 0.581272 | 0.065949 | 1.03E-49 | 0.605794 | 1.76E-76 | 0.924715 | 0.35007 | 0.794811 | 0.003163 | 1.220038 | 0.002562873 |
| N69E V74W L89G A122D | 1 | 0.580676 | 0.050011 | 2.37E-132 | 0.703703 | 4.78E-50 | 0.872723 | 0.001959 | 0.840424 | 6.48E-06 | 1.308724 | 1.42E-09 |
| L89T A122H | 1 | 0.580547 | 0.042967 | 0 | 0.709794 | 0 | 0.814572 | 8.87E-69 | 1.057891 | 0.00191 | 1.368565 | 1.06E-112 |
| L89C V196P | 1 | 0.579881 | 0.057864 | 7.41E-45 | 0.746525 | 1.09E-19 | 1.047866 | 0.13247 | 0.591031 | 1.11E-09 | 1.150891 | 0.000355476 |
| Y63L V74W N81S A122D | 1 | 0.579302 | 0.051397 | 2.78E-139 | 0.644102 | 2.17E-94 | 1.076154 | 0.246658 | 1.218637 | 5.98E-08 | 1.031699 | 0.800292558 |
| Y63L N69E L89T A122H | 1 | 0.578753 | 0.054952 | 0 | 0.609979 | 0 | 0.733163 | 9.01E-140 | 0.974919 | 0.055162 | 1.014386 | 0.055476125 |
| N69E V74E R78H N81S L89C A122D V196P | 1 | 0.578425 | 0.072749 | 9.48E-75 | 0.659675 | 7.33E-101 | 0.992338 | 0.500034 | 0.607956 | 3.64E-28 | 1.241469 | 2.45E-05 |
| V74W N81S L89C V196P | 1 | 0.578356 | 0.043137 | 0 | 0.706499 | 1.11E-271 | 0.88343 | 9.66E-12 | 1.316717 | 1.99E-45 | 1.49459 | 7.94E-146 |
| L89T A122D V196P | 1 | 0.576833 | 0.042396 | 1.03E-161 | 0.703724 | 1.31E-72 | 0.954131 | 0.201765 | 1.255464 | 4.32E-05 | 1.357924 | 7.27E-21 |
| N69E V74W R78H N81S L89G A122H V196P | 1 | 0.576654 | 0.060781 | 1.18E-73 | 0.885189 | 3.74E-05 | 0.553063 | 1.02E-23 | 1.273155 | 6.10E-05 | 1.401206 | 1.31E-10 |
| V74E N81S L89C A122D V196P | 1 | 0.575101 | 0.032171 | 7.79E-62 | 0.722556 | 8.07E-30 | 0.92485 | 0.731755 | 1.759268 | 3.40E-07 | 1.414308 | 1.98E-20 |
| N81S A122H | 1 | 0.574896 | 0.059462 | 0 | 0.711653 | 1.36E-279 | 1.122385 | 3.59E-22 | 1.499311 | 1.51E-42 | 1.282355 | 3.71E-69 |
| Y63L N69E V74E N81S L89C A122D V196P | 1 | 0.574686 | 0.042864 | 1.07E-79 | 0.537539 | 8.97E-80 | 1.462795 | 3.04E-20 | 1.182455 | 0.000861 | 0.700907 | 1.16E-10 |
| Y63L N69E N81S A122H V196P | 1 | 0.573908 | 0.069895 | 0 | 0.532209 | 0 | 0.88281 | 4.51E-12 | 1.172338 | 7.68E-26 | 0.688445 | 3.92E-74 |
| V74E N81S A122D V196P | 1 | 0.573807 | 0.044643 | 2.11E-153 | 0.654728 | 2.62E-99 | 1.17166 | 0.00112 | 1.178737 | 5.80E-07 | 1.14263 | 4.55E-05 |
| Y63L N69E L89A A122D V196P | 1 | 0.573807 | 0.058705 | 0 | 0.765957 | 8.33E-101 | 0.842874 | 1.16E-30 | 0.471169 | 1.35E-274 | 1.783834 | 6.35E-123 |
| V74E R78H L89C A122D V196P | 1 | 0.573377 | 0.037429 | 2.34E-43 | 0.584786 | 2.09E-51 | 4.403305 | 2.07E-17 | 0.158946 | 2.45E-09 | 0.838459 | 0.003037659 |

| | | | | | | | | | | | | |
|---|---|---|---|---|---|---|---|---|---|---|---|---|
| Y63L N69E R78H N81S L89G A122D | 1 | 0.57248 | 0.05927 | 5.30E-30 | 0.547157 | 3.55E-64 | 1.003792 | 0.727945 | 1.058106 | 0.505067 | 0.882259 | 2.69E-05 |
| Y63L N69E V74W N81S L89T A122H | 1 | 0.571715 | 0.046846 | 1.43E-112 | 0.679566 | 5.37E-63 | 1.131179 | 0.002525 | 1.150298 | 0.137872 | 1.235591 | 1.14E-08 |
| R78H L89G V196P | 1 | 0.571284 | 0.060215 | 7.24E-103 | 0.751434 | 2.28E-32 | 0.816891 | 7.52E-05 | 1.590125 | 5.54E-30 | 1.501946 | 1.77E-17 |
| N69E V74E R78H A122H | 1 | 0.570207 | 0.057384 | 4.13E-29 | 0.891307 | 0.000359 | 0.53573 | 2.43E-10 | 1.15035 | 0.471156 | 1.599528 | 9.38E-09 |
| N69E R78H N81S L89G A122D V196P | 1 | 0.569304 | 0.059508 | 4.92E-47 | 0.749872 | 8.86E-34 | 0.916648 | 0.059213 | 0.657201 | 2.14E-18 | 1.003603 | 0.370064752 |
| Y63L N69E V74W L89G A122H V196P | 1 | 0.568853 | 0.051895 | 5.45E-40 | 0.547628 | 2.60E-52 | 0.889089 | 0.317173 | 0.903121 | 0.395452 | 0.743893 | 6.37E-10 |
| N69E V74E N81S L89G | 1 | 0.568258 | 0.045271 | 7.61E-245 | 0.5733 | 1.68E-196 | 1.03077 | 0.172694 | 0.946873 | 0.002013 | 0.763049 | 4.40E-31 |
| Y63L V74E | 1 | 0.567951 | 0.052467 | 3.18E-54 | 0.54879 | 5.59E-90 | 2.039513 | 1.91E-44 | 1.624058 | 6.34E-24 | 0.600249 | 4.38E-09 |
| N69E L89A A122D V196P | 1 | 0.567314 | 0.052056 | 973092054 | 0.577361 | 1.09E-268 | 0.759843 | 1.63E-56 | 0.765017 | 7.15E-51 | 1.074234 | 0.21360961 |
| N69E V74W R78H N81S L89T A122H V196P | 1 | 0.566802 | 0.041405 | 0 | 0.787095 | 2.35E-152 | 0.895223 | 6.22E-07 | 1.31336 | 2.59E-50 | 1.920869 | 3.59587893962301e-318 |
| Y63L N69E R78H N81S L89A A122D | 1 | 0.565537 | 0.038514 | 4.27E-128 | 0.603361 | 2.58E-100 | 0.921037 | 0.002471 | 1.12866 | 0.010139 | 1.069009 | 0.009539904 |
| N69E V74W R78H N81S L89G A122H | 1 | 0.564745 | 0.035769 | 1.10E-95 | 0.642522 | 4.86E-68 | 0.740191 | 7.76E-11 | 0.907787 | 0.022747 | 1.050747 | 0.949103796 |
| N69E V74W L89C A122H V196P | 1 | 0.563119 | 0.041136 | 1.33E-67 | 0.729038 | 4.30E-29 | 0.45995 | 2.88E-34 | 0.957075 | 0.091938 | 1.499473 | 1.10E-10 |
| V74E L89G A122H | 1 | 0.56284 | 0.048475 | 6.70E-88 | 0.684106 | 2.23E-44 | 0.811812 | 0.000332 | 0.84847 | 0.005517 | 1.233126 | 0.002645811 |
| Y63L N69E V74W L89T A122H V196P | 1 | 0.562607 | 0.040045 | 2.16E-171 | 0.684142 | 2.54E-95 | 0.872166 | 0.002565 | 0.690287 | 5.20E-31 | 1.27147 | 3.39E-19 |
| Y63L N69E V74E A122D V196P | 1 | 0.562388 | 0.043407 | 7.16E-56 | 0.601588 | 8.29E-69 | 1.008385 | 0.510819 | 1.415341 | 1.50E-08 | 1.008862 | 0.214814623 |
| N81S L89T A122D V196P | 1 | 0.561268 | 0.058052 | 1.13E-221 | 0.799199 | 3.12E-58 | 0.727834 | 1.11E-25 | 0.615103 | 7.53E-64 | 1.616915 | 5.15E-76 |
| V74W N81S L89C | 1 | 0.560463 | 0.041803 | 2.50E-298 | 0.650504 | 1.92E-140 | 0.881125 | 2.71E-07 | 1.474091 | 4.98E-39 | 1.352353 | 1.92E-34 |
| Y63L N69E V74W A122H V196P | 1 | 0.560113 | 0.056456 | 3.18E-228 | 0.559837 | 8.49E-226 | 0.93947 | 0.000112 | 1.009221 | 0.353165 | 0.7953 | 5.32E-22 |
| V74E L89T V196P | 1 | 0.560039 | 0.045568 | 0 | 0.699545 | 3.89E-149 | 0.811124 | 3.04E-15 | 0.849181 | 1.58E-19 | 1.406462 | 6.79E-51 |
| N69E V74W L89G | 1 | 0.559637 | 0.054846 | 1.78E-209 | 0.539028 | 3.82E-220 | 0.94859 | 0.012708 | 1.238297 | 2.34E-15 | 0.733408 | 6.26E-28 |
| N69E V74E L89T A122D | 1 | 0.558592 | 0.042103 | 1.25E-303 | 0.683936 | 1.73E-166 | 0.950443 | 0.055301 | 1.014048 | 0.744289 | 1.725373 | 5.28E-106 |
| Y63L N69E R78H N81S L89A A122D V196P | 1 | 0.557126 | 0.043435 | 1.53E-62 | 0.560665 | 7.42E-62 | 0.755693 | 0.003146 | 0.70697 | 6.26E-06 | 0.890197 | 0.002808504 |
| V74E N81S L89A V196P | 1 | 0.556958 | 0.070049 | 1.23E-204 | 0.78765 | 1.09E-67 | 0.912287 | 6.60E-05 | 0.69686 | 3.12E-34 | 1.640889 | 1.01E-52 |
| Y63L N69E V74W R78H N81S L89C A122D V196P | 1 | 0.555793 | 0.03785 | 2.85E-104 | 0.588632 | 1.59E-83 | 1.263275 | 0.00021 | 1.338695 | 1.83E-09 | 0.912241 | 0.013133642 |
| N69E V74E N81S L89T A122D | 1 | 0.555012 | 0.055382 | 9.13E-23 | 0.78376 | 1.81E-09 | 0.749599 | 0.002762 | 1.017056 | 0.441559 | 1.338335 | 0.010693553 |
| Y63L V74E N81S L89T | 1 | 0.554963 | 0.059076 | 0 | 0.645572 | 4.69E-290 | 0.781846 | 7.00E-44 | 0.741799 | 3.60E-73 | 1.017335 | 0.832070007 |
| N69E R78H N81S A122H | 1 | 0.554137 | 0.044328 | 0 | 0.598788 | 0 | 1.48907 | 3.35E-108 | 1.145756 | 0.00344 | 1.139921 | 1.87E-15 |
| N69E V74W L89G A122H | 1 | 0.552717 | 0.051507 | 3.26E-51 | 0.815515 | 8.00E-12 | 0.491335 | 1.93E-31 | 0.61264 | 1.26E-13 | 1.570957 | 3.41E-08 |
| V74W R78H N81S L89C | 1 | 0.550603 | 0.054557 | 0 | 0.581615 | 8.63E-295 | 1.077304 | 5.81E-08 | 1.097474 | 9.74E-10 | 1.07779 | 0.006148123 |
| N69E V74E L89A V196P | 1 | 0.549188 | 0.053549 | 7.97E-90 | 0.644524 | 3.24E-50 | 0.97627 | 0.003182 | 1.212801 | 9.00E-05 | 1.006309 | 0.407271059 |
| Y63L N81S L89T A122H | 1 | 0.546513 | 0.044467 | 0 | 0.708983 | 1.21E-112 | 1.164384 | 1.30E-14 | 0.823561 | 3.73E-31 | 1.569636 | 4.19E-93 |
| N69E R78H N81S L89T A122H V196P | 1 | 0.545982 | 0.033504 | 1.98E-74 | 0.722841 | 4.76E-32 | 0.85476 | 0.112512 | 0.831161 | 4.45E-06 | 1.695562 | 3.90E-21 |
| N69E V74W L89C A122D | 1 | 0.545716 | 0.043654 | 3.31E-91 | 0.645399 | 5.89E-50 | 0.753552 | 0.003317 | 0.904483 | 0.013507 | 1.176039 | 0.000266049 |
| N69E N81S L89A V196P | 1 | 0.545507 | 0.035769 | 1.42E-49 | 0.552575 | 1.42E-41 | 0.969554 | 0.361264 | 0.71221 | 3.95E-05 | 0.961035 | 0.035912258 |
| N69E V74W L89C A122H | 1 | 0.545399 | 0.05037 | 1.99E-169 | 0.707068 | 1.56E-90 | 1.068028 | 1.19E-07 | 0.7559 | 8.08E-15 | 1.641756 | 1.31E-51 |
| V74E R78H N81S A122H | 1 | 0.545394 | 0.057486 | 4.89E-121 | 0.668572 | 3.50E-79 | 1.445244 | 7.19E-26 | 1.482142 | 2.60E-41 | 1.24664 | 2.52E-09 |
| V74W L89T V196P | 1 | 0.545255 | 0.061862 | 7.52E-102 | 0.782203 | 5.19E-39 | 0.61214 | 9.60E-31 | 0.583937 | 2.81E-36 | 1.647143 | 1.17E-32 |
| N69E V74E L89T V196P | 1 | 0.544961 | 0.041075 | 0 | 0.69318 | 6.70E-191 | 0.837837 | 1.27E-18 | 0.714568 | 8.69E-65 | 1.525948 | 6.75E-80 |
| Y63L V74E N81S L89A | 1 | 0.543738 | 0.084575 | 5.01E-256 | 0.533218 | 2.28E-300 | 0.683245 | 1.69E-100 | 0.863277 | 2.49E-17 | 0.939386 | 0.000126125 |
| Y63L N69E V74W L89T A122H | 1 | 0.542982 | 0.051334 | 1.57E-143 | 0.585848 | 1.62E-90 | 0.732911 | 3.45E-14 | 1.495687 | 1.38E-21 | 1.089413 | 0.213352692 |
| Y63L N69E V74E N81S L89G V196P | 1 | 0.542815 | 0.056198 | 1.99E-51 | 0.571461 | 4.27E-69 | 0.85944 | 0.157476 | 0.666696 | 3.04E-12 | 0.762258 | 1.51E-07 |
| N69E N81S L89G A122D | 1 | 0.542581 | 0.058103 | 4.26E-272 | 0.662817 | 3.47E-136 | 1.001159 | 0.510352 | 0.774868 | 3.16E-17 | 1.158477 | 4.47E-05 |
| Y63L N69E V74E L89T A122D | 1 | 0.541638 | 0.040825 | 3.78E-209 | 0.582299 | 7.36E-173 | 0.83281 | 3.57E-05 | 1.229293 | 4.81E-08 | 1.031001 | 0.164152178 |
| N69E V74E N81S L89G A122D | 1 | 0.541503 | 0.057983 | 1.18E-50 | 0.584992 | 4.67E-43 | 0.836943 | 0.000291 | 1.081257 | 0.016472 | 1.042851 | 0.957106912 |
| V74W R78H L89A A122H V196P | 1 | 0.540153 | 0.04175 | 7.90E-245 | 0.612627 | 9.69E-152 | 0.986166 | 0.857772 | 1.550771 | 1.11E-35 | 1.113781 | 0.000758575 |
| Y63L V74W N81S L89T A122H V196P | 1 | 0.539445 | 0.047854 | 1.85E-43 | 0.740418 | 2.10E-12 | 0.659503 | 5.43E-05 | 0.694783 | 1.82E-05 | 1.904177 | 5.15E-15 |
| Y63L V196P | 1 | 0.536952 | 0.063093 | 5.27E-148 | 0.621246 | 6.58E-105 | 1.637919 | 3.51E-32 | 1.124092 | 0.000147 | 1.01281 | 0.730840733 |

| | | | | | | | | | | | | |
|---|---|---|---|---|---|---|---|---|---|---|---|---|
| Y63L V74W N81S L89C A122D | 1 | 0.536395 | 0.061825 | 3.92E-87 | 0.564035 | 2.25E-109 | 0.945755 | 0.28666 | 0.899853 | 0.024437 | 0.990516 | 2.14E-07 |
| Y63L V74E L89T A122D | 1 | 0.536013 | 0.047549 | 9.07E-66 | 0.615157 | 3.33E-47 | 1.263915 | 2.57E-06 | 0.89012 | 0.085235 | 1.124377 | 0.103110676 |
| Y63L N69E N81S L89C A122D V196P | 1 | 0.535699 | 0.053995 | 1.36E-267 | 0.617243 | 3.60E-203 | 0.927141 | 0.00018 | 0.944667 | 0.343854 | 1.063148 | 0.001453014 |
| V74E L89G A122H V196P | 1 | 0.534984 | 0.048883 | 1.64E-74 | 0.551796 | 1.65E-72 | 1.131091 | 0.001609 | 1.033148 | 0.630216 | 0.936855 | 0.001522549 |
| L89G A122H V196P | 1 | 0.534949 | 0.047986 | 1.62E-82 | 0.667637 | 1.03E-33 | 1.108177 | 0.144828 | 0.910081 | 0.292952 | 1.54239 | 4.22E-08 |
| N69E V74E N81S A122D V196P | 1 | 0.534826 | 0.048869 | 8.23E-64 | 0.534372 | 1.12E-80 | 1.172217 | 7.19E-08 | 1.349927 | 4.41E-07 | 0.805846 | 3.74E-13 |
| N69E V74E N81S L89A | 1 | 0.533354 | 0.037963 | 1.08E-254 | 0.650049 | 8.47E-99 | 1.11107 | 3.47E-05 | 1.139247 | 2.93E-07 | 1.422819 | 1.07E-57 |
| V74W R78H N81S L89C V196P | 1 | 0.533162 | 0.052829 | 594784200 | 0.492545 | 1.66E-284 | 1.306919 | 1.03E-37 | 1.070564 | 1.65E-06 | 0.778096 | 1.33E-37 |
| Y63L N69E V74W N81S L89C A122H | 1 | 0.532562 | 0.052769 | 0 | 0.52588 | 0 | 1.062337 | 0.000247 | 1.211821 | 1.35E-23 | 0.915781 | 5.31E-08 |
| R78H N81S L89T V196P | 1 | 0.530513 | 0.04347 | 1.15E-133 | 0.672656 | 5.09E-69 | 1.055675 | 0.045099 | 1.209203 | 0.023755 | 1.371443 | 3.45E-18 |
| N69E L89T A122D V196P | 1 | 0.529482 | 0.050274 | 0 | 0.615956 | 0 | 0.692515 | 3.16E-134 | 1.001713 | 0.040231 | 1.510711 | 3.92E-200 |
| N69E V74W L89C | 1 | 0.529222 | 0.048007 | 0 | 0.645264 | 7.65E-219 | 1.023462 | 0.005948 | 1.257687 | 1.05E-28 | 1.109614 | 3.50E-15 |
| Y63L N69E N81S L89A | 1 | 0.528565 | 0.042245 | 1.57E-67 | 0.557714 | 2.66E-65 | 0.720318 | 1.14E-10 | 1.329605 | 6.15E-06 | 0.953168 | 0.008474804 |
| Y63L N81S L89G V196P | 1 | 0.528121 | 0.074613 | 0 | 0.575067 | 2.04E-213 | 0.593419 | 1.15E-229 | 0.555394 | 7.08E-223 | 0.970023 | 0.747585275 |
| Y63L N69E V74W N81S L89T A122H V196P | 1 | 0.527476 | 0.038532 | 8.69E-81 | 0.733887 | 3.90E-28 | 0.726214 | 1.99E-09 | 0.922408 | 0.230723 | 1.617394 | 4.58E-20 |
| V74E R78H N81S A122D V196P | 1 | 0.52727 | 0.048178 | 1.23E-125 | 0.577078 | 1.35E-119 | 1.234951 | 5.61E-09 | 1.025429 | 0.149999 | 0.947333 | 0.000365581 |
| V74E L89A A122D | 1 | 0.52717 | 0.051402 | 1.47E-83 | 0.609114 | 1.71E-70 | 0.877945 | 0.036547 | 1.249392 | 4.71E-05 | 1.052266 | 0.690352721 |
| N69E V74E L89A | 1 | 0.526243 | 0.054519 | 0 | 0.642139 | 0 | 0.846746 | 9.89E-36 | 0.94166 | 0.007189 | 1.050702 | 0.006910833 |
| Y63L N69E R78H N81S L89C A122H V196P | 1 | 0.525953 | 0.033312 | 4.20E-74 | 0.608963 | 7.83E-61 | 0.660168 | 2.72E-10 | 1.007353 | 0.552726 | 0.996409 | 0.999028667 |
| R78H N81S L89T A122H V196P | 1 | 0.523427 | 0.052553 | 1.11E-52 | 0.656859 | 2.69E-25 | 1.886251 | 2.05E-16 | 1.926595 | 4.11E-17 | 1.540832 | 5.22E-07 |
| N69E V74W N81S L89G A122H V196P | 1 | 0.522801 | 0.03989 | 4.86E-35 | 0.74073 | 3.81E-14 | 0.437141 | 4.63E-11 | 0.893984 | 0.204405 | 1.830424 | 2.06E-09 |
| L89T A122H V196P | 1 | 0.522521 | 0.037192 | 0.00E+00 | 0.592203 | 1.54E-226 | 0.755007 | 2.35E-27 | 1.131081 | 6.53E-07 | 1.169019 | 7.50E-11 |
| Y63L V74W L89T A122D | 1 | 0.522238 | 0.038651 | 3.73E-86 | 0.539124 | 4.15E-102 | 0.904358 | 0.048976 | 1.002324 | 0.784507 | 1.06208 | 0.008713536 |
| Y63L V74E N81S L89G V196P | 1 | 0.520612 | 0.055751 | 2.50E-125 | 0.569993 | 4.27E-115 | 0.806896 | 1.71E-06 | 0.856434 | 0.01243 | 0.965681 | 0.117747136 |
| Y63L N69E R78H N81S L89T A122D | 1 | 0.518958 | 0.040111 | 6.51E-62 | 0.55659 | 6.79E-84 | 1.329442 | 1.09E-11 | 1.284477 | 2.34E-06 | 0.887131 | 0.000615621 |
| V74W R78H N81S L89T | 1 | 0.518871 | 0.040369 | 0 | 0.644508 | 3.92E-273 | 1.353859 | 2.16E-34 | 1.691738 | 3.99E-102 | 1.518362 | 1.07E-104 |
| Y63L V74W L89T A122H | 1 | 0.518857 | 0.049265 | .972832126 | 0.570027 | 7.67E-249 | 0.773292 | 1.54E-41 | 1.093948 | 0.0032 | 1.178228 | 1.69E-14 |
| N69E N81S L89A A122D | 1 | 0.518641 | 0.040435 | 1.50E-201 | 0.622765 | 1.14E-106 | 0.981708 | 0.524517 | 0.780949 | 2.00E-11 | 1.424396 | 6.80E-27 |
| V74E N81S L89G A122D | 1 | 0.518517 | 0.053683 | 5.56E-177 | 0.620021 | 1.31E-132 | 1.127047 | 6.53E-06 | 0.914282 | 0.005113 | 1.320731 | 3.83E-15 |
| N69E V74W N81S V196P | 1 | 0.517929 | 0.041431 | 3.85E-239 | 0.569879 | 2.59E-179 | 0.798708 | 1.98E-06 | 1.385657 | 4.34E-14 | 1.092182 | 0.002459969 |
| V74E A122H | 1 | 0.516641 | 0.046344 | 1.36E-189 | 0.689582 | 7.58E-60 | 1.017766 | 0.434683 | 1.221063 | 5.54E-12 | 1.581414 | 1.27E-28 |
| V74W R78H L89T A122H V196P | 1 | 0.516535 | 0.037965 | 1.07E-158 | 0.581699 | 1.71E-106 | 0.569512 | 8.86E-19 | 1.447612 | 4.35E-18 | 1.162274 | 4.18E-05 |
| Y63L N69E V74W R78H N81S L89C | 1 | 0.516301 | 0.053651 | 1.04E-48 | 0.680234 | 3.13E-30 | 1.450106 | 4.94E-10 | 1.280463 | 0.12312 | 1.40894 | 2.08E-05 |
| R78H L89T A122D V196P | 1 | 0.51513 | 0.037861 | 4.25E-268 | 0.616119 | 8.88E-172 | 0.914188 | 2.00E-05 | 1.032979 | 0.940617 | 1.349614 | 1.91E-30 |
| N81S L89T A122H | 1 | 0.514357 | 0.041109 | 4.87E-307 | 0.586839 | 6.40E-235 | 0.949758 | 0.306383 | 1.158465 | 9.40E-08 | 1.144136 | 1.48E-10 |
| Y63L N69E V74W R78H N81S L89T A122H | 1 | 0.513552 | 0.041045 | 1.54E-90 | 0.558674 | 1.05E-79 | 1.7901 | 4.77E-34 | 1.263931 | 0.000199 | 0.940114 | 0.012097574 |
| N69E V74E N81S L89G V196P | 1 | 0.512881 | 0.045497 | 1.42E-37 | 0.587772 | 9.02E-19 | 0.784434 | 0.2829 | 0.87669 | 0.168348 | 1.333501 | 0.004424443 |
| N69E V74E A122H | 1 | 0.512092 | 0.051536 | 8.89E-45 | 0.778584 | 7.12E-13 | 1.018392 | 0.606489 | 1.165715 | 0.025532 | 1.631978 | 5.28E-13 |
| Y63L N69E V74W L89A A122H | 1 | 0.511689 | 0.054469 | 2.03E-247 | 0.553914 | 1.22E-199 | 0.97486 | 0.424083 | 1.31402 | 2.06E-19 | 0.999843 | 0.772400361 |
| Y63L N69E V74W N81S L89C A122D | 1 | 0.511366 | 0.051342 | 7.44E-88 | 0.757277 | 2.89E-32 | 0.984767 | 0.421592 | 1.023053 | 0.392184 | 1.989536 | 9.64E-21 |
| V74E R78H N81S A122H V196P | 1 | 0.511066 | 0.04636 | 1.23E-207 | 0.609132 | 2.74E-143 | 1.072757 | 0.002326 | 1.240648 | 1.70E-15 | 1.094342 | 0.009731672 |
| N81S L89G A122H | 1 | 0.510874 | 0.054338 | 2.44E-139 | 0.682759 | 2.93E-71 | 1.01984 | 0.57318 | 1.328688 | 1.00E-14 | 1.591714 | 1.24E-14 |
| Y63L V74W N81S L89C | 1 | 0.510827 | 0.050616 | 0 | 0.493483 | 894328658 | 1.18068 | 6.53E-17 | 1.214469 | 3.48E-18 | 0.856858 | 4.61E-20 |
| V74E L89G A122D V196P | 1 | 0.509944 | 0.046521 | 5.60E-44 | 0.478349 | 1.89E-72 | 1.001303 | 0.282296 | 1.377063 | 2.94E-05 | 0.696628 | 1.30E-06 |
| V74W N81S L89T V196P | 1 | 0.509465 | 0.051271 | 1.29E-121 | 0.762797 | 3.66E-53 | 0.976262 | 0.24662 | 0.999998 | 0.822725 | 1.621732 | 4.11E-44 |
| Y63L V74W R78H N81S L89T A122H V196P | 1 | 0.509431 | 0.050339 | 6.29E-65 | 0.551145 | 5.68E-94 | 1.045928 | 0.739238 | 1.252717 | 5.39E-07 | 1.195122 | 0.000833908 |
| N69E V74W L89A V196P | 1 | 0.509411 | 0.046743 | 2.04E-137 | 0.478456 | 1.59E-151 | 0.942437 | 0.036416 | 0.932136 | 0.007763 | 0.840391 | 3.66E-09 |
| Y63L V74E L89T V196P | 1 | 0.50704 | 0.047936 | 0 | 0.635825 | 1.70E-217 | 0.796294 | 1.37E-17 | 1.252709 | 6.20E-26 | 1.287892 | 5.91E-41 |

| | | | | | | | | | | | |
|---|---|---|---|---|---|---|---|---|---|---|---|
| Y63L N69E V74E N81S L89A V196P | 1 | 0.506308 | 0.035861 | 4.79E-175 | 0.585903 | 1.13E-119 | 0.997955 | 0.315324 | 0.840117 | 0.000139 | 1.06157 | 0.003941963 |
| N69E R78H N81S L89T | 1 | 0.505795 | 0.039886 | 5.09E-53 | 0.539811 | 1.88E-50 | 0.997112 | 0.920372 | 1.067764 | 0.321085 | 1.092573 | 0.834865672 |
| Y63L N69E R78H N81S A122H | 1 | 0.504406 | 0.045756 | 1.73E-64 | 0.544732 | 1.77E-50 | 1.153779 | 0.000912 | 1.629172 | 1.09E-14 | 0.799176 | 9.99E-07 |
| V74E R78H N81S L89T A122D V196P | 1 | 0.504206 | 0.053867 | 4.34E-91 | 0.669807 | 2.14E-58 | 0.917421 | 0.360761 | 0.939405 | 0.032607 | 1.449949 | 1.76E-16 |
| Y63L V74E R78H N81S V196P | 1 | 0.503669 | 0.044687 | 4.95E-59 | 0.596714 | 8.01E-43 | 1.537635 | 1.10E-16 | 1.277583 | 7.75E-08 | 1.160644 | 0.070376365 |
| Y63L N69E R78H N81S L89C A122H | 1 | 0.503295 | 0.056999 | 1.08E-118 | 0.63595 | 1.44E-127 | 1.067014 | 0.481574 | 1.272661 | 1.56E-09 | 1.244841 | 0.074798434 |
| Y63L N69E R78H N81S A122H V196P | 1 | 0.503231 | 0.061287 | 1.16E-41 | 0.364815 | 1.22E-80 | 1.193638 | 0.000488 | 1.039149 | 0.550041 | 0.372646 | 1.29E-31 |
| V74W L89T A122D V196P | 1 | 0.503039 | 0.036193 | .953842310 | 0.578544 | 1.58E-201 | 0.978358 | 0.391156 | 1.135022 | 2.49E-06 | 1.155475 | 5.95E-13 |
| V74W R78H N81S L89G A122D V196P | 1 | 0.501852 | 0.053742 | 5.54E-188 | 0.598603 | 1.91E-137 | 0.927033 | 0.141029 | 0.895686 | 0.00505 | 1.11484 | 0.152646396 |
| N69E R78H N81S L89T V196P | 1 | 0.501769 | 0.040103 | 1.67E-238 | 0.617968 | 7.27E-117 | 1.619175 | 6.67E-29 | 1.438028 | 2.04E-26 | 1.265057 | 5.56E-24 |
| N69E N81S A122H V196P | 1 | 0.499493 | 0.038607 | 2.96E-51 | 0.526836 | 6.11E-46 | 1.265785 | 0.001297 | 1.547976 | 1.21E-05 | 0.997705 | 0.829776627 |
| Y63L V74W N81S L89A A122H V196P | 1 | 0.498512 | 0.051612 | 1.66E-70 | 0.574993 | 3.09E-59 | 0.918654 | 0.278637 | 0.733701 | 1.23E-07 | 1.488686 | 3.61E-09 |
| N69E R78H L89A | 1 | 0.498236 | 0.038763 | 3.55E-83 | 0.567682 | 1.06E-71 | 1.296943 | 2.98E-06 | 1.461179 | 2.99E-11 | 1.12668 | 0.062847121 |
| Y63L V74W N81S L89T V196P | 1 | 0.497228 | 0.05004 | 1.92E-92 | 0.746633 | 3.66E-42 | 0.915068 | 0.945672 | 0.642843 | 6.50E-21 | 1.541263 | 1.86E-22 |
| Y63L N81S L89T A122H V196P | 1 | 0.497113 | 0.057085 | 3.11E-233 | 0.788166 | 8.05E-58 | 0.770468 | 1.91E-32 | 0.775911 | 2.07E-07 | 1.717948 | 2.90E-71 |
| Y63L V74E N81S L89C A122H | 1 | 0.496993 | 0.037069 | 8.20E-142 | 0.56481 | 4.29E-125 | 1.25495 | 6.74E-17 | 1.32025 | 1.89E-05 | 1.1704 | 0.016339947 |
| Y63L N69E R78H N81S L89G A122D V196P | 1 | 0.495686 | 0.070031 | 5.07E-77 | 0.57937 | 1.46E-27 | 0.804497 | 7.37E-09 | 0.787948 | 2.15E-06 | 1.40375 | 0.004169604 |
| V74W R78H L89G A122H V196P | 1 | 0.49471 | 0.045131 | 3.21E-154 | 0.611402 | 1.53E-106 | 0.860333 | 2.75E-05 | 1.096417 | 0.00012 | 1.293355 | 1.31E-09 |
| V74W R78H N81S L89G A122D | 1 | 0.494302 | 0.052933 | 6.00E-48 | 0.525999 | 9.58E-50 | 0.922975 | 0.378467 | 0.898728 | 0.913498 | 1.069784 | 0.760619667 |
| Y63L V74W L89A A122H | 1 | 0.49393 | 0.051327 | 8.03E-54 | 0.753035 | 1.83E-15 | 0.62127 | 8.51E-05 | 1.18078 | 0.159252 | 1.535392 | 6.68E-13 |
| N69E V74E L89T A122H V196P | 1 | 0.493888 | 0.067417 | 6.32E-42 | 0.602037 | 2.65E-49 | 0.74929 | 3.84E-10 | 0.698797 | 0.001003 | 1.301802 | 5.63E-09 |
| Y63L N81S L89A A122D V196P | 1 | 0.493784 | 0.034974 | 9.52E-266 | 0.571178 | 5.91E-186 | 1.00129 | 0.473061 | 0.928404 | 0.007812 | 1.278496 | 4.09E-27 |
| Y63L R78H N81S L89T | 1 | 0.493392 | 0.051573 | 1.21E-41 | 0.611991 | 1.85E-35 | 0.770322 | 0.001451 | 1.674252 | 0.001612 | 1.057864 | 0.908954222 |
| R78H N81S L89G | 1 | 0.493135 | 0.0272 | 3.11E-44 | 0.557503 | 1.73E-41 | 1.200185 | 0.404362 | 1.472293 | 5.91E-05 | 1.034046 | 0.217306359 |
| R78H L89C A122H | 1 | 0.492935 | 0.049852 | 2.27E-221 | 0.547267 | 3.87E-213 | 1.071833 | 0.060688 | 1.725854 | 2.55E-63 | 0.972347 | 0.343242866 |
| Y63L N81S L89A V196P | 1 | 0.492181 | 0.044647 | 2.62E-181 | 0.565021 | 4.93E-119 | 1.063502 | 0.511956 | 1.366503 | 7.83E-21 | 0.907553 | 0.000780391 |
| V74E L89T A122D | 1 | 0.491954 | 0.04651 | 0 | 0.639733 | 7.52E-230 | 0.723069 | 2.98E-28 | 0.759812 | 3.15E-34 | 1.584014 | 4.36E-118 |
| V74E L89T A122D V196P | 1 | 0.489844 | 0.034866 | 6.14E-259 | 0.584279 | 4.94E-155 | 0.674225 | 7.91E-11 | 1.887686 | 1.10E-69 | 1.36492 | 1.04E-39 |
| Y63L V74W N81S L89A V196P | 1 | 0.489628 | 0.032105 | 1.73E-84 | 0.513355 | 4.99E-66 | 0.97011 | 0.240811 | 1.604514 | 4.24E-08 | 0.986085 | 0.598504775 |
| N69E V74W L89T A122D | 1 | 0.488607 | 0.046194 | 0 | 0.596055 | '412414809 | 0.856215 | 1.14E-16 | 0.902228 | 4.32E-09 | 1.505667 | 3.44E-93 |
| N69E V74E R78H N81S L89C A122D | 1 | 0.488005 | 0.048354 | 0 | 0.50185 | 0 | 1.131231 | 5.83E-10 | 1.413242 | 4.94E-46 | 0.94819 | 1.38E-06 |
| V74E N81S L89T A122D | 1 | 0.487819 | 0.051929 | 1.11E-168 | 0.588588 | 9.22E-132 | 0.848053 | 2.59E-06 | 0.634396 | 6.30E-54 | 1.211019 | 4.98E-07 |
| Y63L V74E L89A | 1 | 0.487145 | 0.050468 | 7.01E-305 | 0.568398 | 1.39E-260 | 0.885989 | 7.63E-10 | 0.885602 | 1.46E-05 | 0.854643 | 1.23E-10 |
| N69E R78H N81S L89T A122H | 1 | 0.48467 | 0.044286 | 0 | 0.626893 | 6.32E-290 | 1.326017 | 1.15E-64 | 1.31102 | 5.33E-34 | 1.464243 | 1.36E-77 |
| N69E V74W R78H N81S L89T A122D | 1 | 0.484609 | 0.053494 | 0 | 0.730558 | 1.12E-220 | 0.908936 | 1.20E-11 | 1.17983 | 1.07E-30 | 1.957582 | 1.83E-242 |
| Y63L V74E N81S L89G | 1 | 0.484489 | 0.038597 | 1.17E-103 | 0.454656 | 4.25E-106 | 1.222597 | 2.27E-06 | 1.036616 | 0.705018 | 0.717016 | 2.80E-20 |
| Y63L N81S L89A | 1 | 0.48412 | 0.038692 | 2.35E-63 | 0.580142 | 2.32E-40 | 1.599012 | 1.02E-07 | 1.087369 | 0.028289 | 1.179058 | 0.012474568 |
| N69E V74W L89T A122D V196P | 1 | 0.483663 | 0.043928 | 0 | 0.562063 | 0 | 0.713009 | 1.68E-98 | 0.61968 | 6.83E-156 | 1.375434 | 9.52E-82 |
| N69E V74W R78H N81S L89G | 1 | 0.48343 | 0.060802 | 3.79E-104 | 0.608336 | 6.88E-86 | 1.267056 | 1.01E-07 | 0.869127 | 0.039301 | 1.405878 | 8.18E-14 |
| V74W R78H N81S L89G | 1 | 0.482111 | 0.030535 | 6.86E-112 | 0.593229 | 1.20E-81 | 1.155622 | 2.17E-05 | 1.012101 | 0.15967 | 1.421103 | 8.65E-15 |
| N69E V74W L89T | 1 | 0.481131 | 0.045683 | 0 | 0.599039 | 0 | 0.820537 | 4.34E-41 | 1.091152 | 0.043986 | 1.513138 | 6.13E-192 |
| N69E V74W R78H N81S L89A V196P | 1 | 0.480747 | 0.034798 | 1.91E-113 | 0.5304 | 3.47E-97 | 0.800648 | 0.002865 | 1.016313 | 0.708298 | 1.11321 | 0.226833416 |
| N69E N81S L89T A122H | 1 | 0.48041 | 0.039089 | 1.72E-85 | 0.657621 | 2.93E-38 | 1.15626 | 0.224258 | 1.048005 | 0.301428 | 1.397784 | 1.27E-10 |
| V74W L89A A122D | 1 | 0.479056 | 0.038322 | 4.62E-270 | 0.627733 | 1.64E-144 | 0.83757 | 3.69E-11 | 1.043125 | 0.693971 | 1.542005 | 5.55E-53 |
| Y63L N69E V74W L89T A122D V196P | 1 | 0.478936 | 0.043498 | 0 | 0.523096 | 0 | 0.680309 | 1.07E-63 | 0.896515 | 2.83E-08 | 1.105563 | 0.000165054 |
| Y63L N69E V74W N81S L89C | 1 | 0.478532 | 0.047416 | 1.96E-161 | 0.381232 | 1.51E-155 | 1.35451 | 9.88E-15 | 1.478711 | 4.83E-18 | 0.634001 | 6.88E-20 |
| Y63L V74E L89T | 1 | 0.477334 | 0.045323 | 0 | 0.54332 | 4.95E-308 | 0.834224 | 6.62E-14 | 1.146173 | 3.75E-08 | 1.071904 | 0.000134118 |
| V74W N81S L89C A122D | 1 | 0.476775 | 0.035561 | 2.95E-112 | 0.530355 | 2.87E-89 | 1.20247 | 2.90E-06 | 1.016032 | 0.933235 | 1.270077 | 0.00025081 |

| | | | | | | | | | | | | |
|---|---|---|---|---|---|---|---|---|---|---|---|---|
| V74W N81S A122H V196P | 1 | 0.476225 | 0.038095 | 2.25E-57 | 0.482989 | 9.13E-64 | 1.260792 | 1.90E-07 | 1.18918 | 5.94E-05 | 0.892698 | 0.002365228 |
| V74W L89A A122H V196P | 1 | 0.475259 | 0.036734 | 1.99E-125 | 0.588484 | 1.14E-79 | 1.114517 | 0.001835 | 1.142797 | 0.000914 | 1.297513 | 6.91E-09 |
| Y63L V74W N81S L89T A122D V196P | 1 | 0.473065 | 0.04893 | 1.34E-99 | 0.698916 | 2.34E-44 | 0.786284 | 0.001321 | 0.768922 | 2.95E-06 | 1.684563 | 4.47E-30 |
| N81S A122H V196P | 1 | 0.472028 | 0.054355 | 5.00E-70 | 0.606587 | 2.21E-75 | 1.19532 | 5.61E-05 | 1.28738 | 1.75E-06 | 1.325406 | 8.98E-06 |
| Y63L V74W L89T A122D V196P | 1 | 0.471023 | 0.064324 | 5.79E-207 | 0.594384 | 3.62E-137 | 0.661609 | 6.84E-15 | 0.60292 | 2.50E-19 | 1.369767 | 3.64E-15 |
| N69E N81S L89G V196P | 1 | 0.470514 | 0.050386 | 0 | 0.620376 | 4.46E-258 | 0.839861 | 1.03E-19 | 0.892436 | 3.20E-06 | 1.353987 | 4.30E-37 |
| N81S L89G A122D V196P | 1 | 0.469752 | 0.050304 | 0 | 0.609773 | 1.64E-196 | 0.914336 | 0.006138 | 0.789004 | 1.37E-13 | 1.328507 | 2.97E-30 |
| N81S L89A A122H | 1 | 0.46971 | 0.046541 | 0 | 0.525427 | 0 | 0.909134 | 1.46E-07 | 1.083383 | 1.85E-07 | 1.346801 | 2.79E-76 |
| V74W L89T A122D | 1 | 0.46849 | 0.044483 | 0 | 0.58958 | 0 | 0.842269 | 5.21E-45 | 0.969759 | 0.007014 | 1.582089 | 5.03E-197 |
| V74W R78H N81S L89A V196P | 1 | 0.468159 | 0.03745 | 8.81E-140 | 0.545772 | 1.41E-114 | 1.370004 | 5.94E-15 | 1.47105 | 3.28E-15 | 1.283887 | 8.40E-08 |
| R78H L89A V196P | 1 | 0.467982 | 0.037436 | 3.55E-209 | 0.52099 | 8.84E-167 | 1.474351 | 3.25E-17 | 2.137922 | 1.03E-48 | 1.142384 | 0.002699292 |
| V74E N81S L89C A122H V196P | 1 | 0.467979 | 0.064575 | 1.88E-275 | 0.646948 | 6.62E-197 | 0.966698 | 0.610021 | 0.901694 | 0.000564 | 1.473254 | 2.31E-50 |
| N69E L89T A122H V196P | 1 | 0.466942 | 0.033596 | 4.49E-228 | 0.540424 | 2.73E-135 | 0.856717 | 0.001043 | 0.770699 | 2.03E-09 | 1.403666 | 2.79E-23 |
| N69E R78H L89T A122D V196P | 1 | 0.465463 | 0.034449 | 4.62E-85 | 0.482832 | 6.35E-96 | 1.508657 | 6.80E-22 | 1.512941 | 2.21E-12 | 0.973749 | 0.217265313 |
| N69E V74E N81S L89A V196P | 1 | 0.465402 | 0.032963 | 8.34E-127 | 0.571529 | 8.45E-77 | 0.901996 | 0.38576 | 1.097163 | 0.07331 | 1.367628 | 1.95E-18 |
| N69E V74W N81S L89C | 1 | 0.464954 | 0.049454 | 1.65E-154 | 0.645002 | 4.91E-93 | 1.334854 | 3.12E-15 | 1.323524 | 1.83E-15 | 1.610509 | 1.60E-15 |
| V74W R78H N81S L89G V196P | 1 | 0.464467 | 0.049734 | 5.21E-263 | 0.623363 | 3.74E-132 | 0.912933 | 0.18171 | 1.038831 | 0.65575 | 1.567179 | 7.14E-39 |
| N69E V74W N81S L89T V196P | 1 | 0.463285 | 0.037695 | 4.20E-58 | 0.561024 | 5.51E-43 | 0.794753 | 0.014528 | 1.26647 | 0.48986 | 1.296337 | 0.047527732 |
| V74W L89A V196P | 1 | 0.461685 | 0.042364 | 500558052 | 0.497326 | 269452537 | 0.915798 | 0.000116 | 1.042371 | 0.277512 | 1.10066 | 0.408791369 |
| Y63L V74E R78H N81S L89C V196P | 1 | 0.460523 | 0.045631 | 7.19E-180 | 0.426356 | 5.70E-172 | 1.534426 | 1.79E-41 | 1.539468 | 5.07E-34 | 0.762719 | 1.85E-17 |
| V74E L89C A122H V196P | 1 | 0.460248 | 0.050805 | 0 | 0.694452 | 8.52E-224 | 0.707909 | 3.94E-102 | 0.578927 | 1.26E-184 | 1.967928 | 5.25E-147 |
| Y63L V74W L89T A122H V196P | 1 | 0.459867 | 0.041766 | 0 | 0.552003 | 0 | 0.941826 | 4.96E-05 | 0.943803 | 5.69E-05 | 1.251704 | 1.11E-23 |
| R78H N81S L89C | 1 | 0.45942 | 0.046235 | 7.22E-74 | 0.649974 | 9.09E-52 | 1.232285 | 0.014268 | 1.624388 | 2.64E-15 | 1.427278 | 6.43E-12 |
| Y63L N69E V74W L89A A122H V196P | 1 | 0.459027 | 0.055464 | 1.60E-208 | 0.581577 | 1.04E-125 | 1.083139 | 0.045654 | 1.039608 | 0.020624 | 1.352719 | 1.59E-13 |
| Y63L R78H N81S L89A A122D | 1 | 0.458694 | 0.035761 | 3.88E-83 | 0.469046 | 4.19E-78 | 0.626113 | 5.08E-11 | 1.063111 | 0.236441 | 0.917613 | 0.010419205 |
| N69E R78H N81S L89G V196P | 1 | 0.45862 | 0.049112 | 4.13E-295 | 0.595649 | 6.91E-162 | 0.758199 | 2.58E-16 | 1.107407 | 0.001824 | 1.390598 | 8.93E-29 |
| Y63L N69E V74W L89T A122D | 1 | 0.458583 | 0.043542 | 0.00E+00 | 0.516499 | 6.17E-243 | 0.894613 | 1.46E-06 | 1.027168 | 0.313567 | 1.027083 | 0.088091539 |
| Y63L N69E V74E N81S L89T V196P | 1 | 0.456804 | 0.037168 | 1.98E-260 | 0.613486 | 4.52E-123 | 1.125089 | 1.13E-05 | 1.229175 | 0.002182 | 1.586015 | 2.55E-44 |
| N69E V74W L89A A122D | 1 | 0.456263 | 0.04668 | 1.54E-245 | 0.67078 | 3.34E-90 | 0.840013 | 6.11E-09 | 0.60534 | 1.65E-36 | 1.971242 | 4.52E-73 |
| Y63L N69E V74W R78H N81S L89C V196P | 1 | 0.455953 | 0.040447 | 6.65E-97 | 0.59723 | 5.34E-68 | 1.385325 | 1.83E-08 | 1.581566 | 1.38E-21 | 1.305488 | 3.69E-07 |
| V74W N81S L89T | 1 | 0.455014 | 0.035169 | 1.01E-237 | 0.560011 | 1.87E-184 | 0.993129 | 0.262025 | 0.980048 | 0.158974 | 1.262857 | 2.58E-13 |
| Y63L N69E V74W N81S L89A | 1 | 0.454862 | 0.032726 | 1.26E-188 | 0.454762 | 5.01E-179 | 1.223726 | 6.31E-10 | 2.150868 | 2.01E-59 | 0.964072 | 0.033442253 |
| N81S L89A A122D V196P | 1 | 0.454648 | 0.051582 | 3.24E-41 | 0.5486 | 3.69E-48 | 0.922479 | 0.496987 | 0.8293 | 0.223255 | 1.351872 | 9.71E-05 |
| Y63L V74E N81S L89A V196P | 1 | 0.454053 | 0.035399 | 3.20E-126 | 0.506142 | 4.35E-104 | 0.968916 | 0.718725 | 0.859583 | 6.41E-07 | 1.156851 | 0.00071342 |
| N69E V74E N81S L89G A122H | 1 | 0.453389 | 0.05144 | 6.58E-45 | 0.6124 | 2.74E-36 | 0.87795 | 0.084263 | 0.966139 | 0.445365 | 1.182236 | 5.17E-05 |
| V74W L89T A122H V196P | 1 | 0.45208 | 0.041059 | 0 | 0.593022 | 0 | 0.72683 | 8.23E-74 | 0.63258 | 2.26E-121 | 1.493489 | 4.41E-93 |
| Y63L V74E L89A A122D V196P | 1 | 0.451575 | 0.0462 | 3.41E-112 | 0.567378 | 9.13E-90 | 1.052584 | 0.007649 | 1.092461 | 0.046713 | 1.503655 | 2.11E-06 |
| N69E N81S L89G A122D V196P | 1 | 0.450568 | 0.048245 | 1.81E-57 | 0.536769 | 2.21E-51 | 0.617554 | 1.61E-10 | 0.722991 | 5.61E-05 | 1.211113 | 0.019570454 |
| N69E V74W N81S L89T A122H V196P | 1 | 0.450059 | 0.047296 | 5.07E-242 | 0.463931 | 1.15E-251 | 1.29854 | 4.99E-19 | 1.059816 | 0.017224 | 1.062791 | 0.721734076 |
| V74W R78H N81S L89A A122D V196P | 1 | 0.449375 | 0.051767 | 7.49E-56 | 0.550221 | 2.12E-52 | 0.925179 | 0.572132 | 1.071992 | 0.764947 | 0.89572 | 0.544699328 |
| Y63L N69E V74E L89A | 1 | 0.448986 | 0.046515 | 0 | 0.537013 | 0 | 1.113372 | 3.25E-07 | 1.521738 | 1.69E-58 | 0.923083 | 6.33E-09 |
| Y63L N69E V74E N81S L89A | 1 | 0.445543 | 0.031713 | 3.24E-275 | 0.529999 | 1.08E-222 | 0.912391 | 0.051135 | 1.565562 | 4.78E-21 | 1.273227 | 2.77E-25 |
| N69E V74E R78H N81S L89T A122H | 1 | 0.445515 | 0.034661 | 1.78E-261 | 0.573003 | 1.54E-172 | 1.316145 | 9.79E-16 | 2.099821 | 6.32E-70 | 1.373463 | 4.92E-33 |
| N69E V74E A122H V196P | 1 | 0.445267 | 0.030414 | 5.99E-32 | 0.473503 | 5.91E-43 | 2.28375 | 7.32E-12 | 1.553254 | 0.054621 | 0.967563 | 0.586481175 |
| R78H N81S L89T | 1 | 0.444962 | 0.046098 | 0 | 0.559484 | 0 | 0.776227 | 2.28E-54 | 0.880631 | 2.02E-09 | 1.065458 | 0.000946352 |
| N69E R78H N81S L89G A122H | 1 | 0.44495 | 0.050391 | 7.83E-68 | 0.570673 | 5.93E-76 | 0.970858 | 0.617603 | 0.96956 | 0.651295 | 1.228412 | 0.000361975 |
| V74E N81S L89T A122D V196P | 1 | 0.444421 | 0.04748 | 2.76E-132 | 0.585631 | 6.62E-115 | 1.295839 | 2.67E-08 | 1.297214 | 7.22E-05 | 1.579242 | 1.27E-27 |
| N69E V74E N81S L89A A122D | 1 | 0.443355 | 0.061178 | 5.18E-32 | 0.549799 | 1.41E-58 | 0.687839 | 1.98E-05 | 1.114968 | 0.826898 | 1.443151 | 2.91E-05 |

| | | | | | | | | | | | | |
|---|---|---|---|---|---|---|---|---|---|---|---|---|
| Y63L N69E V74W N81S L89T V196P | 1 | 0.442803 | 0.04046 | 1.08E-112 | 0.597248 | 2.52E-56 | 0.945693 | 0.806155 | 1.353925 | 2.40E-10 | 1.572618 | 5.15E-14 |
| Y63L V74E N81S L89T A122D V196P | 1 | 0.441362 | 0.047153 | 4.02E-133 | 0.658966 | 2.36E-77 | 1.027348 | 0.175189 | 1.058815 | 0.283332 | 1.819781 | 8.26E-40 |
| N69E V74W R78H N81S L89A A122H V196P | 1 | 0.44019 | 0.050709 | 3.63E-268 | 0.715533 | 5.01E-113 | 0.837241 | 2.08E-19 | 0.901984 | 6.38E-12 | 2.036472 | 1.70E-93 |
| Y63L N69E V74W N81S L89T | 1 | 0.439865 | 0.048555 | 4.93E-231 | 0.599786 | 7.40E-143 | 0.687794 | 2.00E-33 | 0.742227 | 7.73E-23 | 1.544605 | 7.16E-16 |
| N69E N81S L89C A122H V196P | 1 | 0.439656 | 0.035026 | 2.84E-83 | 0.45519 | 1.91E-78 | 1.522863 | 8.93E-11 | 1.080341 | 0.207039 | 0.883031 | 6.74E-05 |
| V74W L89T A122H | 1 | 0.436971 | 0.04149 | 2.29E-277 | 0.520946 | 4.79E-214 | 0.78449 | 2.14E-15 | 1.151678 | 0.000672 | 1.090086 | 0.000108691 |
| R78H A122H | 1 | 0.436834 | 0.038751 | 1.13E-147 | 0.5444 | 3.50E-123 | 2.839292 | 3.02E-73 | 1.614644 | 2.14E-23 | 1.424815 | 5.05E-21 |
| N69E R78H N81S L89C | 1 | 0.435109 | 0.06004 | 2.64E-261 | 0.559754 | 1.23E-290 | 1.339527 | 3.13E-32 | 0.904097 | 2.74E-05 | 1.588062 | 4.08E-51 |
| N69E N81S L89A A122D V196P | 1 | 0.434663 | 0.031463 | 1.28E-105 | 0.508465 | 8.21E-83 | 0.900468 | 0.770675 | 0.893977 | 0.05794 | 1.448333 | 2.39E-12 |
| N69E V74E N81S L89T A122H V196P | 1 | 0.434324 | 0.031727 | 6.21E-133 | 0.582709 | 6.34E-100 | 1.002483 | 0.003002 | 1.140767 | 0.137378 | 1.611116 | 9.62E-24 |
| Y63L N69E R78H N81S L89T | 1 | 0.434251 | 0.044989 | 1087646913 | 0.546849 | 1.38E-279 | 0.842021 | 4.06E-12 | 0.749441 | 3.06E-21 | 0.918253 | 4.82E-06 |
| V74W R78H N81S L89A A122H | 1 | 0.432378 | 0.03225 | 1.91E-205 | 0.545111 | 2.02E-141 | 0.719659 | 9.69E-17 | 0.927758 | 0.413233 | 1.505237 | 1.26E-39 |
| Y63L N69E R78H N81S L89T V196P | 1 | 0.432036 | 0.03453 | 5.33E-39 | 0.519536 | 8.39E-42 | 1.803522 | 2.48E-17 | 1.352357 | 0.000189 | 1.192933 | 0.205749127 |
| Y63L V74W R78H N81S A122H | 1 | 0.431499 | 0.04463 | 2.21E-63 | 0.645631 | 4.12E-31 | 1.185688 | 2.85E-05 | 1.104891 | 0.512204 | 1.483613 | 1.82E-06 |
| Y63L N69E V74W L89A A122D | 1 | 0.428629 | 0.038882 | 3.92E-54 | 0.494883 | 9.34E-49 | 1.208182 | 1.15E-05 | 0.929904 | 0.608455 | 0.815435 | 0.000406319 |
| N69E V74W N81S L89C V196P | 1 | 0.427547 | 0.042364 | 0 | 0.440532 | 2.54E-301 | 1.188373 | 4.46E-15 | 1.368357 | 2.73E-28 | 0.964534 | 0.012912748 |
| Y63L N69E L89T A122H V196P | 1 | 0.422858 | 0.038405 | 0 | 0.592937 | 1.31E-234 | 0.696109 | 7.40E-75 | 0.697476 | 1.33E-66 | 1.569403 | 9.43E-77 |
| Y63L N69E V74E R78H N81S L89C A122D V196P | 1 | 0.422512 | 0.065719 | 5.38E-97 | 0.407202 | 2.31E-116 | 1.10586 | 0.111554 | 1.591927 | 1.57E-24 | 0.706562 | 4.33E-08 |
| V74W R78H N81S L89T V196P | 1 | 0.422242 | 0.042493 | 4.76E-295 | 0.775681 | 5.61E-69 | 0.992269 | 0.222646 | 1.338166 | 7.16E-24 | 2.486707 | 7.35E-196 |
| Y63L V74W L89A A122D V196P | 1 | 0.422175 | 0.041164 | 1.94E-34 | 0.524513 | 1.51E-36 | 0.890538 | 0.303553 | 1.994533 | 3.83E-06 | 1.049243 | 0.858287573 |
| N69E V74W L89T A122H | 1 | 0.419642 | 0.039674 | 2.08E-85 | 0.560314 | 1.63E-54 | 0.95164 | 0.92238 | 0.690868 | 1.74E-11 | 1.309482 | 4.01E-09 |
| N69E V74W N81S L89A A122D | 1 | 0.419179 | 0.027486 | 2.13E-61 | 0.483565 | 2.83E-51 | 1.467564 | 9.01E-07 | 0.875774 | 0.225103 | 1.372533 | 1.99E-09 |
| N69E V74W N81S L89A V196P | 1 | 0.419064 | 0.029681 | 4.54E-38 | 0.466101 | 1.39E-43 | 1.425427 | 0.004954 | 1.609045 | 0.002595 | 1.029354 | 0.732630037 |
| N69E V74E L89A A122D V196P | 1 | 0.418719 | 0.042604 | 1.72E-298 | 0.540604 | 9.93E-225 | 1.13623 | 6.03E-10 | 1.362769 | 6.35E-20 | 1.175765 | 1.19E-12 |
| N69E V74E L89T A122D V196P | 1 | 0.418608 | 0.031552 | 5.01E-75 | 0.490715 | 9.09E-76 | 1.260048 | 8.16E-05 | 1.523434 | 8.97E-08 | 1.118147 | 0.008544859 |
| N69E V74E L89A A122D | 1 | 0.417078 | 0.038271 | 4.44E-58 | 0.408785 | 2.14E-79 | 1.09885 | 0.003664 | 1.089872 | 0.096928 | 0.801352 | 0.000135976 |
| Y63L N69E V74E L89T V196P | 1 | 0.41507 | 0.039241 | 4.50E-116 | 0.501912 | 8.66E-112 | 1.034955 | 0.140488 | 1.225098 | 4.24E-06 | 1.038195 | 0.149934106 |
| V74W N81S L89T A122D V196P | 1 | 0.412176 | 0.042632 | 4.27E-60 | 0.594737 | 9.31E-46 | 0.948918 | 0.697738 | 1.063944 | 0.988559 | 1.375961 | 4.33E-09 |
| N69E R78H N81S L89G | 1 | 0.412065 | 0.05686 | 1.26E-80 | 0.536076 | 4.78E-99 | 1.042627 | 0.010017 | 0.816911 | 0.000197 | 1.35363 | 2.33E-10 |
| V74E N81S L89A A122D | 1 | 0.411681 | 0.029158 | 8.48E-55 | 0.518705 | 5.30E-36 | 0.999317 | 0.195534 | 1.311679 | 0.144159 | 1.349359 | 6.92E-07 |
| V74E N81S L89G A122D V196P | 1 | 0.4113 | 0.044041 | 5.11E-55 | 0.530619 | 2.02E-55 | 1.479473 | 1.49E-15 | 0.984193 | 0.50229 | 1.198231 | 0.057059035 |
| N69E V74W R78H L89A A122D V196P | 1 | 0.410936 | 0.040068 | 1.89E-106 | 0.61583 | 8.91E-68 | 1.437503 | 5.50E-17 | 1.976641 | 2.01E-33 | 1.354988 | 7.95E-09 |
| N69E V74W R78H L89A A122H V196P | 1 | 0.410651 | 0.048488 | 0 | 0.524153 | 1.16E-288 | 0.69898 | 2.17E-37 | 0.893439 | 0.020133 | 1.552391 | 4.78E-61 |
| Y63L V74E R78H N81S L89C A122H | 1 | 0.406758 | 0.040304 | 3.25E-200 | 0.429207 | 3.84E-170 | 1.095557 | 0.000868 | 1.63145 | 2.25E-30 | 0.970993 | 0.324151824 |
| Y63L V74W L89A A122H V196P | 1 | 0.405656 | 0.049404 | 0 | 0.461347 | 0 | 0.83652 | 5.26E-26 | 0.74206 | 1.77E-37 | 1.192636 | 4.45E-09 |
| Y63L V74W R78H N81S L89C A122D | 1 | 0.404657 | 0.040096 | 3.00E-223 | 0.404054 | 2.01E-213 | 2.162266 | 3.50E-109 | 1.924669 | 4.97E-65 | 0.829706 | 1.30E-13 |
| Y63L N69E V74E N81S L89T A122D | 1 | 0.404424 | 0.044643 | 0 | 0.585219 | 1.90E-201 | 0.831371 | 4.10E-12 | 1.168779 | 2.64E-07 | 1.587681 | 1.30E-39 |
| V74E L89T A122H V196P | 1 | 0.404 | 0.029067 | 8.99E-235 | 0.440197 | 2.23E-197 | 1.235561 | 9.49E-08 | 1.763653 | 2.93E-42 | 1.064689 | 0.019072022 |
| N69E V74W R78H N81S L89T V196P | 1 | 0.404 | 0.036915 | 0 | 0.549847 | 0 | 1.36283 | 1.76E-65 | 1.595852 | 3.98E-92 | 1.80917 | 3.26E-196 |
| N69E V74W R78H N81S L89A A122D V196P | 1 | 0.40392 | 0.028609 | 1.81E-48 | 0.452109 | 1.14E-49 | 1.675381 | 2.46E-07 | 1.771165 | 5.74E-07 | 1.082235 | 0.90584963 |
| Y63L N69E V74E N81S L89G A122H V196P | 1 | 0.403562 | 0.048762 | 1.26E-38 | 0.569531 | 7.61E-29 | 0.623998 | 5.50E-06 | 0.93307 | 0.585275 | 1.585091 | 4.03E-05 |
| V74E N81S A122H V196P | 1 | 0.399532 | 0.044103 | 2.58E-217 | 0.54104 | 1.67E-177 | 1.107381 | 5.50E-05 | 1.288583 | 3.89E-15 | 1.211138 | 0.007956549 |
| N69E V74W R78H N81S L89G A122D | 1 | 0.399011 | 0.056372 | 0 | 0.458937 | 553441792 | 0.902759 | 4.12E-09 | 0.739856 | 4.07E-27 | 0.94967 | 0.000272857 |
| N81S L89C A122H V196P | 1 | 0.398436 | 0.031742 | 6.03E-67 | 0.439628 | 7.32E-56 | 1.175738 | 0.007062 | 0.869064 | 0.081116 | 0.927074 | 0.450739712 |
| R78H N81S L89C A122H | 1 | 0.398399 | 0.029078 | 0 | 0.458153 | 0 | 1.327893 | 7.13E-25 | 1.815977 | 1.64E-47 | 1.133913 | 6.39E-12 |
| N69E R78H L89T | 1 | 0.397389 | 0.042268 | 2.65E-84 | 0.661843 | 4.10E-34 | 0.835268 | 0.000194 | 1.175976 | 0.000766 | 2.945687 | 1.59E-24 |
| V74E R78H N81S L89T V196P | 1 | 0.397349 | 0.040475 | 1.66E-26 | 0.479264 | 6.48E-41 | 1.173769 | 0.064155 | 1.354195 | 5.06E-06 | 1.728129 | 5.05E-13 |
| Y63L R78H N81S L89T A122H | 1 | 0.395011 | 0.036093 | 2.03E-51 | 0.487111 | 1.19E-46 | 1.194178 | 0.01821 | 1.762336 | 4.66E-12 | 1.282389 | 0.005801413 |

| | | | | | | | | | | | | |
|---|---|---|---|---|---|---|---|---|---|---|---|---|
| N69E N81S L89A A122H | 1 | 0.394649 | 0.029006 | 0 | 0.53865 | 9.27E-294 | 1.348761 | 2.20E-34 | 1.671432 | 2.87E-61 | 1.841858 | 1.12E-114 |
| Y63L V74W R78H N81S L89C V196P | 1 | 0.393706 | 0.039011 | 0 | 0.440156 | 5.07E-291 | 1.466196 | 5.62E-42 | 1.697443 | 2.02E-70 | 1.161207 | 2.18E-06 |
| N69E V74E N81S L89C A122D V196P | 1 | 0.393243 | 0.039636 | 7.73E-126 | 0.494788 | 7.39E-110 | 1.293767 | 1.28E-06 | 1.147018 | 0.010333 | 1.19794 | 0.000302733 |
| N69E V74W L89A A122H V196P | 1 | 0.392864 | 0.04747 | 1.60E-112 | 0.509441 | 3.55E-93 | 1.150378 | 0.000311 | 0.823018 | 9.53E-06 | 1.242794 | 2.72E-07 |
| Y63L N69E V74W N81S L89A A122D V196P | 1 | 0.39276 | 0.027818 | 3.28E-70 | 0.466794 | 1.28E-61 | 1.252454 | 0.000476 | 1.136432 | 0.311872 | 1.250103 | 4.50E-08 |
| V74E R78H N81S L89T A122D | 1 | 0.392632 | 0.04636 | 0 | 0.487591 | 3.35E-274 | 0.966935 | 0.245138 | 1.209848 | 0.000702 | 1.30094 | 1.10E-17 |
| Y63L N69E N81S L89C A122H V196P | 1 | 0.389883 | 0.054037 | 3.23E-81 | 0.477356 | 3.50E-122 | 0.99713 | 0.994866 | 1.034713 | 0.921421 | 0.941605 | 0.946634873 |
| Y63L N69E V74W N81S L89G V196P | 1 | 0.389871 | 0.041746 | 3.66E-42 | 0.519027 | 3.81E-46 | 1.436616 | 1.18E-05 | 1.040889 | 0.364021 | 1.195646 | 0.093585255 |
| N69E V74W N81S L89G A122D V196P | 1 | 0.389102 | 0.041668 | 6.60E-143 | 0.527952 | 5.69E-114 | 0.711817 | 4.77E-11 | 0.778463 | 2.45E-05 | 1.387723 | 5.42E-12 |
| N69E V74W N81S L89C A122D | 1 | 0.38697 | 0.028244 | 8.90E-167 | 0.444996 | 3.82E-143 | 1.382411 | 6.21E-11 | 1.330205 | 2.51E-06 | 1.050058 | 0.067183538 |
| V74W L89A A122H | 1 | 0.386281 | 0.039304 | 8.14E-173 | 0.506545 | 1.29E-146 | 0.805375 | 4.41E-08 | 0.764784 | 4.07E-09 | 1.30335 | 3.87E-09 |
| Y63L V74W L89T V196P | 1 | 0.386019 | 0.036652 | 0 | 0.455192 | 5.18E-300 | 0.977002 | 0.547871 | 1.300416 | 1.80E-16 | 1.136616 | 1.08E-11 |
| N69E V74E R78H N81S L89T A122H V196P | 1 | 0.385931 | 0.028192 | 3.70E-148 | 0.646647 | 2.02E-66 | 0.658998 | 4.60E-14 | 1.128227 | 0.099397 | 2.320941 | 1.78E-65 |
| V74W N81S L89G | 1 | 0.385832 | 0.030738 | 2.17E-66 | 0.417948 | 1.82E-62 | 1.429397 | 2.02E-06 | 1.244745 | 0.00205 | 0.784755 | 0.000750202 |
| Y63L R78H L89T | 1 | 0.38486 | 0.036385 | 1.30E-58 | 0.442188 | 7.36E-52 | 1.204513 | 0.000134 | 1.521859 | 0.001371 | 1.083561 | 0.026869366 |
| V74E R78H N81S L89T | 1 | 0.383361 | 0.042317 | 0 | 0.599131 | 2.16E-221 | 0.794387 | 3.43E-27 | 1.106476 | 0.002884 | 2.096037 | 5.78E-125 |
| V74E R78H N81S L89A V196P | 1 | 0.38331 | 0.027149 | 1.67E-67 | 0.515776 | 5.28E-49 | 1.371266 | 8.49E-05 | 1.521175 | 0.003462 | 1.859619 | 5.12E-31 |
| N69E R78H N81S L89A V196P | 1 | 0.383145 | 0.029871 | 4.13E-209 | 0.455082 | 1.04E-164 | 1.289627 | 1.77E-18 | 1.394282 | 2.23E-09 | 1.401614 | 2.23E-25 |
| N69E L89A A122H V196P | 1 | 0.383131 | 0.029233 | 1.24E-48 | 0.592254 | 3.38E-39 | 1.419039 | 0.00011 | 1.34718 | 0.265725 | 1.631026 | 1.27E-08 |
| V74E R78H L89A | 1 | 0.382285 | 0.040694 | 4.47E-184 | 0.446839 | 1.20E-151 | 0.984966 | 0.657536 | 1.886279 | 3.23E-24 | 1.100775 | 0.011825644 |
| R78H N81S L89A V196P | 1 | 0.380897 | 0.034552 | 2.67E-90 | 0.531526 | 4.49E-68 | 1.029208 | 0.117583 | 1.534038 | 4.21E-09 | 1.275424 | 1.47E-05 |
| V74W R78H L89T A122D | 1 | 0.380026 | 0.028126 | 6.36E-127 | 0.462979 | 8.54E-126 | 1.547193 | 6.25E-18 | 2.949752 | 2.97E-47 | 1.238057 | 2.92E-08 |
| N69E V74W L89A A122H | 1 | 0.379976 | 0.039366 | 2.76E-293 | 0.504165 | 4.43E-276 | 1.004827 | 0.29325 | 0.773105 | 8.10E-13 | 1.113644 | 0.015692124 |
| N69E V74E R78H N81S L89T A122D | 1 | 0.379725 | 0.046246 | 0 | 0.452324 | 0 | 0.961122 | 0.027817 | 0.89741 | 1.69E-08 | 1.206761 | 1.19E-15 |
| N69E V74W L89A | 1 | 0.379721 | 0.041916 | 0 | 0.614064 | 1.96E-214 | 0.810588 | 7.03E-28 | 0.933395 | 0.011957 | 2.035987 | 5.18E-112 |
| N69E V74W N81S L89C A122H | 1 | 0.378771 | 0.042974 | 1.16E-50 | 0.532756 | 1.25E-46 | 0.781476 | 0.002756 | 1.928912 | 3.25E-09 | 1.79207 | 3.58E-16 |
| V74E N81S L89A A122D V196P | 1 | 0.37688 | 0.024712 | 1.04E-54 | 0.461572 | 3.14E-47 | 2.055224 | 6.78E-15 | 1.24736 | 0.967825 | 1.426688 | 1.84E-07 |
| N69E V74E N81S L89G A122D V196P | 1 | 0.372482 | 0.048916 | 1.03E-48 | 0.55064 | 2.70E-38 | 0.977248 | 0.82491 | 0.74099 | 0.000173 | 1.210501 | 0.004126145 |
| R78H L89T A122H | 1 | 0.371262 | 0.027983 | 2.25E-108 | 0.391637 | 4.16E-121 | 1.838067 | 4.86E-15 | 2.684327 | 1.72E-20 | 0.960743 | 0.768067832 |
| R78H L89A | 1 | 0.370127 | 0.045077 | 0 | 0.472718 | 0 | 0.872856 | 2.66E-16 | 1.118518 | 3.61E-15 | 1.388536 | 3.13E-59 |
| N69E V74E N81S L89T A122H | 1 | 0.368449 | 0.033423 | 1.46E-181 | 0.416915 | 2.82E-177 | 1.479538 | 7.47E-21 | 1.883766 | 1.15E-21 | 1.107046 | 5.21E-05 |
| N69E V74W R78H L89A A122H | 1 | 0.361371 | 0.038468 | 8.47E-259 | 0.450342 | 1.10E-225 | 1.051396 | 0.000622 | 1.239927 | 6.17E-05 | 1.232765 | 5.27E-09 |
| V74W L89A | 1 | 0.360706 | 0.038397 | 3.85E-103 | 0.497945 | 4.48E-81 | 0.855258 | 0.153019 | 0.896988 | 0.035976 | 1.303147 | 2.33E-07 |
| N69E R78H N81S L89A A122H V196P | 1 | 0.360614 | 0.025541 | 2.29E-147 | 0.437403 | 6.99E-132 | 1.557625 | 2.85E-12 | 1.866963 | 9.83E-14 | 1.415461 | 2.20E-23 |
| L89A A122H V196P | 1 | 0.359723 | 0.043465 | 0 | 0.514044 | 0 | 1.31055 | 1.92E-52 | 1.202704 | 3.26E-23 | 1.689335 | 1.99E-111 |
| V74W N81S L89A A122D V196P | 1 | 0.359709 | 0.028044 | 3.06E-56 | 0.430897 | 3.55E-48 | 1.413034 | 1.70E-06 | 0.84028 | 0.00487 | 1.427136 | 5.69E-10 |
| N69E V74W R78H N81S L89G V196P | 1 | 0.359688 | 0.050817 | 6.07E-302 | 0.455446 | 2.47E-225 | 0.911029 | 0.011438 | 0.809976 | 5.37E-08 | 1.203041 | 0.000905951 |
| N69E V74E N81S L89A A122D V196P | 1 | 0.358506 | 0.02795 | 1.46E-95 | 0.441839 | 4.51E-88 | 1.135667 | 0.00049 | 1.394794 | 0.010729 | 1.336358 | 7.61E-12 |
| Y63L N69E V74W N81S L89A A122H | 1 | 0.358294 | 0.032541 | 1.62E-197 | 0.410218 | 3.56E-183 | 1.24651 | 1.27E-10 | 1.065308 | 0.899692 | 1.056211 | 0.461475313 |
| Y63L N69E V74E N81S L89A A122D | 1 | 0.357361 | 0.044677 | 1.95E-46 | 0.499852 | 1.22E-35 | 0.785071 | 0.081819 | 1.06233 | 0.303279 | 1.529489 | 0.002348308 |
| V74E N81S L89A A122H | 1 | 0.354762 | 0.025251 | 1.46E-98 | 0.426415 | 3.12E-99 | 1.81158 | 5.40E-29 | 2.775617 | 5.49E-42 | 1.239412 | 8.86E-09 |
| N69E V74W N81S L89A A122D V196P | 1 | 0.353953 | 0.027595 | 2.44E-174 | 0.435943 | 9.27E-153 | 1.431907 | 2.86E-20 | 1.217779 | 0.018167 | 1.426037 | 6.83E-31 |
| V74W N81S L89C A122H | 1 | 0.353036 | 0.034981 | 1.54E-176 | 0.455512 | 1.03E-138 | 0.989363 | 0.170052 | 1.029161 | 0.853933 | 1.465644 | 6.84E-21 |
| N69E V74W R78H L89T A122D V196P | 1 | 0.350856 | 0.033171 | 1.34E-125 | 0.448882 | 1.05E-103 | 1.600886 | 1.54E-17 | 1.824332 | 1.84E-18 | 1.468869 | 5.29E-15 |
| N69E V74W R78H N81S L89T | 1 | 0.350709 | 0.042712 | 0 | 0.484773 | 0 | 0.837334 | 1.15E-23 | 1.162733 | 6.94E-20 | 1.584696 | 1.13E-117 |
| N69E R78H N81S L89G A122H V196P | 1 | 0.350592 | 0.048591 | 7.16E-36 | 0.502742 | 3.21E-49 | 0.771519 | 7.89E-12 | 0.983339 | 0.234279 | 1.653517 | 3.51E-09 |
| V74W N81S L89G V196P | 1 | 0.349857 | 0.036221 | 4.06E-74 | 0.481834 | 3.07E-60 | 0.944011 | 0.69761 | 1.585033 | 0.001899 | 1.472973 | 2.90E-07 |
| N69E R78H N81S L89A A122H | 1 | 0.349361 | 0.025678 | 5.91E-179 | 0.421176 | 2.16E-173 | 1.581529 | 4.87E-25 | 2.092048 | 3.18E-29 | 1.344812 | 1.39E-16 |

| | | | | | | | | | | | |
|---|---|---|---|---|---|---|---|---|---|---|---|
| V74E N81S L89T A122H | 1 | 0.3458 | 0.031597 | 2.76E-264 | 0.532681 | 2.04E-174 | 1.129102 | 1.71E-05 | 1.289673 | 8.14E-13 | 1.977195 | 3.15E-71 |
| N69E R78H N81S L89A | 1 | 0.345177 | 0.034656 | 4.49E-108 | 0.798088 | 1.94E-21 | 0.677409 | 4.49E-13 | 0.741797 | 4.89E-07 | 4.45933 | 2.01E-83 |
| R78H L89T V196P | 1 | 0.343276 | 0.025874 | 9.43E-77 | 0.393337 | 8.05E-85 | 1.97088 | 1.61E-19 | 2.173173 | 9.91E-22 | 1.165127 | 0.001000767 |
| Y63L V74E L89T A122H V196P | 1 | 0.341489 | 0.031015 | 1.35E-201 | 0.426576 | 8.05E-165 | 1.073419 | 0.220326 | 0.940571 | 0.103379 | 1.349673 | 1.71E-15 |
| N69E V74W N81S L89A | 1 | 0.339688 | 0.02444 | 1.61E-92 | 0.374359 | 9.37E-90 | 1.186258 | 0.001088 | 2.464104 | 1.79E-29 | 1.188703 | 0.000226499 |
| R78H N81S L89A A122H | 1 | 0.338236 | 0.02486 | 2.88E-234 | 0.484199 | 2.16E-192 | 1.591874 | 5.26E-37 | 1.591514 | 7.33E-29 | 2.09999 | 8.21E-76 |
| V74W N81S L89A A122H V196P | 1 | 0.337925 | 0.034986 | 6.88E-53 | 0.412031 | 1.75E-54 | 1.963481 | 3.63E-14 | 1.407175 | 0.003141 | 1.386503 | 0.000132644 |
| Y63L N69E V74W N81S L89T A122D | 1 | 0.332359 | 0.03538 | 4.99E-160 | 0.442093 | 2.42E-141 | 1.079778 | 0.001253 | 0.839586 | 0.000378 | 1.256234 | 8.66E-08 |
| R78H N81S L89T A122H | 1 | 0.330697 | 0.026907 | 3.76E-63 | 0.436126 | 4.90E-62 | 2.358157 | 1.44E-14 | 1.546671 | 0.000549 | 1.423742 | 5.16E-06 |
| V74W R78H L89T A122D V196P | 1 | 0.328343 | 0.029821 | 1.01E-125 | 0.382847 | 8.23E-118 | 0.815899 | 0.009374 | 0.835355 | 0.02897 | 1.14887 | 0.004303563 |
| Y63L N69E V74E A122H V196P | 1 | 0.328182 | 0.051047 | 3.68E-78 | 0.420512 | 7.71E-87 | 0.896358 | 0.009963 | 0.915094 | 0.462479 | 1.081601 | 0.695421097 |
| V74W L89A A122D V196P | 1 | 0.325505 | 0.03312 | 7.39E-252 | 0.429689 | 9.25E-230 | 1.361524 | 8.79E-19 | 1.408844 | 1.80E-12 | 1.502784 | 1.29E-29 |
| Y63L V74W N81S L89G A122D | 1 | 0.325145 | 0.045937 | 9.15E-139 | 0.414711 | 5.99E-140 | 0.976449 | 0.858313 | 0.719272 | 1.80E-13 | 1.148199 | 0.456284709 |
| V74W N81S L89A V196P | 1 | 0.32509 | 0.023531 | 7.60E-57 | 0.382528 | 9.80E-62 | 1.172074 | 0.041322 | 1.628402 | 0.003092 | 1.212347 | 0.002391269 |
| N81S L89T A122H V196P | 1 | 0.324701 | 0.042642 | 2.76E-57 | 0.550676 | 1.61E-54 | 1.106597 | 0.005721 | 1.021351 | 0.002044 | 1.606518 | 7.56E-13 |
| Y63L N69E V74W N81S L89G A122D | 1 | 0.323139 | 0.045653 | 2.09E-154 | 0.415659 | 2.14E-132 | 0.831417 | 6.28E-05 | 0.674858 | 7.93E-17 | 1.242016 | 0.002560396 |
| N69E V74W R78H N81S L89A | 1 | 0.323138 | 0.029348 | 3.61E-245 | 0.41094 | 4.82E-213 | 0.896331 | 0.025977 | 0.836056 | 5.50E-06 | 1.429925 | 7.22E-19 |
| R78H L89A A122H V196P | 1 | 0.315683 | 0.037274 | 7.09E-139 | 0.450654 | 2.26E-124 | 1.304166 | 8.28E-12 | 0.886685 | 0.004959 | 1.379713 | 1.64E-12 |
| R78H N81S L89A | 1 | 0.313765 | 0.023061 | 2.45E-127 | 0.432347 | 4.36E-110 | 1.386607 | 3.03E-10 | 2.041993 | 5.29E-19 | 1.841224 | 8.55E-32 |
| Y63L N69E V74W N81S L89G A122D V196P | 1 | 0.307064 | 0.032883 | 8.59E-46 | 0.398146 | 3.04E-58 | 0.741451 | 0.033178 | 1.577895 | 6.24E-05 | 1.211314 | 0.092918579 |
| V74E N81S L89G A122H V196P | 1 | 0.305737 | 0.036942 | 1.85E-66 | 0.49062 | 5.08E-57 | 1.138837 | 0.001667 | 1.204966 | 0.00209 | 1.845305 | 7.74E-11 |
| N69E V74W R78H N81S L89A A122H | 1 | 0.303643 | 0.04723 | 4.61E-78 | 0.418318 | 9.49E-88 | 0.749491 | 2.25E-11 | 0.597677 | 7.69E-31 | 1.40641 | 1.32E-05 |
| R78H N81S L89A A122H V196P | 1 | 0.300335 | 0.042431 | 0.00E+00 | 0.403357 | 1.91E-252 | 1.201434 | 6.74E-11 | 1.002571 | 0.829731 | 1.571358 | 2.84E-28 |
| N69E V74W N81S L89A A122H | 1 | 0.29749 | 0.021404 | 1.68E-132 | 0.359326 | 1.51E-132 | 2.487893 | 3.19E-27 | 2.57894 | 2.78E-51 | 1.44249 | 5.96E-18 |
| V74W R78H N81S L89A | 1 | 0.297073 | 0.026981 | 0 | 0.482892 | 8.46E-303 | 1.070538 | 0.002345 | 1.028518 | 0.561613 | 2.373599 | 2.36E-139 |
| N69E V74E L89A A122H | 1 | 0.29167 | 0.035522 | 0 | 0.409442 | 5466679935 | 0.93435 | 0.186269 | 1.114629 | 0.000451 | 1.453225 | 5.01E-27 |
| Y63L R78H N81S L89T A122H V196P | 1 | 0.289973 | 0.038081 | 5.28E-72 | 0.438299 | 2.92E-66 | 1.155696 | 0.003149 | 2.320649 | 3.34E-24 | 1.721276 | 2.43E-15 |
| V74W N81S L89A | 1 | 0.27065 | 0.024581 | 2.67E-117 | 0.35997 | 6.10E-112 | 1.069066 | 0.525627 | 1.409486 | 0.000163 | 1.565275 | 2.15E-14 |
| V74W N81S L89G A122D | 1 | 0.269474 | 0.038071 | 4.47E-147 | 0.423767 | 2.97E-119 | 0.976768 | 0.263053 | 0.866989 | 4.75E-05 | 1.853192 | 5.34E-22 |
| Y63L N81S L89A A122H V196P | 1 | 0.265713 | 0.03754 | 7.52E-53 | 0.313646 | 1.35E-48 | 1.363615 | 0.003288 | 0.831129 | 0.003044 | 1.137941 | 0.120770353 |
| R78H L89A A122H | 1 | 0.246833 | 0.026276 | 1.16E-89 | 0.34734 | 4.05E-89 | 3.245092 | 2.60E-31 | 2.264578 | 1.96E-17 | 1.319216 | 1.31E-06 |
| Y63L N69E V74W R78H N81S L89T | 1 | 0.241881 | 0.029458 | 1.72E-82 | 0.352592 | 1.95E-76 | 1.355002 | 1.39E-06 | 2.407961 | 4.07E-25 | 1.554389 | 9.88E-11 |
| N69E V74E L89T A122H | 1 | 0.236778 | 0.036829 | 2.95E-94 | 0.445543 | 7.65E-76 | 0.819962 | 4.76E-06 | 0.696568 | 3.68E-15 | 2.10009 | 4.94E-18 |
| Y63L N69E V74E L89T A122H | 1 | 0.191408 | 0.029772 | 3.24E-129 | 0.393236 | 6.77E-124 | 0.735524 | 1.21E-11 | 0.536718 | 6.02E-39 | 2.830423 | 8.09E-54 |
