## Supplementary Table 3 for "Sensitivity optimization of a rhodopsin-based fluorescent voltage indicator"

| Number of mutations | Total variants | Variants with A122D | % |
|---|---|---|---|
| 2 | 64 | 10 | 15.6 |
| 3 | 167 | 43 | 25.7 |
| 4 | 247 | 88 | 35.6 |
| 5 | 219 | 86 | 39.3 |
| 6 | 118 | 52 | 44.1 |
| 7 | 30 | 11 | 36.7 |
| 8 | 3 | 2 | 66.7 |
| **Sum** | **848** | **292** | **34.4** |
