## Supplementary Table 4 for "Sensitivity optimization of a rhodopsin-based fluorescent voltage indicator"

**Supplementary Table 4**: Custom primers used for library tagmentation and NextSeq sequencing

| primer | sequence (5'-3') |
|---|---|
| **custom read 1 sequencing primer** | GCGATCGAGGACGGCAGATGTGTATAAGAGACAG |
| **custom index 1 (i7) sequencing primer** | CTGTCTCTTATACACATCTGAGGCGGAGACGGTG |
| **custom index 2 (i5) sequencing primer** | ATCGCAGGCTATAGCGTGGAGACGCTGCCGACGA |
| **custom read 2 sequencing primer** | CACCGTCTCCGCCTCAGATGTGTATAAGAGACAG |
| **T7 well BC adapter oligo (1-96)** | GTCTCGTGGGCTCGGCTGTCCCTGTCCNNNNNNNNCACCGTCTCCGCCTCAGATGTGTATAAGAGACAG |
| **i5_1 adapter oligo** | TCGTCGGCAGCGTCTCCACGCTATAGCCTGCGATCGAGGACGGCAGATGTGTATAAGAGACAG |
| **i7_1 PCR primer** | CAAGCAGAAGACGGCATACGAGATATCCGCATGTCTCGTGGGCTCGG |
| **i5 plate BC PCR primer (1-288)** | AATGATACGGCGACCACCGAGATCTACACNNNNNNNNTCGTCGGCAGCGTC |
| **pMENTS** | CTGTCTCTTATACACATCT |
