## Supplementary methods for "Sensitivity optimization of a rhodopsin-based fluorescent voltage indicator"

### **Deep Sequencing**

#### *Tn5 Transposase Purification*

The purification protocol is derived from the methods of Picelli et al. pMXB1-Tn5-intein was transformed into BL21(DE3)pLysS (Novagen) and grown in medium containing 100 µg/ml carbenicillin and 60 µg/ml chloramphenicol (Picelli et al. 2014). One liter of culture was grown to $OD_{600}$ = 0.9 at 37°C, chilled to 23°C, induced with 0.25 mM IPTG, and grown for 16 hours at 23°C. The cell pellet was stored at -80°C after harvesting. The pellet was resuspended in 100 ml "HEGX" buffer (20 mM HEPES-KOH, 0.8 M NaCl, 1 mM EDTA, 10% glycerol, 0.2% Triton X-100, pH 7.2 at 4°C) with the addition of a cOmplete protease inhibitor cocktail (Roche), and lysed by sonication while on ice. The lysate was clarified by centrifuging at 35,000xg for 45 min at 4°C. To precipitate bacterial DNA from the clarified lysate, 2 mL of 10% polyethyleneimine (PEI) was slowly added dropwise while stirring at 4°C. The resulting turbidity was removed by centrifuging the lysate again in 50 mL conical tubes.

The re-clarified lysate was passed over a 10 mL column of chitin resin (NEB #S6651L), which binds to the intein tag, and washed with 200 mL of HEGX buffer. For elution, one column volume of 100 mM dithiothreitol (DTT) in HEGX was passed through the column, then the column was closed for at least 48 hours for DTT to cleave the intein tag at 4°C. After 48 hours, approximately 15 ml of eluate was collected from the column, concentrated to 3 mL, and dialyzed in 2x Dialysis Buffer (100 mM HEPES, 0.2 M NaCl, 0.2 mM EDTA, 2 mM DTT, 0.2% Triton X-100, 20% glycerol, pH 7.2 at 4°C). Finally, the glycerol content of the protein was adjusted to 50%. The protein was stored at -30°C until use.

#### *Sequence library preparation:*

Each batch of Tn5 transposase was serially diluted (50% glycerol, 50 mM Tris-HCl pH 7.5, 100 mM NaCl, 0.1 mM EDTA, 0.1% Triton X100, 1mM DTT), reacted and the fragmentation profiles analyzed (Agilent Bioanalyzer: HT DNA analysis kit) to determine the best dilution factor. Optimal DNA fragmentation reactions will yield a peak between 300 and 700 base pairs with minimal high molecular weight fragments.

Voltron plasmid libraries (96 well format) were diluted to 250 pg/ul and consolidated into 384 well plates. Unique 8mer barcodes were designed and inserted into ninety-six 'T7 well BC adapter' oligos (Supplementary Table 4) and 288 'i5 plate BC PCR' oligos. Free-end adapter pairs were annealed (100uM each oligo, reassociation buffer: 50mM NaCl, 10 mM TE pH 8.0, 1mM EDTA). Oligos pMENTS and i5_1 made double-stranded universal adapter #1. Oligo pMENTS and oligo set i7_1-96 made a 96 well plate of barcoded #2 adapters. Tn5 was diluted (dilution factor determined empirically for each batch) with reassociation buffer and was precharged with adapter #1 and each of the adapter #2's in a 96 well PCR plate at 37oC for 30 minutes (1uM of each adapter, 20% glycerol final conc.). The precharging reaction was scaled to tagment 24 x 96 well plasmid plates at a time. The 96 well plate of precharged Tn5 was diluted with 5 volumes of water and 2 volumes of TAPS-DMF buffer (50 mM TAPS-NaOH, 25 mM MgCl2, 50% v/v DMF pH 8.5) to activate the Tn5. Eight microliters of activated Tn5 set (96 wells) was robotically dispensed into each quadrant of six 384 well plates and 2 uL of plasmid DNA (0.5 ng) from each of six diluted 384 well plasmid plate was robotically added and mixed. Each plate was reacted at 55oC for 15 minutes and then immediately stopped by adding 2.5 uL 0.2% SDS/well and incubating at 70$^{o}$C for 10 minutes.

After tagmentation each 96 well quadrant (barcodes T7_1 to 96) of the 384 well reaction plate was robotically pooled into a single well. Each pool contained a mix of 96 tagmented plasmids each with a unique well barcode incorporated in its adapters. The tagmentation pools were collected in 96-well deep well plates and stored at -20$^{o}$C. Each 96 well plate of tagmentation pools was PCR amplified with universal i7_1 plate PCR primer and a barcoded “i5 plate BC PCR” primer for each well (12 cycles; OneTaq Hot Start 2X; NEB). Every DNA fragment now has two barcodes associated (plate and well) that will map its sequencing reads back to original plasmid library plates. PCR products were purified and eluted (ZR-96; Zymo Research). Random samples were analyzed (Agilent High Sensitivity DNA Kit) to ensure optimal fragmentation for each tagmentation run. All purified PCR products were quantified using Qubit double-stranded assay kit (Invitrogen). PCR samples with yields above 1 ng/ul were pooled (4ng each) into a library for NextSeq sequencing.

#### *Deep Sequencing*

Library concentration was determined by qPCR (Kapa Biosystems) using library size obtained by Agilent Bioanalyzer. Libraries sequenced on a NextSeq 550 using custom sequencing primers shown in Supplementary Table 4, with 150 bp in read 1 and read 2 and 8 bp for each library barcode.

#### *Analysis*

We chose to use a two-track approach to detect variants in the samples: an alignment-based method to map the reads to the expected reference, and a de novo assembly method to determine the sequence of Voltron2 independent of the plasmid reference.

Samples were analyzed using custom shell and Python scripts. Paired reads were trimmed with cutadapt v1.18 (Martin 2011) with the following parameters: --trim-n -n 4 -a CTGTCTCTTATACACATCT -a GAGGCGGAGACGGTG -a GGACAGGGACAGCCGAGCCCACGAGAC -a ATCTCGTATGCCGTCTTCTGCTTG -a AGATGTGTATAAGAGACAG -m36 -A CTGTCTCTTATACACATCT -A GCCGTCCTCGATCGCAGGCT -A ATCGCAGGCTATAGCGTGGAGACGCTGCCGACGA -A GTGTAGATCTCGGTGGTCGC -A AGATGTGTATAAGAGACAG.

For the alignment approach, trimmed reads mapped to the reference plasmid Voltron_421dot1.fasta using bwa mem v0.7.17-r1188 (Li 2013) with -a and -M flags. Secondary alignments were removed using samtools v1.10 (Li et al. 2009) and per-base coverage across the Voltron2 CDS was calculated using bedtools genomecov (Quinlan and Hall 2010). Duplicate alignments were then removed with samtools rmdup and mpileup was calculated using the following parameters: -C50 -Q20 -q10 -l Voltron_421dot1.fasta.cds.bed -Bf Voltron_421dot1.fasta. The consensus sequence for the Voltron2 CDS was extracted with bcftools.

For the de novo assembly approach, trimmed reads were downsampled using reservoir sampling without replacement to approximately 100x coverage and were interleaved for assembly. Assembly was performed using SPAdes v3.9.0 (Bankevich et al. 2012) with the following parameters: --careful -t 1 -m 6.

A custom python script was used to translate the CDS sequences determined by alignment and assembly into amino acid sequences for variant calling. For the alignment approach, the start of the CDS is known based on the plasmid reference. For the assembly approach, the start of the CDS was discovered in the contigs product by SPAdes by searching for either a 27 base sequence upstream of the CDS or a 27 base sequence at the start of the CDS (which should be unaffected by the mutagenesis). Variants at each amino acid position were called by comparing the translated sequences from alignment and assembly approaches to the Voltron2 translation, and only samples where the set of variants determined by each approach were identical were considered to be a consensus and reported in Supplementary Tables 1,2.

Once the analysis is complete for all samples, the results can be merged into a combined_summary.txt file using the example Linux code below:

```
ls -1 *_1.fastq.gz | sed 's/_1\.fastq\.gz//' | sed 's/Sample_//' >
all_samples
cd alignments/
ln -s */*.cov .
ls -1 *.cov > coverage
combine_genie_wiggle_files.py -l coverage
for i in *.cov; do unlink $i; done
ln -s ../all_samples .
echo "sample plate well starting_reads primary_aligns unmapped
mapped_ratio PCR_duprate uniq_mapped uniq_unmap uniq_ratio asmbl_reads
Total_records Total_bases Total_Ns Total_not-N Longest_record
Second_longest Shortest_record Median_length N50 map2ref_var denovo_var
common_var dn_CDS_rec dn_CDS_start map2ref_len denovo_len map2ref_seq
map2ref_qual denovo_seq" | tr " " "\t" > combined_summary.txt
for i in $(cat all_samples | sed 's/trim\.//'); do echo -n $i" " | tr " "
"\t" >> combined_summary.txt;
echo $i | cut -d"_" -f1 | tr "\n" "\t" >> combined_summary.txt
echo $i | cut -d"_" -f3 | tr "\n" "\t" >> combined_summary.txt
grep -A 9 "Begin map2ref" ../logs/${i}.out | cut -f2 | grep -v "Begin" |
tr "\n" "\t" >> combined_summary.txt
```

```
(test -r ${i}/scaffolds.fasta.summary.txt && grep -A 9 "Begin assembly
reporting" ../logs/${i}.out | cut -f2 | grep -v "Begin" | tr "\n" "\t" >>
combined_summary.txt) || echo -n "          " | tr " " "\t"  >>
combined_summary.txt
(test -s ${i}/${i}.map2ref.fq.txt && paste ${i}/${i}.map2ref.fq.txt >>
combined_summary.txt) || echo "           " | tr " " "\t" >>
combined_summary.txt
done
```
